# Recombinant adenovirus causes prolonged mobilization of macrophages in the anterior chamber of mice

**DOI:** 10.1101/2021.01.13.426423

**Authors:** Kacie J. Meyer, Danielle Pellack, Adam Hedberg-Buenz, Nicholas Pomernackas, Dana Soukup, Kai Wang, John H. Fingert, Michael G. Anderson

**Author notes:** Correspondence to: Michael G. Anderson, Department of Molecular Physiology and Biophysics, 3123 Medical Education and Research Facility, 375 Newton Road, Iowa City, IA 52242.

## Abstract

**Purpose:** Ocular tissues of mice have been studied in many ways using replication deficient species C type 5 adenoviruses (Ad5) as tools for manipulating gene expression. While refinements to injection protocols and tropism have led to several advances in targeting cells of interest, there remains a relative lack of information concerning how Ad5 may influence other ocular cell types capable of confounding experimental interpretation. Here, a slit-lamp is used to thoroughly photodocument the sequelae of intraocular Ad5 injections over time in mice, with attention to potentially confounding indices of inflammation.

**Methods:** A cohort of C57BL/6J mice was randomly split into 3 groups (Virus, receiving unilateral intracameral injection with 5×10^7^ pfu of a cargo-less Ad5 construct; Saline, receiving unilateral balanced salt solution injection; and Naïve, receiving no injections). From this initial experiment, a total of 52 eyes from 26 mice were photodocumented via slit-lamp at four time points (baseline, 1, 3, and 10 weeks following initiation of the experiment) by an observer masked to treatments and other parameters of the experimental design. Following the last in vivo exam, tissues were collected. Based on the slit-lamp data, tissues were studied via immunostaining with the macrophage marker F4/80. Subsequently, three iterations of the original experiment were performed with otherwise identical experimental parameters testing the effect of age, intravitreal injection, and A195 buffer, adding slit-lamp photodocumentation of an additional 32 eyes from 16 mice.

**Results:** The masked investigator was able to use the sequential images from each mouse in the initial experiment to assign each mouse into its correct treatment group with near perfect fidelity. Virus injected eyes were characterized by corneal damage indicative of intraocular injection and a prolonged mobilization of clump cells on the surface of the iris. Saline injected eyes had only transient corneal opacities indicative of intraocular injections, and Naïve eyes remained normal. Immunostaining with F4/80 was consistent with ascribing the clump cells visualized via slit-lamp imaging as a type of macrophage. Experimental iterations using Ad5 indicate that all virus injected eyes had the distinguishing feature of a prolonged presence of clump cells on the surface of the iris regardless of injection site. Mice receiving an intraocular injection of Ad5 at an advanced age displayed a protracted course of corneal cloudiness that prevented detailed visualization of the iris at the last time point.

**Conclusions:** Because the eye is often considered an “immune privileged site”, we suspect that several studies have neglected to consider that the presence of Ad5 in the eye might evoke strong reactions from the innate immune system. Ad5 injection caused a sustained mobilization of clump cells, i.e. macrophages. This change is likely a consequence of either direct macrophage transduction or a secondary response to cytokines produced locally by other transduced cells. Regardless of how these cells were altered, the important implication is that the adenovirus led to long lasting changes in the environment of the anterior chamber. Thus, these findings describe a caveat of Ad5-mediated studies involving macrophage mobilization, which we encourage groups to use as a bioassay in their experiments and consider in interpretation of their ongoing experiments using adenoviruses.

## INTRODUCTION

Recombinant adenoviruses have many advantages—and some notable disadvantages—for application as gene transfer vehicles [1-4]. One reason that adenoviruses were initially developed for gene transfer is that a great deal of their basic biology has long been well understood [5]. All adenoviruses are non-enveloped double stranded DNA viruses with a linear double-stranded genome encased along with core proteins into an icosahedral capsid [6]. Adenoviruses can be classified into seven species (A-G), with multiple serotypes per sub-group [7]. In humans, adenovirus infection is typically mild, with the notable exception of immunocompromised patients, for which it can be life-threatening. To promote safety of recombinant adenoviruses used in laboratories, the E1 region of the adenoviral genome is typically deleted, rendering the virus replication incompetent and creating a location for insertion of transgene cassettes [8]. Recombinant adenoviruses typically have broad tropism, high efficiency of gene delivery, and can transduce both dividing and quiescent cell populations. Upon entry into the nucleus, adenovirus can initiate gene transcription without integrating into the host genome, circumventing problematic insertional mutagenesis. Replication deficient species C type 5 (Ad5) was among the first vectors of this type studied, and following the refinement of protocols for its efficient production [9, 10], has grown in popularity to become one of the most popular gene transfer tools used in research.

Ocular tissues of mice have been studied in many ways with Ad5 [11, 12]. Most studies using Ad5 have desired transduction of two ocular tissues, retinal photoreceptors in the posterior segment and trabecular meshwork cells in the anterior segment. In both cases, effective transfer to these cell-types is challenged by the high transduction efficiencies of neighboring tissues. For the retina, retinal pigmented epithelium and Muller cells tend to be more readily transduced than photoreceptors [11], and in the case of the anterior chamber, corneal endothelium tends to be transduced more than trabecular meshwork cells [13]. Transduction with replication-deficient adenoviruses is transient, with Ad5-driven reporter expression typically described as lasting a period of 2–7 weeks following intraocular injections [13-15]—which can be extended with use of anti-inflammatory treatments [14]. Refinements to injection protocols [16] and tropism [17, 18] continue to improve apparent outcomes, though with more success for transfer to trabecular meshwork cells than for photoreceptors. While the efficiency of transfer to desired cell-types is important, it is equally important to consider how to prevent Ad5 from influencing unwanted cell-types, especially cells of the immune system. While some progress in averting adenoviral immune responses has been made in other tissues [1], less has been studied or attempted in the eye [14].

Here, we used slit-lamp imaging to describe the consequences of intraocular Ad5 injection in healthy C57BL/6J mice that were photodocumented at 4 time points (baseline, 1, 3, and 10 weeks following initiation of the experiment). We were led to conduct this study in a comprehensive fashion following sporadic observations from pilot experiments indicating that Ad5 injected eyes had adverse reactions in the anterior chamber that were more common and severe than suggested by the existing literature—which has largely been based on histologic sampling. Our slit-lamp data indicate a highly predictable response involving corneal opacity, which resolves, and a prolonged mobilization of clump cells on the surface of the iris, which did not resolve up to the oldest time points examined. These long-lived changes to macrophages of the anterior chamber could have a confounding influence in studies using adenoviral vectors and we suggest that future experiments should include screening for them as part of their experimental design.

## METHODS

### Experimental animals

All experiments were performed at the University of Iowa, conducted in accordance with the Association for Research in Vision and Ophthalmology Statement for the Use of Animals in Ophthalmic and Vision Research, and approved by the Institutional Animal Care Use and Committee of the University of Iowa. C57BL/6J mice were obtained from The Jackson Laboratory (Stock 000664; Bar Harbor, ME, USA) and subsequently bred and housed at the University of Iowa Research Animal Facility.

### Slit-lamp examination

Slit-lamp examination and photo documentation were performed by MGA (masked to treatment for the entire course of the study) at baseline (3 days prior to when some eyes received injections), and 1, 3, and 10 weeks following initiation of the experiment. Anterior chamber phenotypes were assessed in conscious mice, using a slit-lamp at 25X and 40X magnifications (SL-D7; Topcon, Tokyo, Japan), and photodocumented using a digital camera (D800; Nikon, Tokyo, Japan). All photographs were taken with identical slit-lamp settings and documented using identical camera settings and image processing. Following the final exam, eyes were grouped by MGA (still masked to treatment) based on common ocular phenotypes.

### Adenovirus injection

A cargo-less Ad5 stock construct was purchased from the University of Iowa Viral Vector Core (Ad5CMVempty; Catalog #: VVC-U of Iowa-272, Iowa City, IA, USA). The University of Iowa Viral Vector Core purifies adenoviral vectors by standard ultracentrifugation using a double cesium chloride step gradient and dialyzes extensively against A195 formulation buffer [19].

Mice with normal slit-lamp examinations at baseline were randomly divided into sex-matched groups: 1) Virus mice (N=11) had one eye injected intracamerally with 5×10^7^ pfu of Ad5CMVempty and one eye remaining naïve; 2) Saline mice (N=11) had one eye injected intracamerally with balanced salt solution (BSS; Alcon Laboratories, Fort Worth, TX, USA) and one eye remaining naïve; and 3) Naïve mice (N=5) did not receive an injection in either eye but were otherwise treated identically to the other groups. Treatments were randomized between right and left eyes. One mouse from the Saline cohort died during the study. To test iterations of the original experiment, a second cohort of mice with normal slit-lamp examinations at baseline were randomly divided into sex-matched groups: 1) Virus mice (N=4) had one eye injected intracamerally with 5×10^7^ pfu of Ad5CMVempty and one eye remaining naïve, identical to “Virus mice” in the original experiment; 2) Buffer mice (N=4) had one eye injected intracamerally with A195 Buffer provided by the University of Iowa Viral Vector Core and one eye remaining naïve; 3) Intravitreal mice (N=4) had one eye injected intravitreally with 5×10^7^ pfu of Ad5CMVempty and one eye remaining naïve; and 4) Aged mice (N=4) had one eye injected with 5×10^7^ pfu of Ad5CMVempty and one eye remaining naïve; three eyes in this group received intracameral injections and one eye received an intravitreal injection.

For injections, mice were anesthetized with a mixture of 87.5 mg/kg ketamine (VetaKet®, AKORN, Lake Forest, IL, USA) and 12.5 mg/kg xylazine (Anased, Lloyd Laboratories®, Shenandoah, IA, USA). All eyes were dilated with 2% cyclopentolate eye drops (Alcon Laboratories, Inc., Fort Worth, TX, USA). Upon full anesthesia, 0.5% proparacaine eye drops (Bausch & Lomb, Rochester, NY, USA) were applied to all eyes to provide additional local anesthesia. For each eye designated to receive an intracameral injection, the cornea was first punctured using a 33G needle and the aqueous humor allowed to drain from the anterior chamber. Treatment was delivered as a 2.0 µl intracameral injection using a 30G needle with reentry via the initial puncture site. For each eye designated to receive an intravitreal injection, the eye was punctured slightly posterior to the limbus using a 33G needle. Treatment was delivered as a 2.0 µl intravitreal injection using a 30G needle with reentry via the initial puncture site. For recovery, all mice received artificial tear ointment in both eyes (AKORN Animal Health, Inc., Lake Forest, IL, USA), an injection of 1 mg/kg antisedan (Zoetis, Inc.; Kalamazoo, MI, USA), and exogenous warmth.

### Histochemistry

Mice were euthanized by carbon dioxide inhalation followed by cervical spine dislocation. Eyes were collected, fixed in 4% paraformaldehyde in 1X phosphate buffered saline (PBS) for 4 h at 4°C with agitation. Anterior cups were rinsed in PBS, cryoprotected in increasing concentrations of sucrose solutions in PBS (5 to 30%), embedded in a mixture of two parts 30% sucrose and one part Optimal Cutting Temperature compound (Tissue-Tek®; Sakura Finetek USA, Inc.; Torrance, CA, USA), and cut in 7 µm sagittal cryosections using a cryostat. Cryosections underwent either histochemical or immunohistochemical staining. For the histochemical staining, slides were stained with hematoxylin and eosin using standard methods and imaged by light microscopy (BX52 equipped with a DP72 camera; Olympus) using identical settings. For immunohistochemical staining, cryosections were blocked in 1% bovine serum albumin in PBS, incubated with primary antibody (monoclonal rat anti-F4/80, 1:200 dilution, MCA497; BIO-RAD; Hercules, CA, USA) for one hour, rinsed, and incubated with secondary antibody (conjugated goat anti-rat Alexa fluoro 546, 1:1,000 dilution, A11081; Invitrogen by ThermoFisher Scientific; Waltham, MA, USA) for 30 minutes, all of which were done at room temperature. Both antibodies were diluted in antibody diluent (IHC-Tek™; IHC WORLD, LLC; Ellicott City, MD, USA). Samples were rinsed, counterstained with DAPI, and imaged at 200X magnification by confocal microscopy (DM2500 SPE; Leica Microsystems, Inc.; Buffalo Grove, IL, USA) and dark field light microscopy using identical settings.

## RESULTS

### Cohort generation

To study the consequences of adenovirus following ocular injection, we used a slit-lamp to document the anterior chamber over time following Ad5 injection into the anterior chamber. To collect baseline data, we first imaged 29 naïve C57BL/6J mice (male N=11; female N=18) at 12.5 weeks of age. Consistent with previous reports on sporadic ocular abnormalities in C57BL mouse strains [20], 2 of 58 eyes were found to be microphthalmic, leading to 2 mice being excluded from further study. At 13 weeks of age, remaining mice (male N=11; female N=16) were randomly assigned to one of three treatment groups: Virus, Saline, or Naïve. Mice in the Virus group had one eye injected with Ad5CMVempty and the other eye remained naïve. Mice in the Saline group served as a control for ocular injection of a fluid, with one eye injected with BSS and the other eye remaining naïve. Both eyes of mice in the Naïve group remained naïve; they otherwise received identical treatment to the experimental animals. In mice receiving injections, treatment was randomized between left and right eyes.

Our initial study design was to have at least 10 mice in the Virus and Saline groups, and 5 mice in the Naïve group; to exclude any mice with obvious initial complications from the injection; to expect some attrition during aging; and to not exclude any mice meeting criteria, which might result in cohorts with greater than 10 mice. There were no obvious complications with injections and one mouse in the Saline group died early in the course of the study. Thus, the final number of mice in each group available for the masked study were: Virus (N=11), Saline (N=10), and Naïve N=5).

### Slit-lamp examinations

A masked investigator examined and photodocumented all eyes (N=52) at the 1-, 3-, and 10-week time points. The entirety of this data set (208 images at 25X and 52 images at 40X) is available in Appendix 1–8. Following completion, the masked investigator used the four sequential image pairs from each mouse to assign mice into similar groupings based on common ocular phenotypes. Three mice were not able to be categorized because each had one eye that appeared to have been manipulated but exhibited severe corneal opacity that prevented imaging of any other anterior chamber structures. In these eyes, which were opaque throughout the entirety of the study, it was not possible to discern whether they were mice that had inadvertent damage from the injection procedures or mice that had severe reactions to the materials injected (Appendix 1-2). The remaining mice could be grouped into one of three groups with common presentations.

Group 1 (N=8) was characterized by one normal appearing eye, and one eye indicative of intraocular injection and inflammation (Figure 1, Appendix 3-4). Eyes in this group had small corneal opacities at 1 week, which typically worsened (larger and more opaque) through 3 weeks and resolved to various degrees by 10 weeks. Affected eyes sometimes had small lenticular opacities. Most notably, and completely unique to this group, all the affected eyes in this group had irides with focally “rough” appearing surfaces. When the iris could be observed through the cloudy cornea, rough appearing irides were visible in some eyes at the 3-week time point and were present in all eyes at the 10-week time point. These changes were evident with the aid and magnification of a slit-lamp, but imperceptible to the naked eye. Upon closer examination, these areas were due to clusters of clump cells on the surface of the iris—cells identical to those we have previously observed in mice with iris diseases [21, 22], and resembling reports of similar macrophage-like cells of the iris that have been described by others [23-25]. Clump cells have a characteristic small round shape, which is best noted if observed via slit-lamp with an extreme angle with respect to the light source, in which their location on the surface of the iris is evident from the small crescent shadow they cast on the surface. Aged mice with normal eyes will sometimes have 1–2 spots where these cells are clearly present or suspected [22], but this will not give the iris a rough appearance. Thus, the rough appearing iris and striking accumulation of these clump cells was a clear indication of a shared aberration among affected eyes of this group.

**Figure 1.**
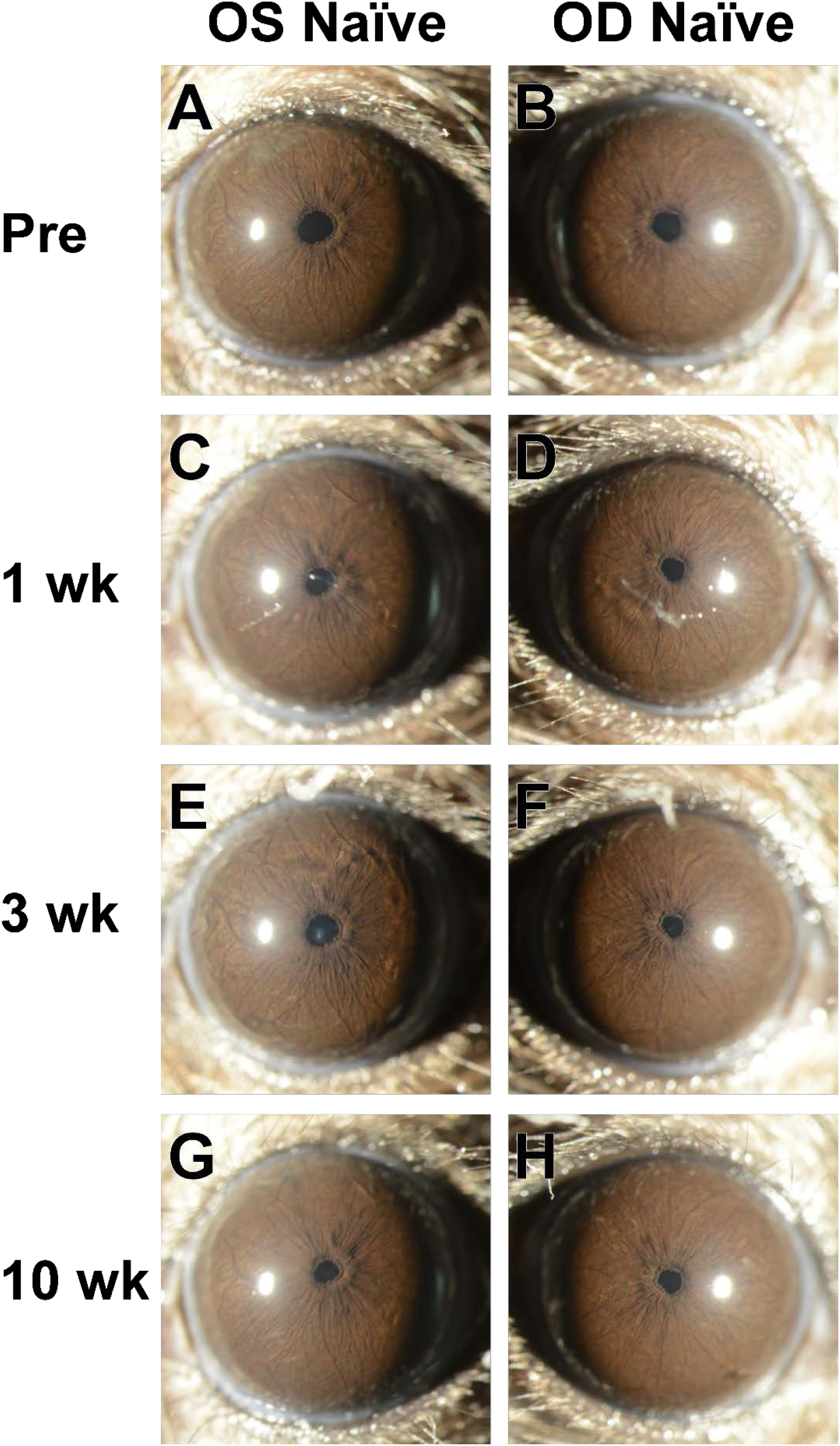
Slit-lamp images from a mouse in the Naïve group. Images from the left and right eyes of a single mouse at progressive ages showing normal appearance of the anterior chamber. **A-B**: Pre-treatment images were collected in mice that were 12.5 weeks old. The cornea is clear, and the iris vessels are the main notable feature of the iris. With subsequent aging in these unmanipulated mice, the same healthy appearance is maintained at **C-D**: 1 week, **E-F**: 3 weeks, and **G-H**: 10 weeks following initiation of the experiment. See Figure 4A for a different view of the same eye and time point shown in panel G. Images at 25X magnification were collected by an investigator who was masked to treatment status at the time they were photographed.

Among the other groups, Group 2 (N=9 mice) was characterized by one eye with a normal appearance and one eye with mild focal corneal cloudiness at the 1-week time point, which resolved at later, and sometimes with small focal lenticular opacities in the same eye, which did not resolve (Figure 2, Appendix 5-6). Group 3 (N*=*6 mice) was characterized by both eyes with a consistently normal appearance (Figure 3, Appendix 7-8).

**Figure 2.**
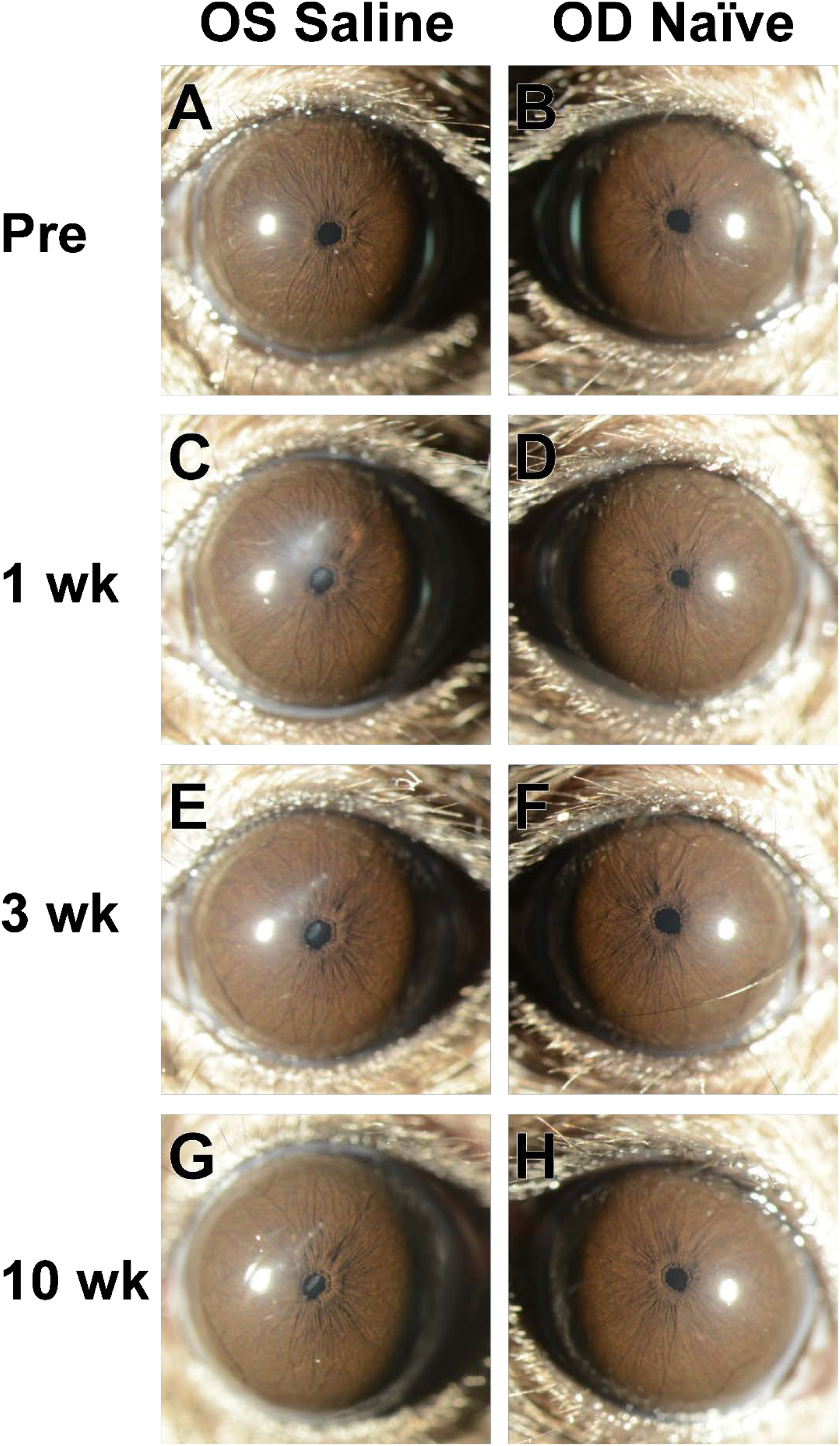
Slit-lamp images from a mouse in the Saline group. Images from the left and right eyes of a single mouse at progressive ages showing consequences of intraocular injection in the left eye and normal appearance of the anterior chamber in the right eye. **A-B**: Pre-treatment images were collected in mice that were 12.5 weeks old. The left eye was subsequently injected with BSS. **C-D**: At the 1-week time point, the injected left eye has a mild corneal opacity where the needle had been inserted and a small lenticular opacity; the naïve right eye has a normal appearance. **E-F**: At the 3-week and **G-H**: 10-week time points, the corneal opacity in the injected left eye became progressively less severe and the lenticular opacity appeared to be unchanged; meanwhile, the naïve right eye maintained a normal appearance. See Figure 4C for a different view of the same eye and time point shown in panel G. Images at 25X magnification were collected by an investigator who was masked to treatment status at the time they were photographed.

**Figure 3.**
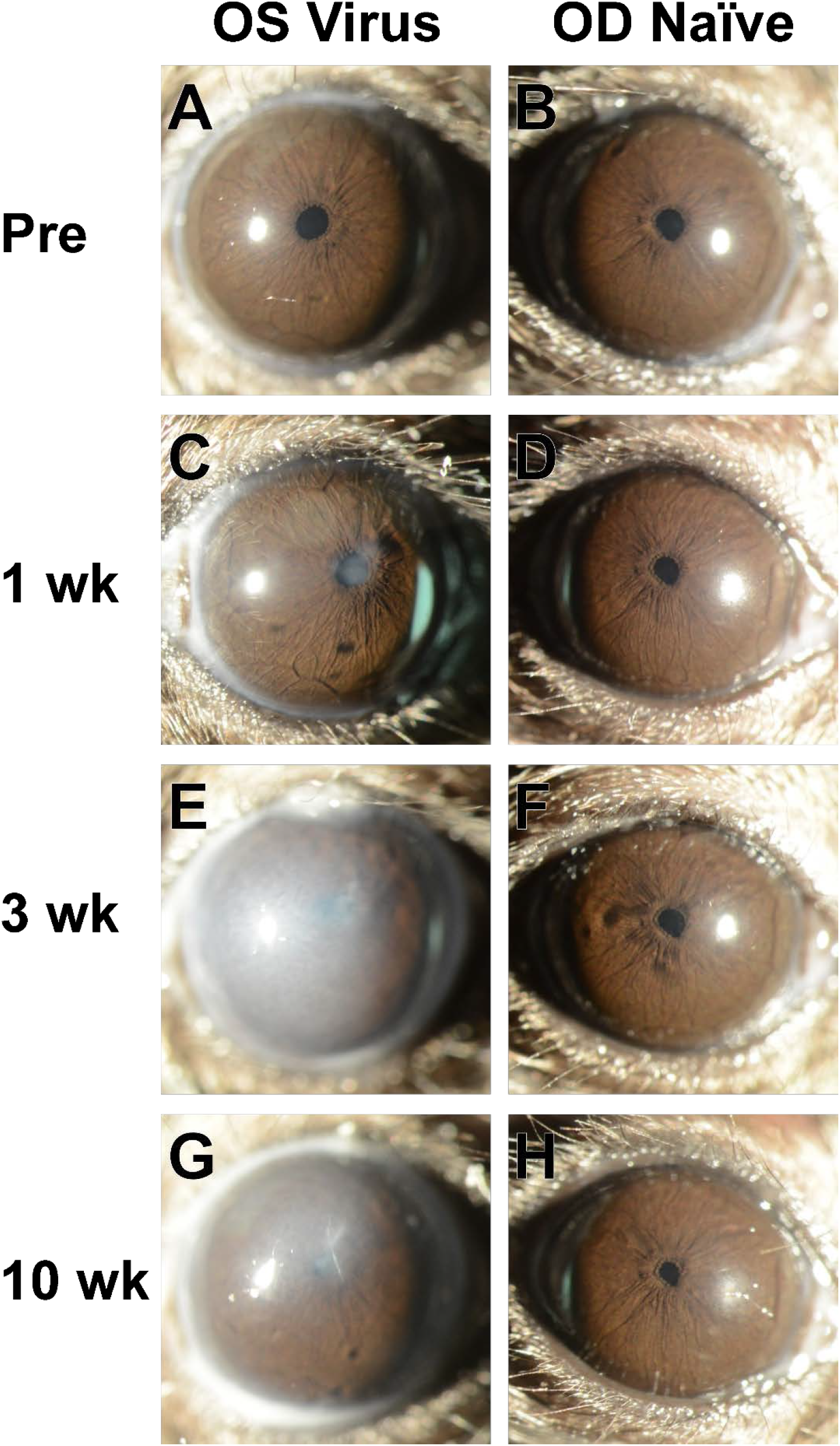
Slit-lamp images from a mouse in the Virus group. Images from the left and right eyes of a single mouse at progressive ages showing consequences of intraocular Ad5 injection in the left eye and normal appearance of the anterior chamber in the right eye. **A-B**: Pre-treatment images were collected in mice that were 12.5 weeks old. The left eye subsequently received an intraocular injection of Ad5. **C-D**: At the 1-week time point, the injected left eye has several areas showing mild corneal opacity and lenticular opacity; the naïve right eye has a normal appearance. **E-F**: At the 3-week time point, the corneal opacity of the injected left eye had become significantly more severe, blocking visualization of the remainder of the anterior chamber; the uninjected right eye remained normal in appearance. **G-H**: At the 10-week time point, the corneal opacity in the injected left eye lessened in severity and where areas of the iris can be viewed, rough appearing areas are present. The naïve right eye maintained a normal appearance Note that the apparent cloudiness of the cornea is also dependent on the reflectivity of the light source, see Figure 4E for a different view of the same eye and time point shown in panel G. Images at 25X magnification were collected by an investigator who was masked to treatment status at the time they were photographed.

At completion, the masked observer (who was masked to treatment and the precise number of mice per group) assigned Group 1 to Virus, Group 2 to Saline, and Group 3 to Naïve. After unmasking, all 8 eyes predicted to be in the Virus group were accurately assigned. There was 1 mismatch among the controls, in which a mouse in the Saline group was inaccurately predicted to be in the Naïve group. There is a significant association between the predicted treatment groups and the true treatment groups (P=4.08E-9; True Group N=8:10:5 [Virus:Saline:Naïve] vs. Predicted Group N=8:9:6 [Virus:Saline:Naïve]; Fisher’s Exact Test). All 3 of the non-categorized mice with severe corneal opacity were in the Virus group.

### Appearance and immunostaining of clump cells

The most uniquely distinguishing feature between groups was the prolonged presence of clump cells on the surface of the iris (Figure 4). Clump cells are typically considered to be a type of macrophage [23, 24]. To confirm that the cells we have observed responding to Ad5 share features of macrophages, we performed immunostaining with the macrophage marker F4/80 [26, 27] (Figure 5) and H&E histochemical staining (Supplementary Figure 1). Small round F4/80-positive cells on top of the iris stroma, the same location as we observed the clump cells by slit-lamp, were uniquely visible in most, but not all, sections of eyes from the Virus group. F4/80-positive cells were additionally observed within the ciliary body and deep within the iridocorneal angle of eyes from the Virus group.

**Figure 4.**
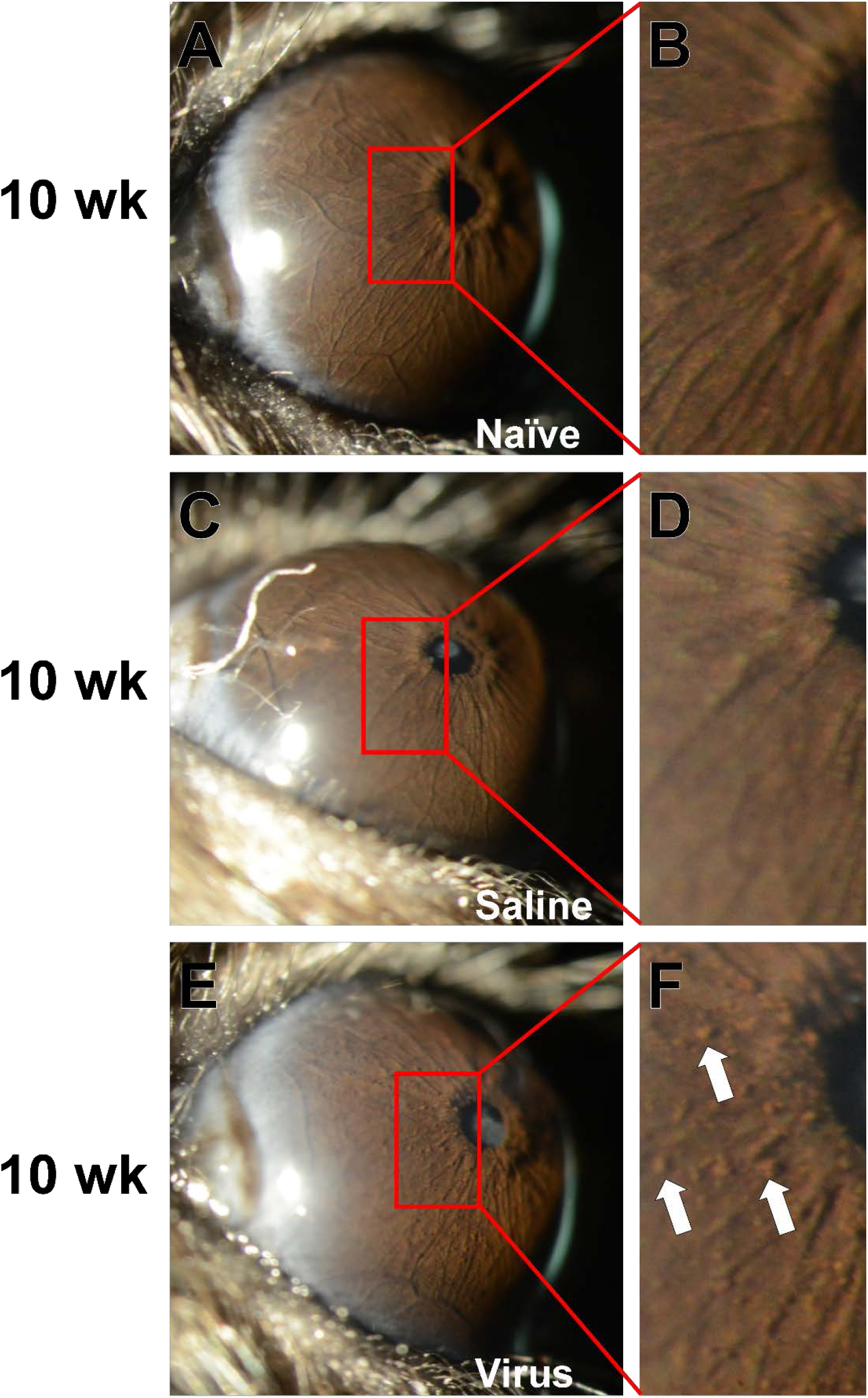
High-magnification slit-lamp images showing the unique presence of clump cells in Ad5 injected eyes. Images are from the same eyes shown at the 10-week time point in Figures 1-3, but photographed at a higher magnification and with the mouse held at a more severe angle with respect to the light source. A digital enlargement of the same areas immediately to the left of each pupil are shown in the right-hand column. **A-B**: The iris of mice in the Naïve group retained a normal morphology with no visible clump cells throughout the study. The same eye is also shown in Figure 1G. **C-D**: The iris of mice in the Saline group also maintained a normal morphology lacking visible clump cells. A lenticular opacity is also visible. The same eye is also shown in Figure 2G. **E-F**: Unique to only treated eyes of the Virus group, intraocular injection of Ad5 led to a notable accumulation of clump cells (*white arrow*, several additional also visible but unmarked) on the surface of the iris. Lenticular and corneal opacity is also apparent. The same eye is also shown in Figure 3G. Images at 40X magnification were collected by an investigator who was masked to treatment status at the time they were photographed.

**Figure 5.**
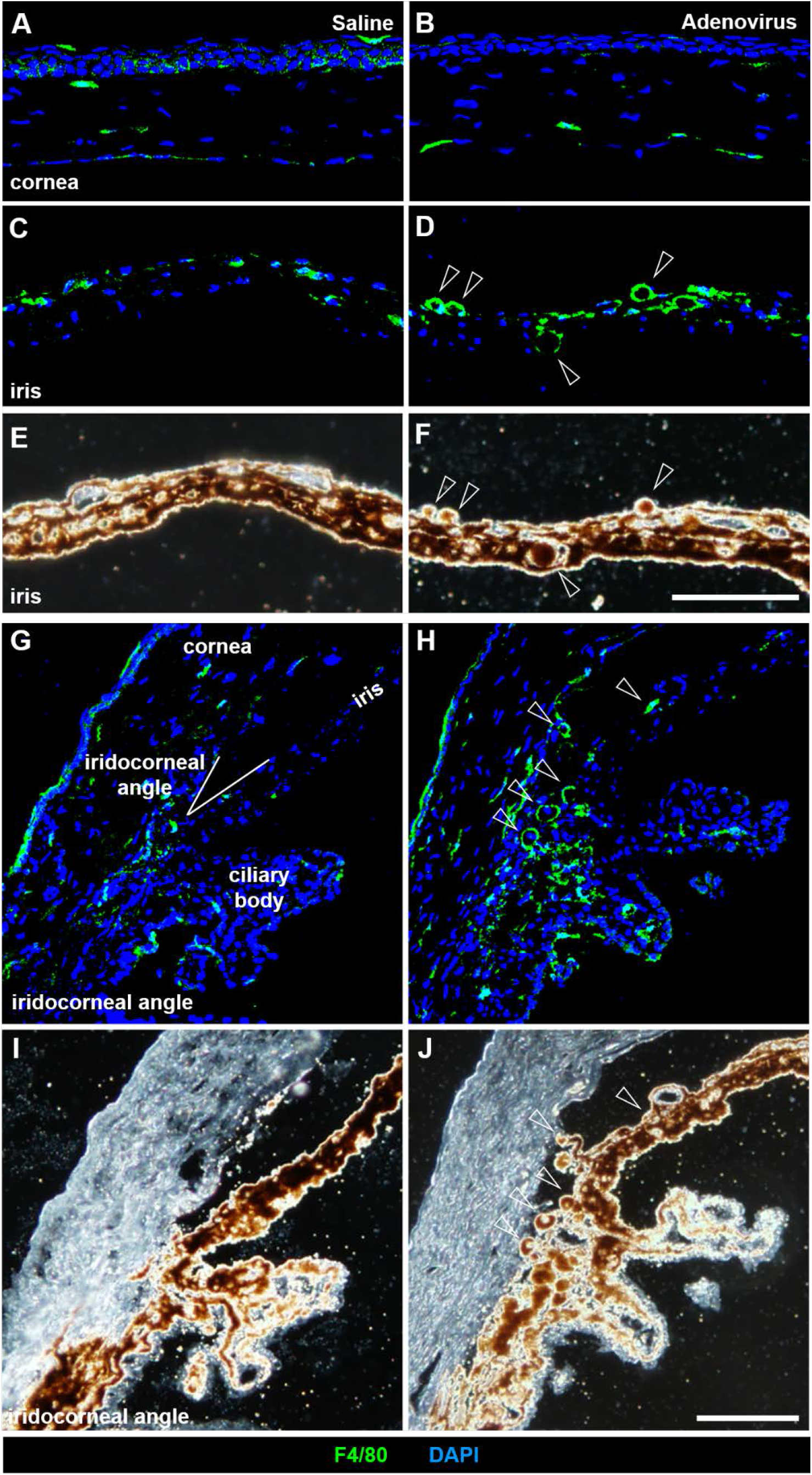
Localization of the macrophage marker F4/80 to clump cells in Ad5-injected eyes. Fluorescent and light micrographs of eyes immunostained with F4/80 and counterstained with DAPI (nuclear stain) in treated eyes of mice from the Saline cohort (left column) compared to those of mice from the Virus cohort (right column). **A-B**: The central cornea shows a similar localization and prevalence of F4/80^+^ cells. **C-F**: the mid-peripheral iris and **G-J**: iridocorneal angle show an increased prevalence of F4/80^+^ cells localized along the anterior iris stroma, matching the location of clump cells visualized via slit-lamp exam (*white arrowheads*). Note that F4/80^+^ cells also appear to be pigment laden, which is an additional feature ascribed to clump cells. F4/80^+^ cells are also prominent in the posterior iris pigmented epithelium and iridocorneal angle of Ad5-injected eyes. Notation of the prominent intraocular structures are indicated (*white text*) in panel G. Scale bar = 100 µm for (A-F) and (G-J).

### Testing experimental iterations

To study the effect of commonly used iterations of experiments involving Ad5 ocular injection, we used a slit-lamp to document the anterior chamber over time of a second cohort of C57BL/6J mice. To collect baseline data, we first imaged 12 naïve C57BL/6J mice (male N=6; female N=6) at 12.5 weeks of age and 4 naïve C57BL/6J mice (male N=2; female N=2) at 23.5 weeks of age. No mice in this cohort were excluded from further study. At 13 weeks of age and 24 weeks of age, respectively, mice were randomly assigned to one of four treatment groups with treatment randomized between left and right eyes: Virus, Buffer, Buffer, or Aged. Mice in the Virus group had treatment identical to the “Virus” group in the original experiment, having one eye injected intracamerally with Ad5CMVempty and the other eye remaining naïve, and serve as a positive control. Mice in the Buffer group had one eye injected intracamerally with A195 Buffer, a common Adenovirus vehicle, and the other eye remained naïve. Mice in the Intravitreal group had one eye injected intravitreally with Ad5CMVempty and the other eye remained naïve. Last, mice in the Aged group were 24-weeks-old and had one eye injected with Ad5CMVempty and the other eye remained naïve; three eyes in this group received an intracameral injection and one received an intravitreal injection.

A masked investigator examined and photodocumented all eyes (N=32) at the 1-, 3-, and 10-week post-injection time points. The entirety of this data set (128 images at 25X and 32 images at 40X) is available in Appendix 9-16. Following completion, the masked investigator used the four sequential image pairs from each mouse to assign mice into four groupings based on common ocular phenotypes.

Group 1 (N=4) was characterized by one normal appearing eye, and one eye indicative of intracameral injection and inflammation (Figure 6, Appendix 9-10). Eyes in this group had small focal corneal opacities at 1 week, which typically worsened (larger and more opaque) through 3 weeks and resolved to various degrees by 10 weeks. Most notably, all the affected eyes in this group had irides with focally “rough” appearing surfaces. When the iris could be observed through the cloudy cornea, rough appearing irides were present in all eyes at the 10-week time point. Group 2 (N=4) was characterized by one normal appearing eye, and one eye indicative of intracameral injection (Figure 6, Appendix 11-12). Eyes in this group have a mild focal corneal cloudiness at the 1-week time point that resolved over time and have normal appearing irides. Group 3 (N=4) was characterized by one normal appearing eye, and one eye indicative of inflammation (Figure 6, Appendix 13-14). Eyes in this group have various degrees of diffuse corneal cloudiness. When the iris could be observed through the cloudy cornea, rough appearing irides were present in all eyes at the 10-week time point. Group 4 (N=4) was characterized by one normal appearing eye, and one eye indicative of intraocular injection and inflammation (Figure 6, Appendix 15-16). Eyes in this group were characterized by corneal cloudiness to an extent that visualization of the iris was not possible at the 10-week time point. Of note, all eyes in this group (including naïve eyes) are larger than the eyes in Groups 1-3.

**Figure 6.**
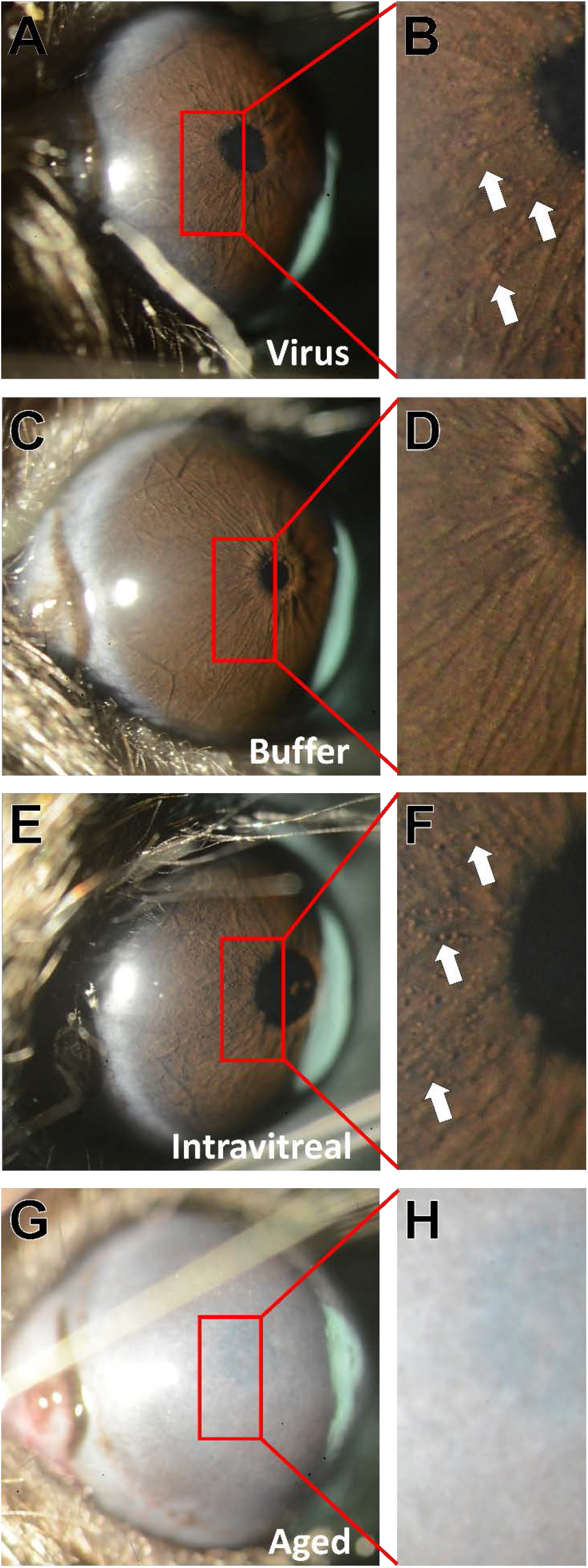
High-magnification slit-lamp images showing the results of experimental iterations for Ad5 intraocular injections. Images are from the same eyes shown at the 10-week time point in Appendices 9, 11, 13, and 15, but photographed at a higher magnification and with the mouse held at a more severe angle with respect to the light source. Images at 40X magnification were collected by an investigator who was masked to treatment status at the time they were photographed. A digital enlargement of the same areas immediately to the left of each pupil are shown in the right-hand column. **A-B**: In the Virus positive control group, anterior chamber injection of Ad5 led to a notable accumulation of clump cells (*white arrow*, several additional also visible but unmarked) on the surface of the iris, replicating the experiment shown in Figure 4E-F. **C-D**: The iris of mice in the Buffer group (anterior chamber injection of A195 buffer) maintained a normal morphology throughout the study lacking visible clump cells. **E-F**: In the Intravitreal group, an intravitreal injection of Ad5 led to a notable accumulation of clump cells (*white arrow*, several additional also visible but unmarked) on the surface of the iris. **G-H**: The eyes of the mice in the Aged group that received an intraocular injection of Ad5 all had corneal cloudiness at the 10-week time point that precluded visualization of the iris.

At completion, the masked observer (who was masked to treatment and the precise number of mice per group) assigned Group 1 to Virus (positive control), Group 2 to Buffer, Group 3 to Intravitreal, and Group 4 to Aged. After unmasking, all 32 eyes were assigned to the correct group.

## DISCUSSION

Although the ophthalmic slit-lamp has long been the primary instrument used in clinical ophthalmology, its use in research with mice remains uncommon. To our knowledge, we are the first to thoroughly characterize the sequelae of Ad5 injection in the anterior chamber of mice via slit-lamp photodocumentation. The results show that injection of the Ad5 vector itself—empty of any gene therapy inserts—results in changes to the anterior chamber involving transient corneal opacity and a persistent change in the localization of macrophages on the surface of the iris. The timeframe of these changes suggests that they are likely to be longer lived than viral gene expression itself. In describing the appearance and location of these Ad5-responsive macrophages, our results establish how slit-lamp exams can be used as a bioassay for inflammatory events in ongoing research using viral vectors.

The biological events allowing Ad5 to deliver gene therapy constructs to cells has been extensively studied. Capsid proteins typically mediate cellular internalization of Ad5 via attachment to the cell surface Coxsackie and adenovirus receptor (CAR; in mice encoded by the *Cxadr* gene) [28, 29], which is followed by interaction with integrins [30, 31] (Reviewed in [32-34]). CAR-negative cells are typically described as being poorly transduced by Ad5. However, several CAR-independent pathways mediating Ad5 transduction are also known. Ad5 can bind with multiple blood factors, such as blood coagulation factor X [35], which helps protect the adenovirus-complex from attack by the classical complement pathway [36], and facilitates transduction by bridging the virus to cell surface heparin sulfate proteoglycans. In cultured cells, this mechanism mediated by sulfated glycans has a prominent role in transduction of hepatocytes, but there are conflicting data regarding its importance to transduction of the mouse liver by Ad5 in vivo [37] and a species-dependent difference in the influence of human versus mouse blood coagulation factor X proteins in transduction that complicates the interpretation of some studies [38]. In macrophages, scavenger receptor A (in mice, encoded by the *Msr1* gene) is yet another Ad5 receptor [39]. The possibility of additional receptors awaiting characterization has been implicated by several studies [38, 40, 41]. Once bound to a receptor, Ad5 is internalized via clathrin-mediated endocytosis [42] or macropinocytosis [43]. After additional trafficking steps, the viral genome is ultimately inserted into the host nucleus [44]. The viral genome remains episomal and expresses any cargo gene transcripts until cell division has diluted the adenovirus to insignificant levels or the immune system has destroyed the transduced cells.

There is also a broad framework for predicting the concurrent immunologic events associated with the presence of recombinant Ad5 in a tissue. Adenoviruses elicit strong immune responses, even when they are initially injected into “immune privileged” sites in the eye. Viral antigens, as well as transgene products, are both capable of activating adaptive immune responses [45]. Consequently, intraocular injections of Ad5 are followed by intraocular presence of CD4^+^ and CD8^+^ T cells and generation of new circulating antibodies [35]. Hamilton and colleagues previously showed that repeated injections of Ad5 with an empty cassette did not interfere with subsequent expression of Ad5 with a luciferase reporter, suggesting that the immune privilege of the eye was likely protecting it from mounting an immune response against Ad5 [46]. However, since the time of Hamilton’s study it has become increasingly clear that innate immunity also needs to be considered. Adenoviruses elicit strong innate immune responses [47, 48]. In cells associated with innate immunity, such as macrophages and dendritic cells, Ad5 interacts with pathogen recognition receptors (PRRs) and stimulates release of numerous cytokines and chemokines [49]. There are many classes of PRRs, with toll-like receptors being one of the most studied [50, 51]. Many serotypes of adenovirus interact with PRRs and the biology of these interactions have been studied with particular intensity at the ocular surface where they are critically important to adenoviral keratoconjunctivitis [52]. Interestingly, innate responses to Ad5 may be quite broad and evoke pro-inflammatory responses from “non-immune” cell-types, including epithelial cells [53-55]. For example, cultured human conjunctival epithelial cells transduced with Ad5 have an upregulation of interleukin-6 (IL-6), interleukin-8 (IL-8), and intracellular adhesion molecule-1 (ICAM-1) [55].

In the specific context of intraocular Ad5 injections, relevant biological and immunologic consequences can be predicted, but much remains untested. The Ad5 receptor(s) for tissues transduced in the anterior chamber have not been defined. Because blood coagulation factor X seems to not be abundant in aqueous humor [56], the sulfated glycan mediated pathway for transduction seems unlikely for Ad5 injected directly into the anterior chamber. The sustained mobilization of clump cells in our experiment could represent either a consequence of direct macrophage transduction or a secondary response to cytokines produced locally by other transduced cells. Regardless of how these cells were altered, the important implication is that the adenovirus led to long lasting changes in the environment of the anterior chamber. Had the Ad5 we injected been carrying transgene cargo, it would not have been possible to discern whether phenotypic changes were specific to the cargo gene versus the cargo gene in the context of the now-altered environment of the anterior chamber. Could different injection techniques, concentrations, tropism, serotypes, or genetic backgrounds of mice have made a difference in our findings? Undoubtedly these are relevant variables, but our findings and the existing literature indicate that cautious interpretations are appropriate—many adenovirus transduced cells are likely to have an altered expression of cytokines induced by PRR signaling. We suspect that multiple studies have overlooked the clump cell phenotype because they have primarily used histology as an assay, which in any given section may show few, if any macrophages altered by Ad5, but that are readily seen with the broader view from slit-lamp examination. It is also relevant that small changes in cytokines in vivo may be undetectable by biochemical assays, but still physiologically important.

There are some caveats relevant to our study that should be considered. First, we did not directly measure any cytokines, chemokines, or PRR signaling in Ad5 affected eyes. We suggest their presence based on the literature and our hypothesis has not been tested. Second, we have reliably demonstrated that Ad5 injection caused a prolonged mobilization of clump cells with detailed photodocumentation of the iris that has been correlated with immunolabeling. However, much remains unknown about the nature of these cells. Visibly, the cells are identical in appearance and similar in apparent abundance to those we have previously studied in mice with various forms of iris disease [21, 22]. Several studies have shown that gene expression changes driven by intraocular injection of Ad5 typically last only 2–7 weeks [13-15]. Assuming a similar time course occurred in our experiments, the prolonged mobilization of clump cells at 10 weeks following injection predicts that the iris changes caused by Ad5 injection are likely longer lived than the Ad5 itself. Third, to avoid potential complications from anesthesia and/or rebound tonometry, our current experiments did not collect any IOP data which might be relevant considering the localization of F4/80-positive cells deep in the iridocorneal angle and proximity to the aqueous humor drainage structures.

There remains much promise for viral-mediated approaches in studying anterior chamber physiology using mice [57, 58]. Our current findings describe a caveat of Ad5-mediated studies involving macrophage mobilization, which we encourage groups to monitor and consider in their ongoing experiments using adenoviruses.

## Supporting information

Appendix 1 unassigned 25X

Appendix 2 unassigned 40X

Appendix 3 virus 25X

Appendix 4 virus 40X

Appendix 5 saline 25X

Appendix 6 saline 40X

Appendix 7 naive 25X

Appendix 8 naive 40X

Appendix 9 Virus 25X

Appendix 10 Virus 40X

Appendix 11 Buffer 25X

Appendix 12 Buffer 40X

Appendix 13 Intravitreal 25X

Appendix 14 Intravitreal 40X

Appendix 15 Aged 25X

Appendix 16 Aged 40X

## ACKNOWLEDGMENTS

This work was supported in part by Merit Review Award (I01 RX001481) from the U.S. Department of Veterans Affairs RR&D Service and an NIH/NEI Center Support Grant to the University of Iowa (P30 EY025580). AHB was supported by Training Grant Award (T32DK112751). The contents do not represent the views of the U.S. Department of Veterans Affairs or the U.S. Government.

---

**Supplementary Figure 1.**
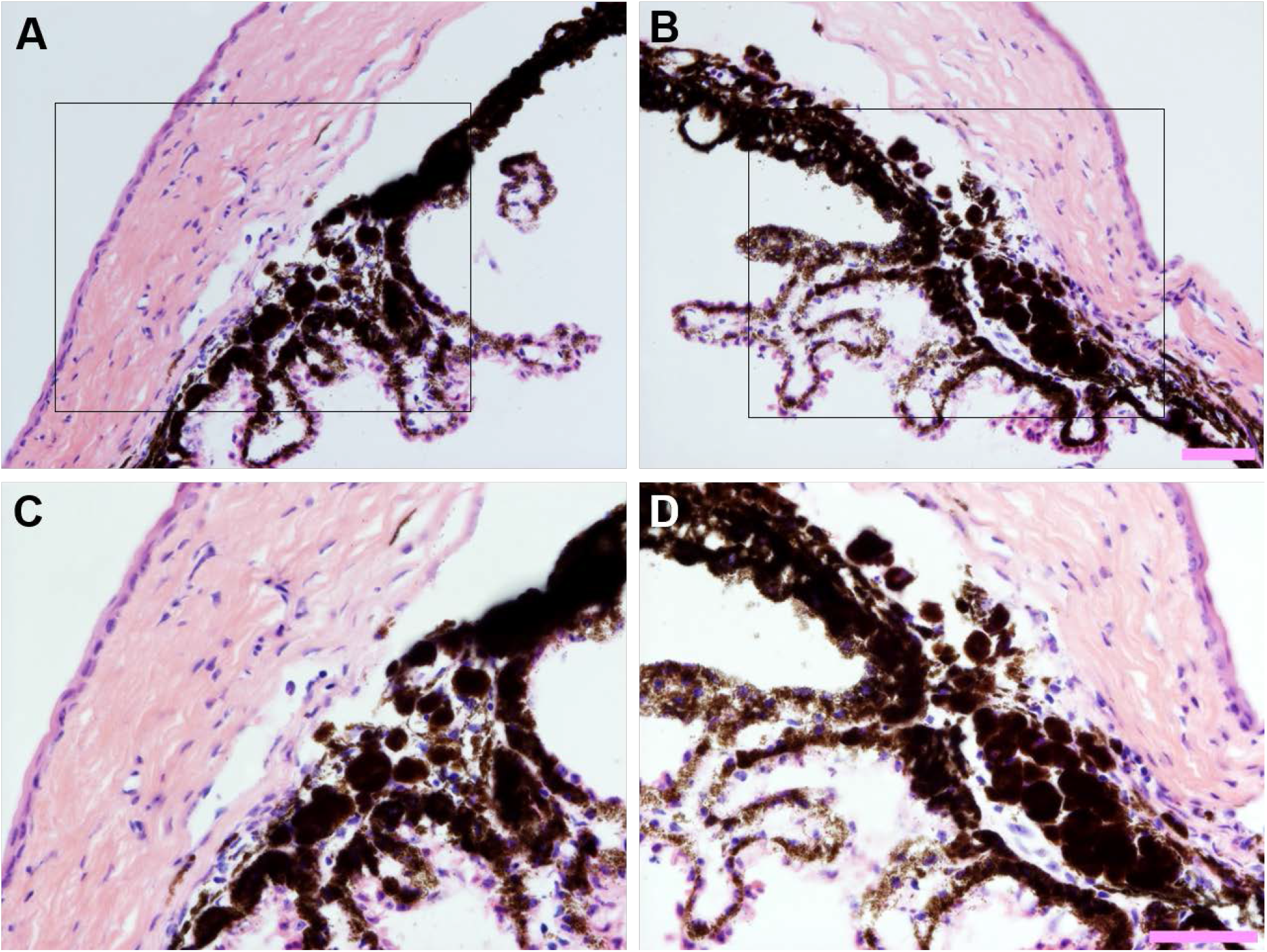
Persistence of clump cells in the iridocorneal angle of eyes following intraocular Ad5 injection. Light micrographs collected from opposite poles (*left vs. right column*) of hematoxylin and eosin-stained histological sections from the same eye shown at (A–B) 200X and (C–D) 400X total magnification. Areas within inset boxes (*top row*) are show at higher magnification below (*bottom row*). Note that despite some variability, there is a persistent localization of clump cells deep within the iridocorneal angle with proximity to the drainage structures. Scale bar = 50 µm.

## APPENDICES

**Appendix 1**. Complete dataset of 25X slit-lamp images from three mice at multiple ages which were not able to be categorized in the masked study because of severe corneal opacity.

**Appendix 2**. Complete dataset of 40X slit-lamp images from three mice at the 10-week time point which were not able to be categorized in the masked study because of severe corneal opacity.

**Appendix 3**. Complete dataset of 25X slit-lamp images from eight mice at multiple ages assigned to the Virus group in the masked study.

**Appendix 4**. Complete dataset of 40X slit-lamp images from eight mice at the 10-week time point assigned to the Virus group in the masked study.

**Appendix 5**. Complete dataset of 25X slit-lamp images from nine mice at multiple ages assigned to the Saline group in the masked study.

**Appendix 6**. Complete dataset of 40X slit-lamp images from nine mice at the 10-week time point assigned to the Saline group in the masked study.

**Appendix 7**. Complete dataset of 25X slit-lamp images from six mice at multiple ages assigned to the Naïve group in the masked study.

**Appendix 8**. Complete dataset of 40X slit-lamp images from six mice at the 10-week time point assigned to the Naïve group in the masked study.

**Appendix 9**. Complete dataset of 25X slit-lamp images from four mice at multiple ages assigned to the Virus positive group in the masked study of experimental iterations.

**Appendix 10**. Complete dataset of 40X slit-lamp images from four mice at the 10-week time point assigned to the Virus positive group in the masked study of experimental iterations.

**Appendix 11**. Complete dataset of 25X slit-lamp images from four mice at multiple ages assigned to the Buffer group in the masked study of experimental iterations.

**Appendix 12**. Complete dataset of 40X slit-lamp images from four mice at the 10-week time point assigned to the Buffer group in the masked study of experimental iterations.

**Appendix 13**. Complete dataset of 25X slit-lamp images from four mice at multiple ages assigned to the Intravitreal group in the masked study of experimental iterations.

**Appendix 14**. Complete dataset of 40X slit-lamp images from four mice at the 10-week time point assigned to the Intravitreal group in the masked study of experimental iterations.

**Appendix 15**. Complete dataset of 25X slit-lamp images from four mice at multiple ages assigned to the Aged group in the masked study of experimental iterations.

**Appendix 16**. Complete dataset of 40X slit-lamp images from four mice at the 10-week time point assigned to the Aged group in the masked study of experimental iterations.

