## Appendix 1 unassigned 25X for "Recombinant adenovirus causes prolonged mobilization of macrophages in the anterior chamber of mice"

Appendix 1
Mouse ID: 28453

Masked assignment: **Unassigned**
Actual group: **Virus** (OS injected)

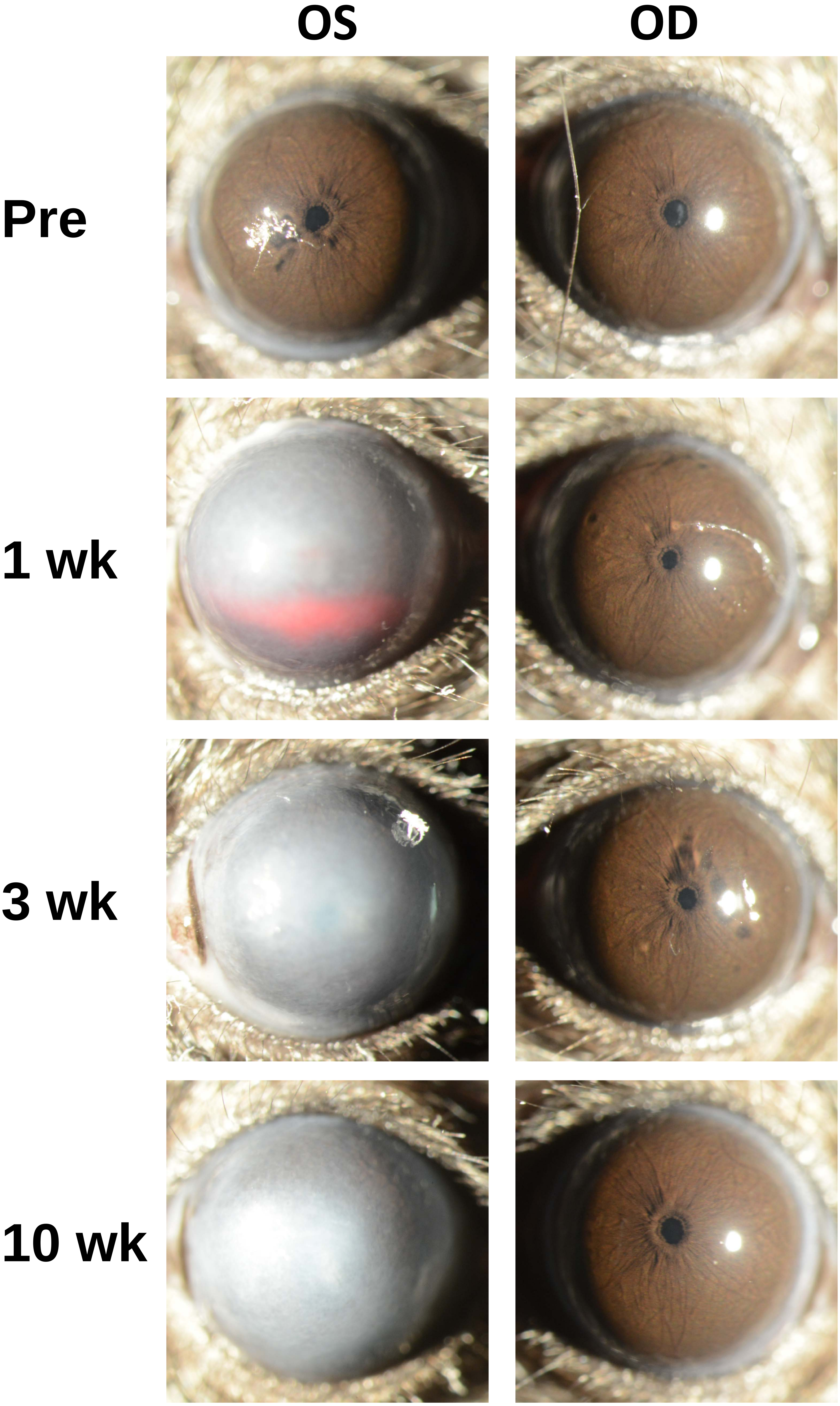

Appendix 1
Mouse ID: 28455

Masked assignment: **Unassigned**
Actual group: **Virus** (OD injected)

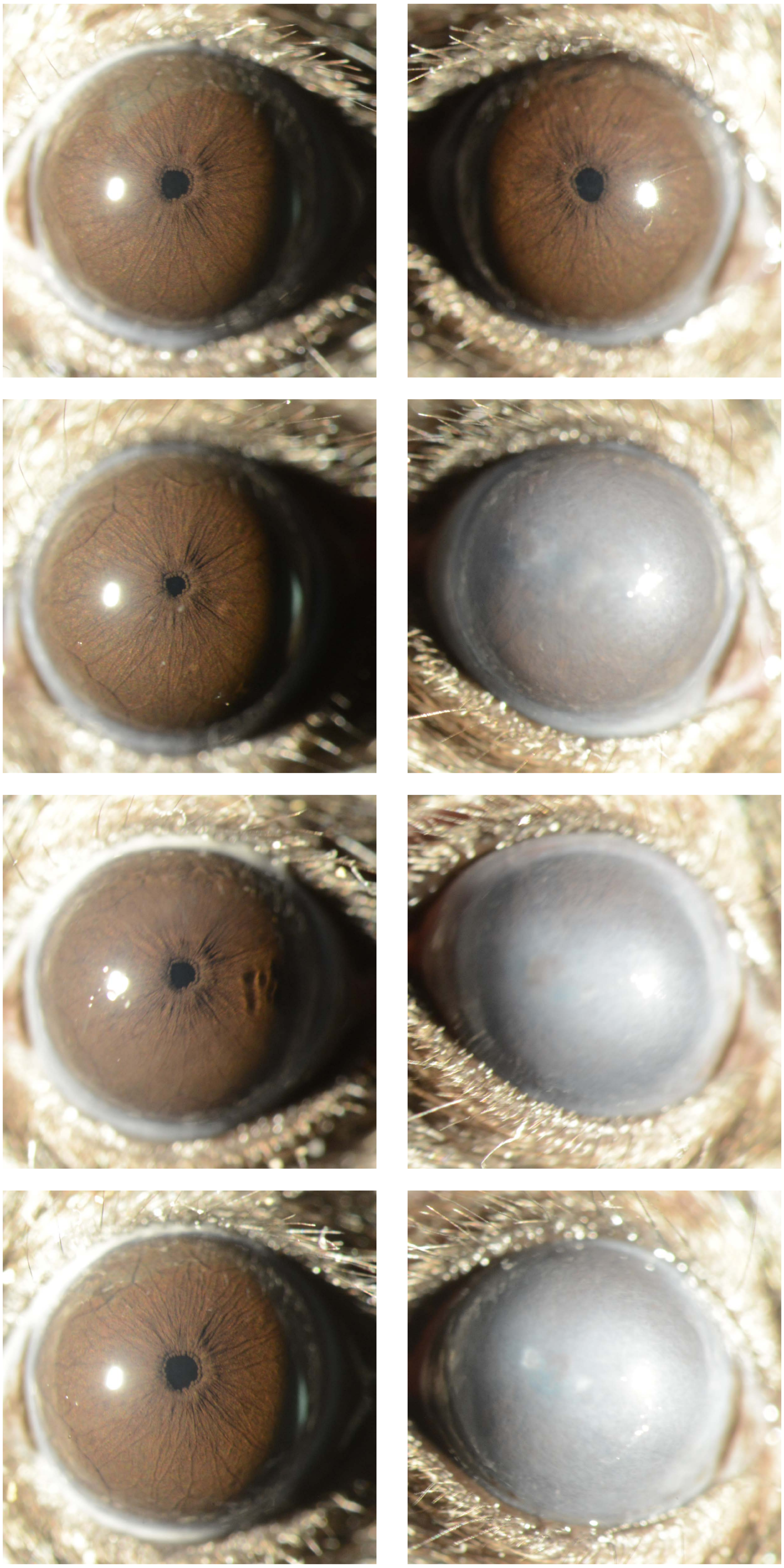

Appendix 1
Mouse ID: 28469

Masked assignment: **Unassigned**
Actual group: **Virus** (OD injected)

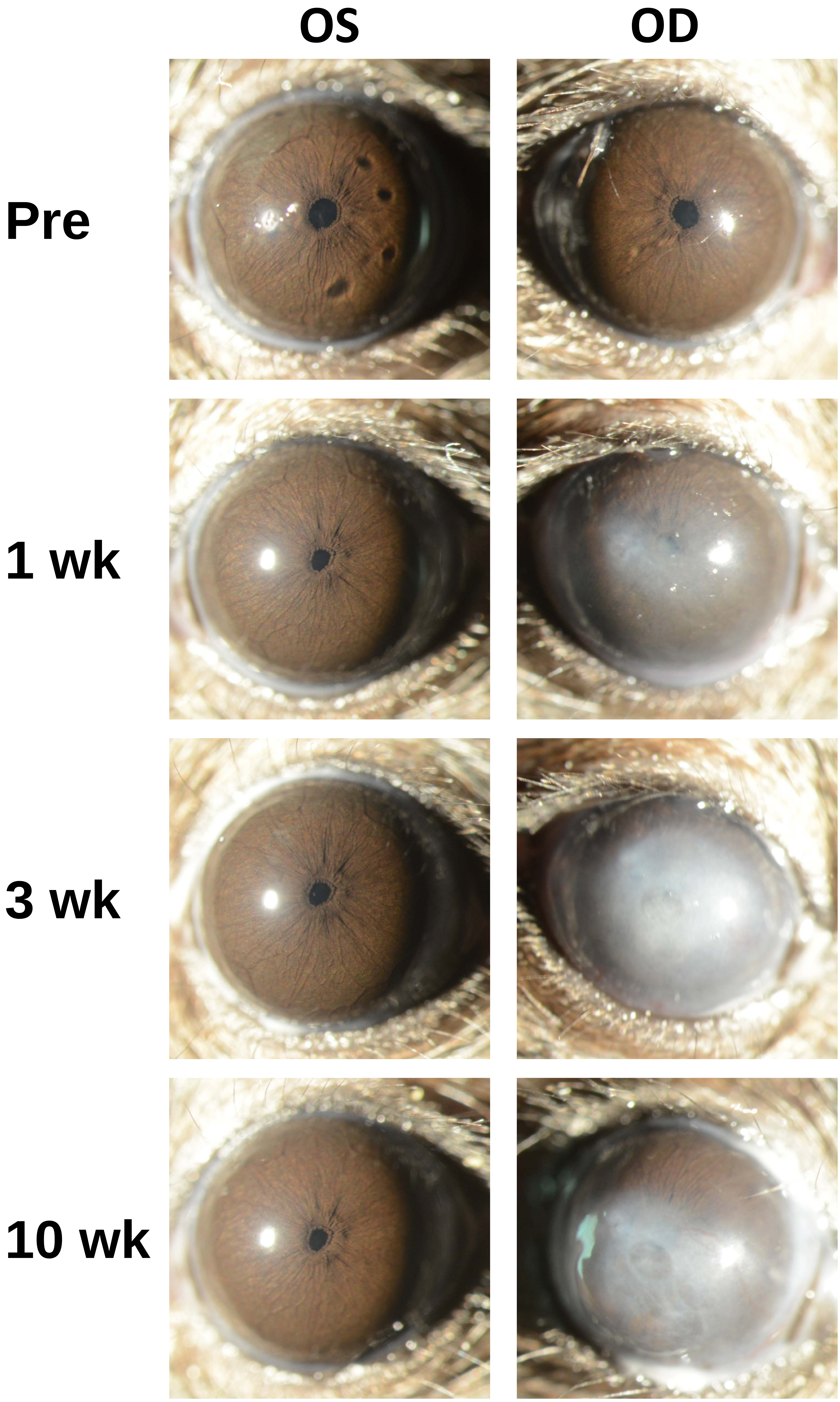
