## Appendix 2 unassigned 40X for "Recombinant adenovirus causes prolonged mobilization of macrophages in the anterior chamber of mice"

Appendix 2
Mouse ID: 28453
10-week timepoint

Masked assignment: **Unassigned**
Actual group: **Virus** (OS injected)

**OS**

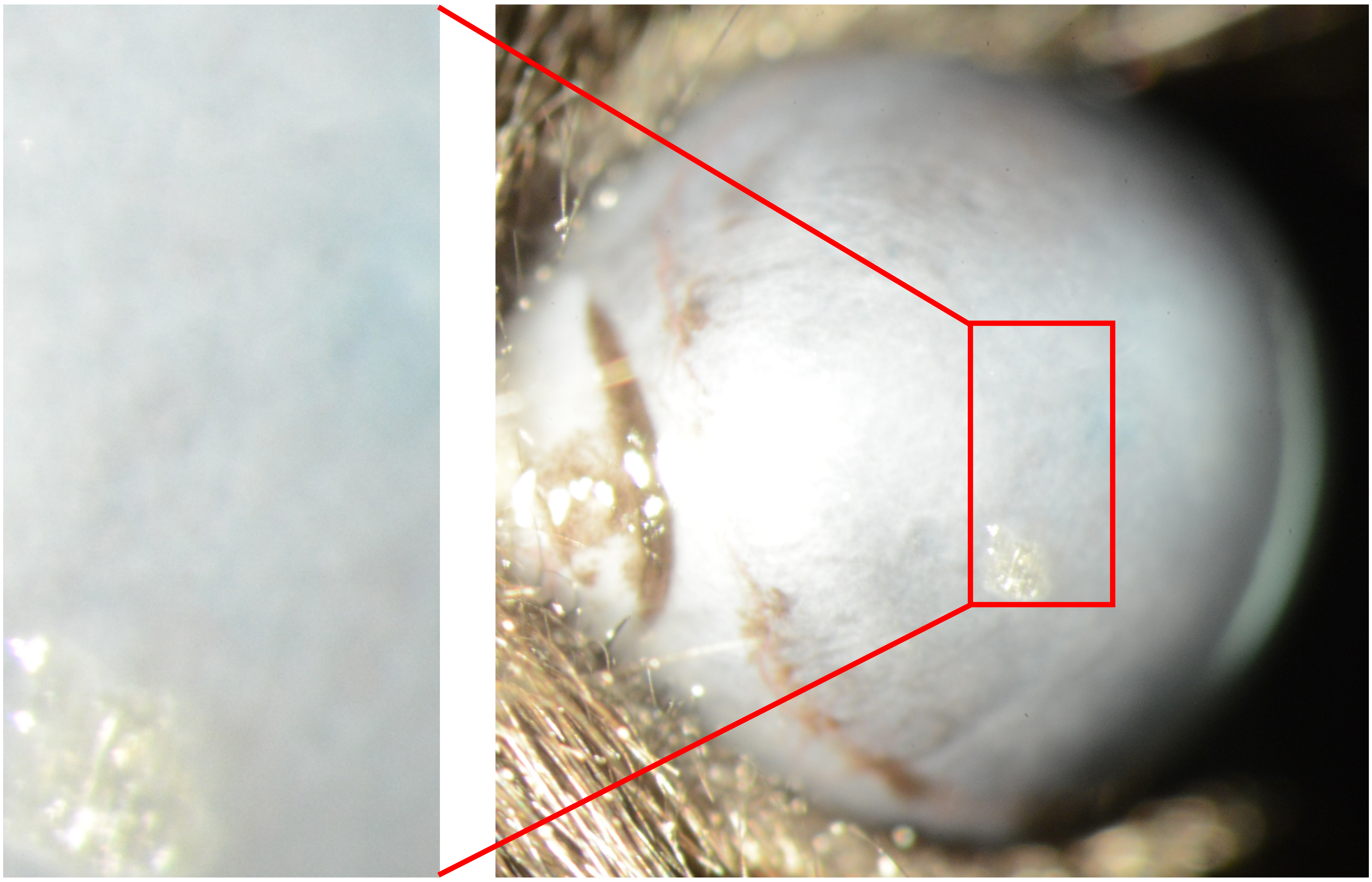

**OD**

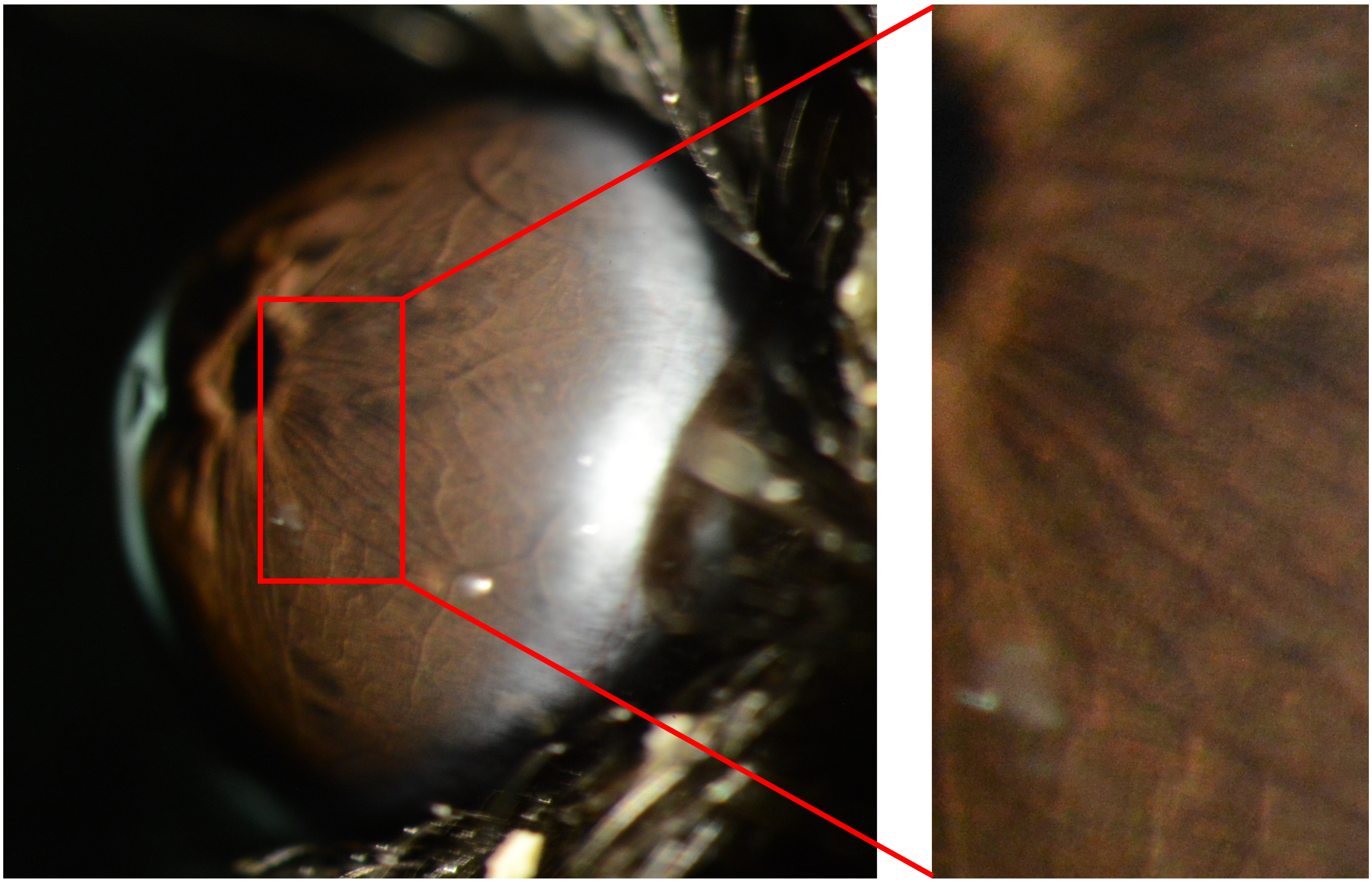

Appendix 2
Mouse ID: 28455
10-week timepoint

Masked assignment: **Unassigned**
Actual group: **Virus** (OD injected)

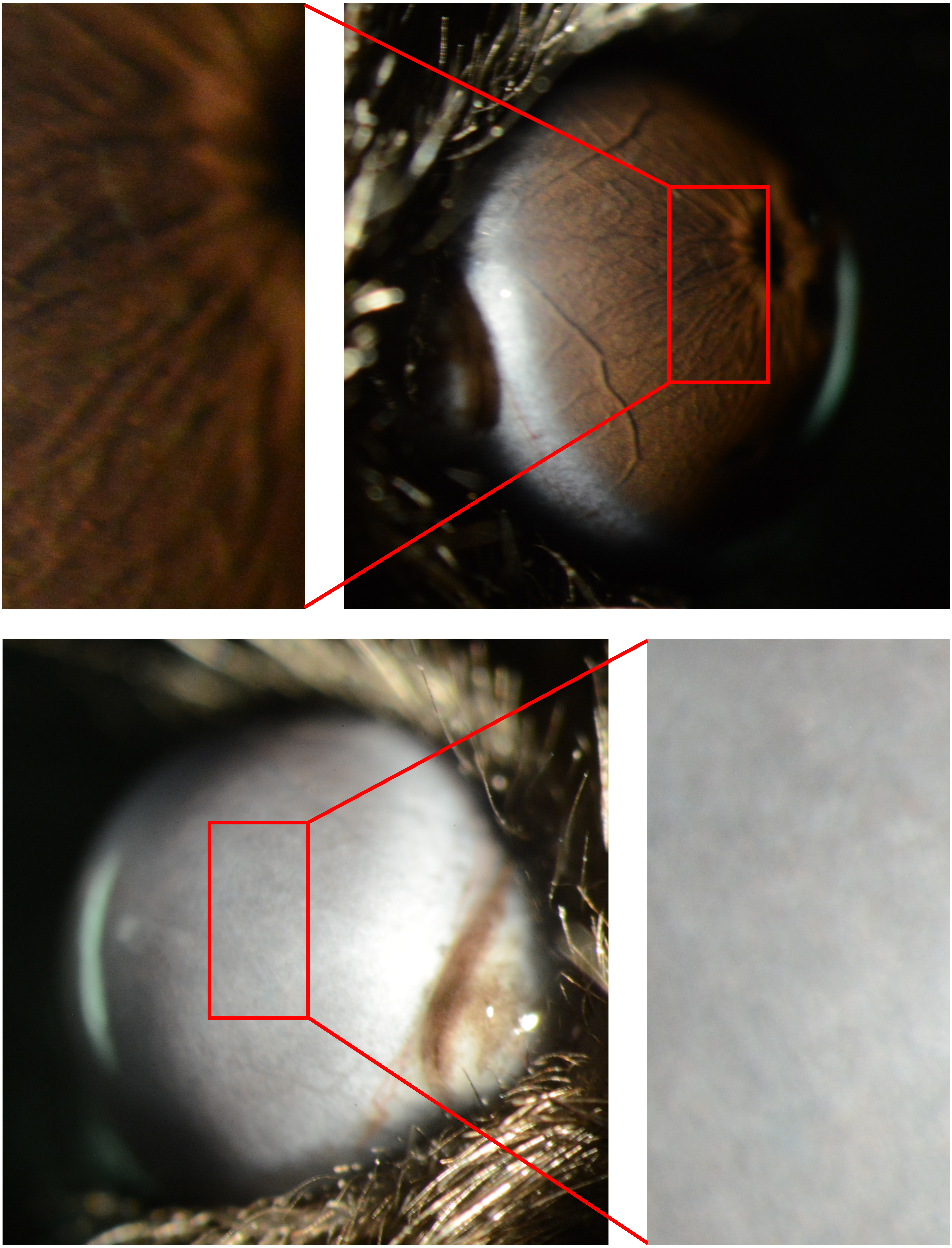

**OS**

**OD**

Appendix 2
Mouse ID: 28469
10-week timepoint

Masked assignment: **Unassigned**
Actual group: **Virus** (OD injected)

**OS**

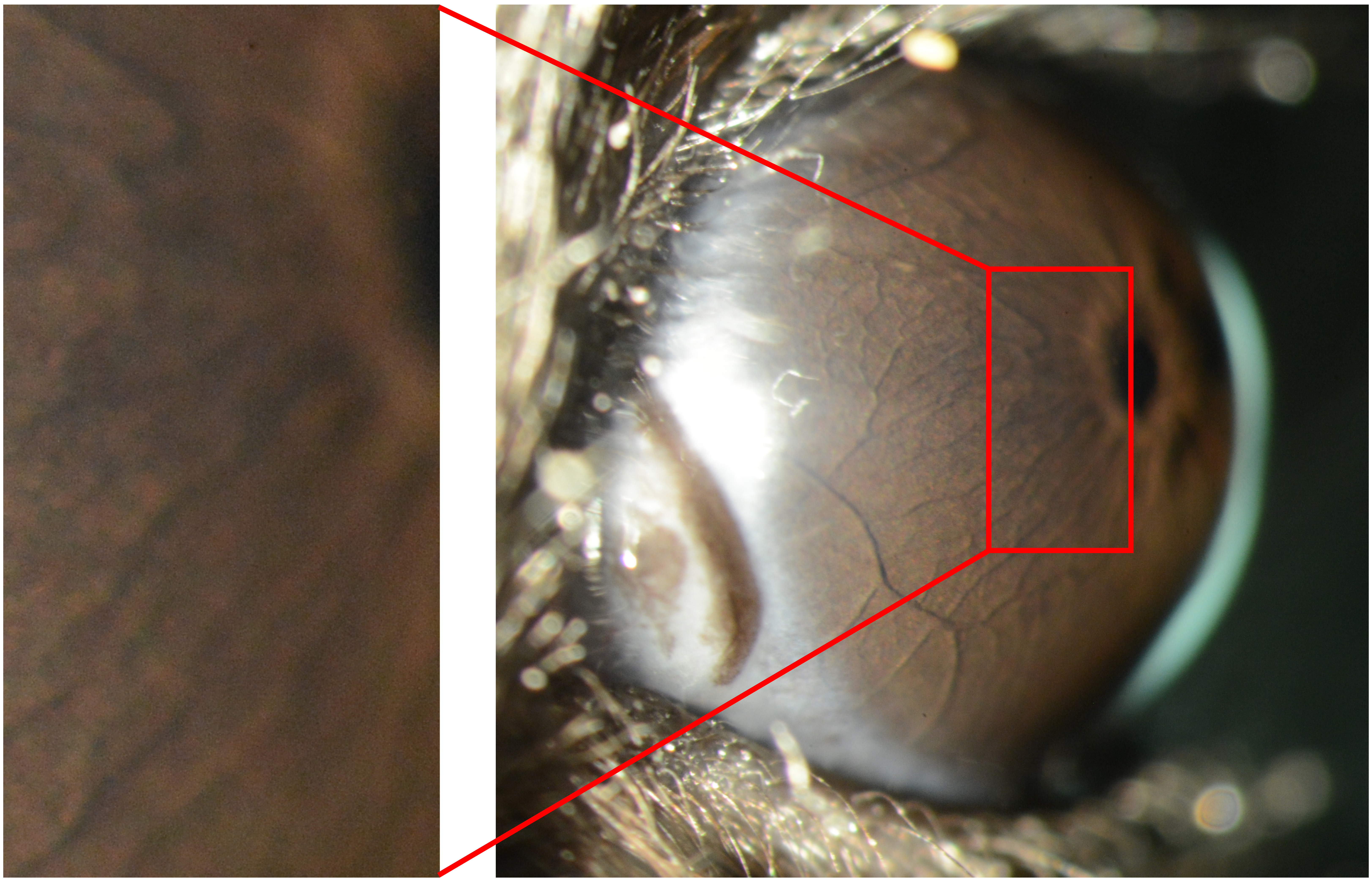

**OD**

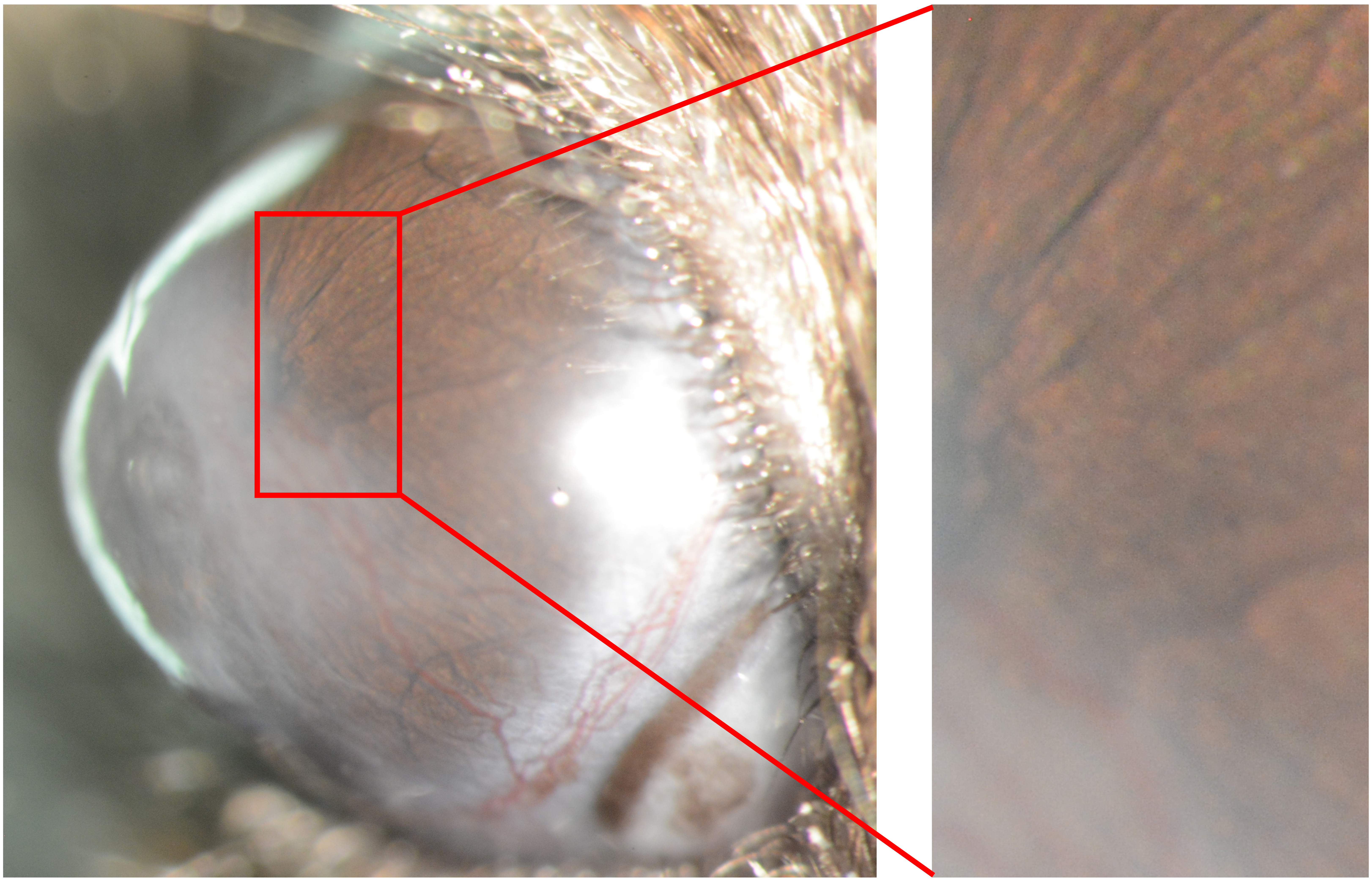
