## Appendix 3 virus 25X for "Recombinant adenovirus causes prolonged mobilization of macrophages in the anterior chamber of mice"

Appendix 3
Mouse ID: 28444

Masked assignment: **Virus**
Actual group: **Virus** (OS injected)

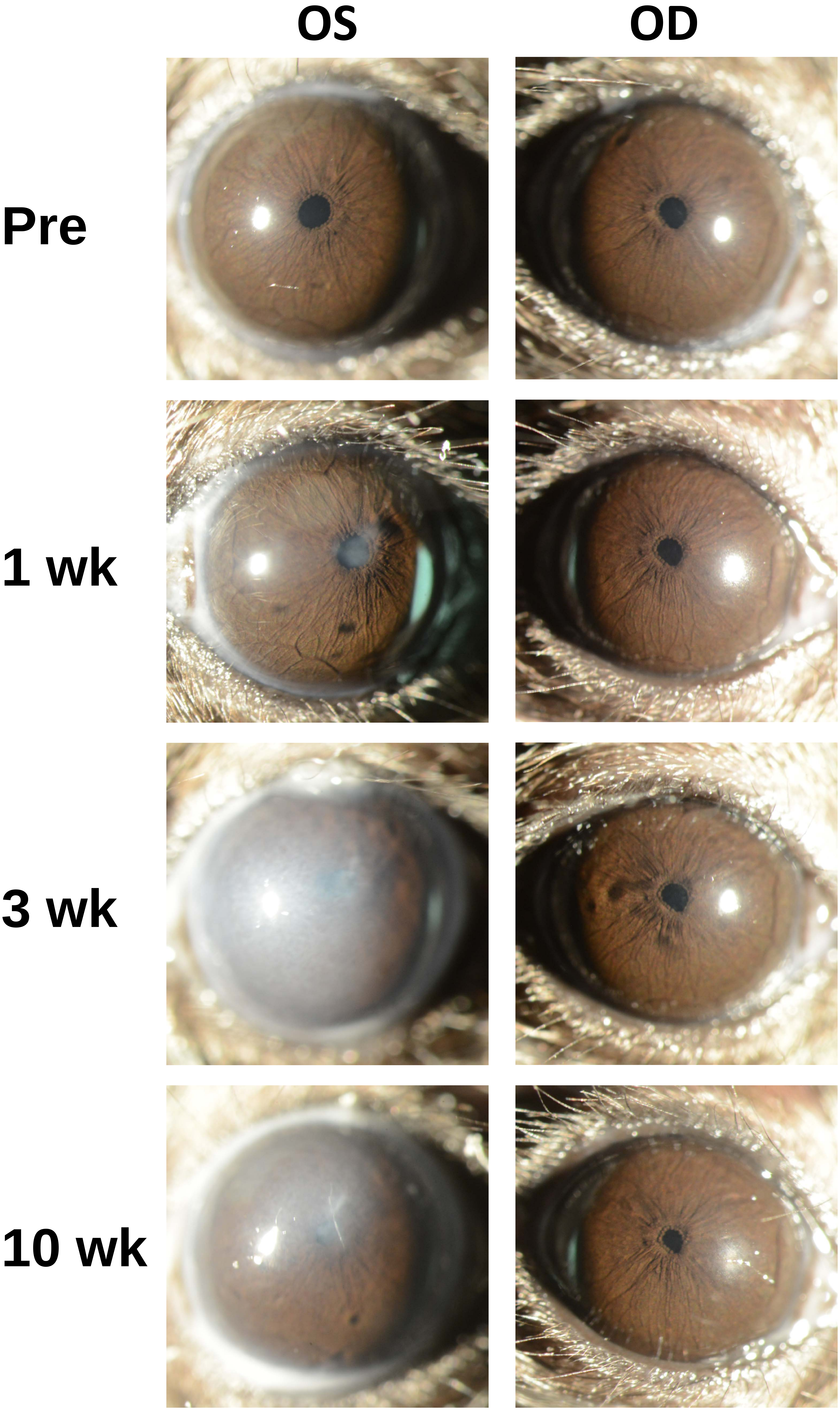

Appendix 3
Mouse ID: 28447

Masked assignment: **Virus**
Actual group: **Virus** (OD injected)

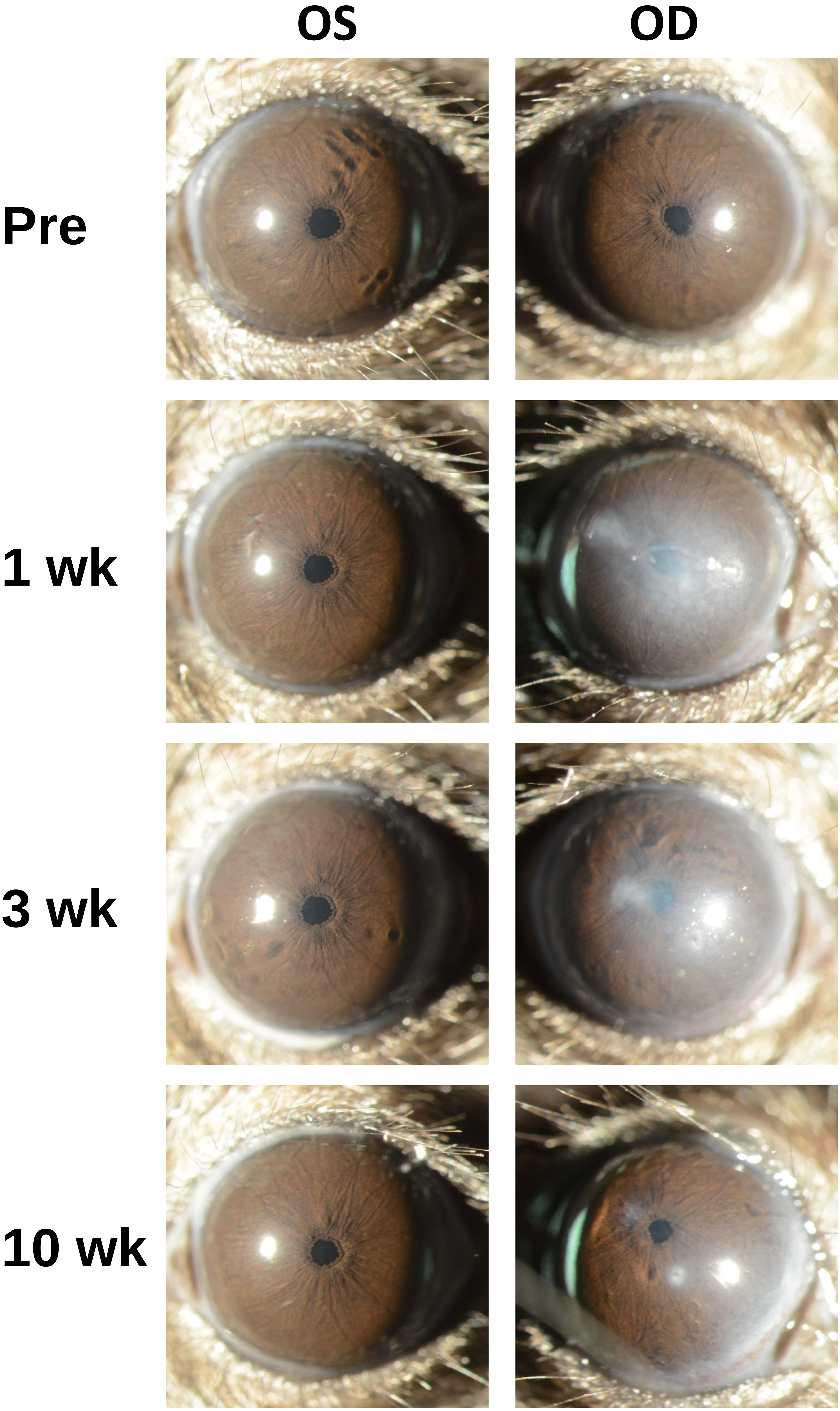

Appendix 3
Mouse ID: 28449

Masked assignment: **Virus**
Actual group: **Virus** (OD injected)

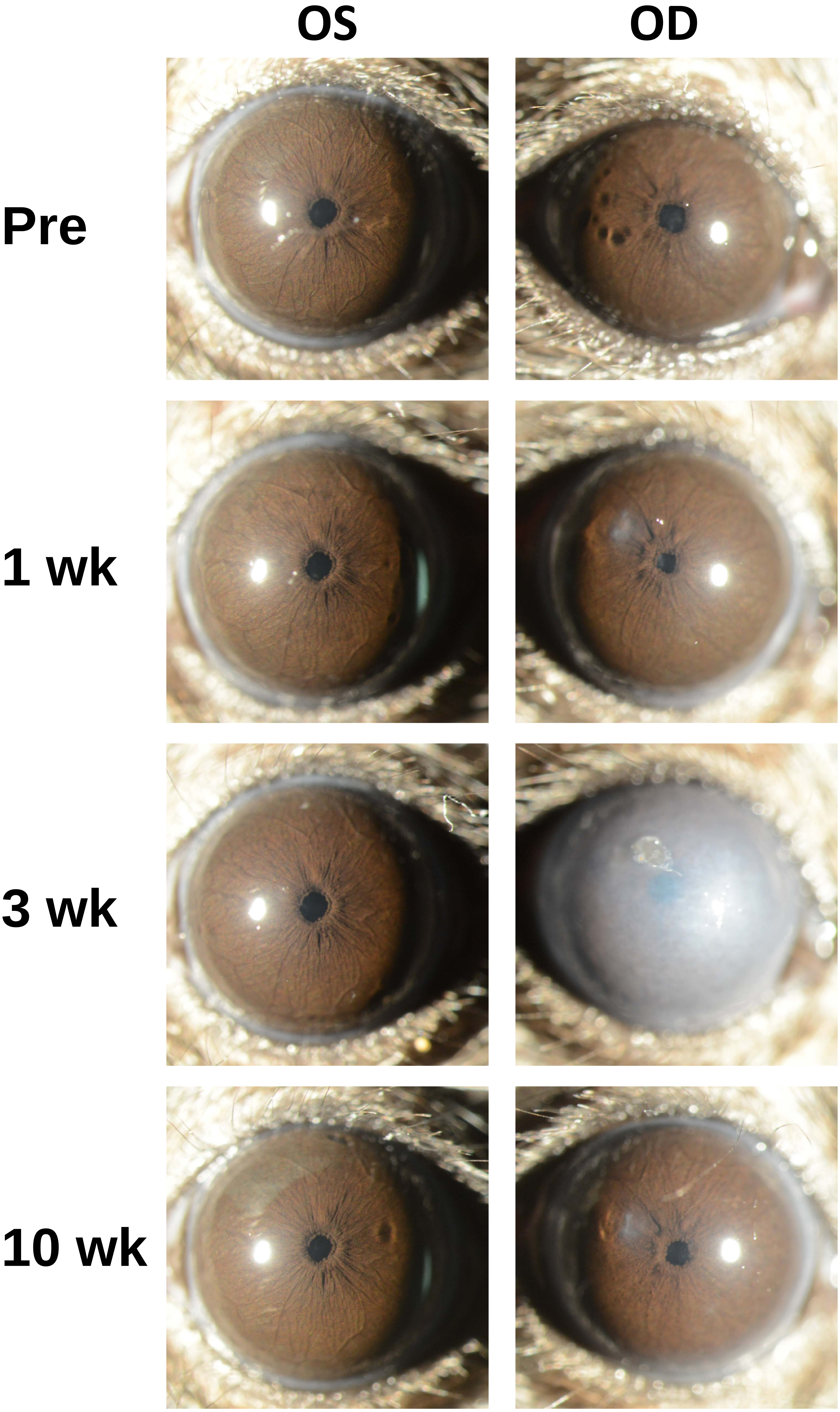

Appendix 3
Mouse ID: 28454

Masked assignment: **Virus**
Actual group: **Virus** (OS injected)

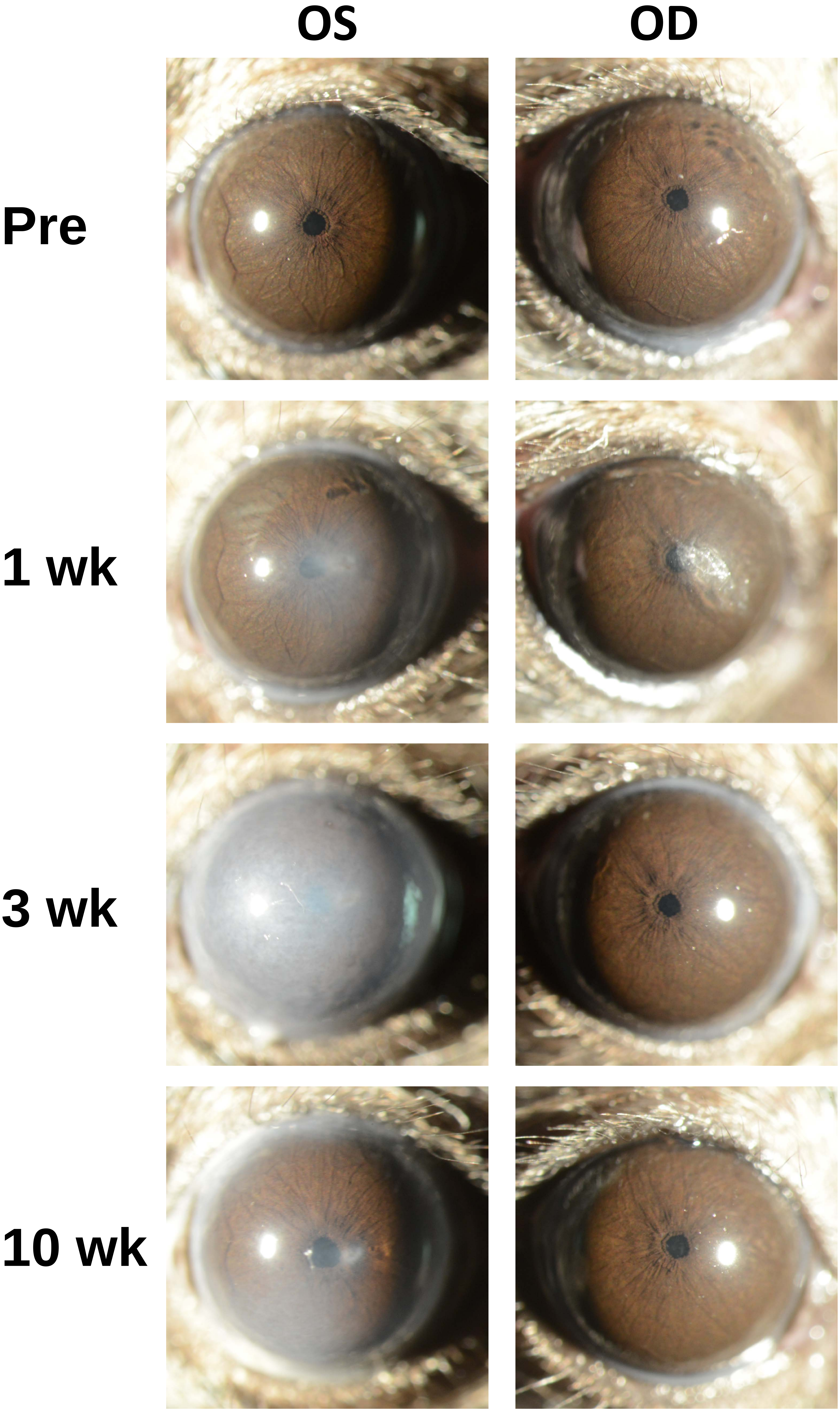

Appendix 3
Mouse ID: 28462

Masked assignment: **Virus**
Actual group: **Virus** (OS injected)

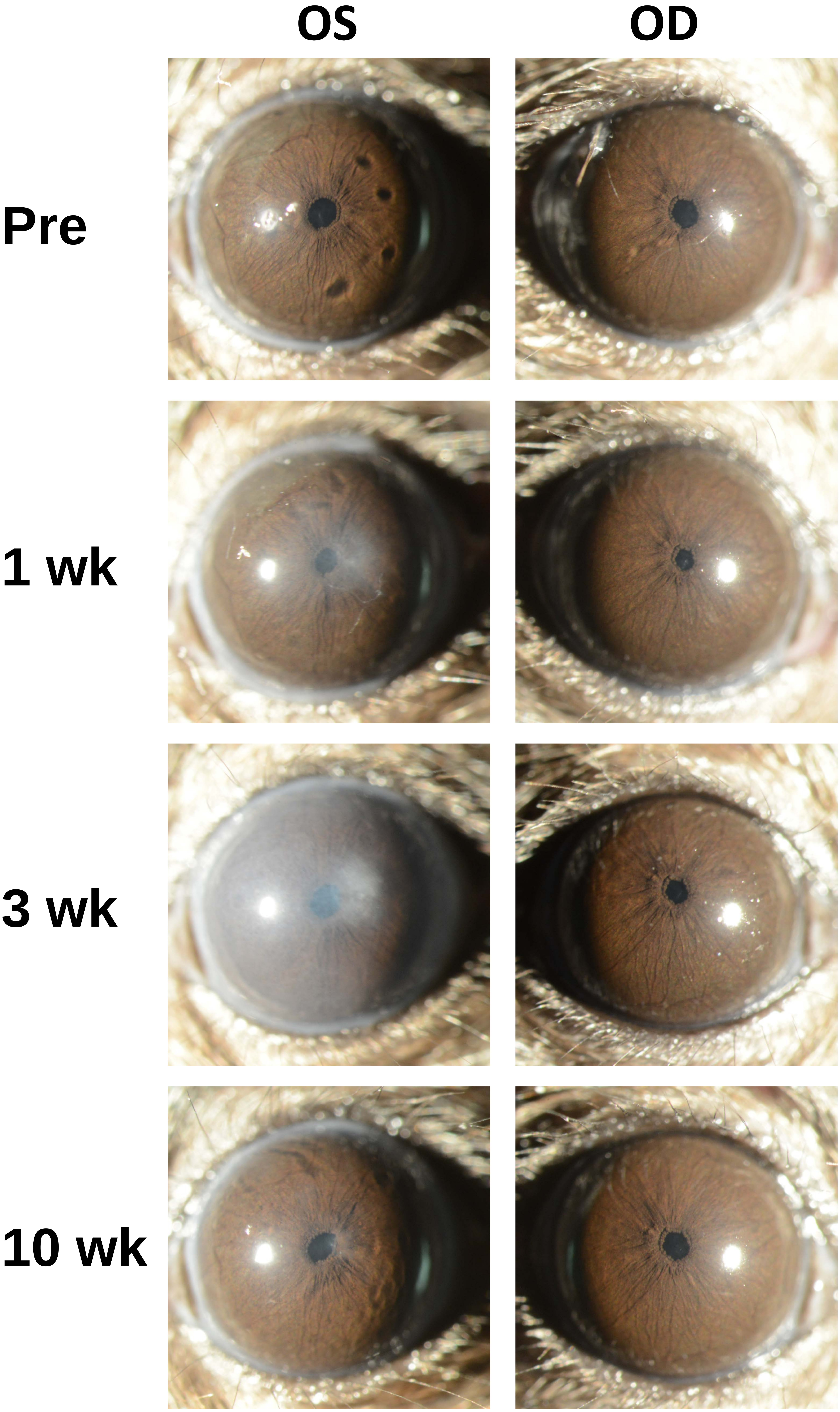

Appendix 3
Mouse ID: 28465

Masked assignment: **Virus**
Actual group: **Virus** (OS injected)

**OS** **OD**

**Pre**

**1 wk**

**3 wk**

**10 wk**

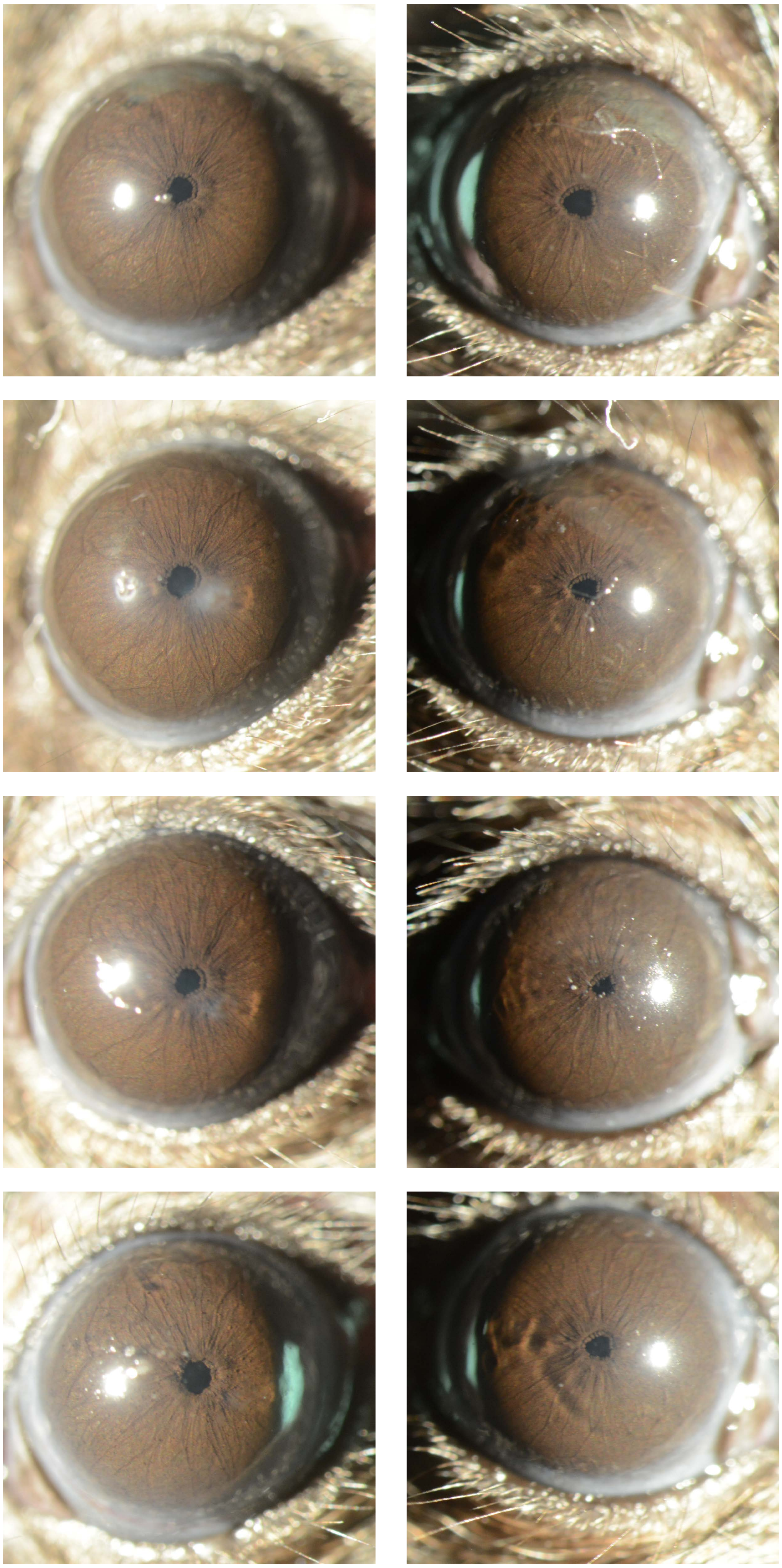

Appendix 3
Mouse ID: 28467

Masked assignment: **Virus**
Actual group: **Virus** (OD injected)

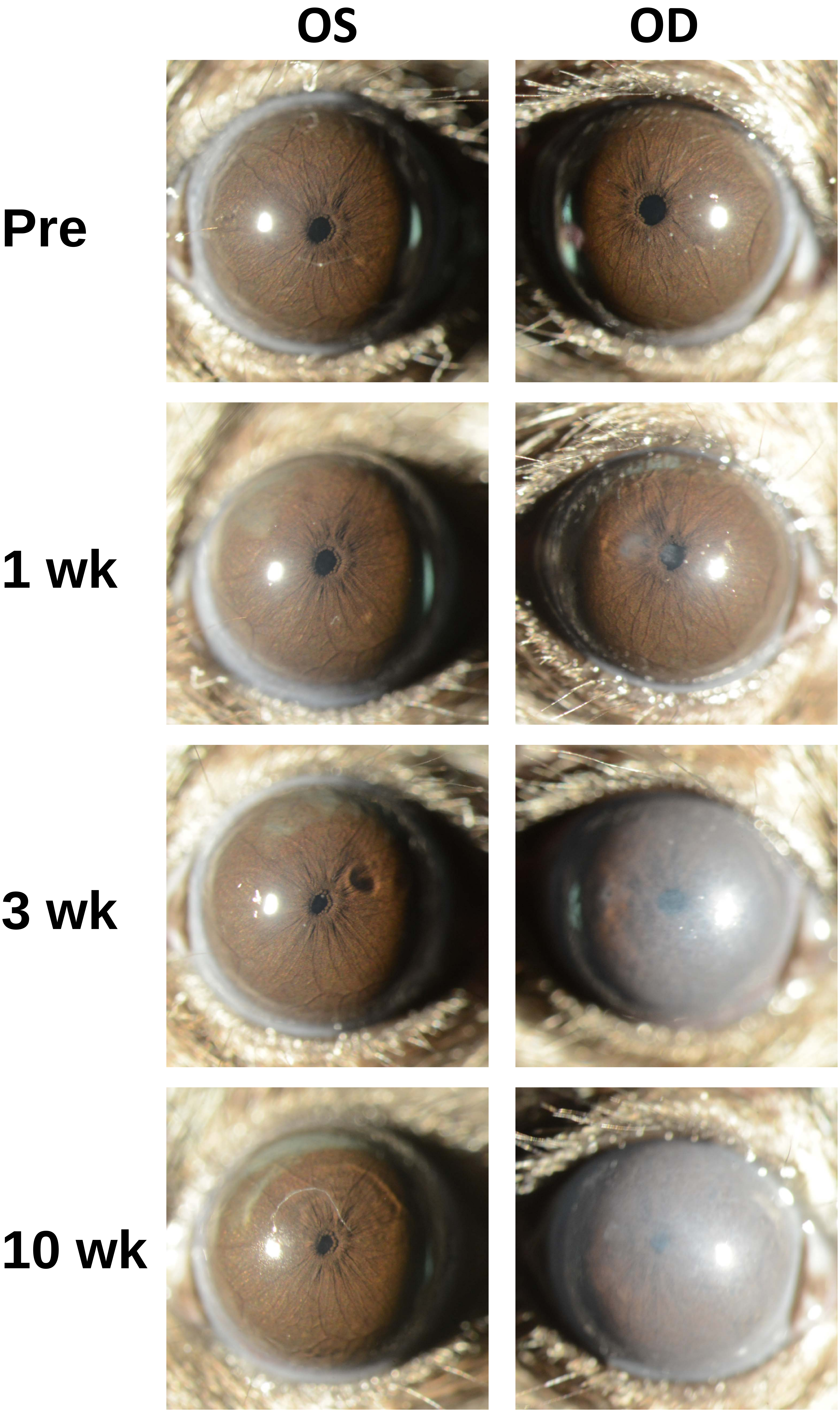

Appendix 3
Mouse ID: 28486

Masked assignment: **Virus**
Actual group: **Virus** (OD injected)

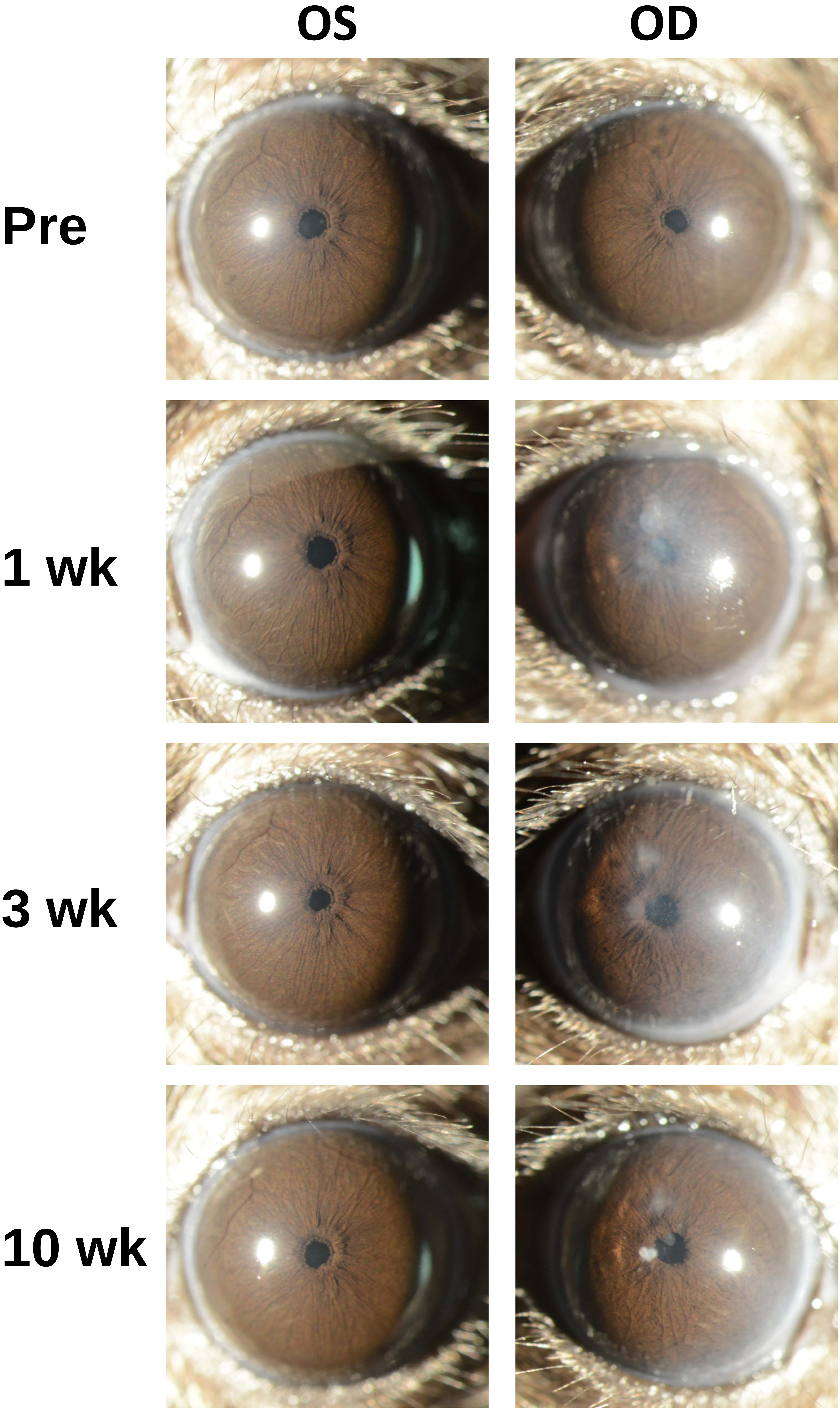
