## Appendix 4 virus 40X for "Recombinant adenovirus causes prolonged mobilization of macrophages in the anterior chamber of mice"

Appendix 4
Mouse ID: 28444
10-week timepoint

Masked assignment: **Virus**
Actual group: **Virus** (OS injected)

**OS**

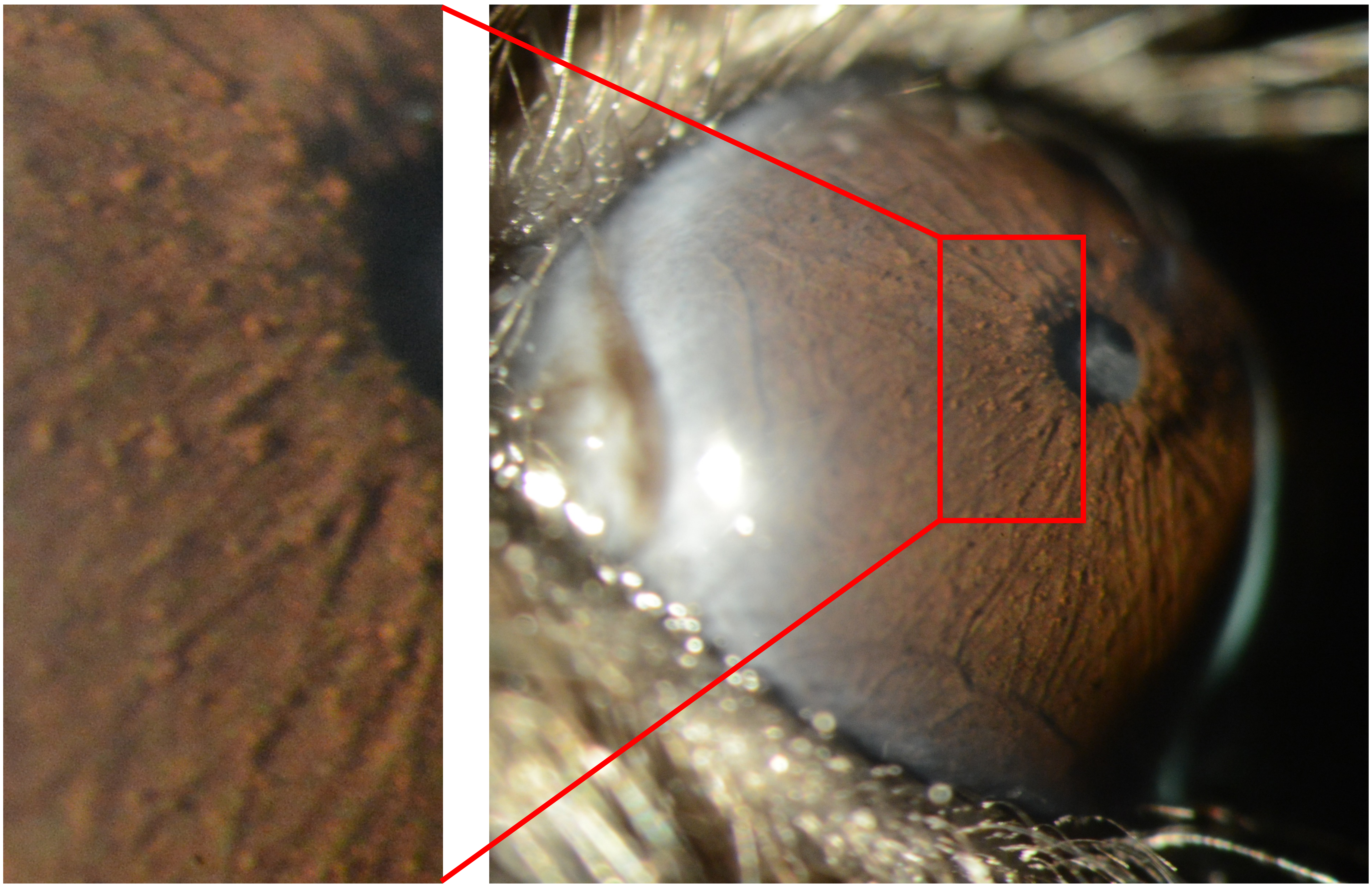

**OD**

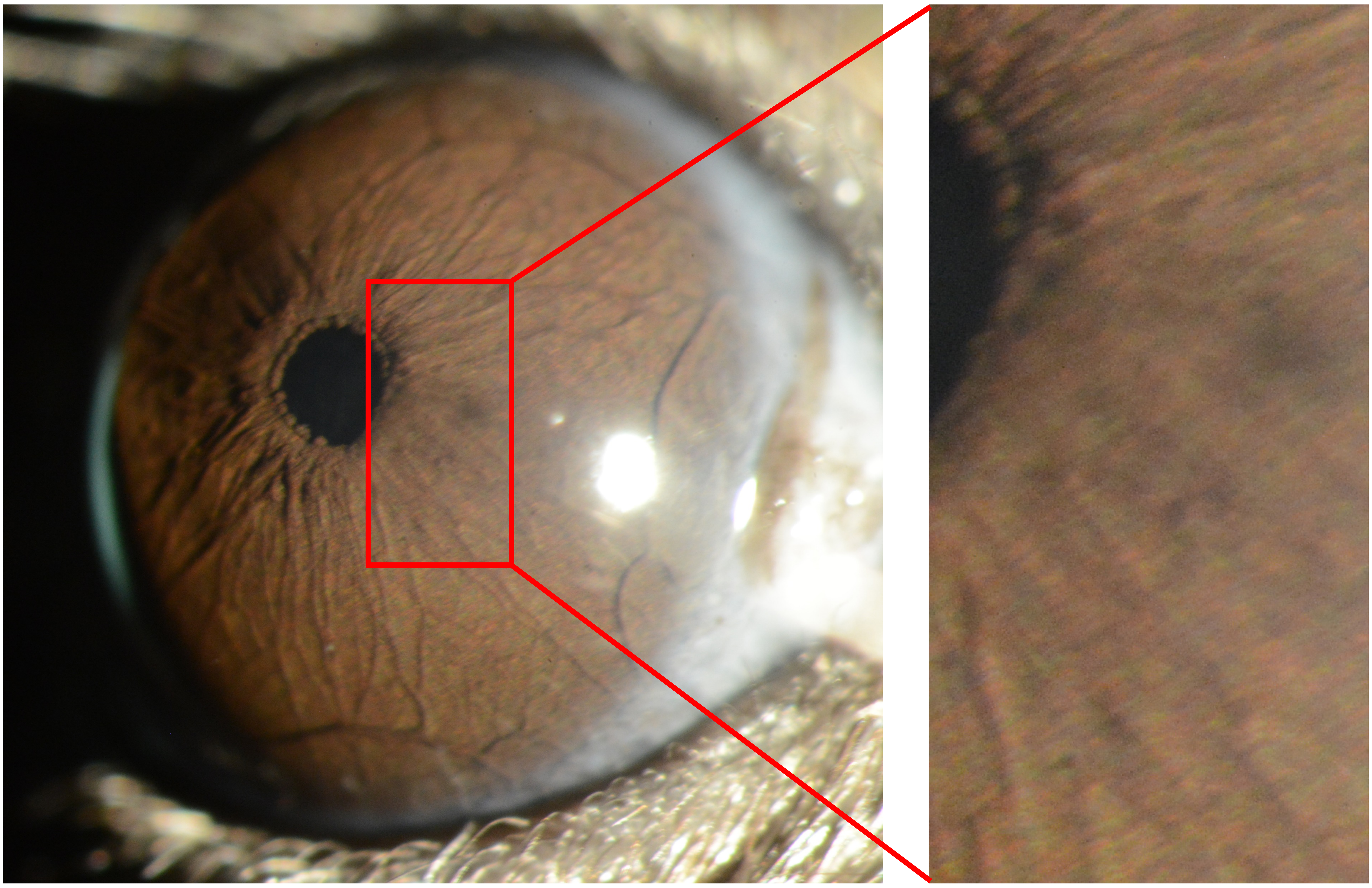

Appendix 4
Mouse ID: 28447
10-week timepoint

Masked assignment: **Virus**
Actual group: **Virus** (OD injected)

**OS**

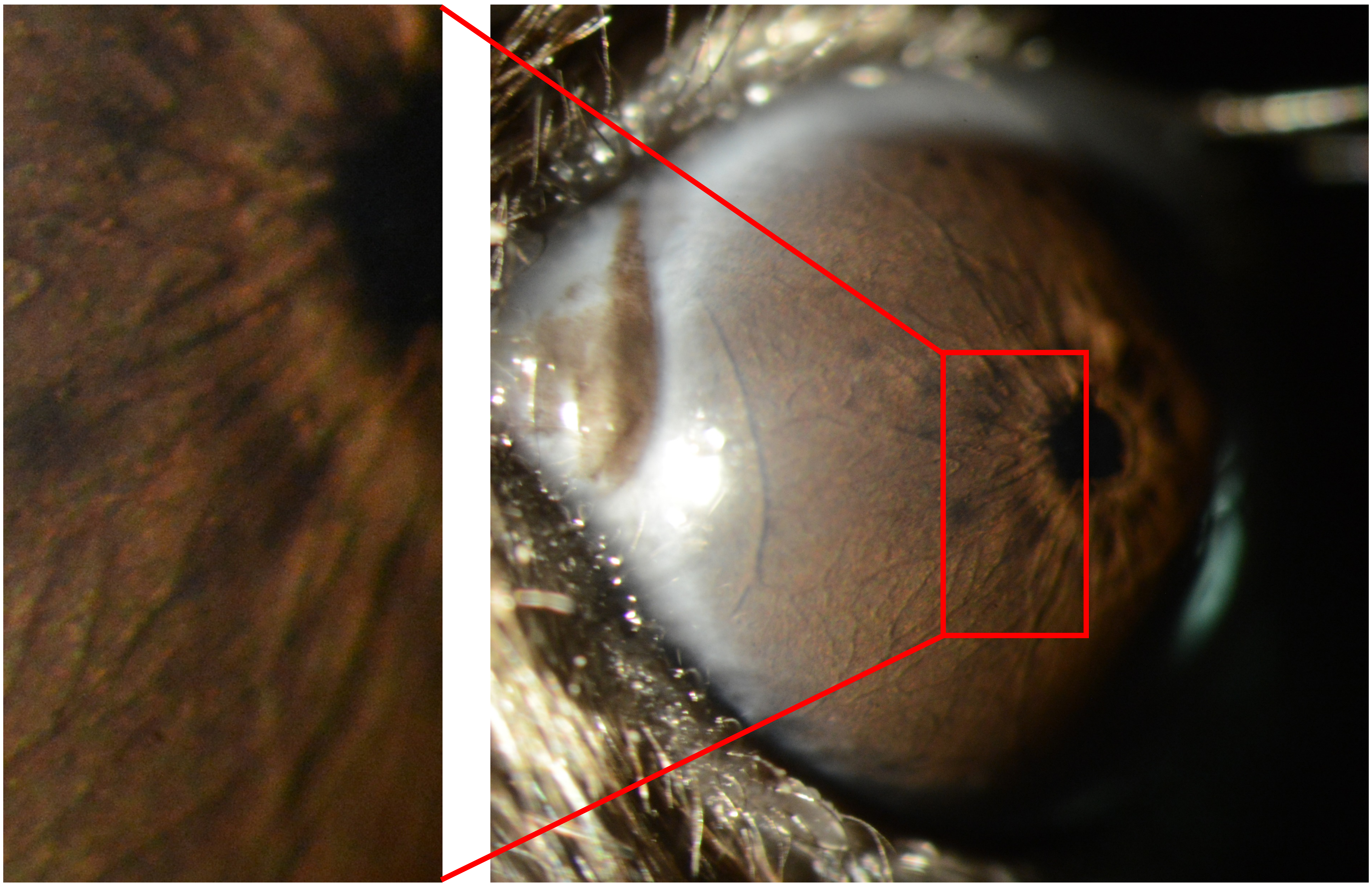

**OD**

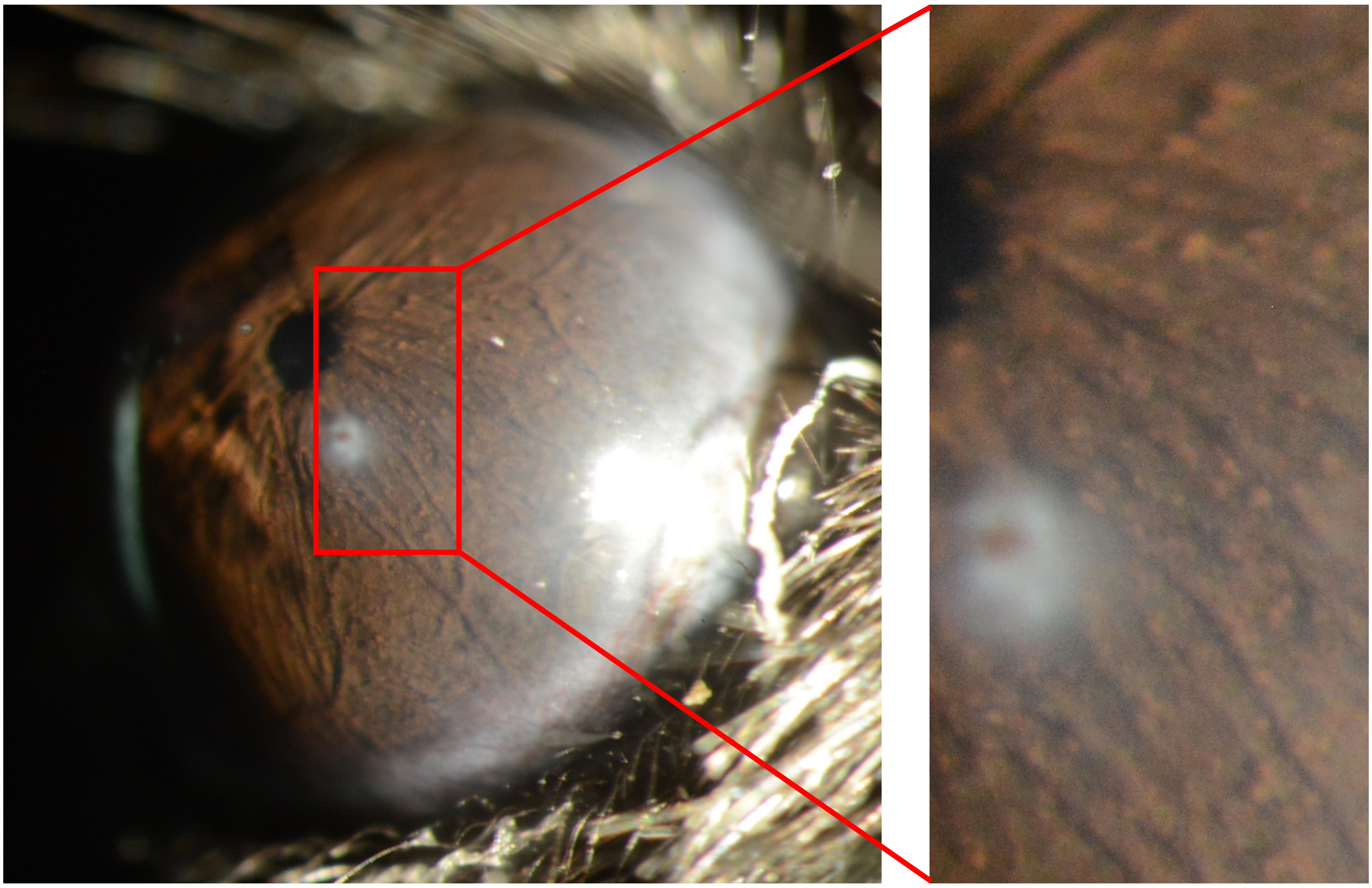

Appendix 4
Mouse ID: 28449
10-week timepoint

Masked assignment: **Virus**
Actual group: **Virus** (OD injected)

**OS**

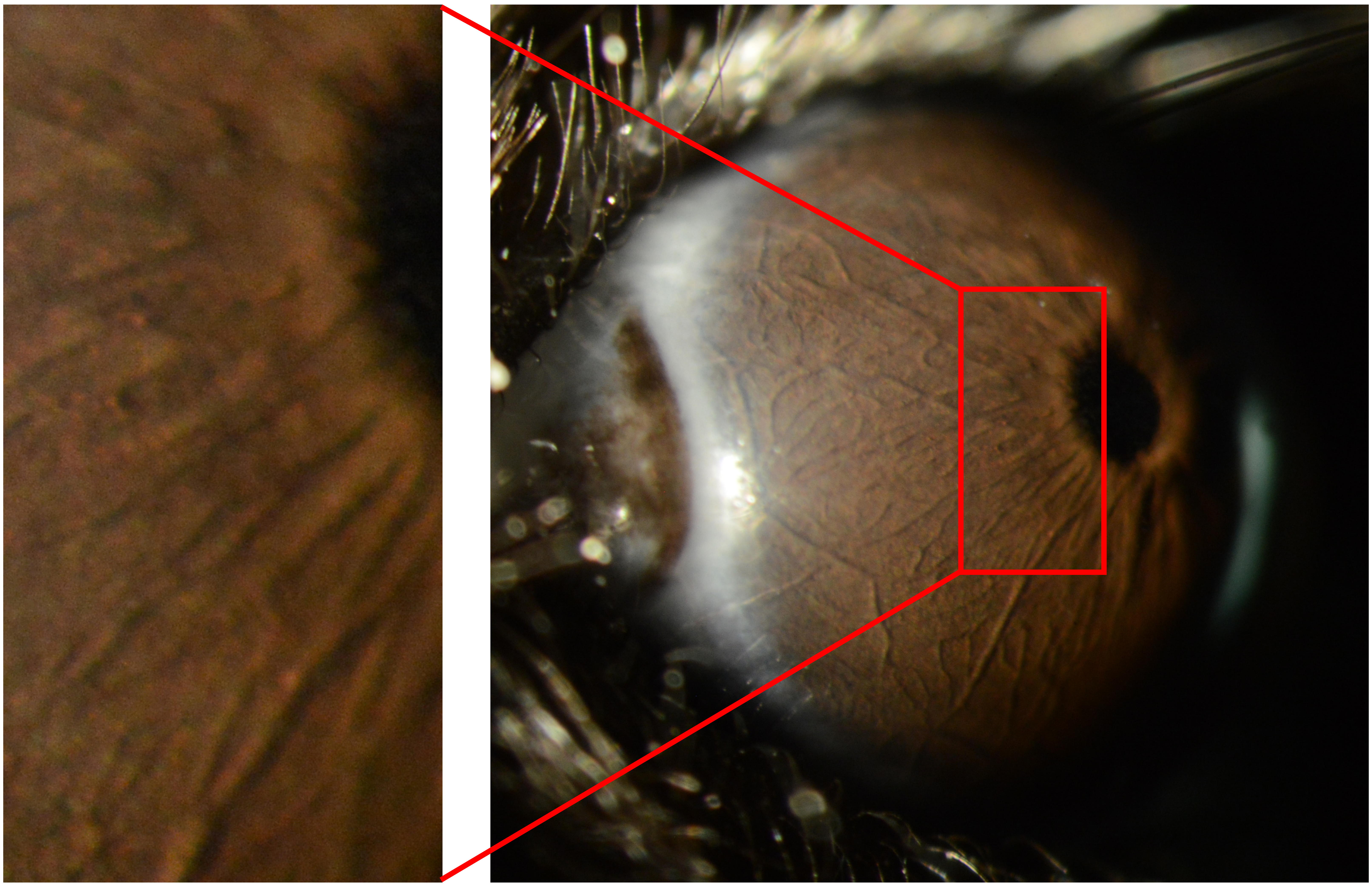

**OD**

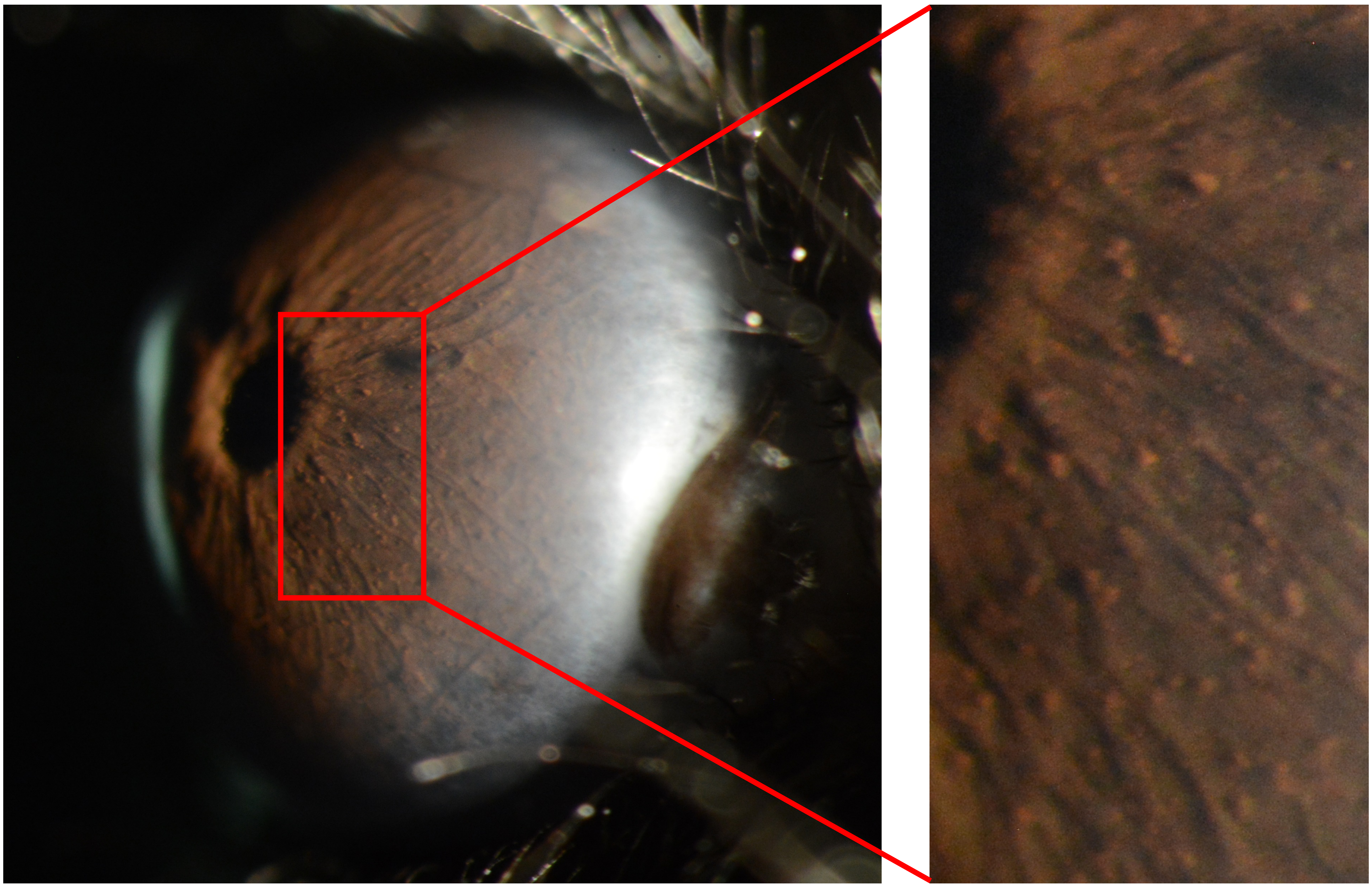

Appendix 4
Mouse ID: 28454
10-week timepoint

Masked assignment: **Virus**
Actual group: **Virus** (OS injected)

**OS**

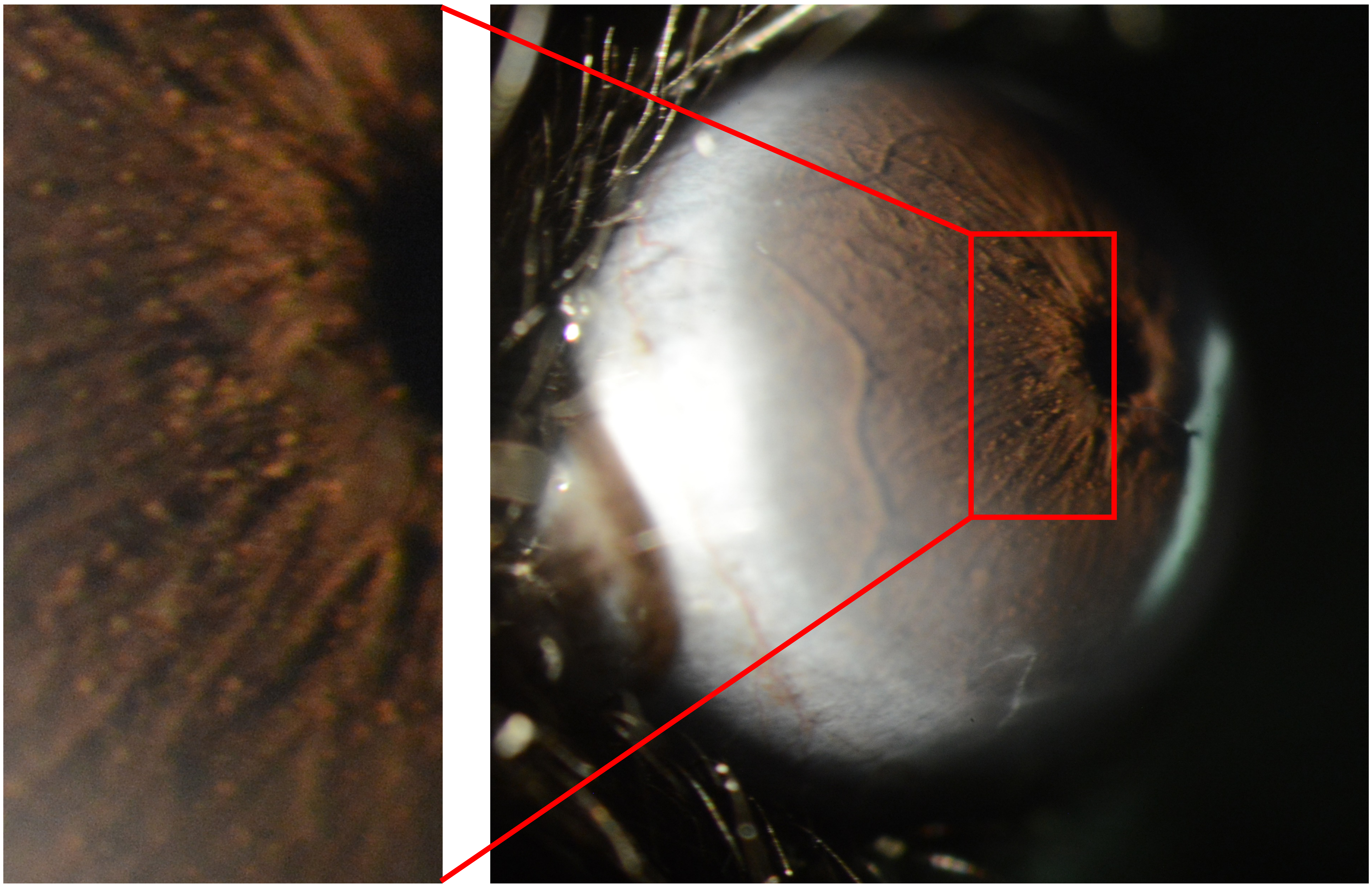

**OD**

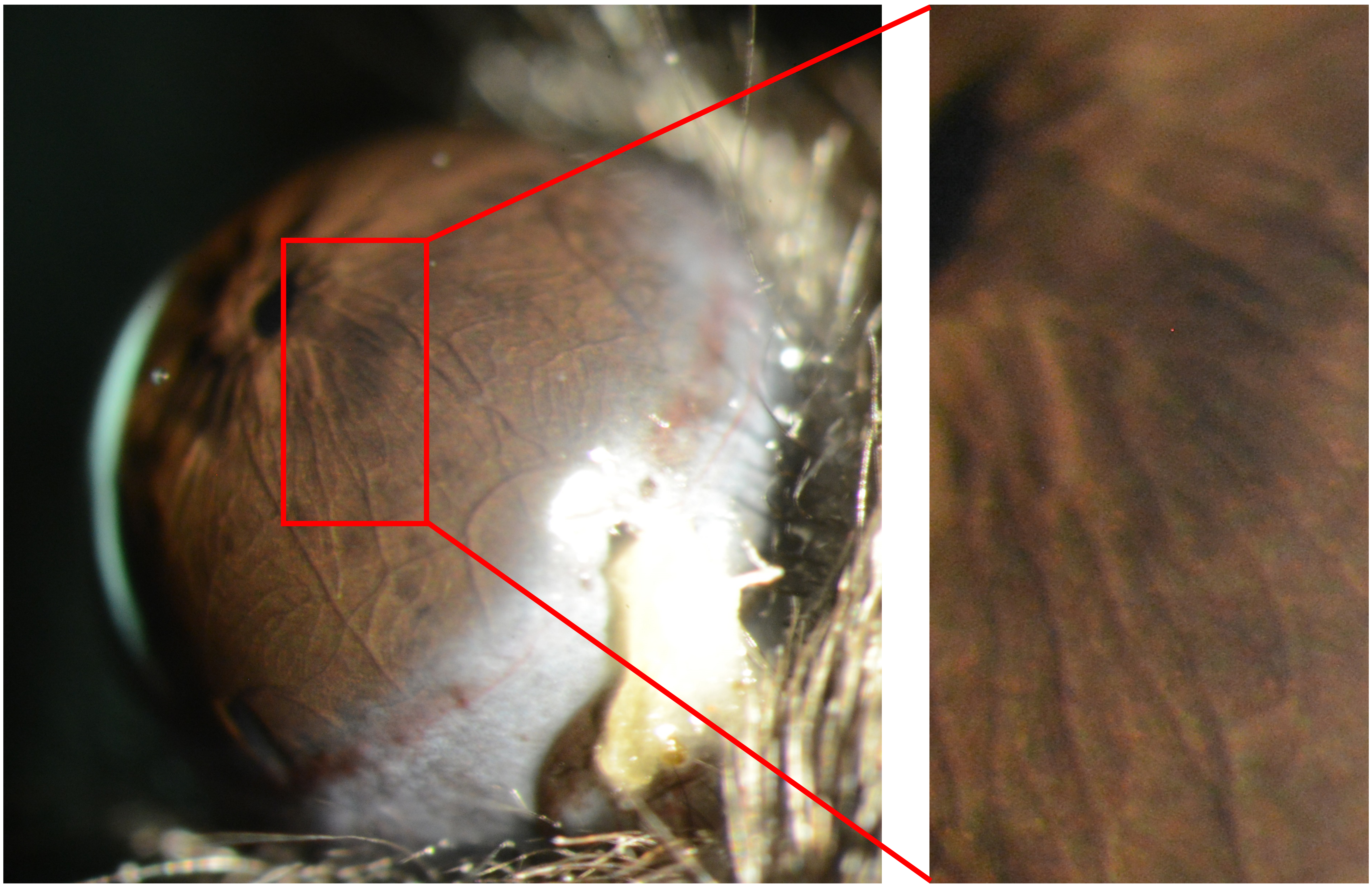

Appendix 4
Mouse ID: 28462
10-week timepoint

Masked assignment: **Virus**
Actual group: **Virus** (OS injected)

**OS**

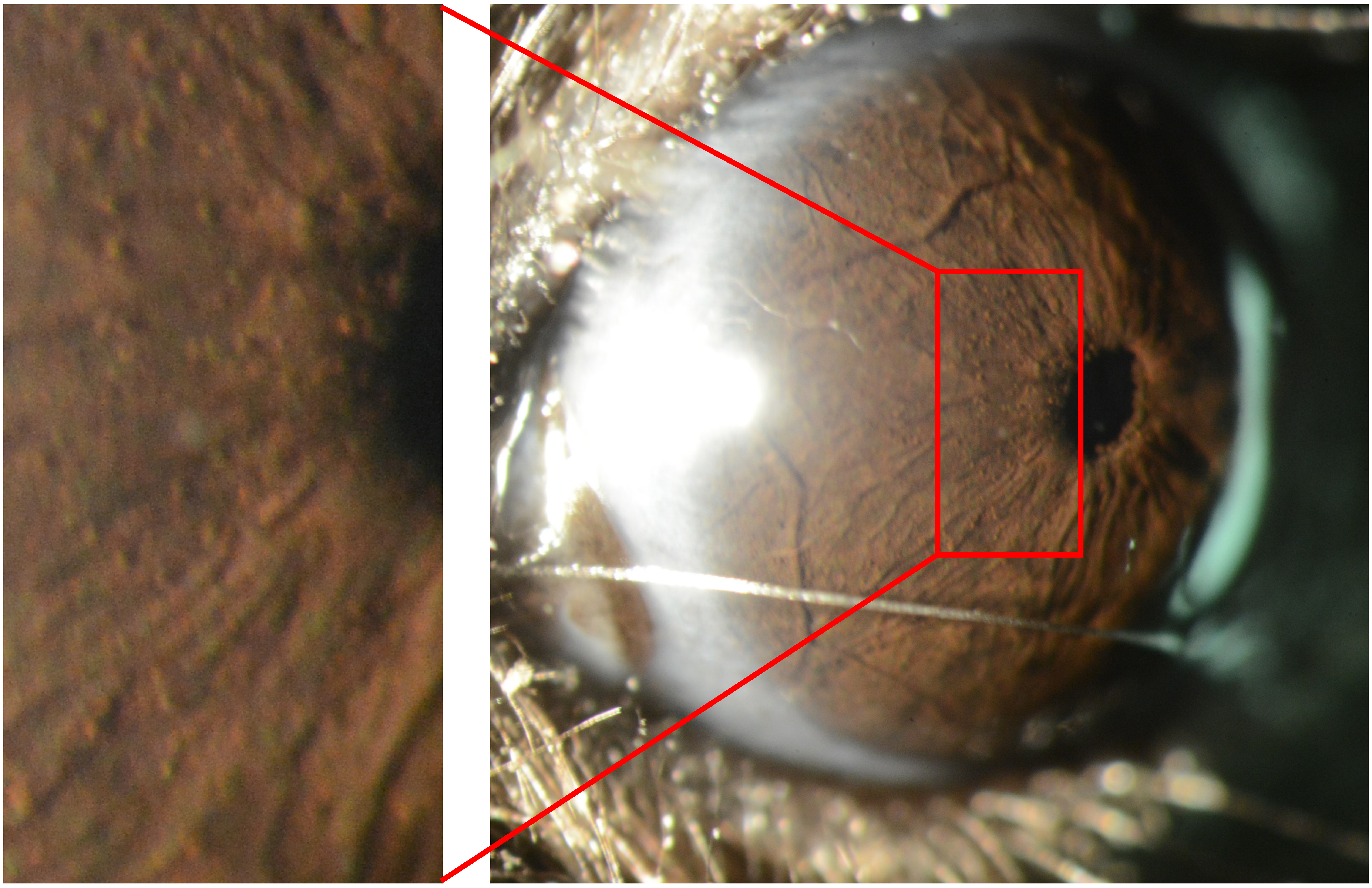

**OD**

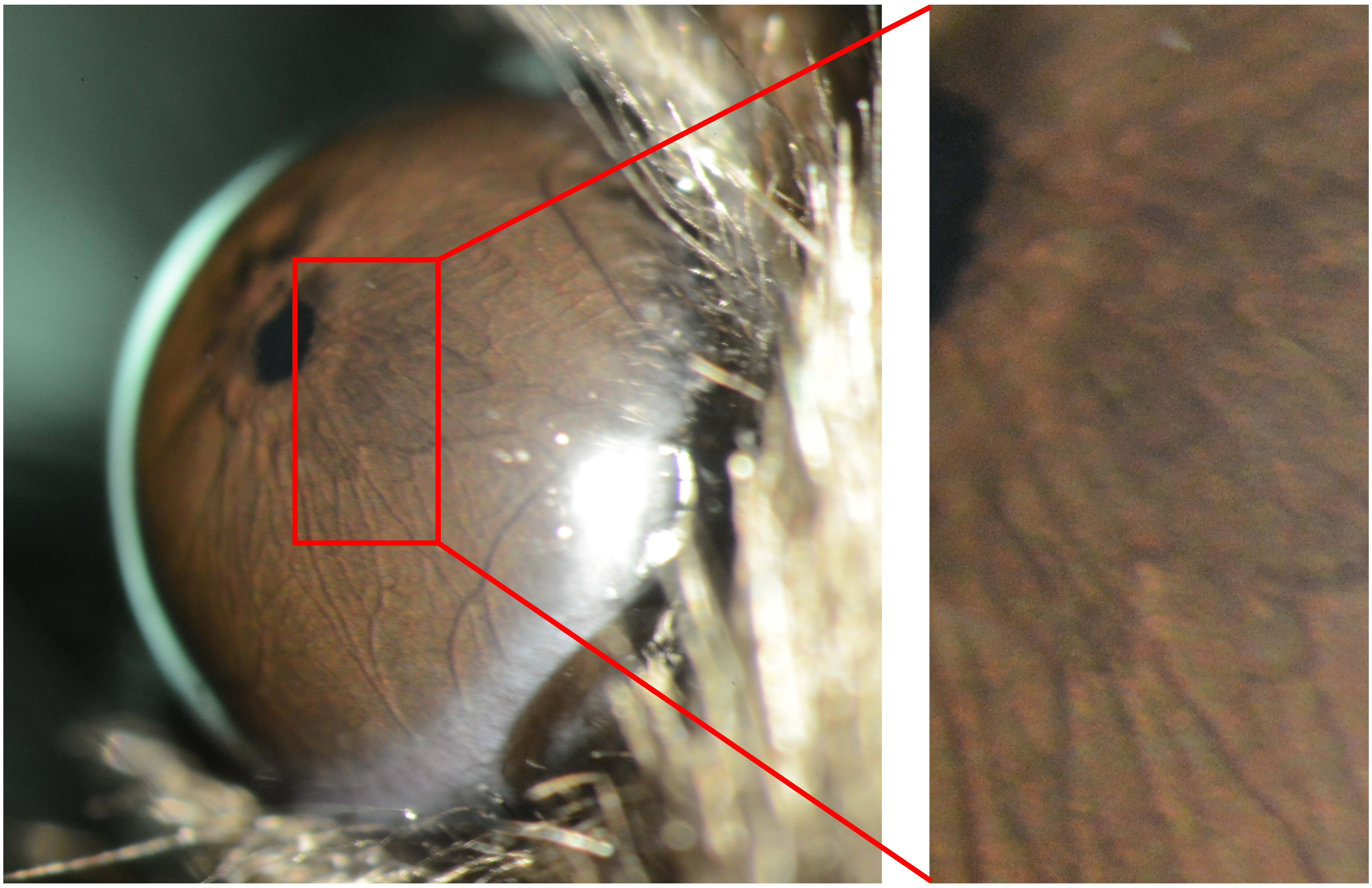

Appendix 4
Mouse ID: 28465
10-week timepoint

Masked assignment: **Virus**
Actual group: **Virus** (OS injected)

**OS**

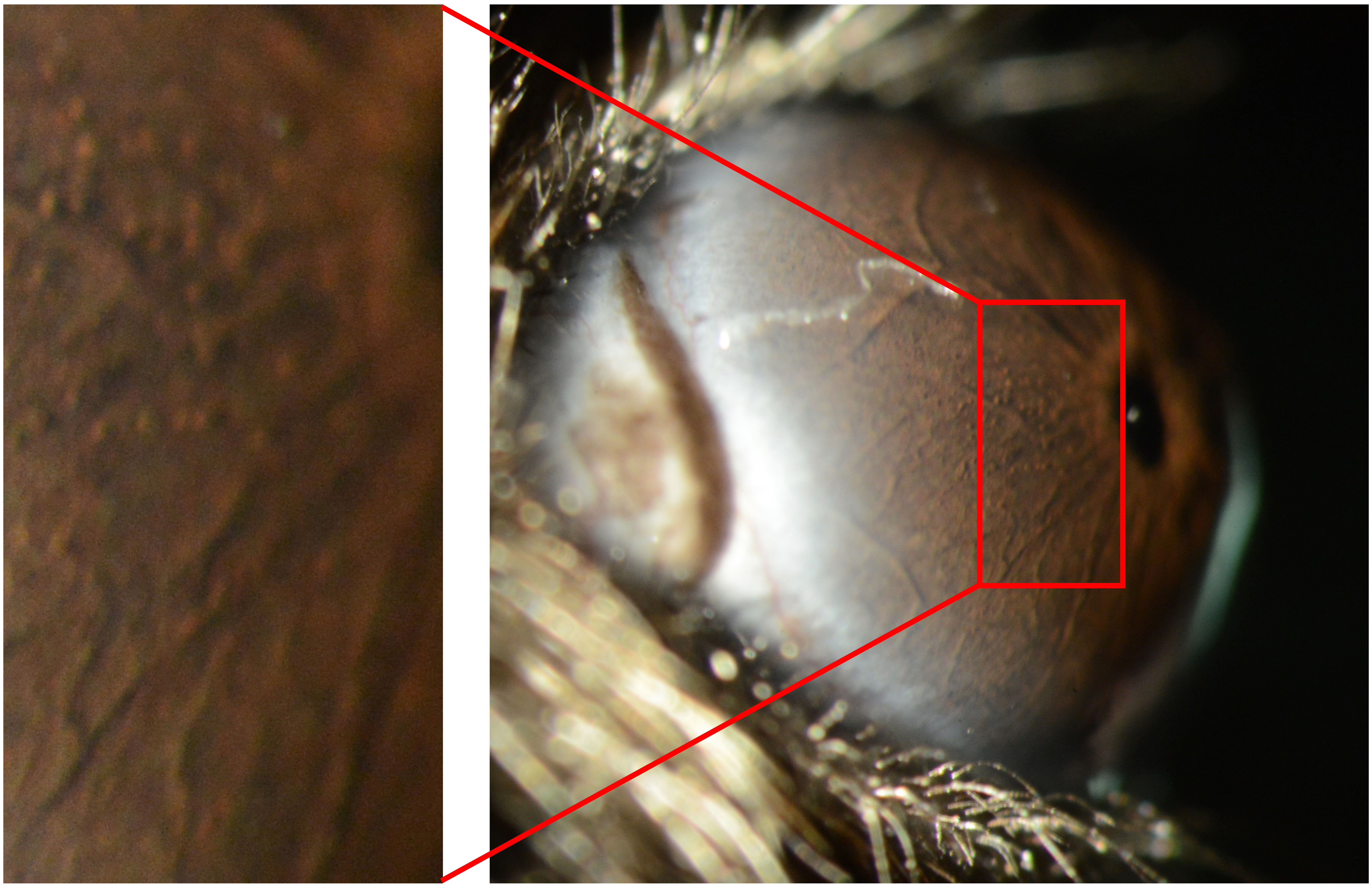

**OD**

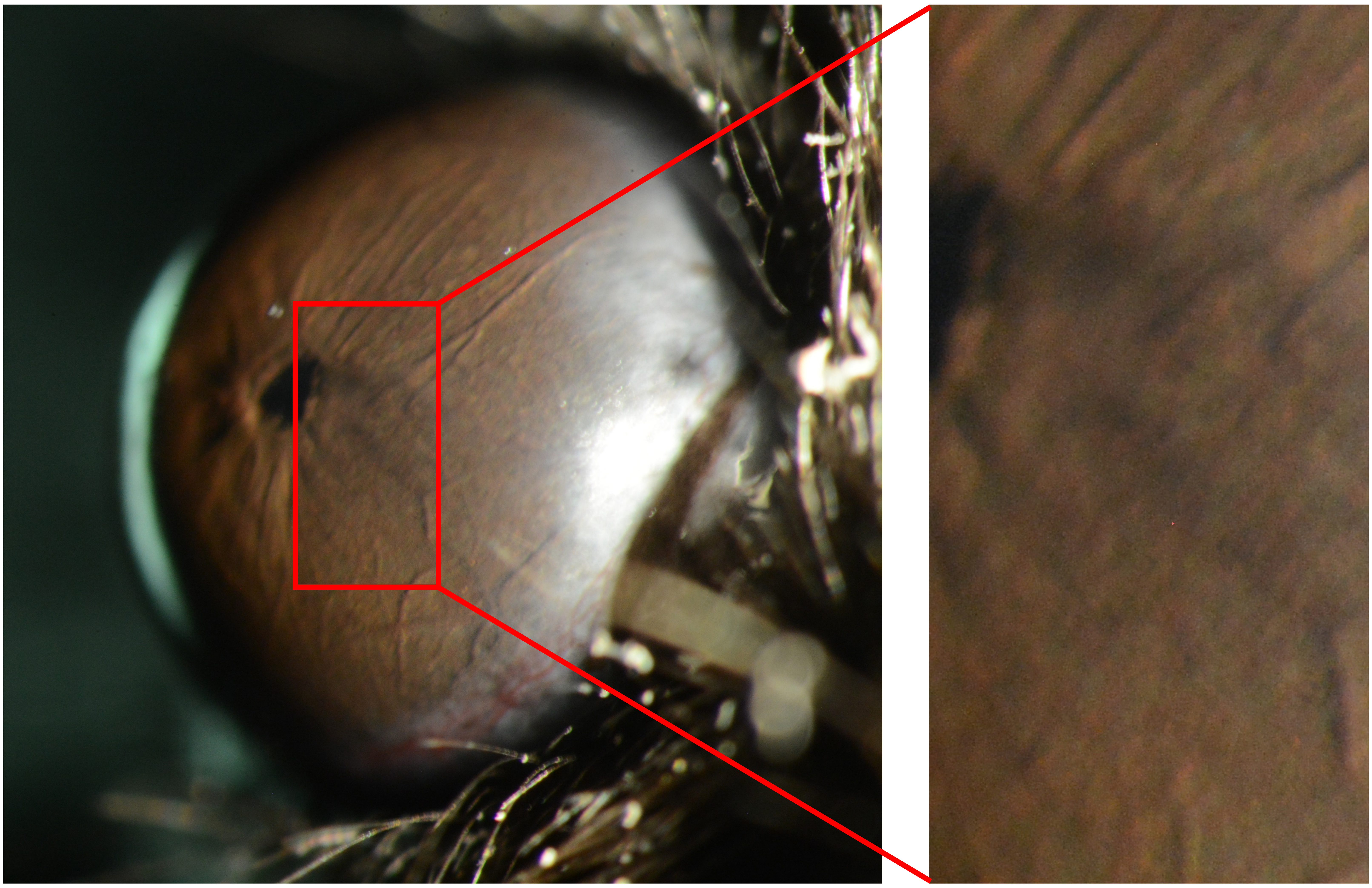

Appendix 4
Mouse ID: 28467
10-week timepoint

Masked assignment: **Virus**
Actual group: **Virus** (OD injected)

**OS**

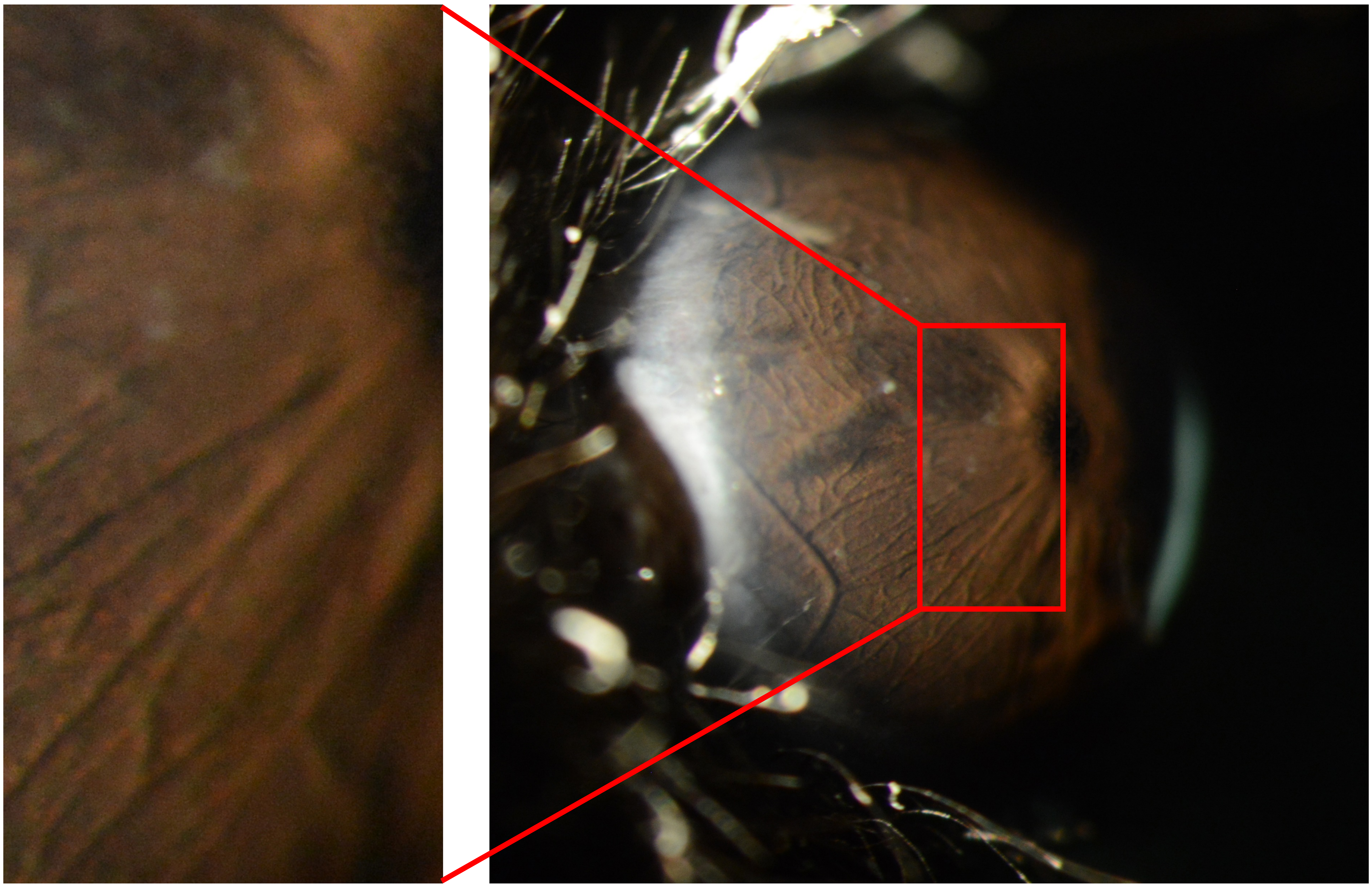

**OD**

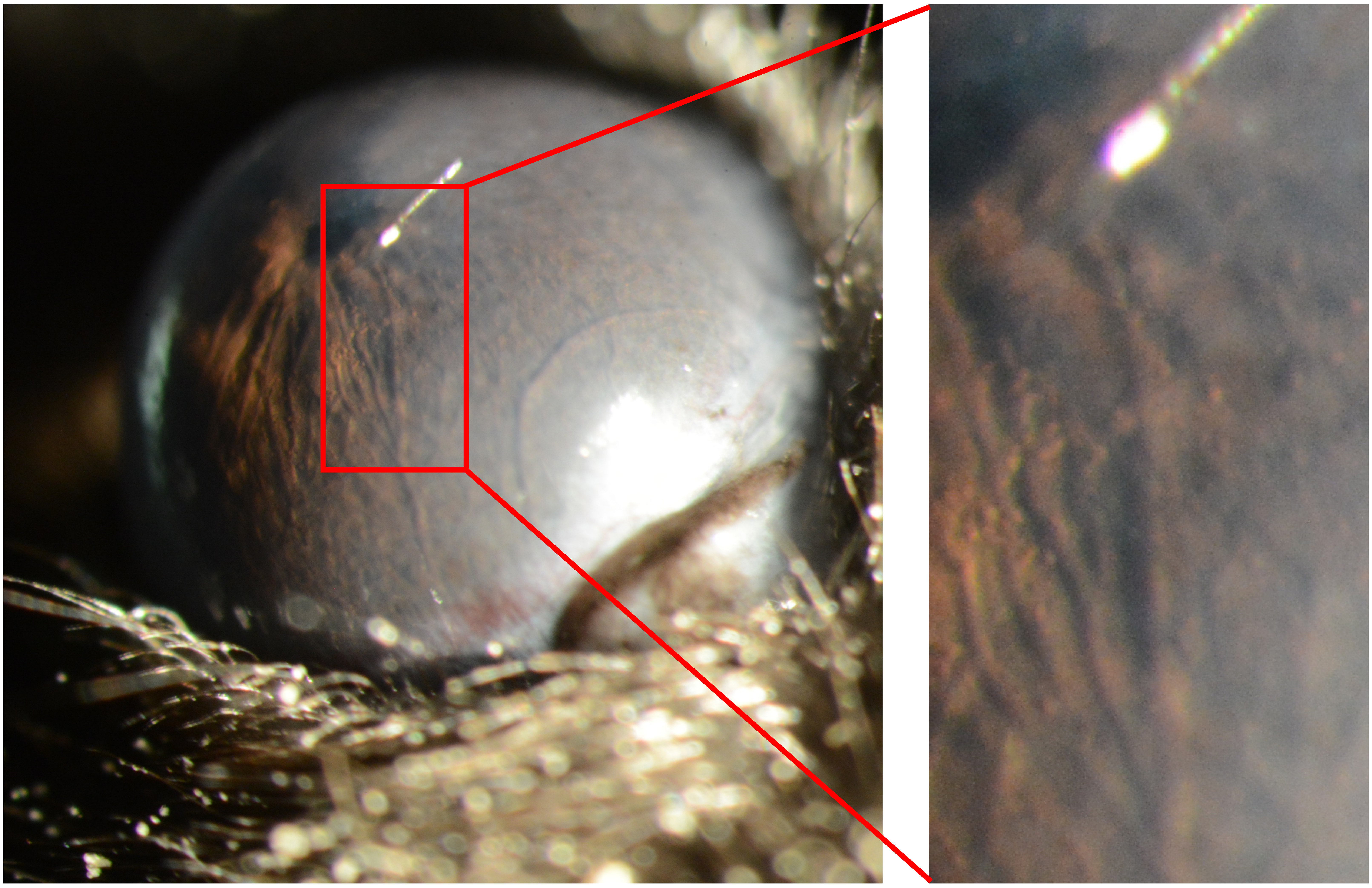

Appendix 4
Mouse ID: 28486
10-week timepoint

Masked assignment: **Virus**
Actual group: **Virus** (OD injected)

**OS**

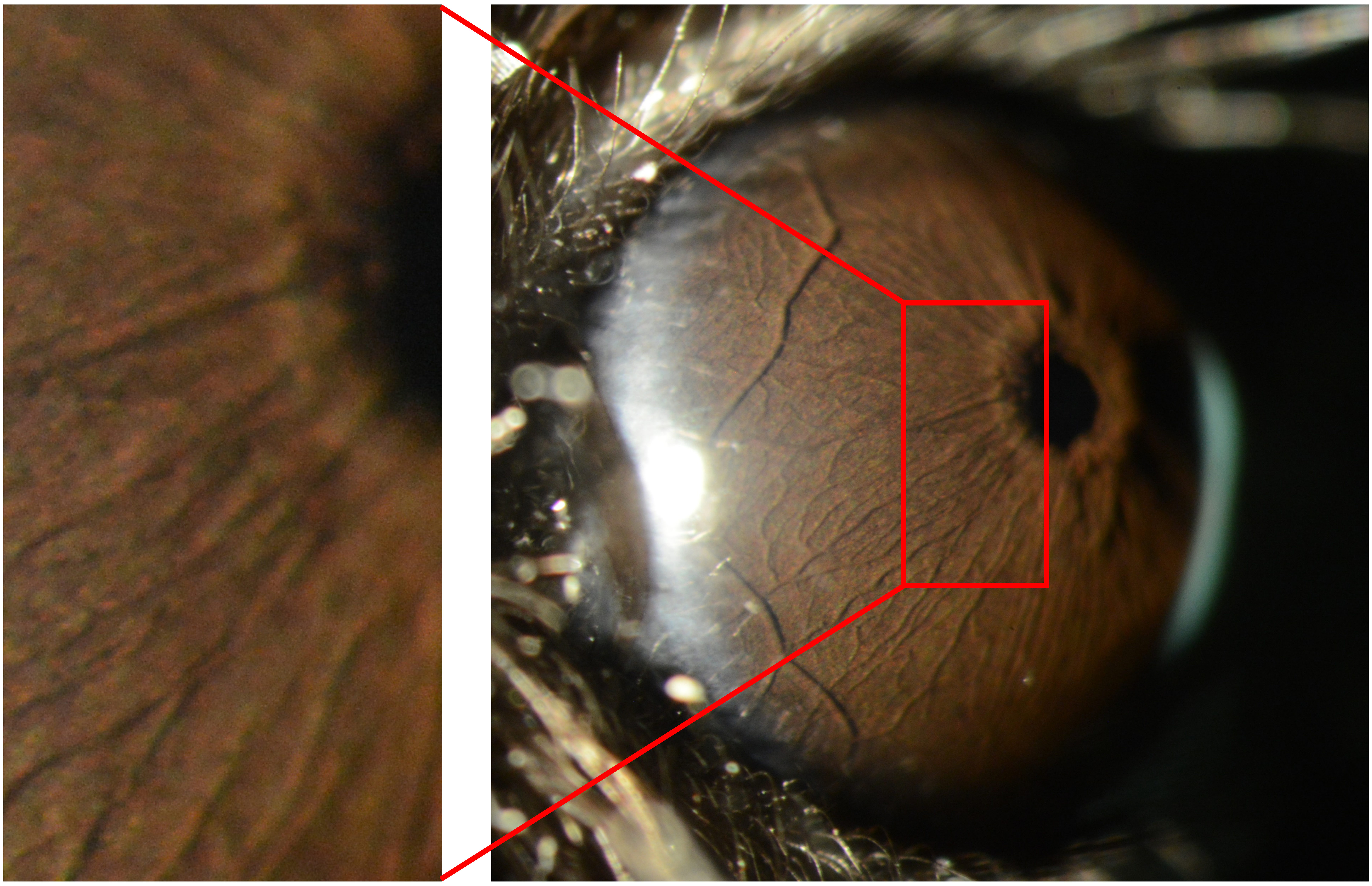

**OD**

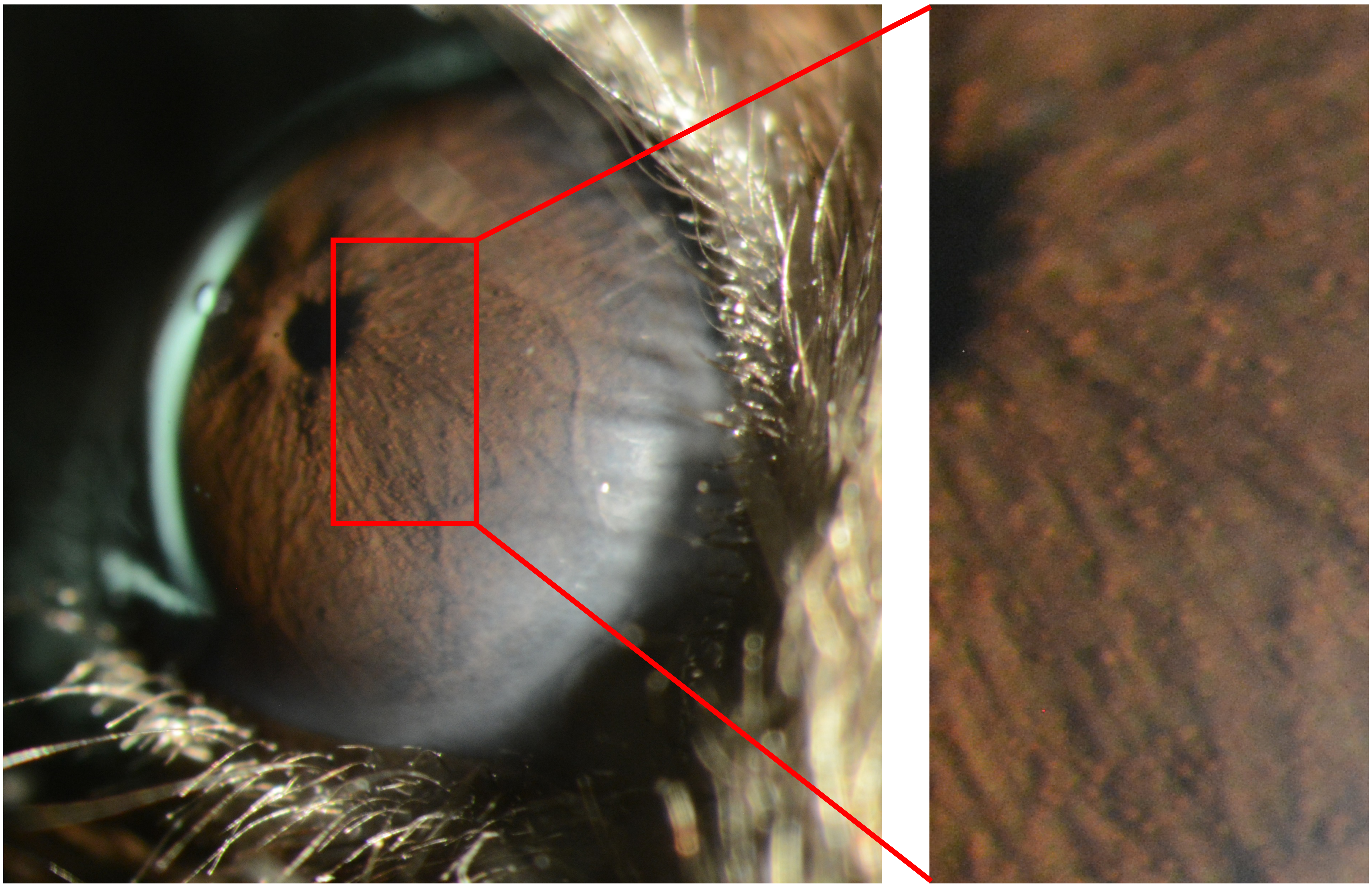
