## Appendix 5 saline 25X for "Recombinant adenovirus causes prolonged mobilization of macrophages in the anterior chamber of mice"

Appendix 5
Mouse ID: 28445

Masked assignment: **Saline**
Actual group: **Saline** (OS injected)

**OS** **OD**

**Pre**

**1 wk**

**3 wk**

**10 wk**

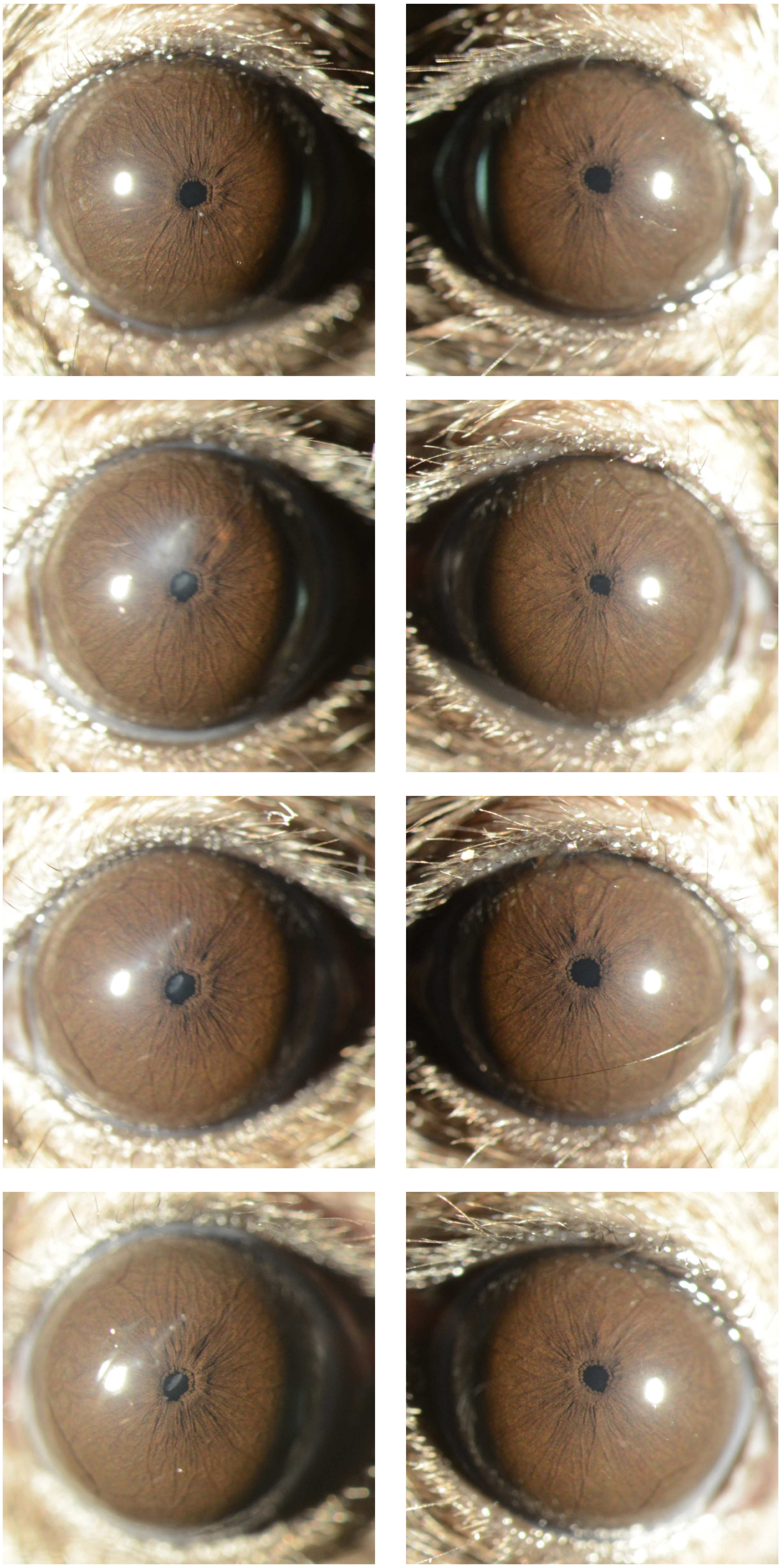

Appendix 5
Mouse ID: 28448

Masked assignment: **Saline**
Actual group: **Saline** (OD injected)

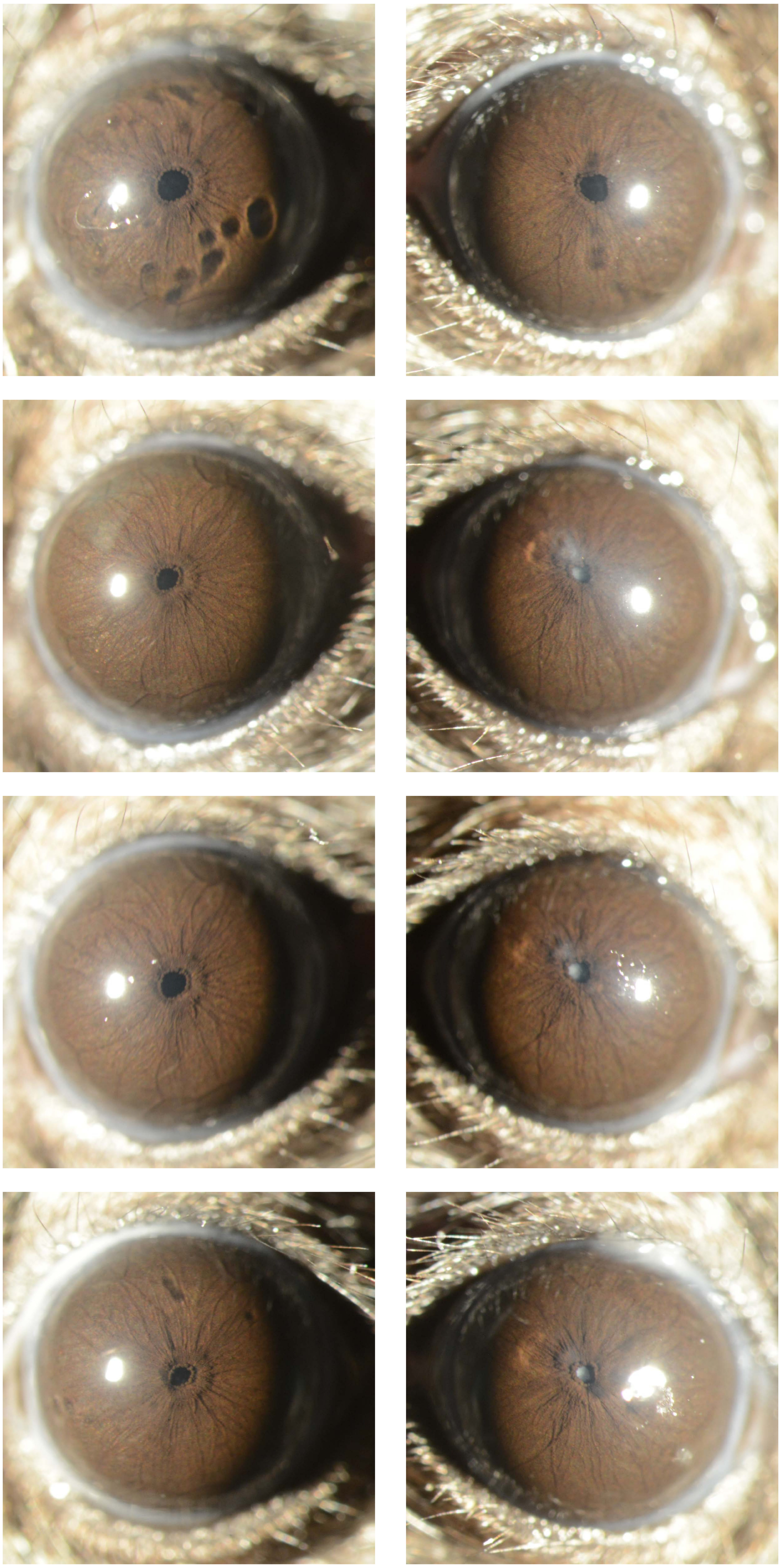

Appendix 5
Mouse ID: 28450

Masked assignment: **Saline**
Actual group: **Saline** (OS injected)

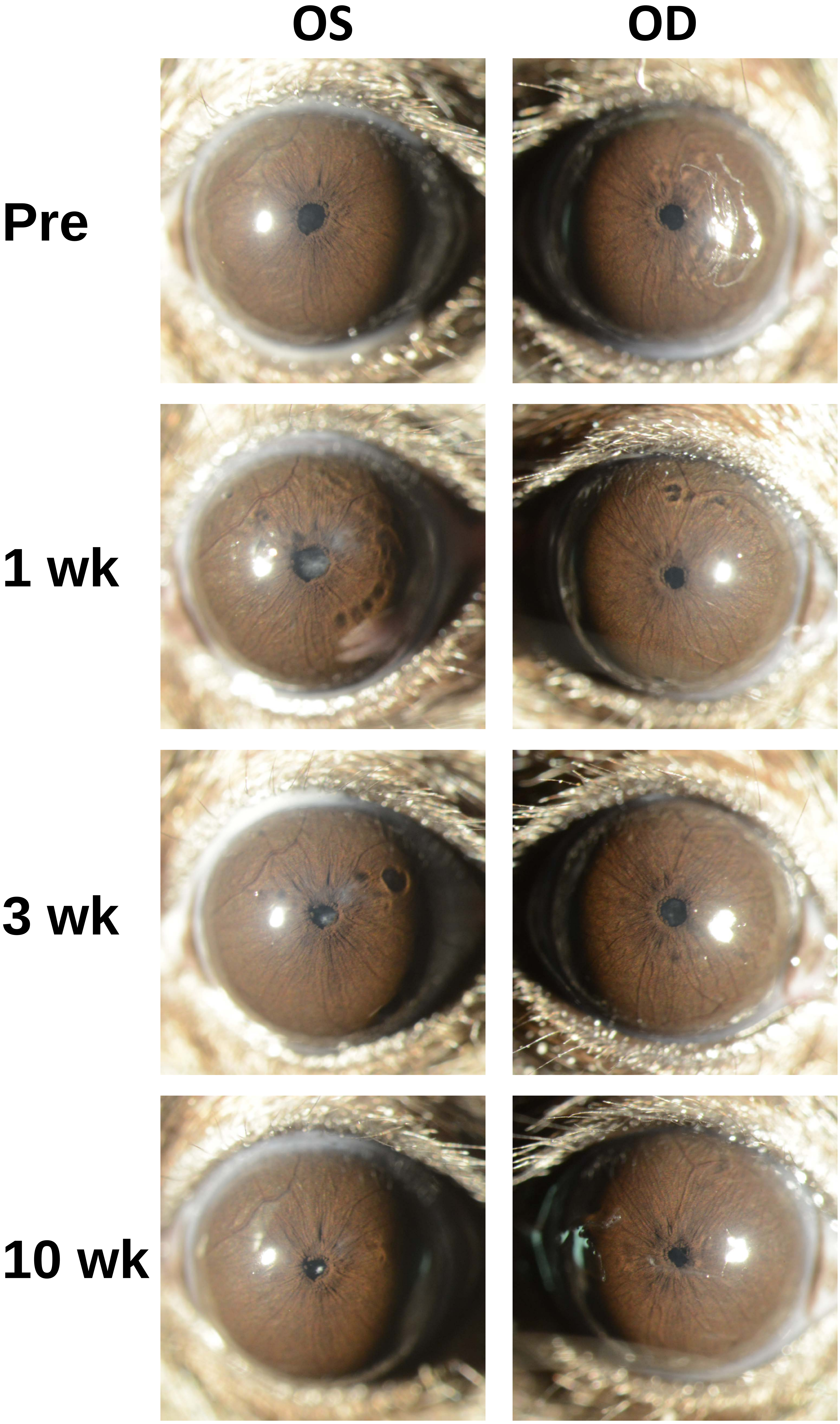

Appendix 5
Mouse ID: 28459

Masked assignment: **Saline**
Actual group: **Saline** (OS injected)

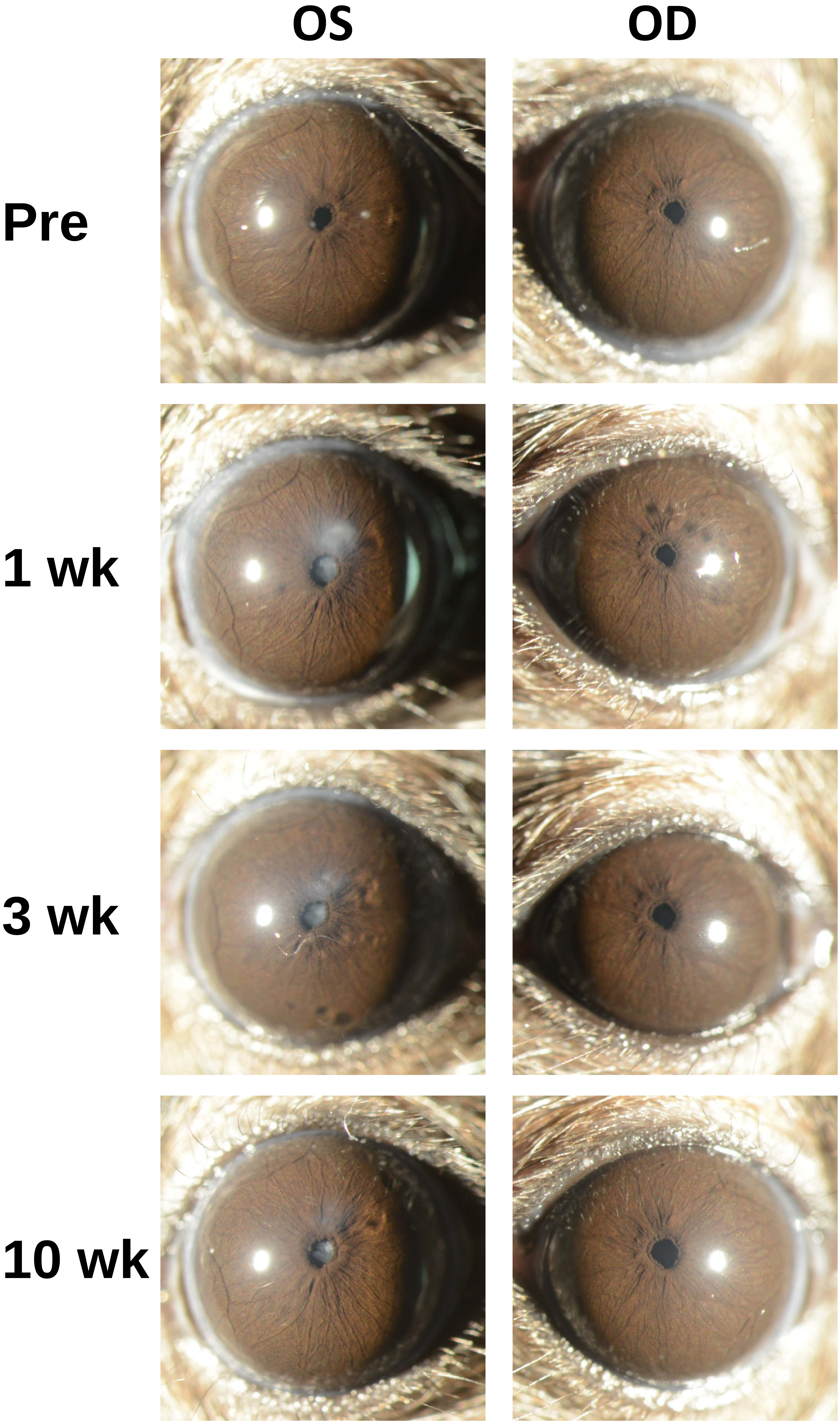

Appendix 5
Mouse ID: 28461

Masked assignment: **Saline**
Actual group: **Saline** (OS injected)

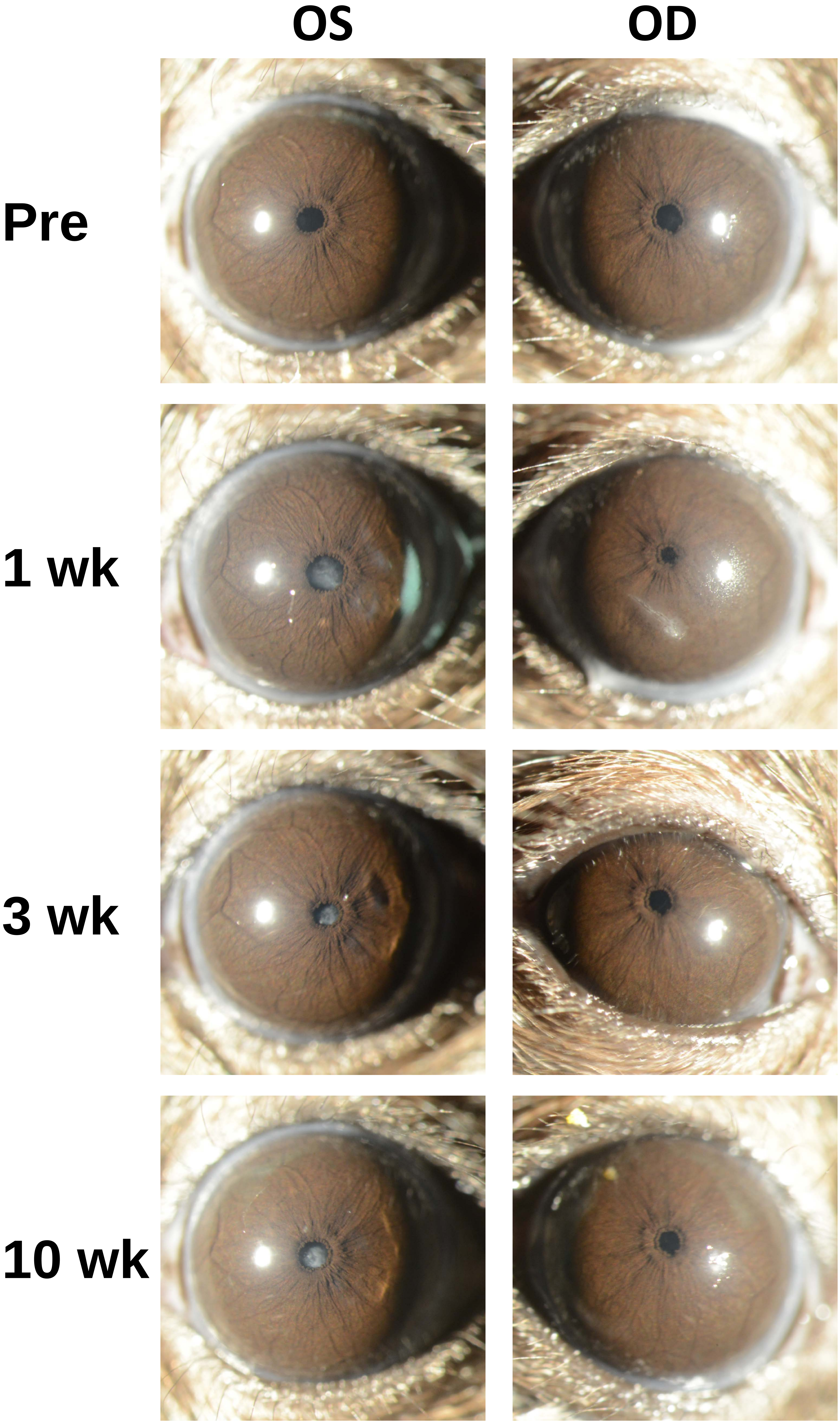

Appendix 5
Mouse ID: 28464

Masked assignment: **Saline**
Actual group: **Saline** (OS injected)

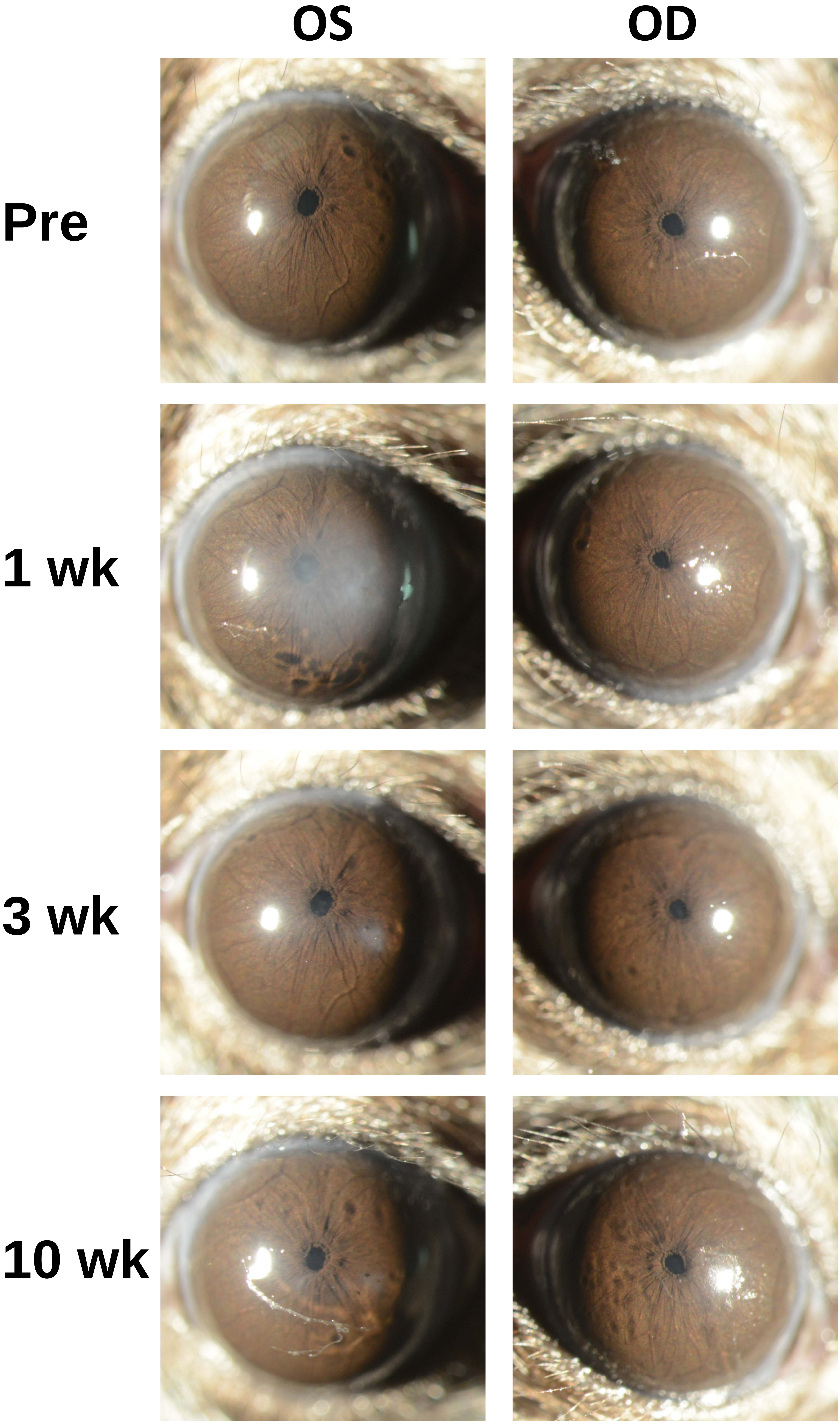

Appendix 5
Mouse ID: 28466

Masked assignment: **Saline**
Actual group: **Saline** (OD injected)

**OS** **OD**

**Pre**

**1 wk**

**3 wk**

**10 wk**

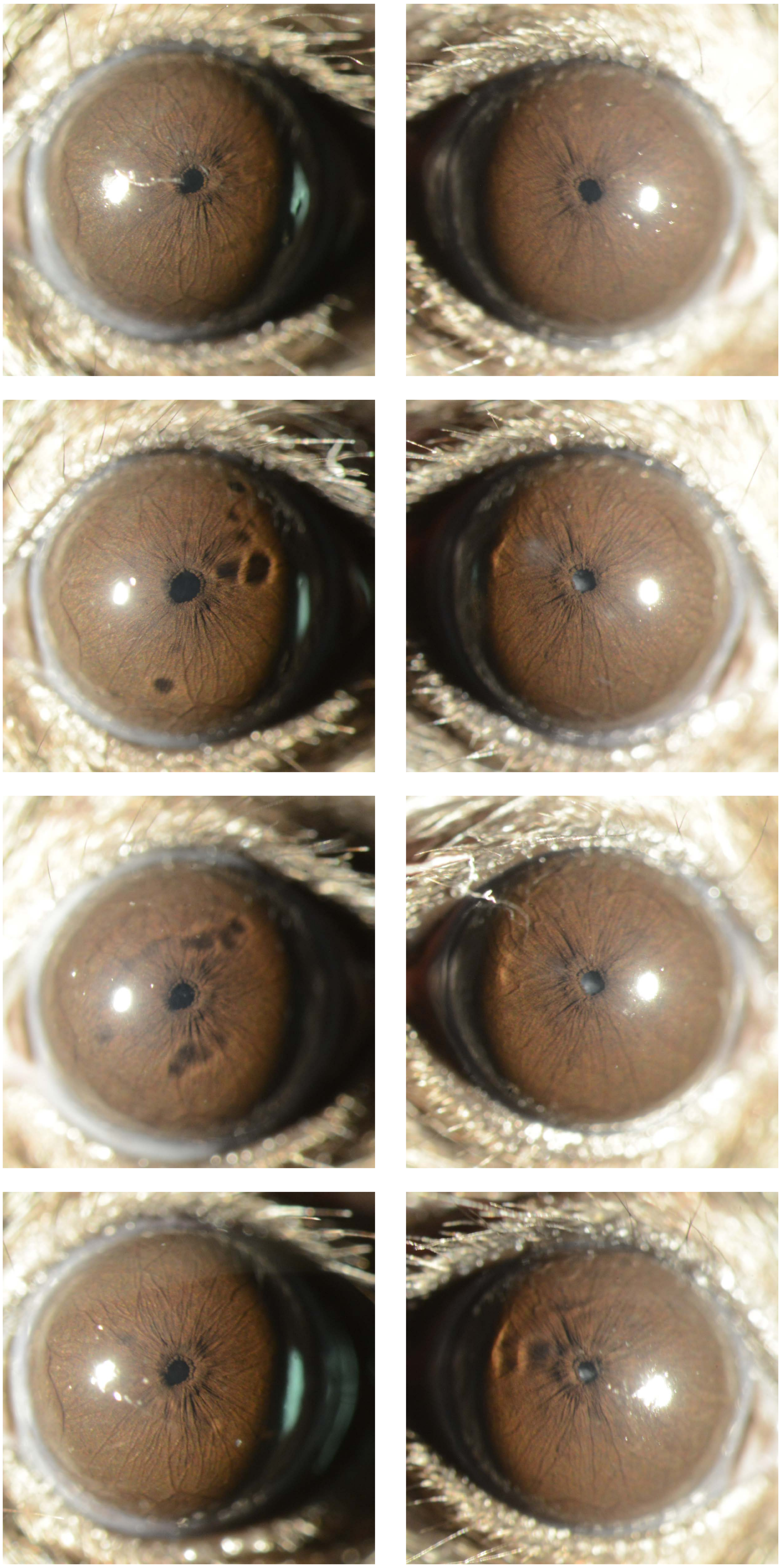

Appendix 5
Mouse ID: 28487

Masked assignment: **Saline**
Actual group: **Saline** (OD injected)

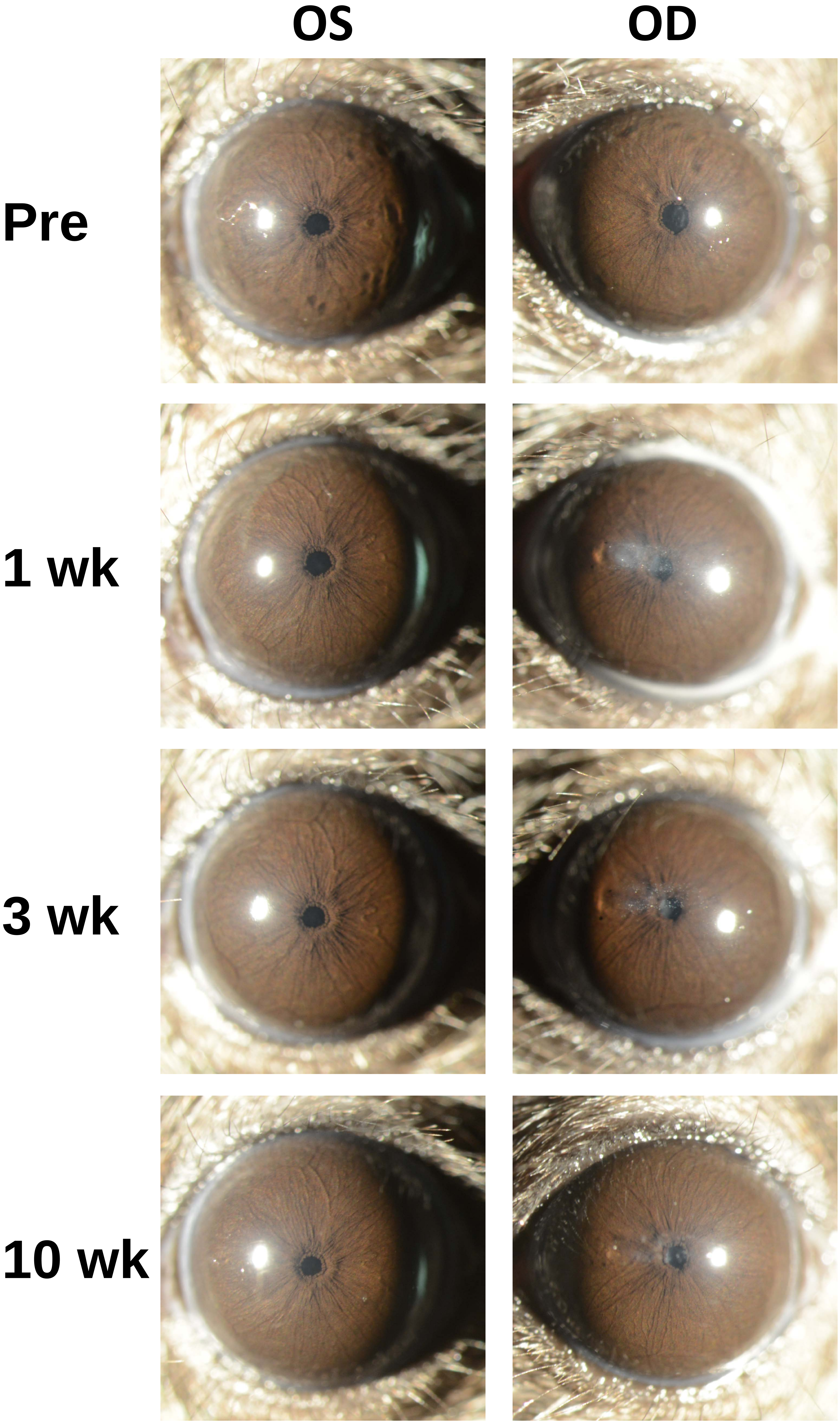

Appendix 5
Mouse ID: 28490

Masked assignment: **Saline**
Actual group: **Saline** (OD injected)

| | OS | OD |
|---|---|---|
| **Pre** | | |
| **1 wk** | | |
| **3 wk** | | |
| **10 wk** | | |

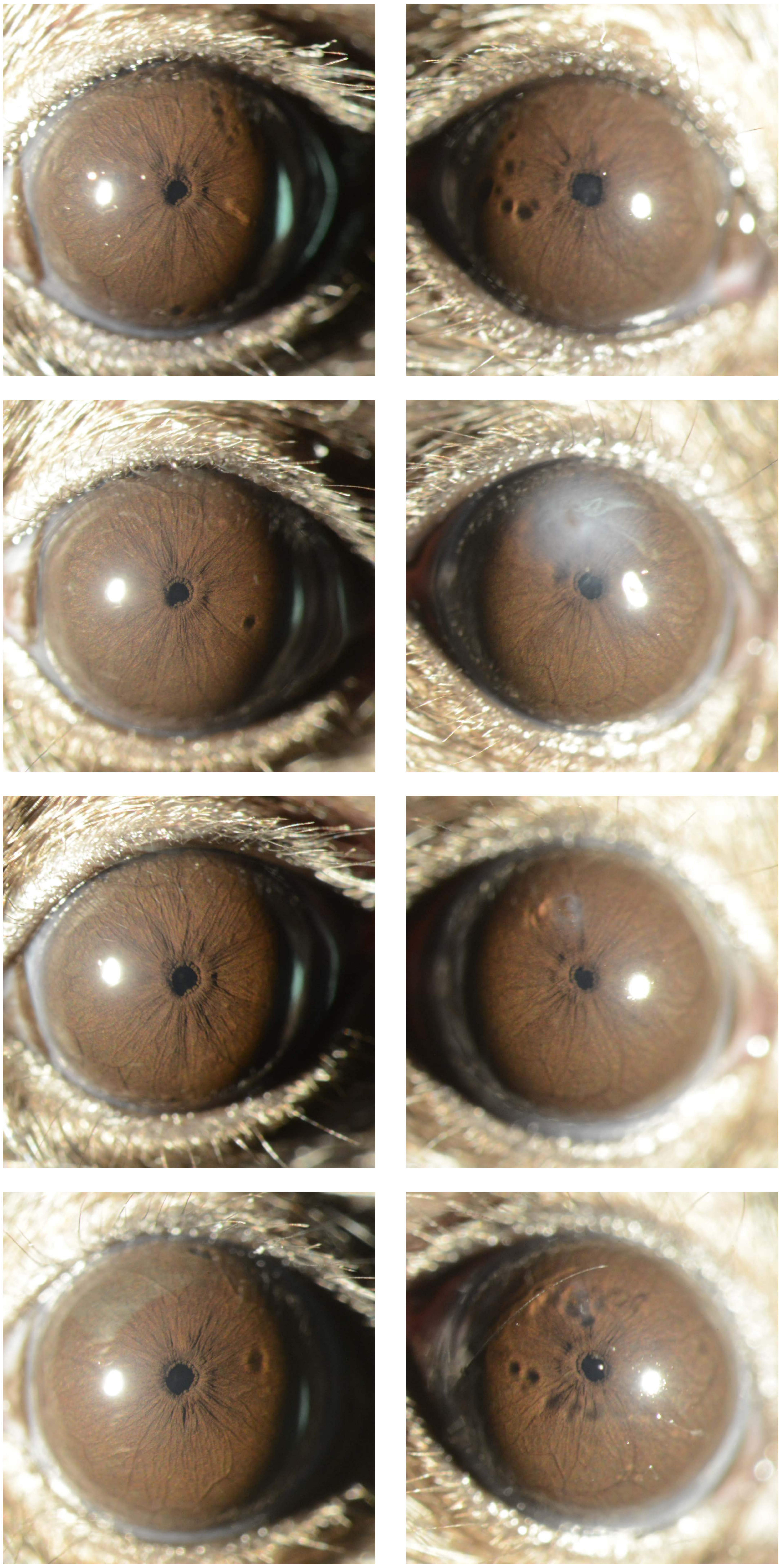
