## Appendix 6 saline 40X for "Recombinant adenovirus causes prolonged mobilization of macrophages in the anterior chamber of mice"

Appendix 6
Mouse ID: 28459
10-week timepoint

Masked assignment: **Saline**
Actual group: **Saline** (OS injected)

**OS**

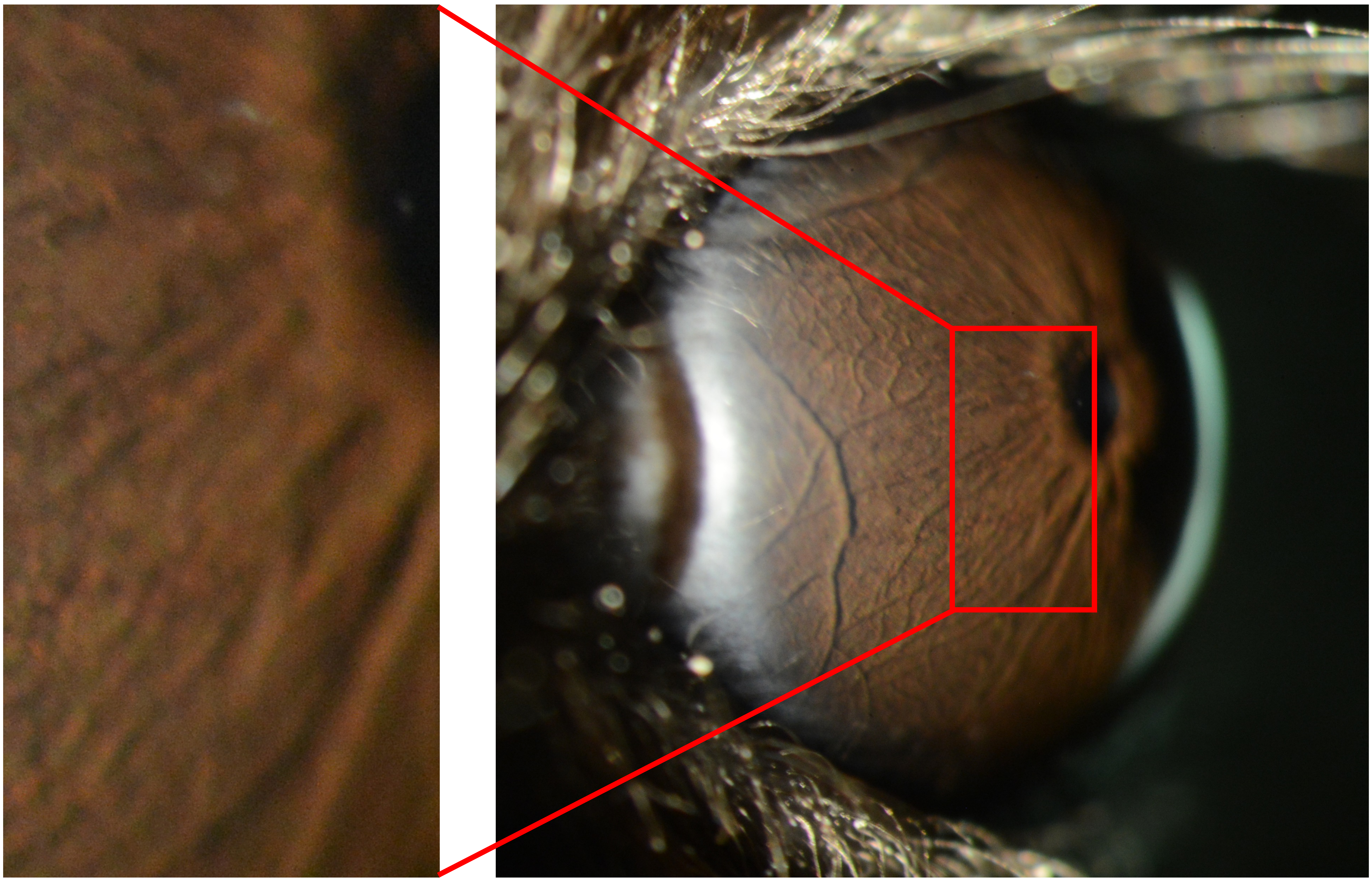

**OD**

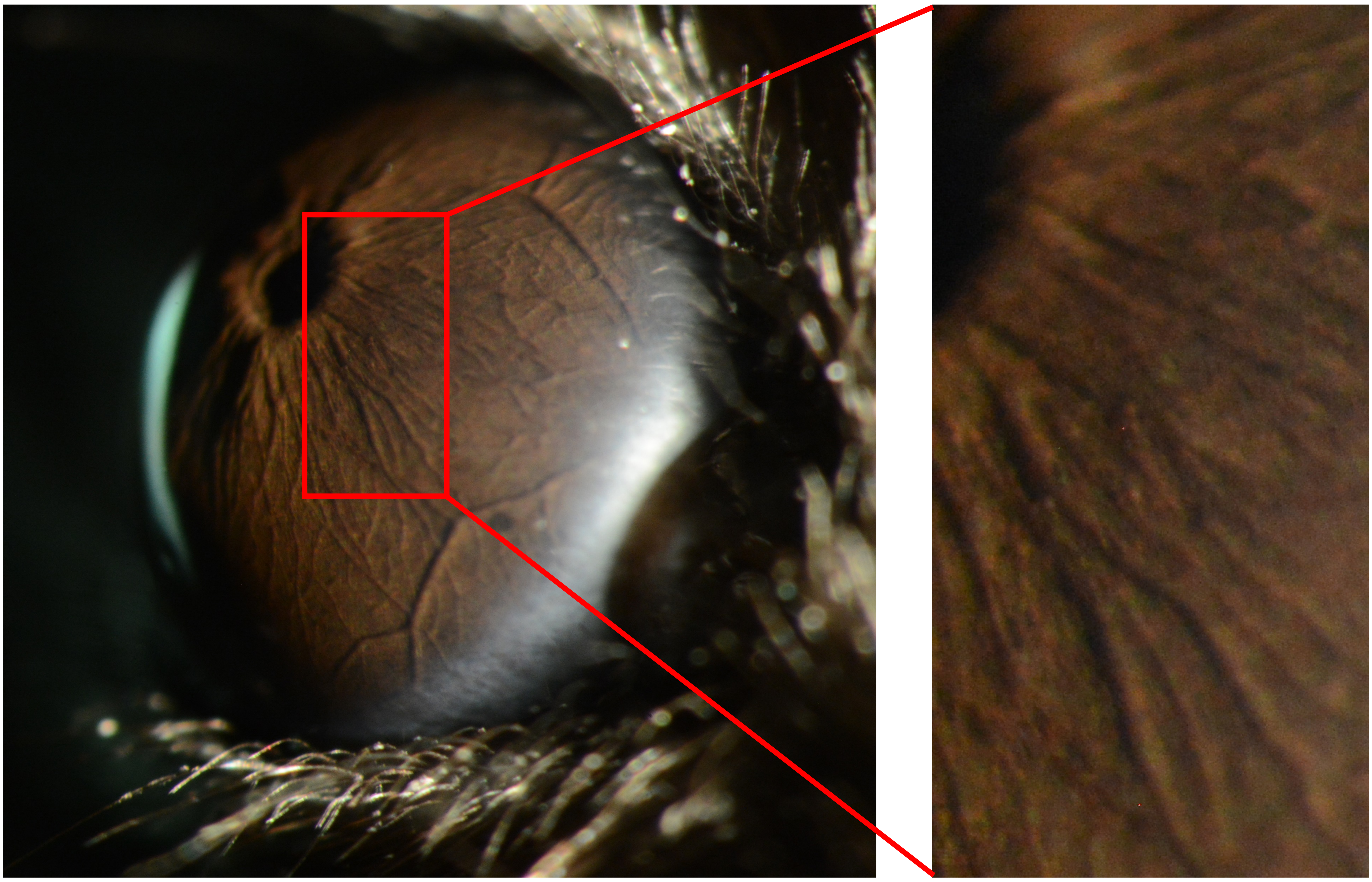

Appendix 6
Mouse ID: 28461
10-week timepoint

Masked assignment: **Saline**
Actual group: **Saline** (OD injected)

**OS**

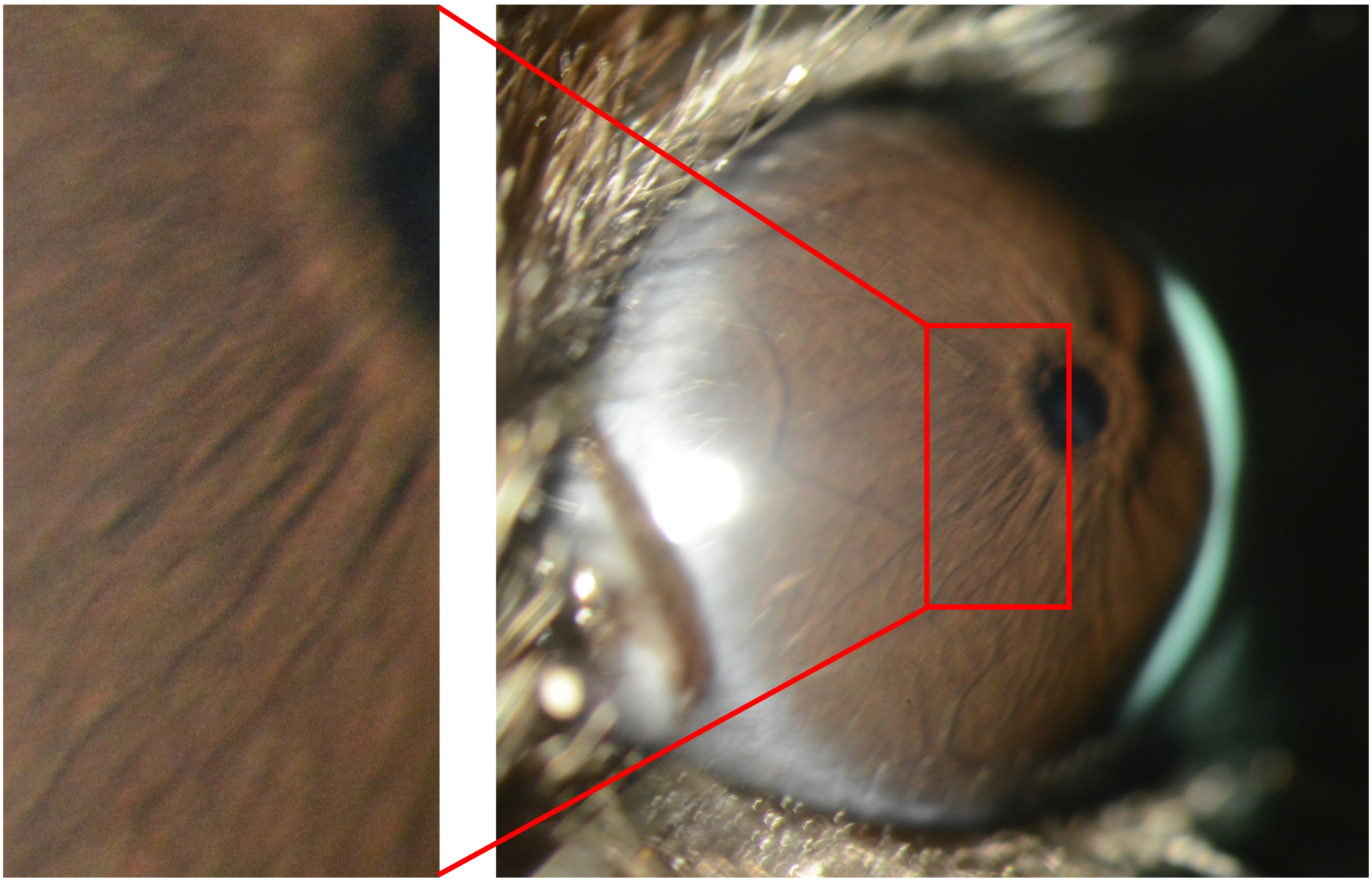

**OD**

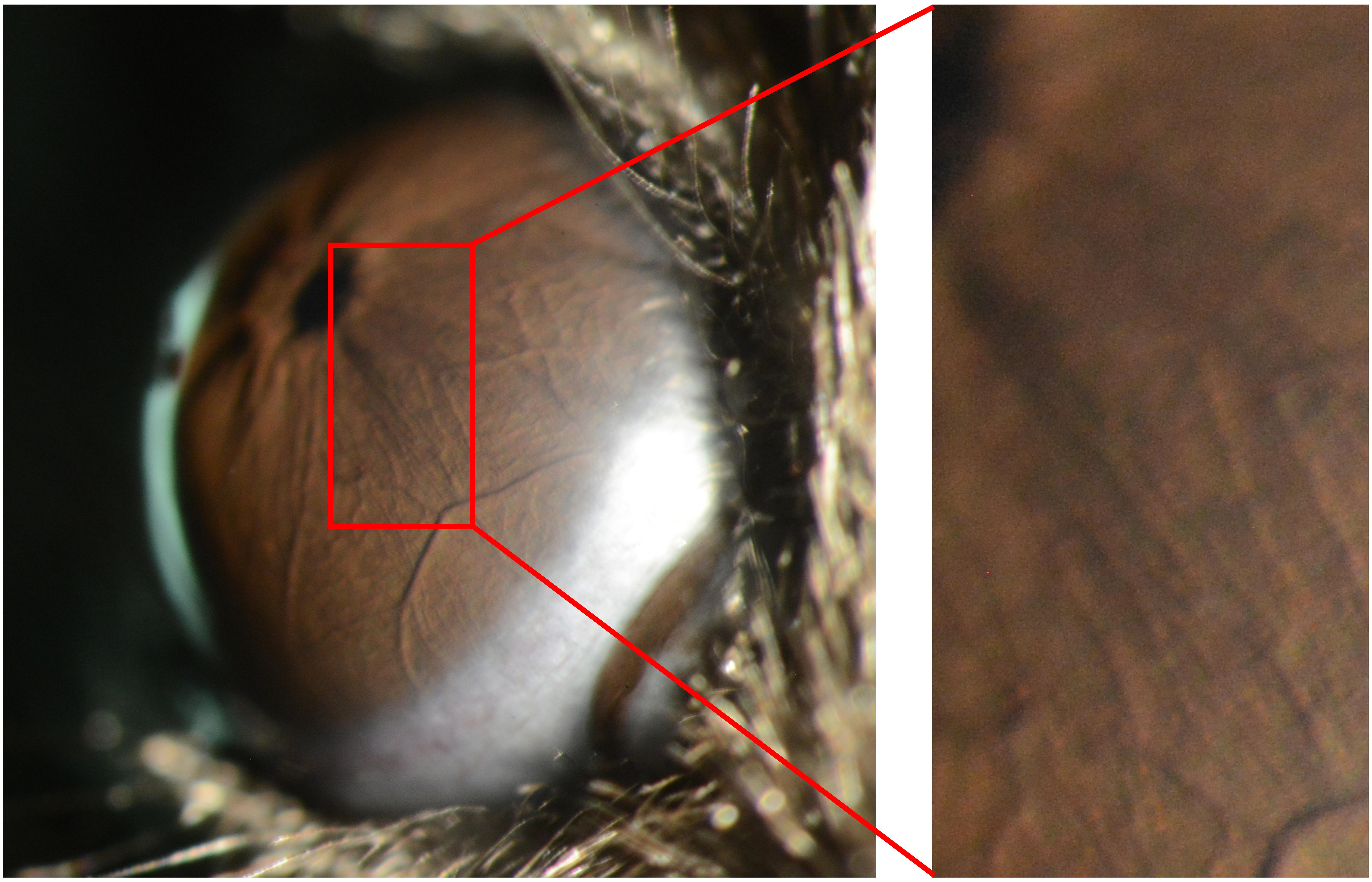

Appendix 6
Mouse ID: 28445
10-week timepoint

Masked assignment: **Saline**
Actual group: **Saline** (OS injected)

**OS**

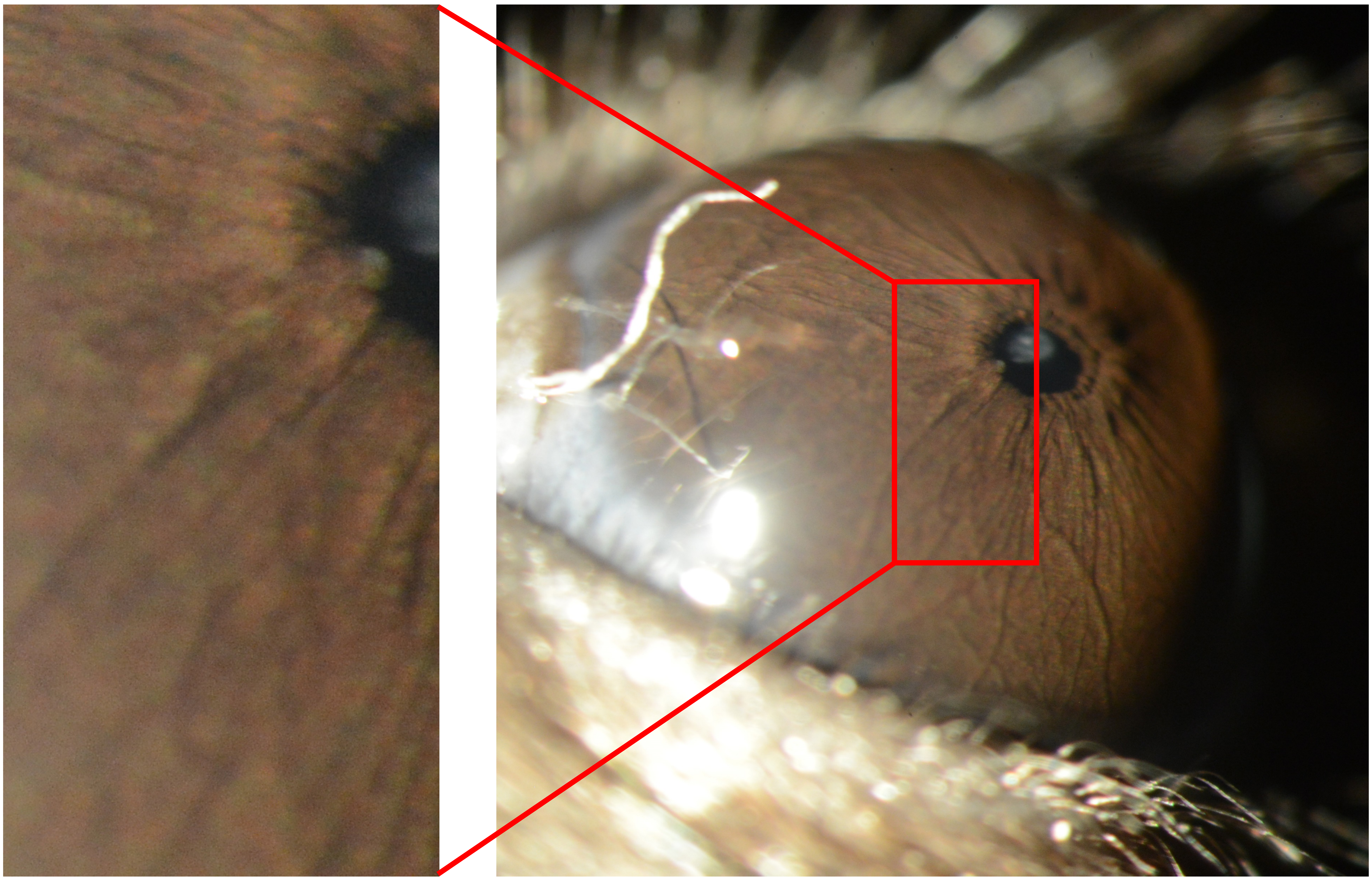

**OD**

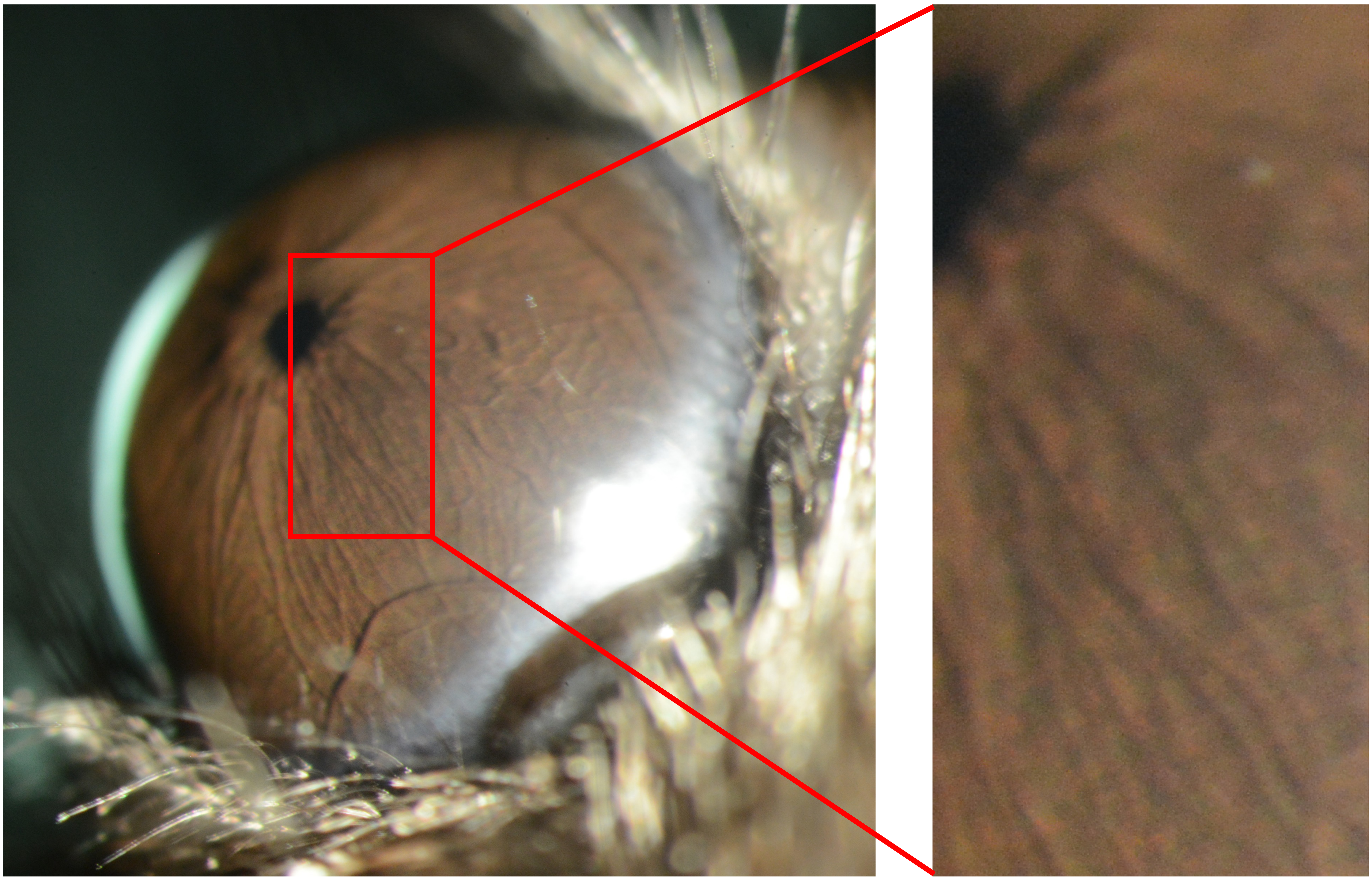

Appendix 6
Mouse ID: 28448
10-week timepoint

Masked assignment: **Saline**
Actual group: **Saline** (OS injected)

**OS**

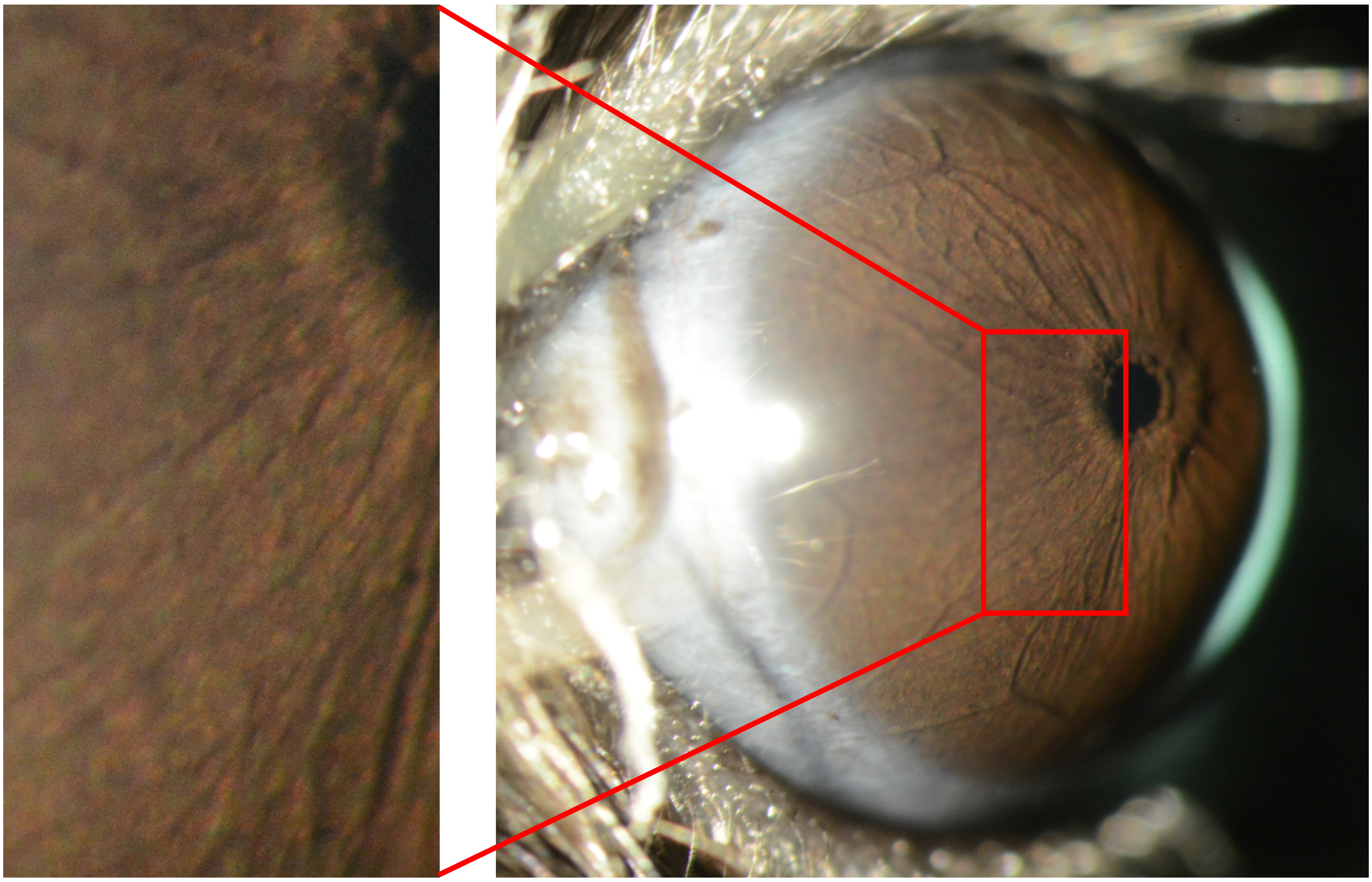

**OD**

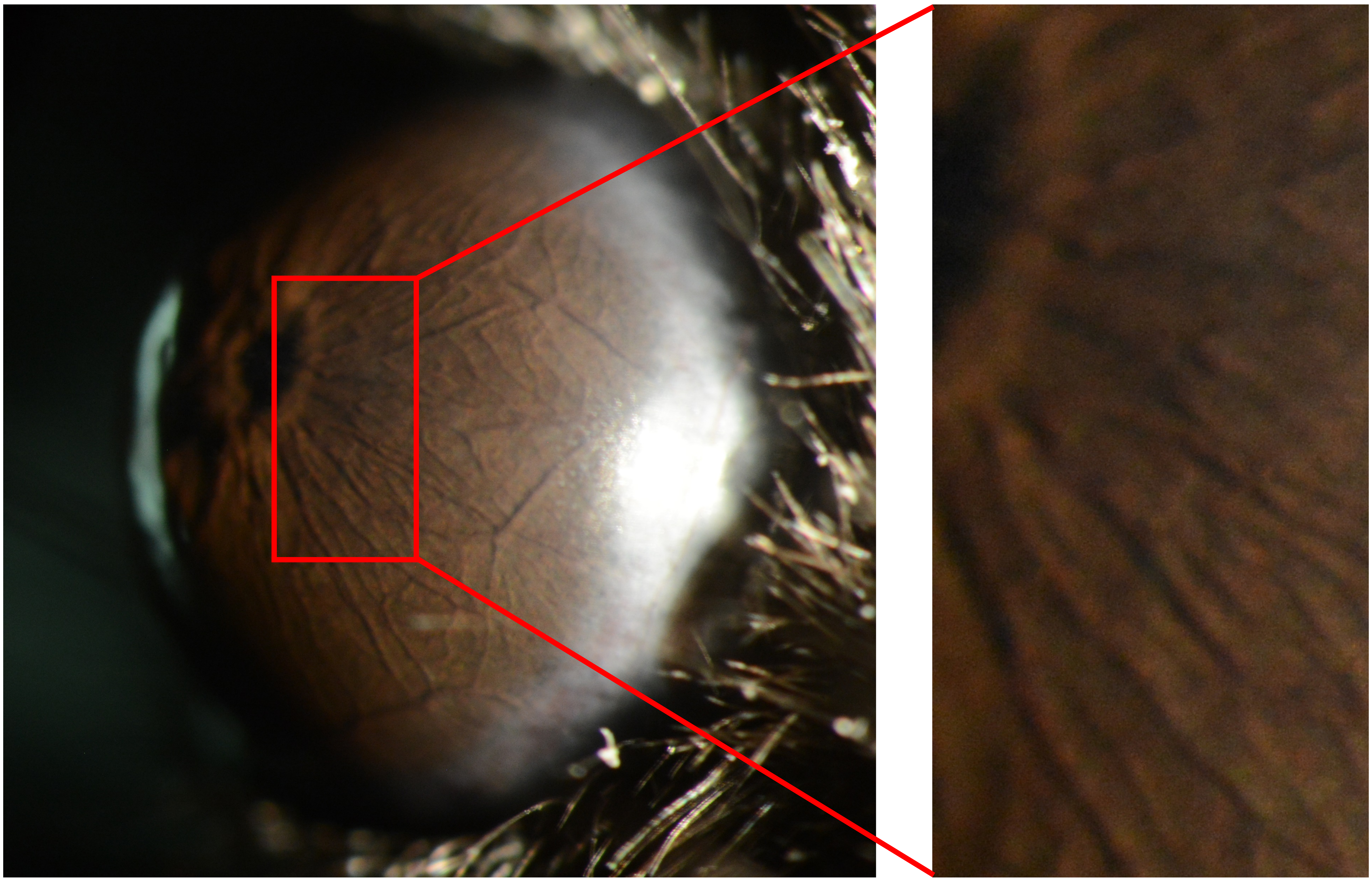

Appendix 6
Mouse ID: 28450
10-week timepoint

Masked assignment: **Saline**
Actual group: **Saline** (OS injected)

**OS**

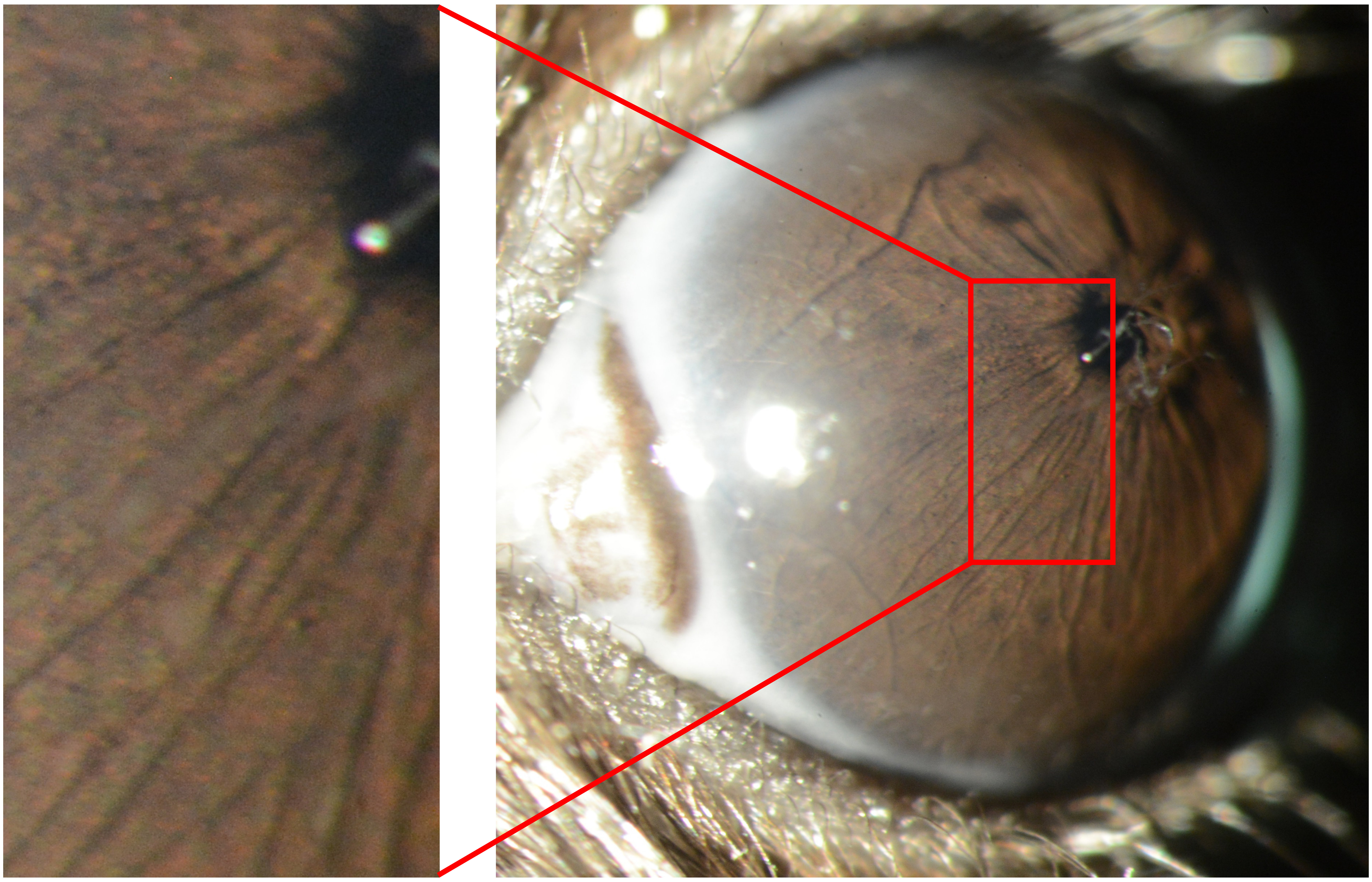

**OD**

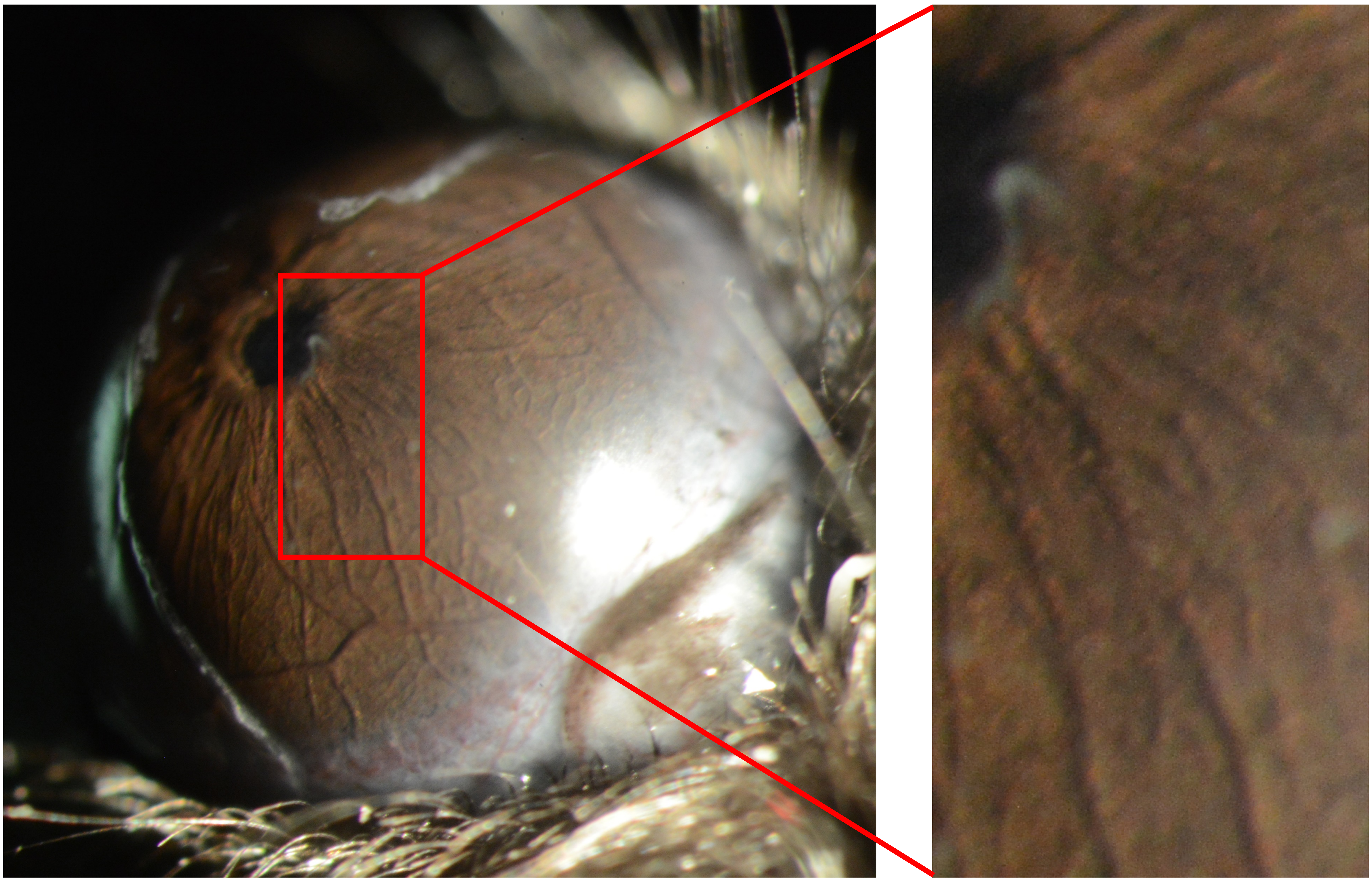

Appendix 6
Mouse ID: 28464
10-week timepoint

Masked assignment: **Saline**
Actual group: **Saline** (OS injected)

**OS**

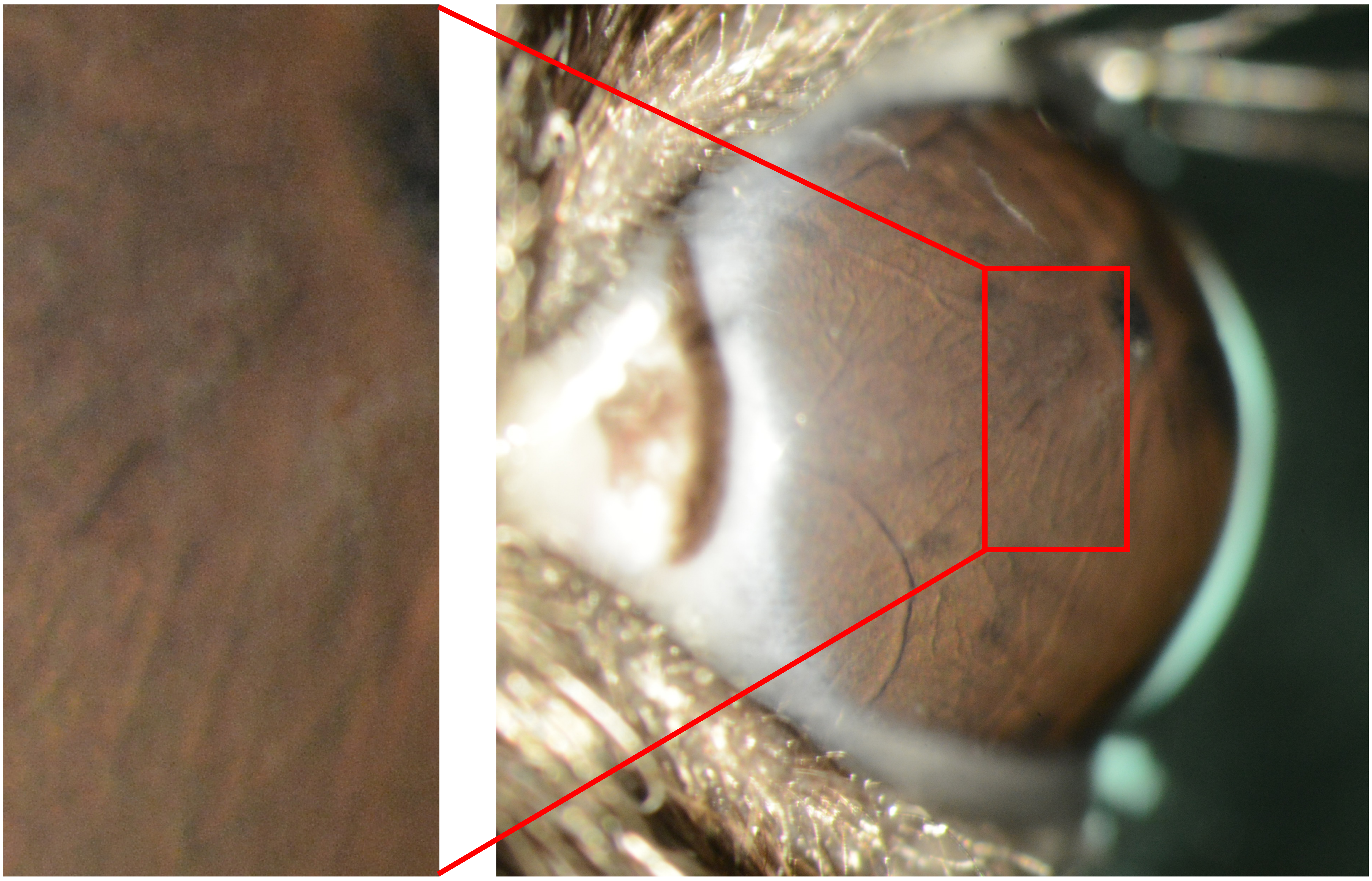

**OD**

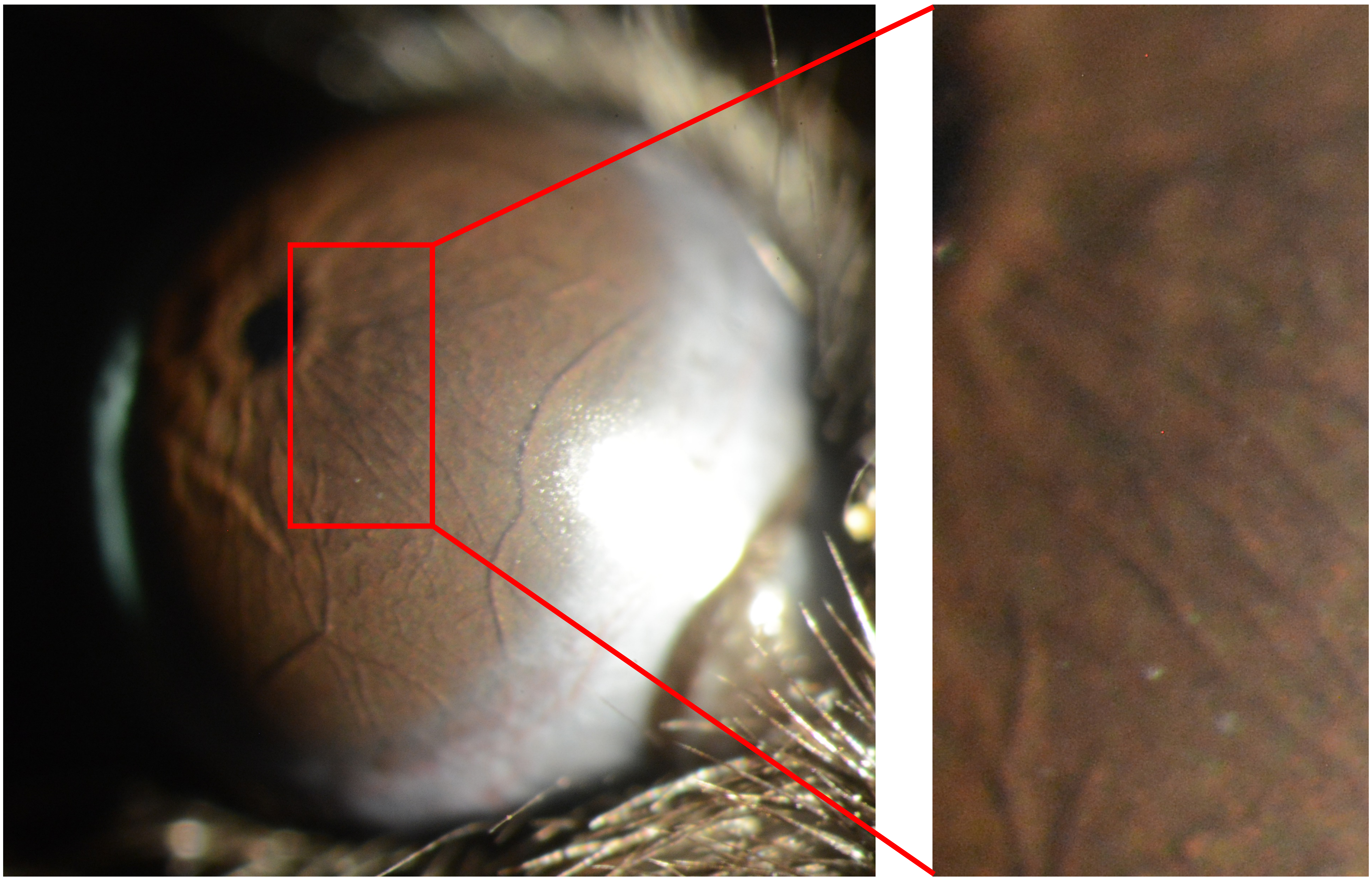

Appendix 6
Mouse ID: 28466
10-week timepoint

Masked assignment: **Saline**
Actual group: **Saline** (OD injected)

**OS**

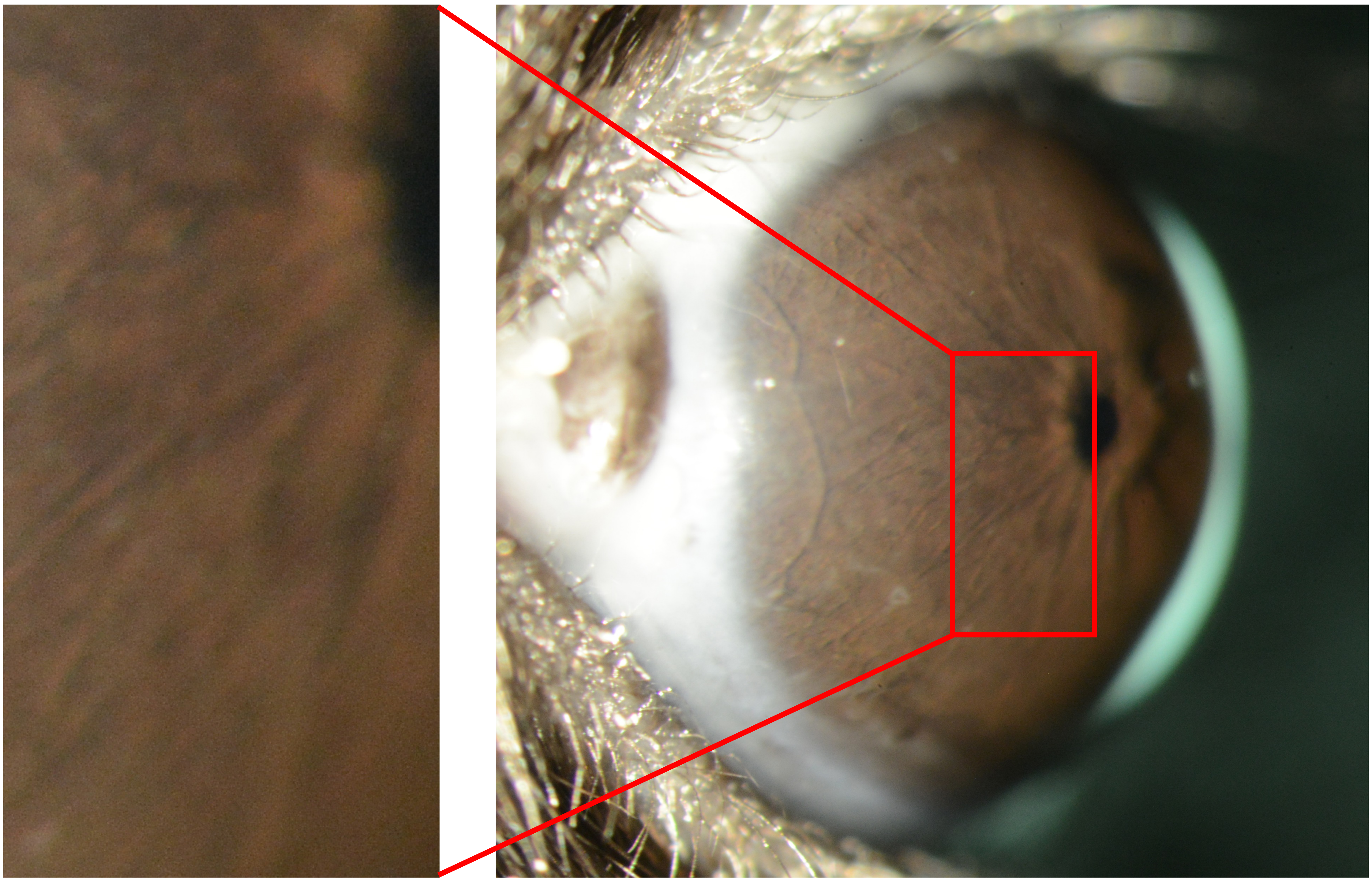

**OD**

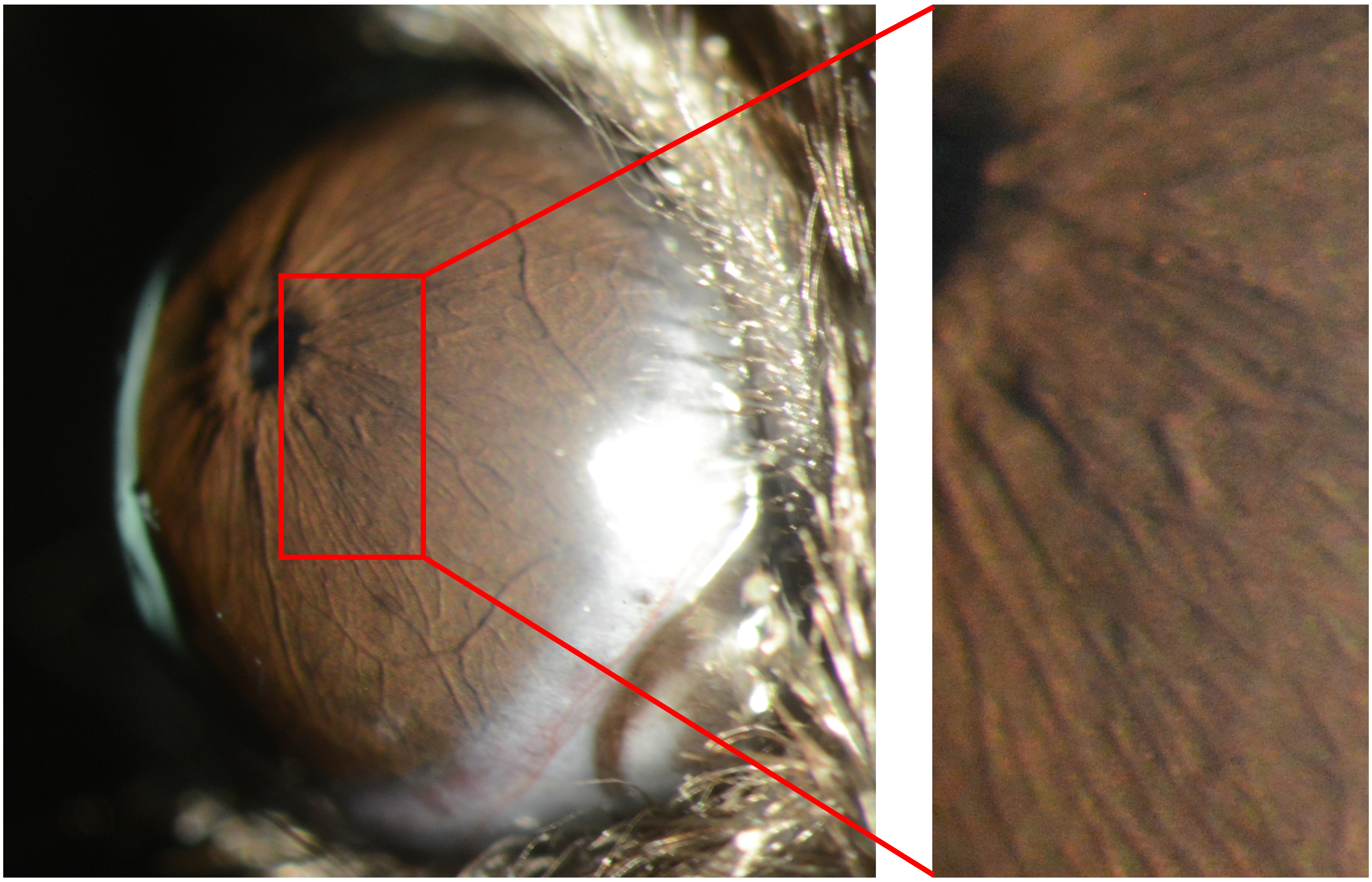

Appendix 6
Mouse ID: 28487
10-week timepoint

Masked assignment: **Saline**
Actual group: **Saline** (OD injected)

**OS**

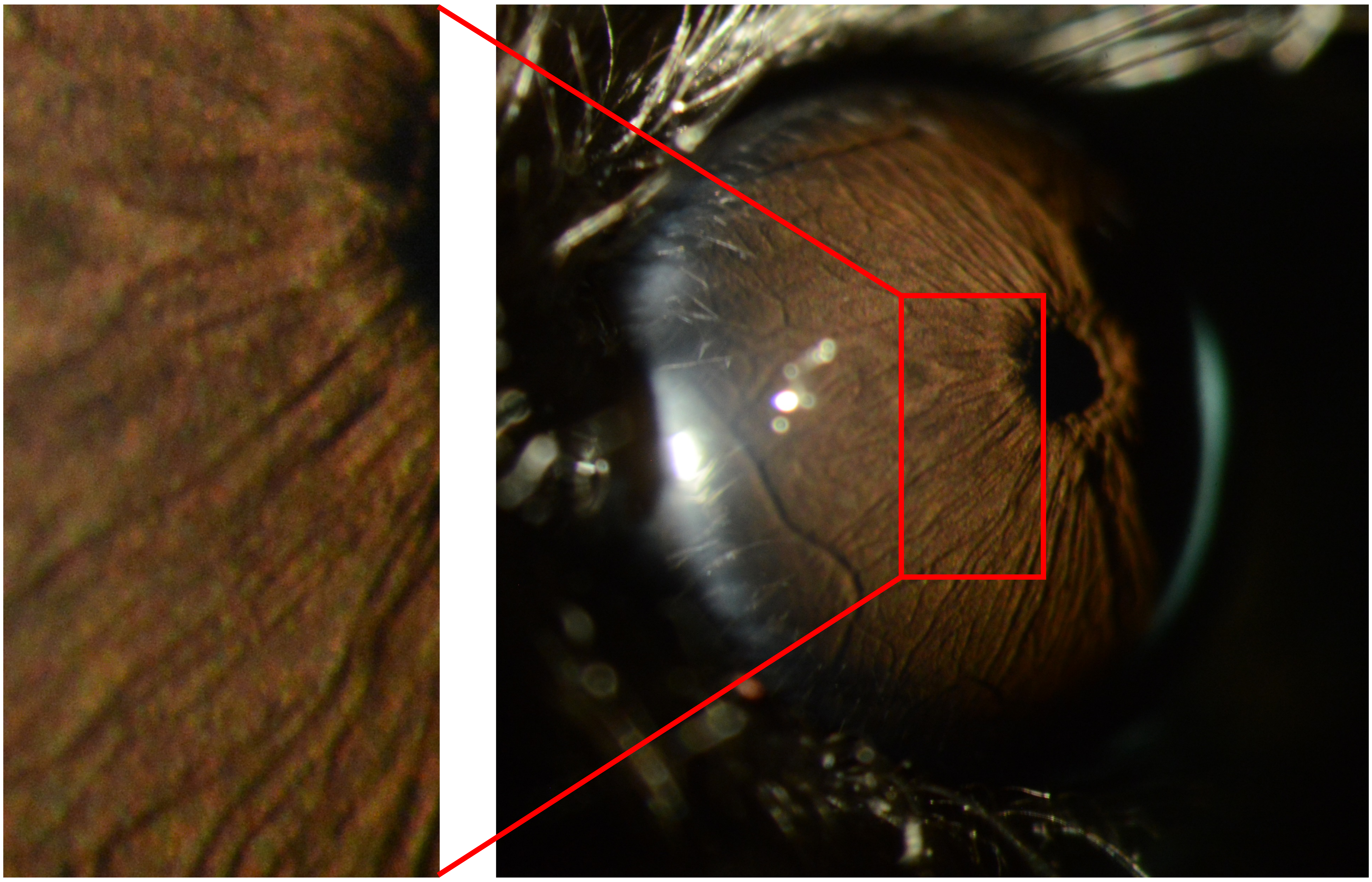

**OD**

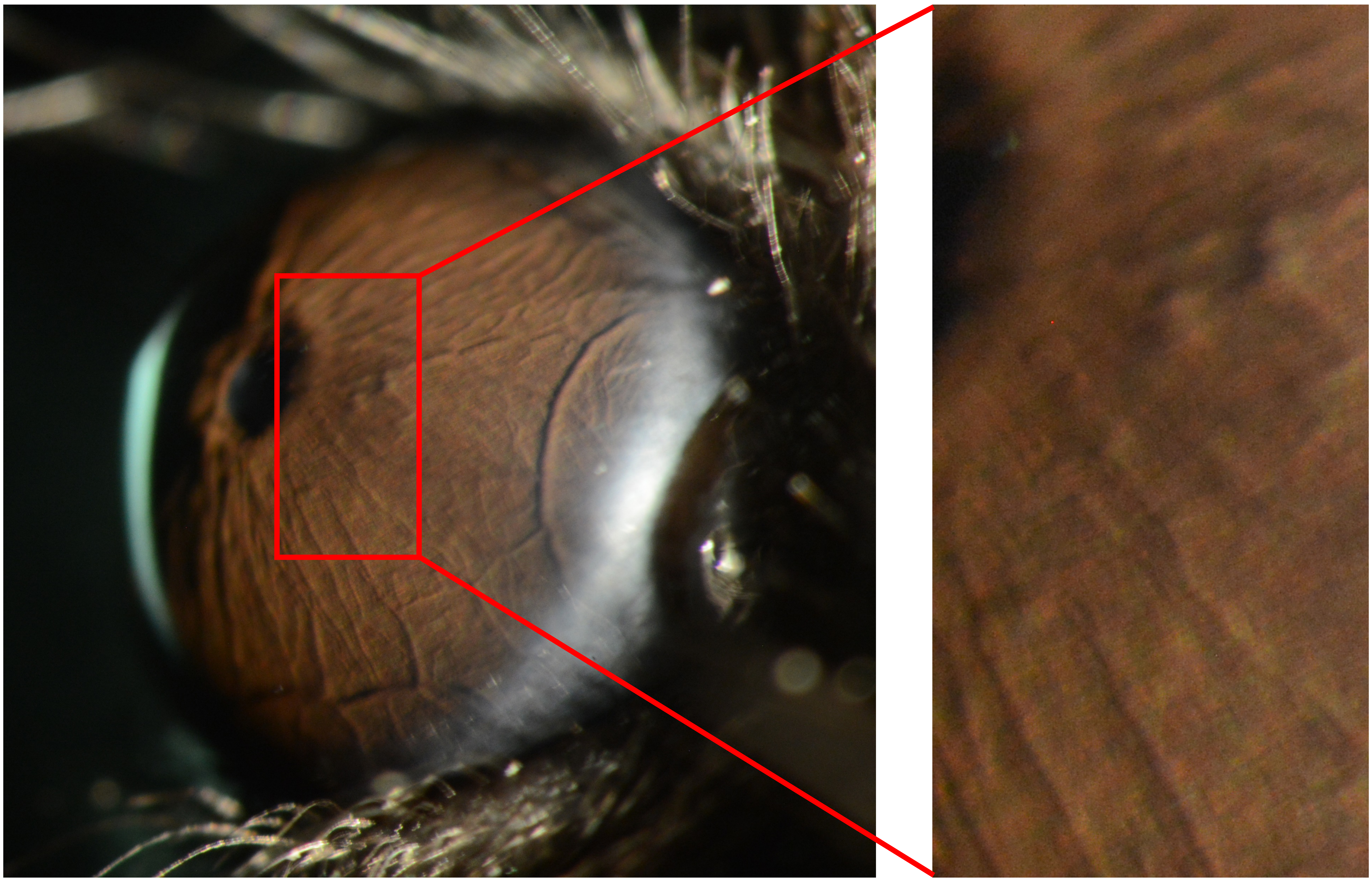

Appendix 6
Mouse ID: 28490
10-week timepoint

Masked assignment: **Saline**
Actual group: **Saline** (OD injected)

**OS**

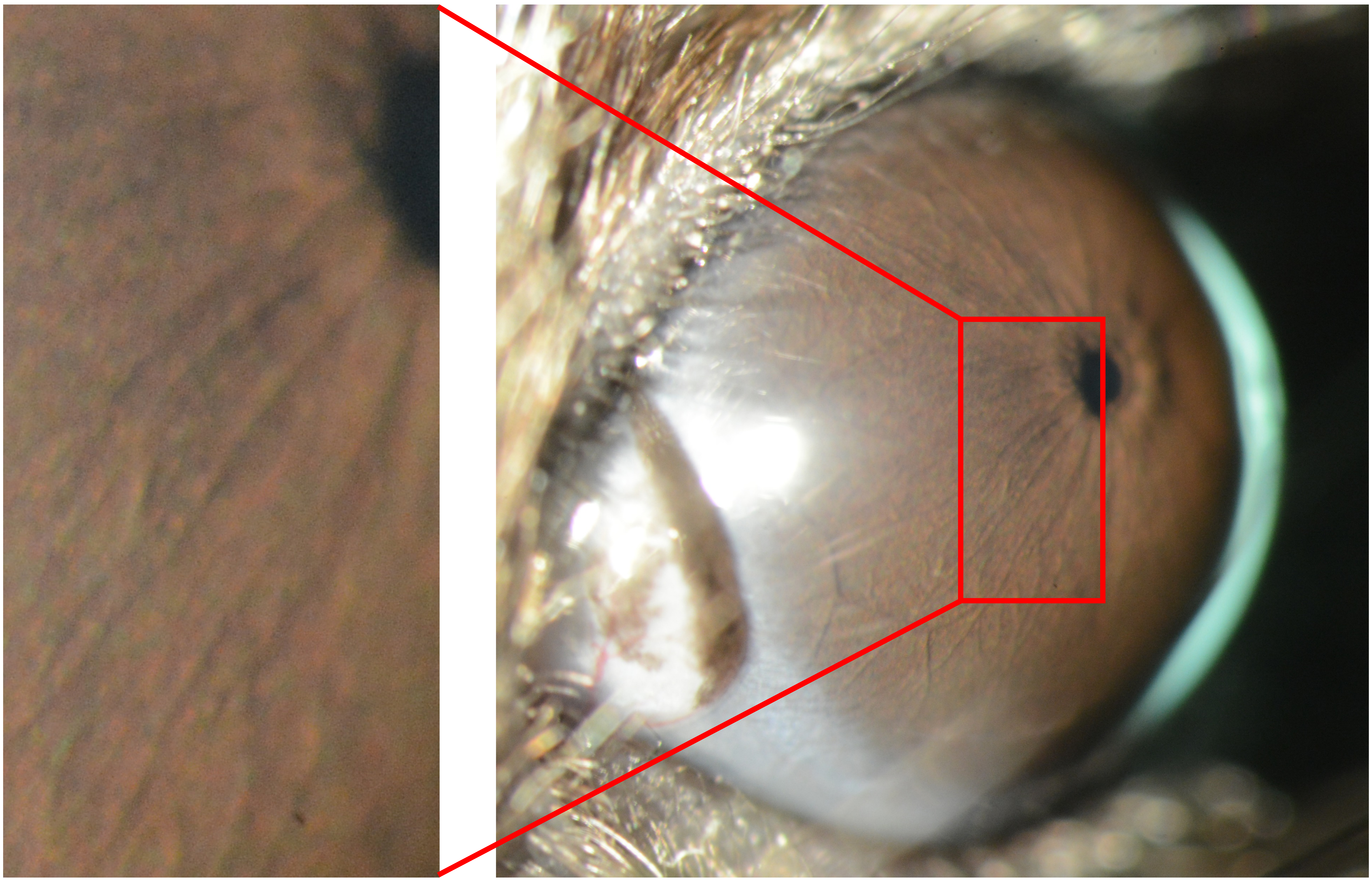

**OD**

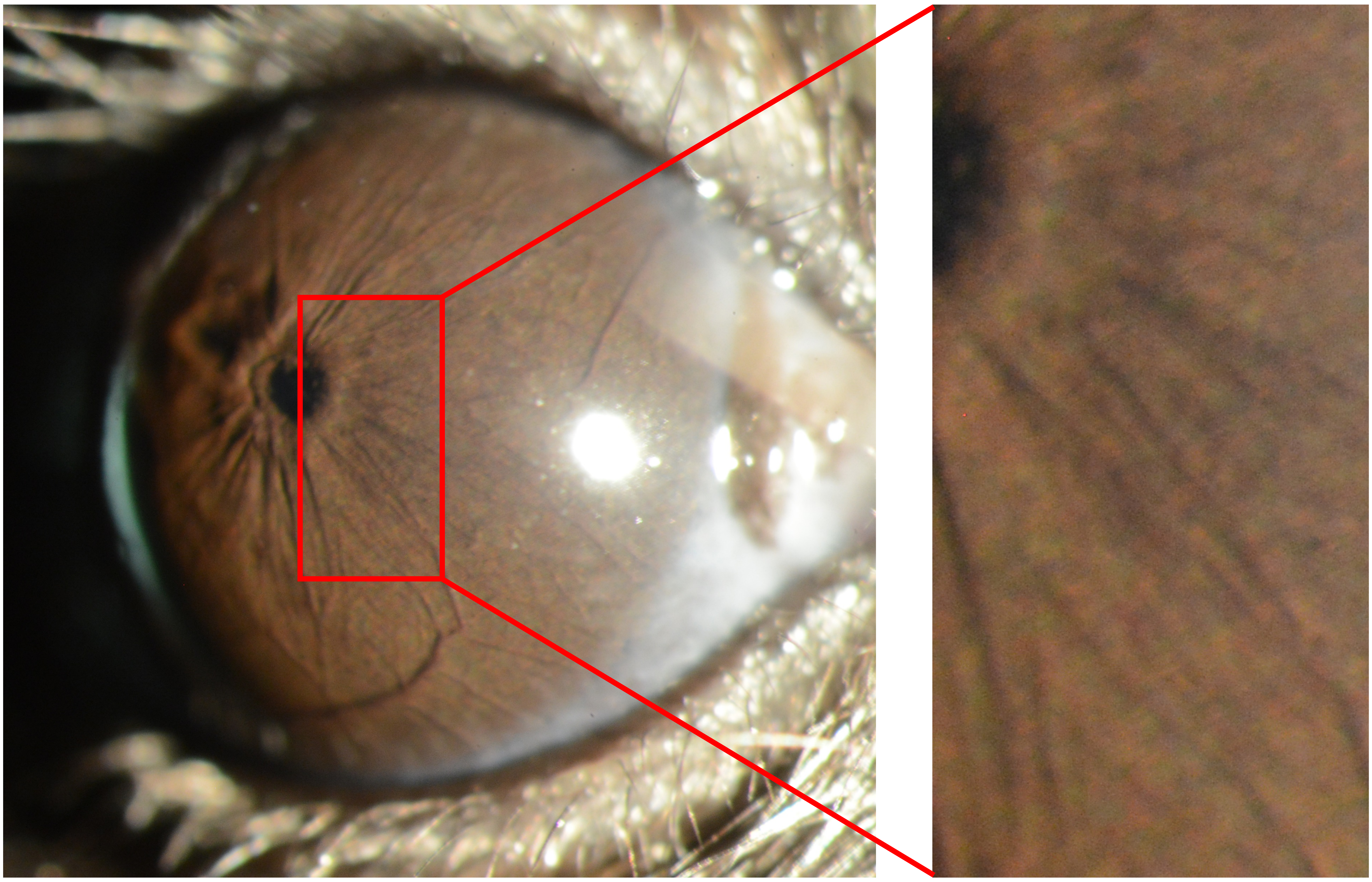
