## Appendix 7 naive 25X for "Recombinant adenovirus causes prolonged mobilization of macrophages in the anterior chamber of mice"

Appendix 7
Mouse ID: 28443

Masked assignment: **Naïve**
Actual group: **Saline** (OS injected)

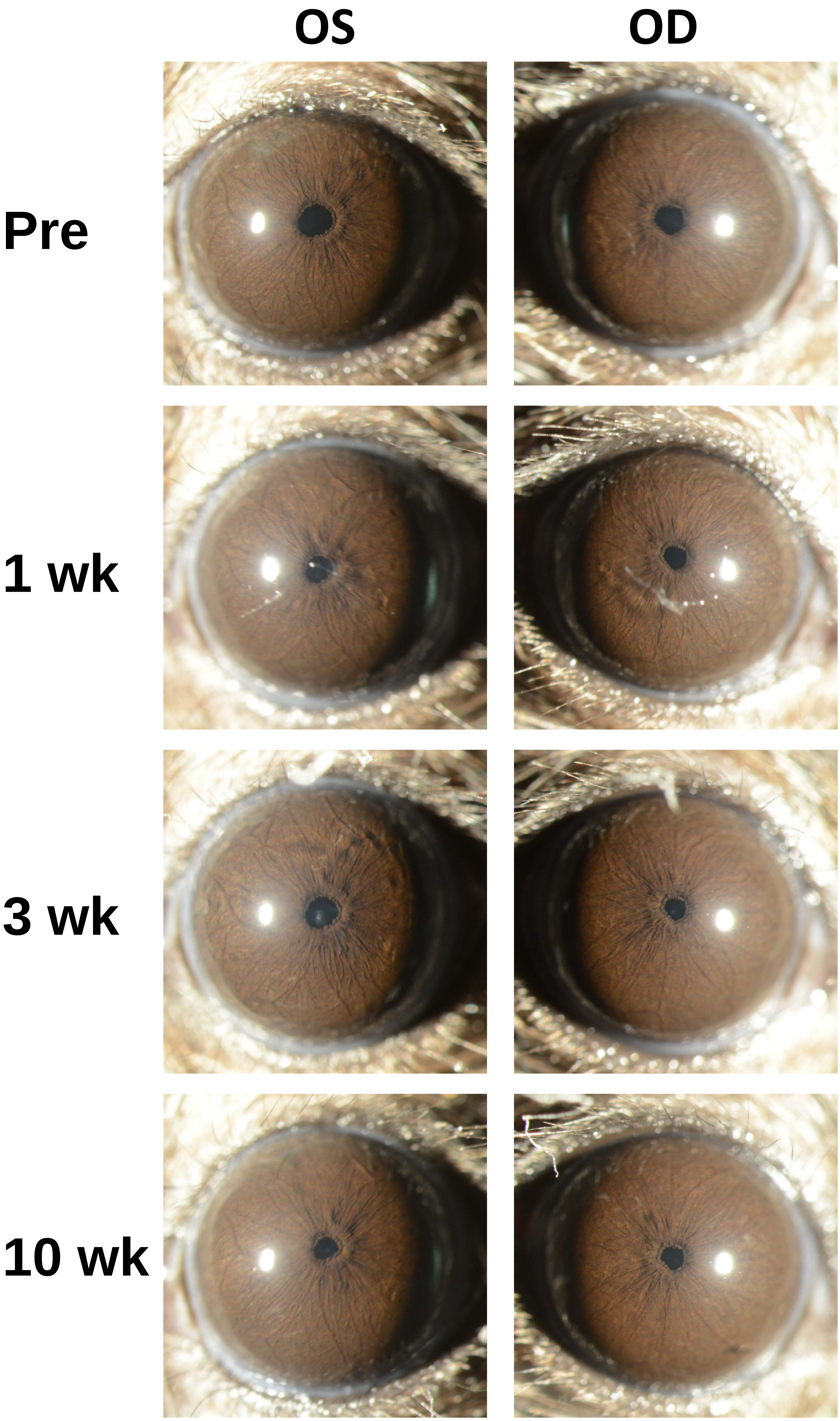

Appendix 7
Mouse ID: 28446

Masked assignment: **Naïve**
Actual group: **Naïve**

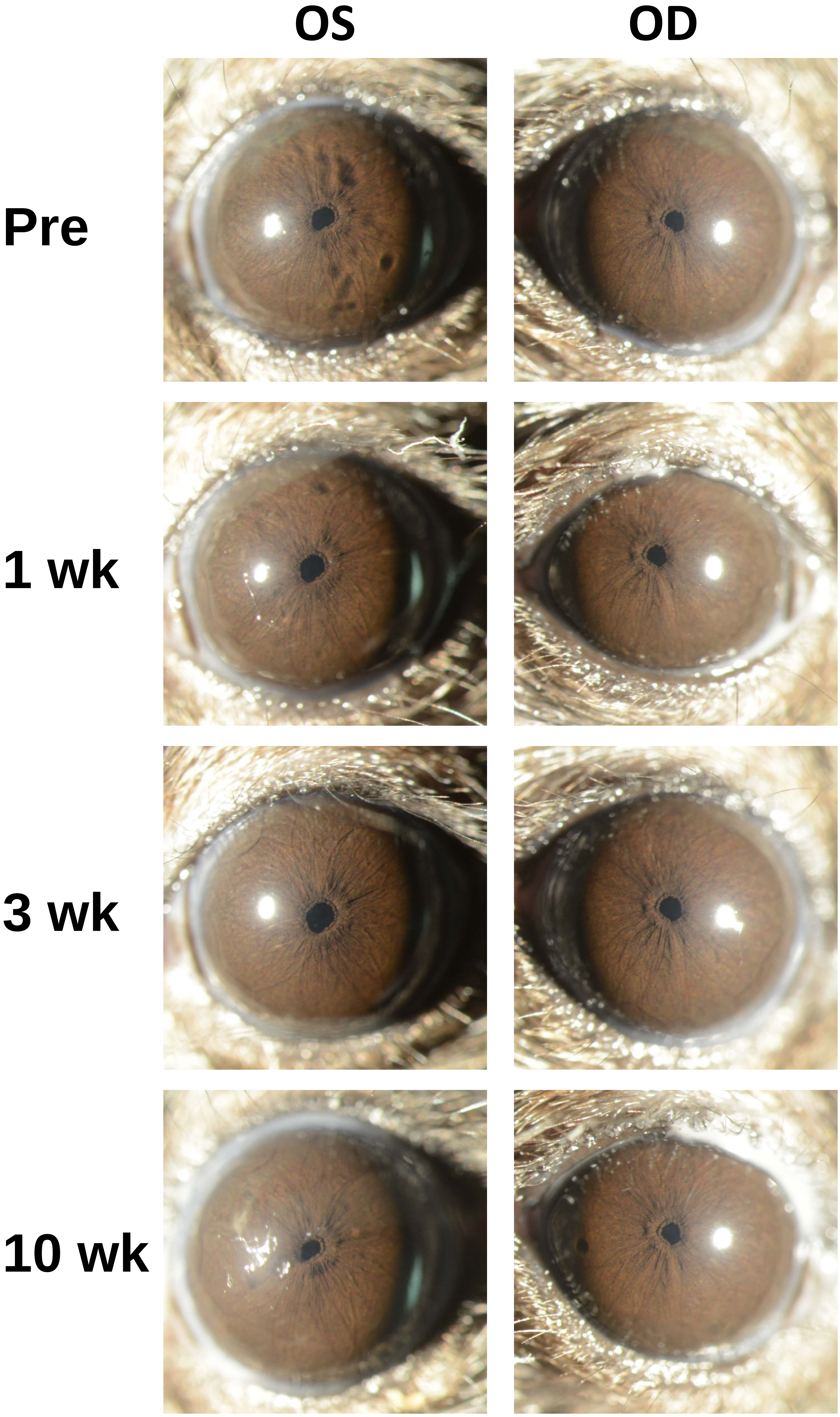

Appendix 7
Mouse ID: 28457

Masked assignment: **Naïve**
Actual group: **Naïve**

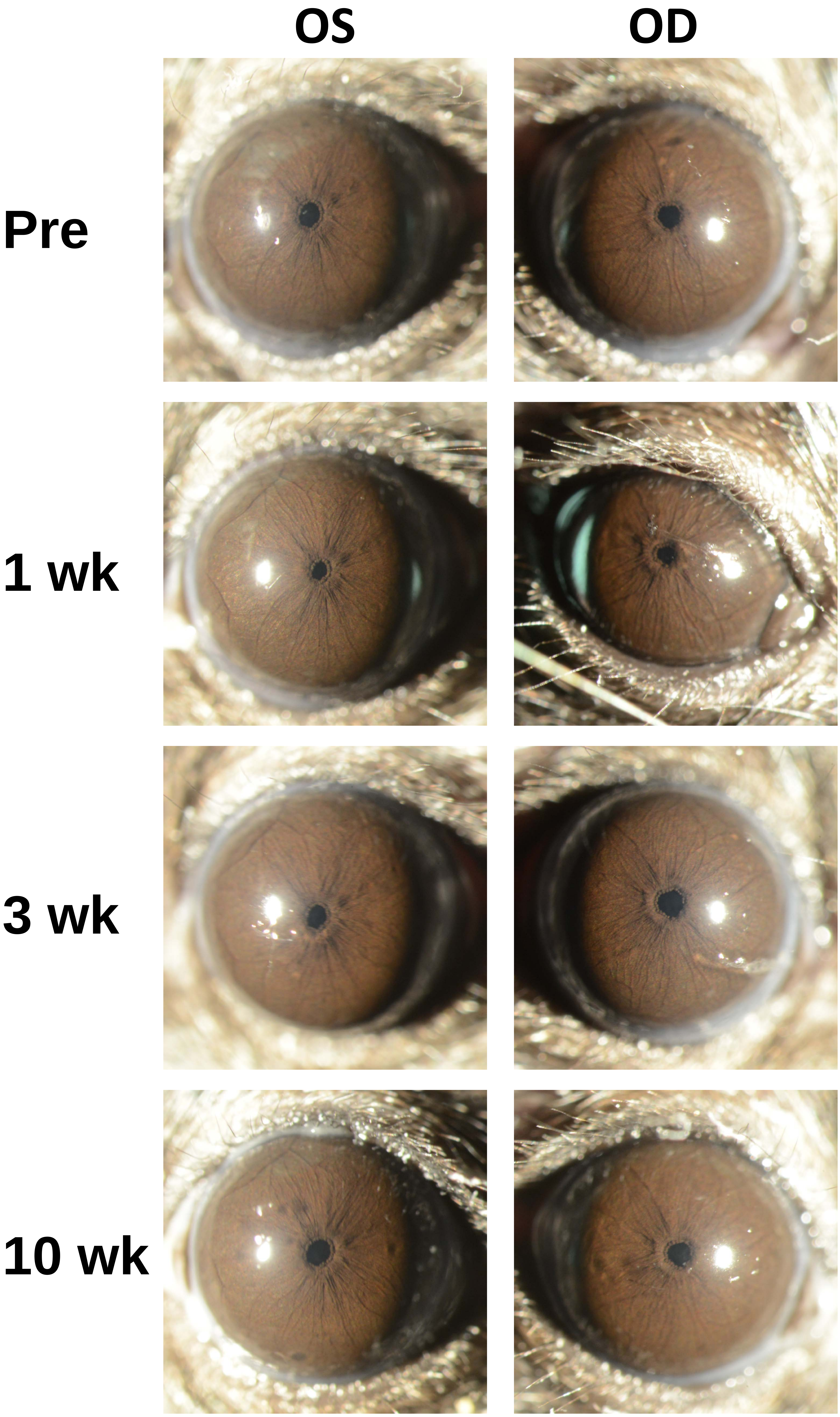

Appendix 7
Mouse ID: 28460

Masked assignment: **Naïve**
Actual group: **Naïve**

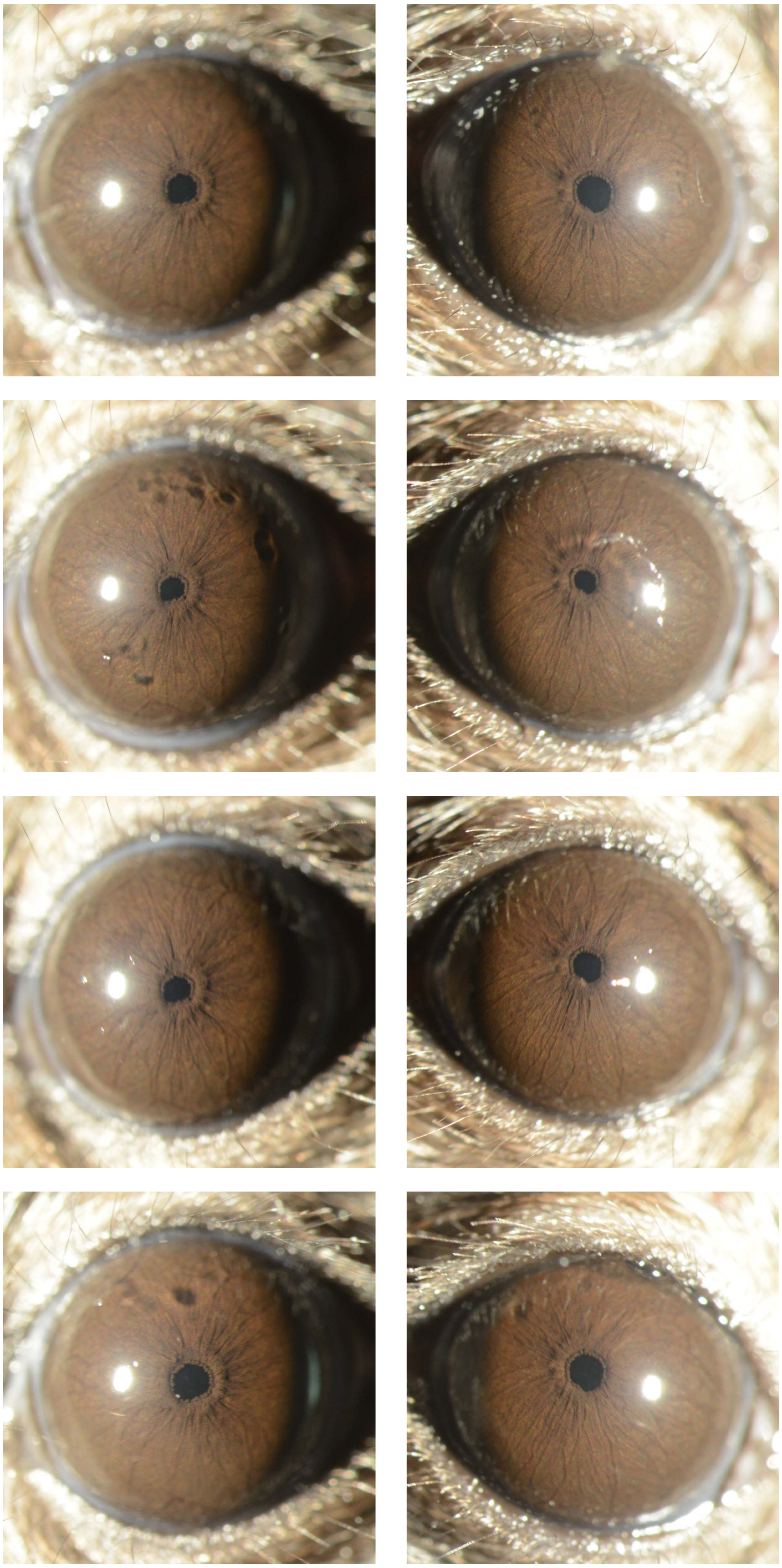

Appendix 7
Mouse ID: 28463

Masked assignment: **Naïve**
Actual group: **Naïve**

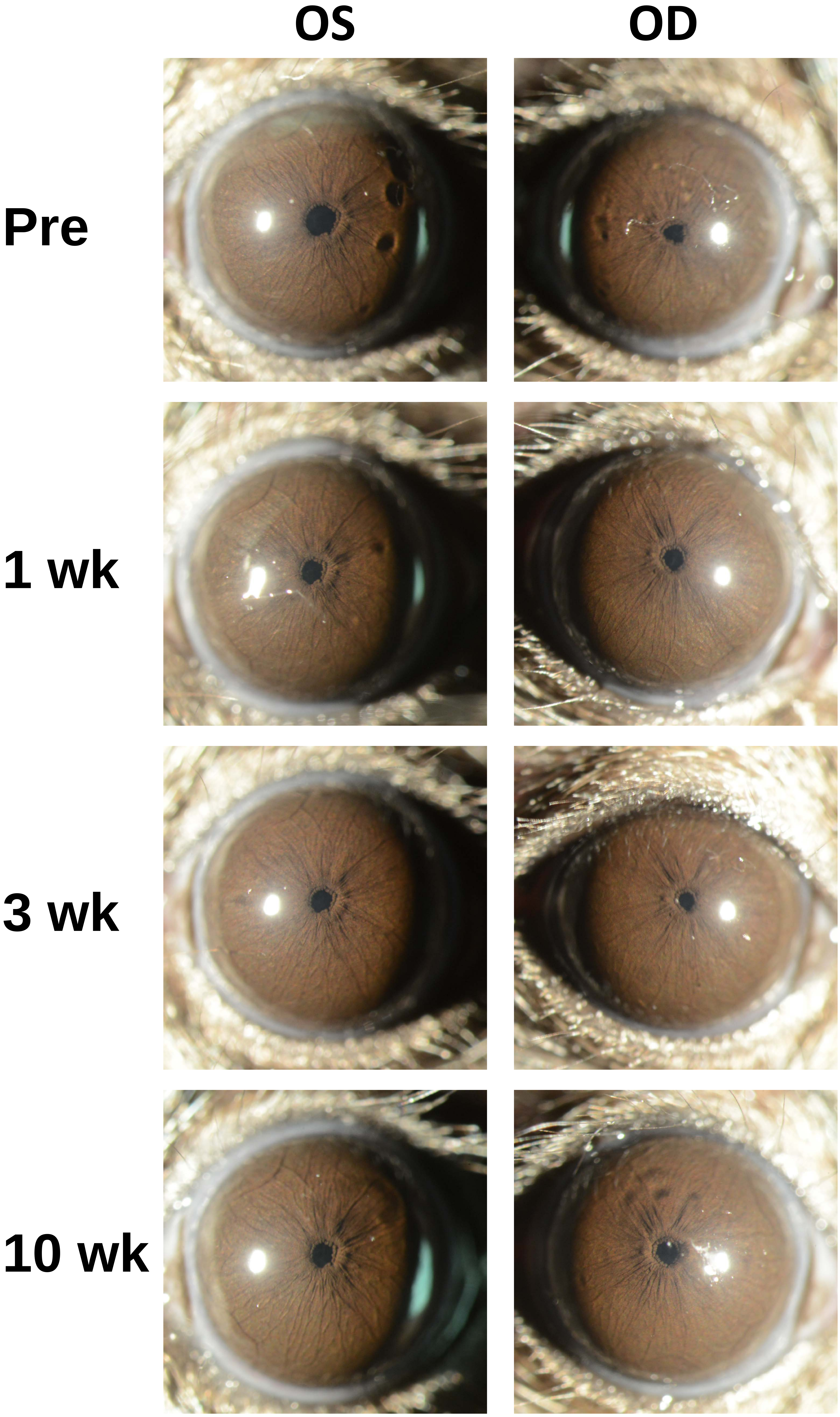

Appendix 7
Mouse ID: 28489

Masked assignment: **Naïve**
Actual group: **Naïve**

**OS** **OD**

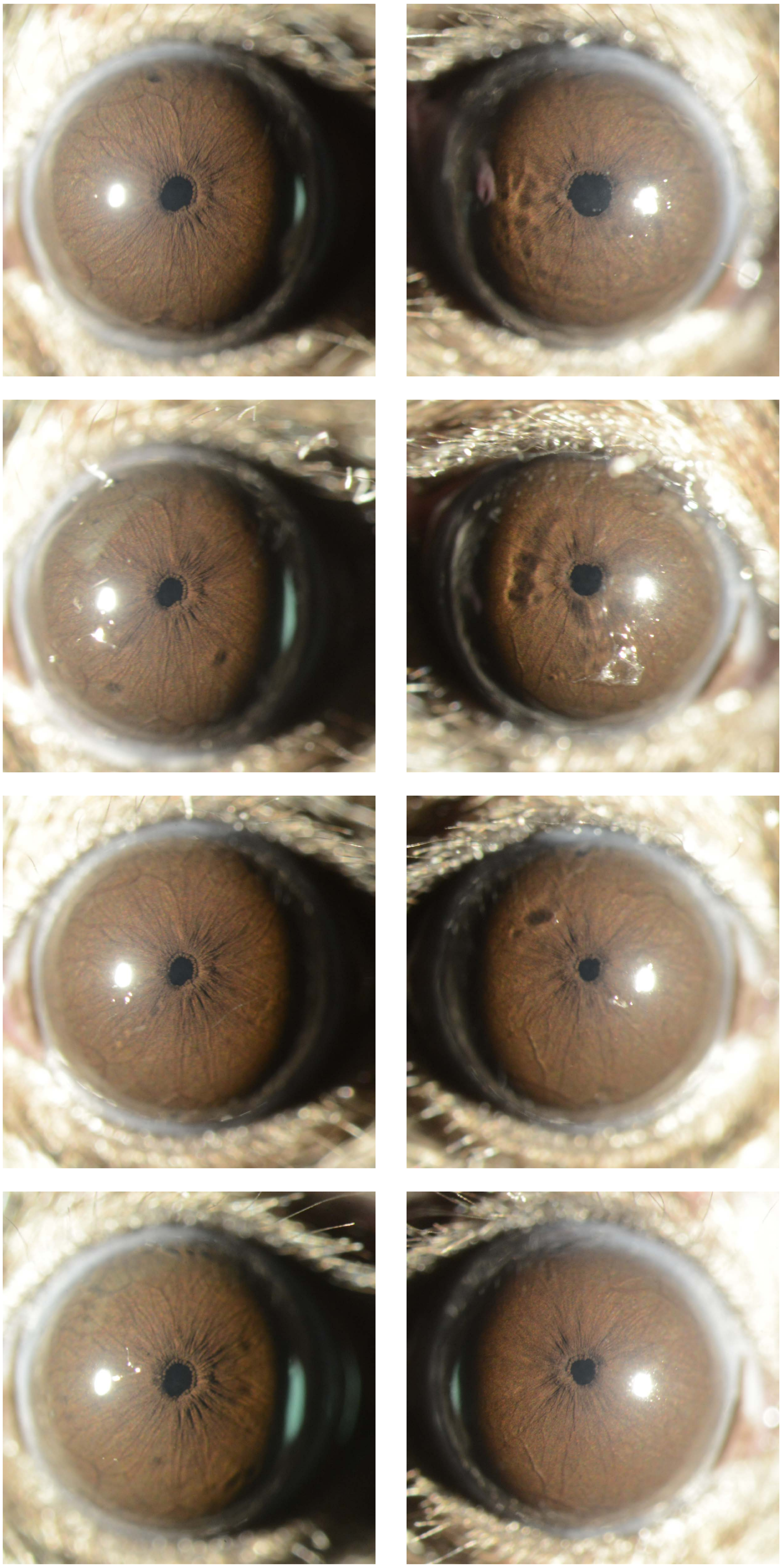

**Pre**

**1 wk**

**3 wk**

**10 wk**
