## Appendix 8 naive 40X for "Recombinant adenovirus causes prolonged mobilization of macrophages in the anterior chamber of mice"

Appendix 8
Mouse ID: 28443
10-week timepoint

Masked assignment: **Naïve**
Actual group: **Saline** (OS injected)

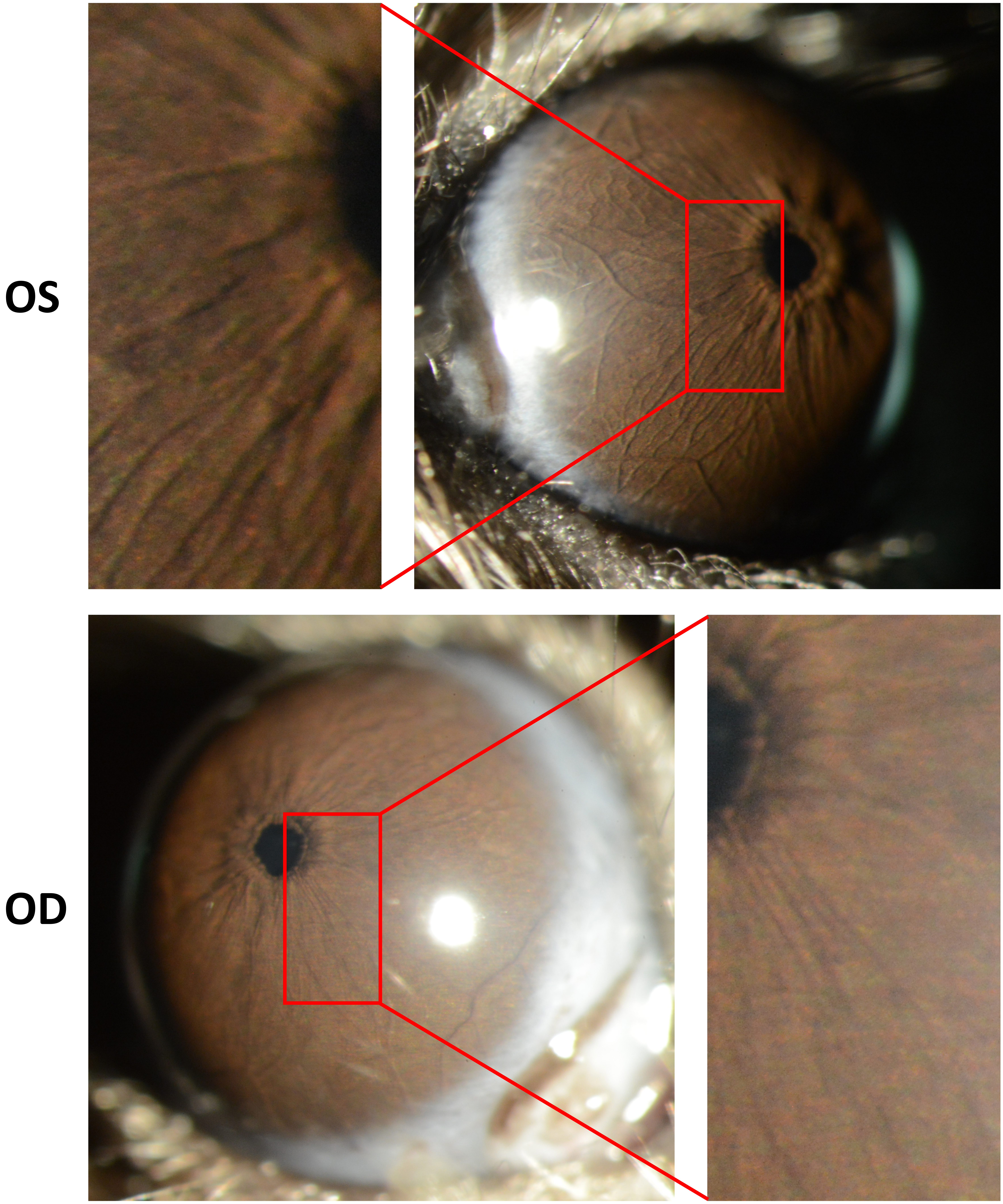

Appendix 8
Mouse ID: 28446
10-week timepoint

Masked assignment: **Naïve**
Actual group: **Naïve**

**OS**

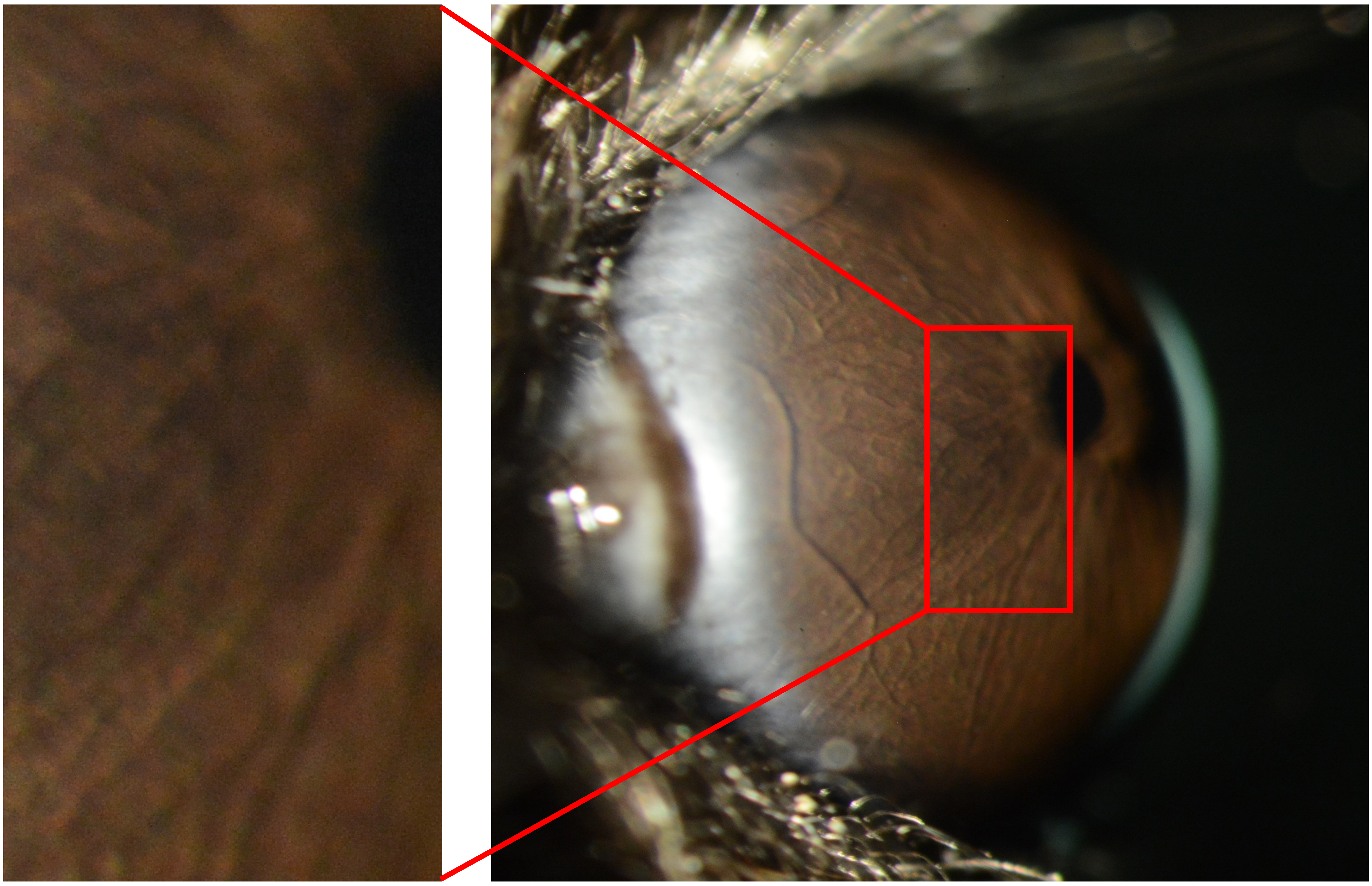

**OD**

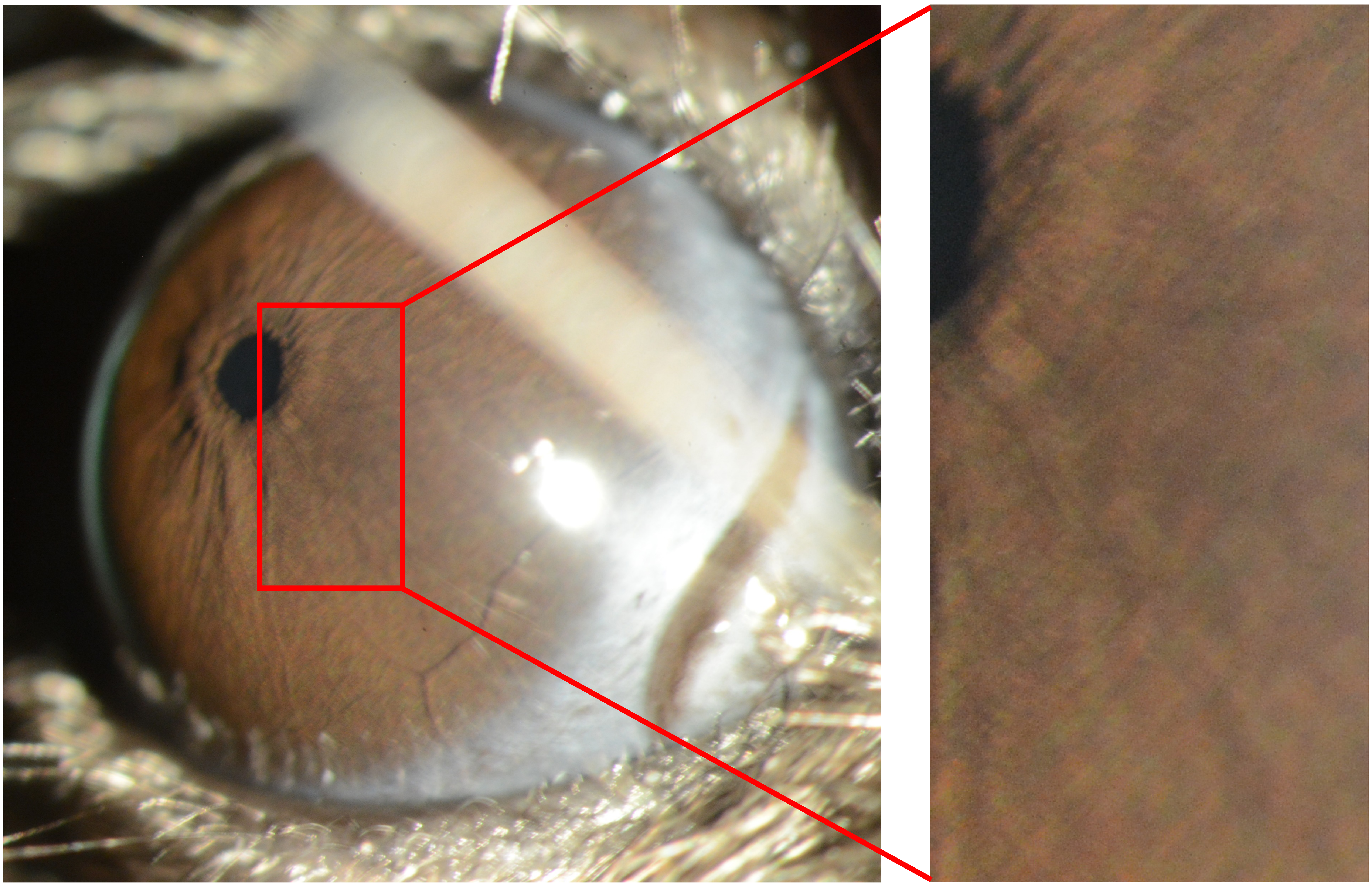

Appendix 8
Mouse ID: 28457
10-week timepoint

Masked assignment: **Naïve**
Actual group: **Naïve**

**OS**

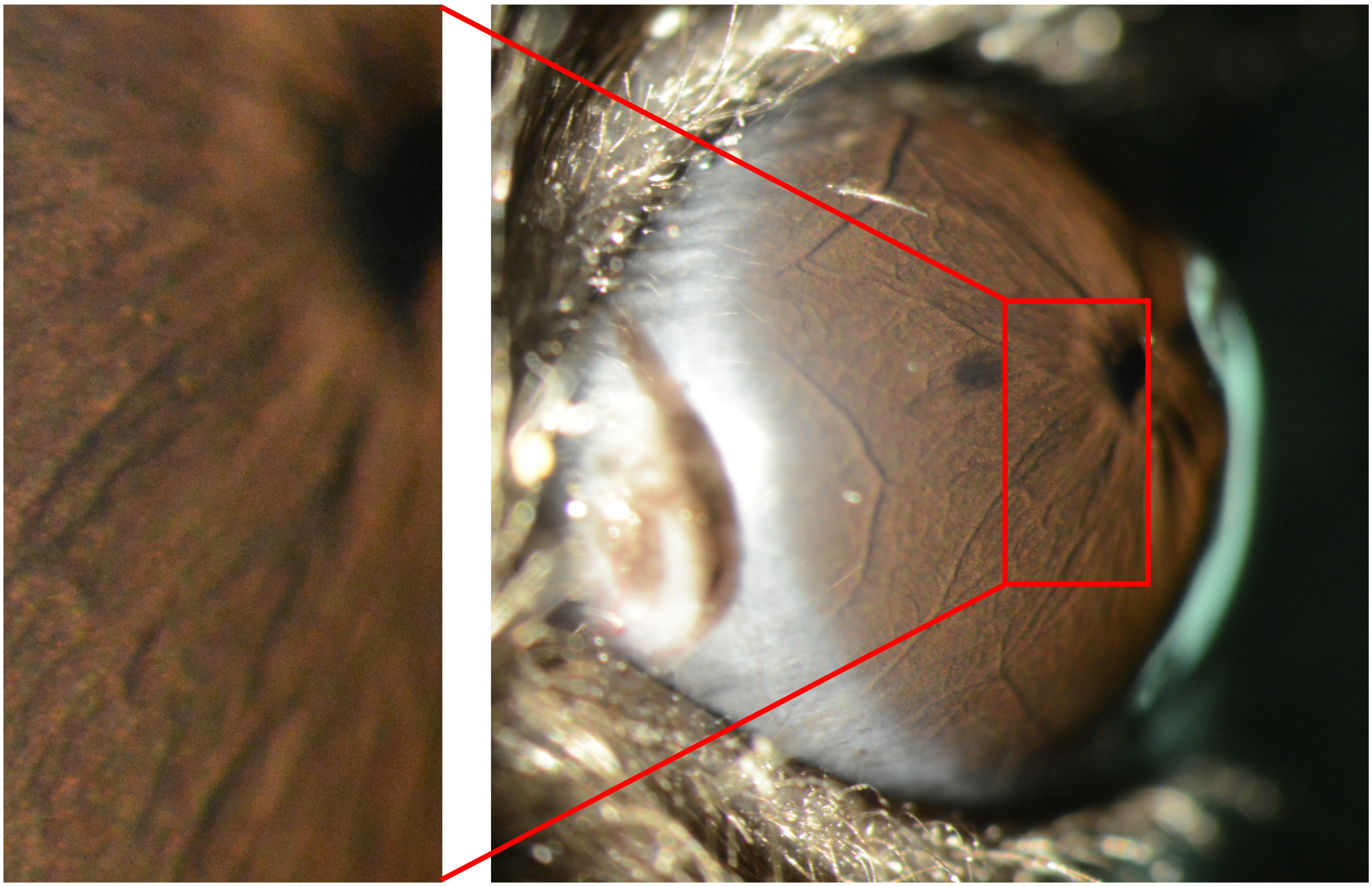

**OD**

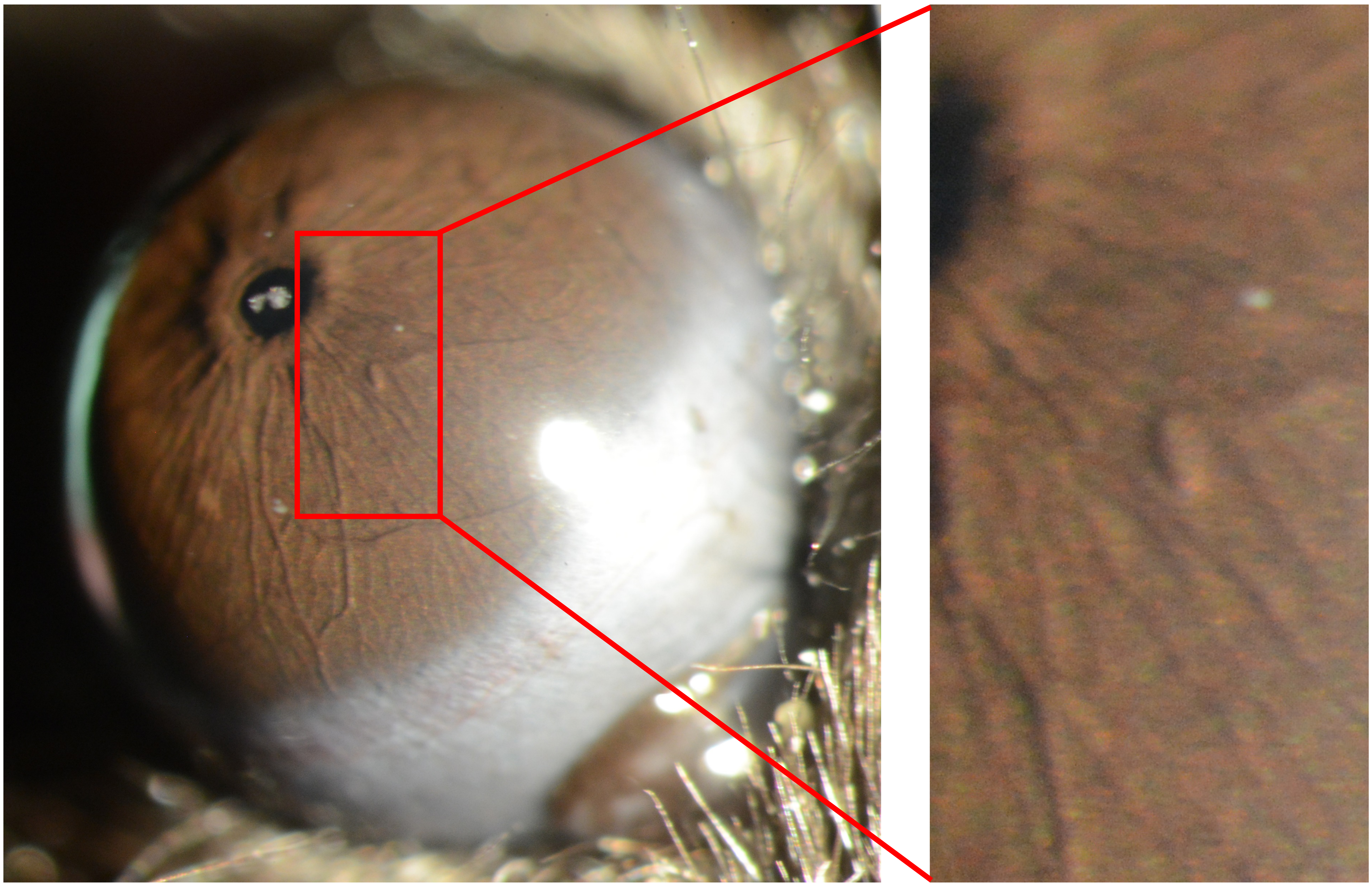

Appendix 8
Mouse ID: 28460
10-week timepoint

Masked assignment: **Naïve**
Actual group: **Naïve**

**OS**

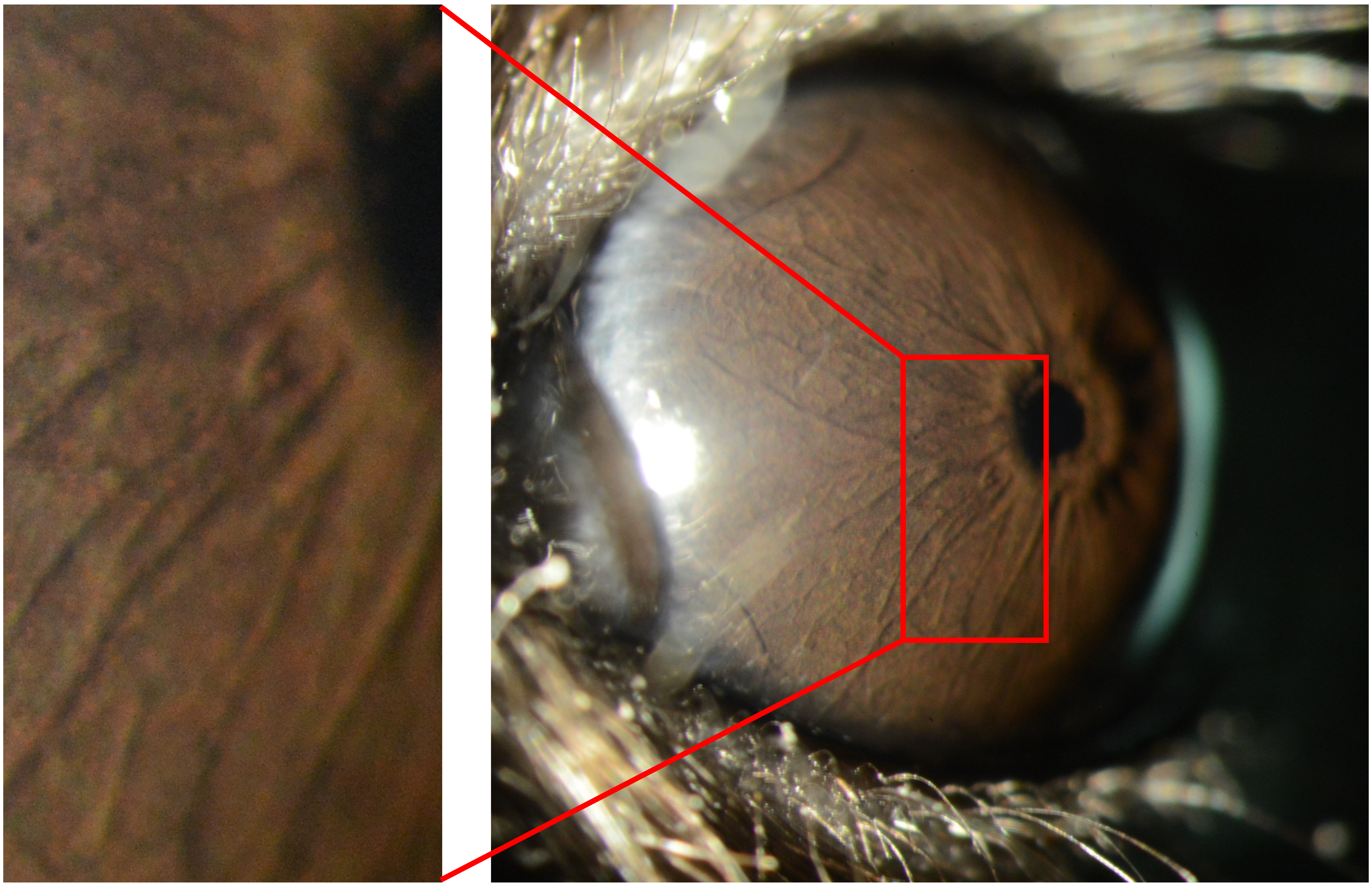

**OD**

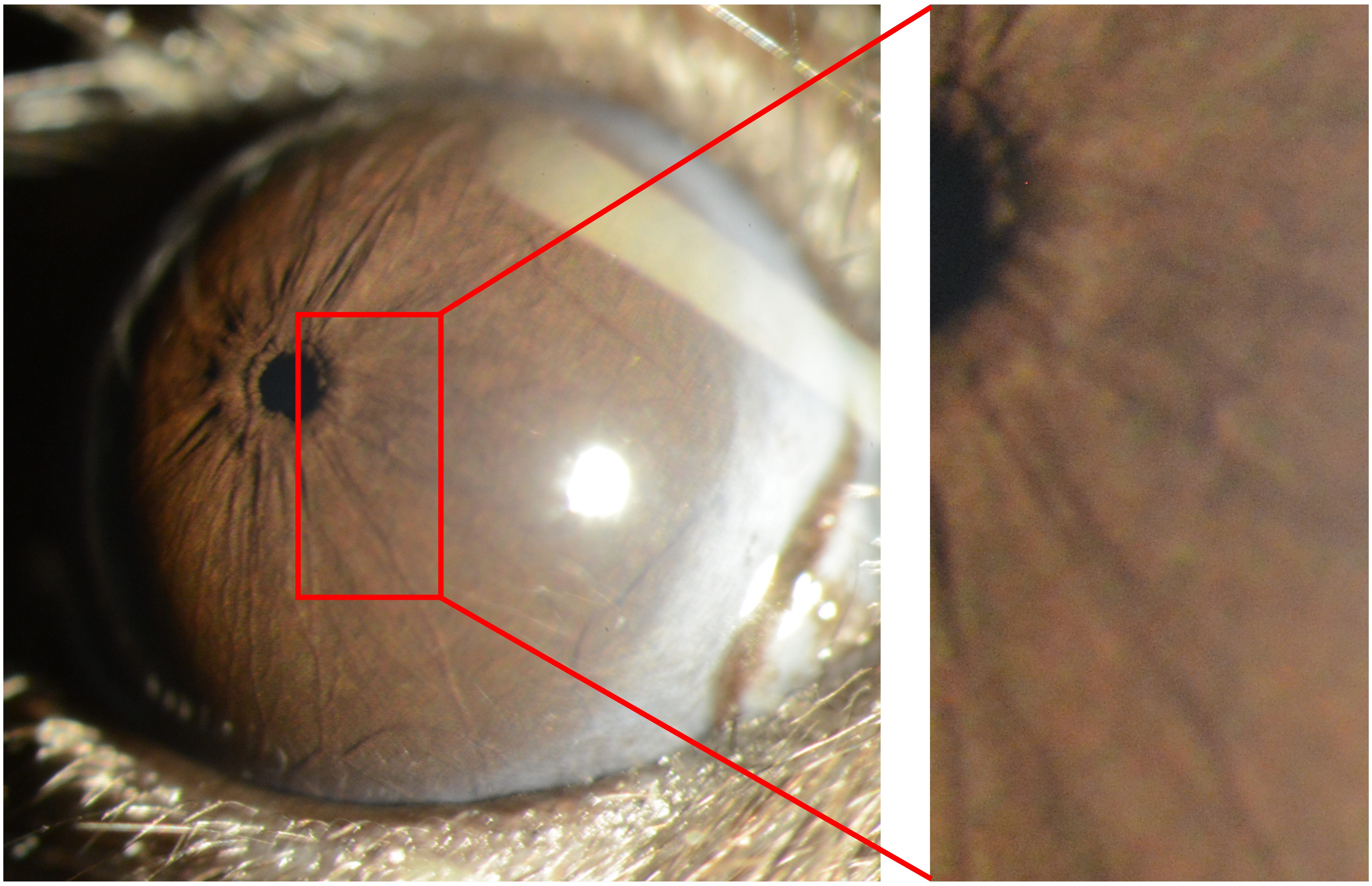

Appendix 8
Mouse ID: 28463
10-week timepoint

Masked assignment: **Naïve**
Actual group: **Naïve**

**OS**

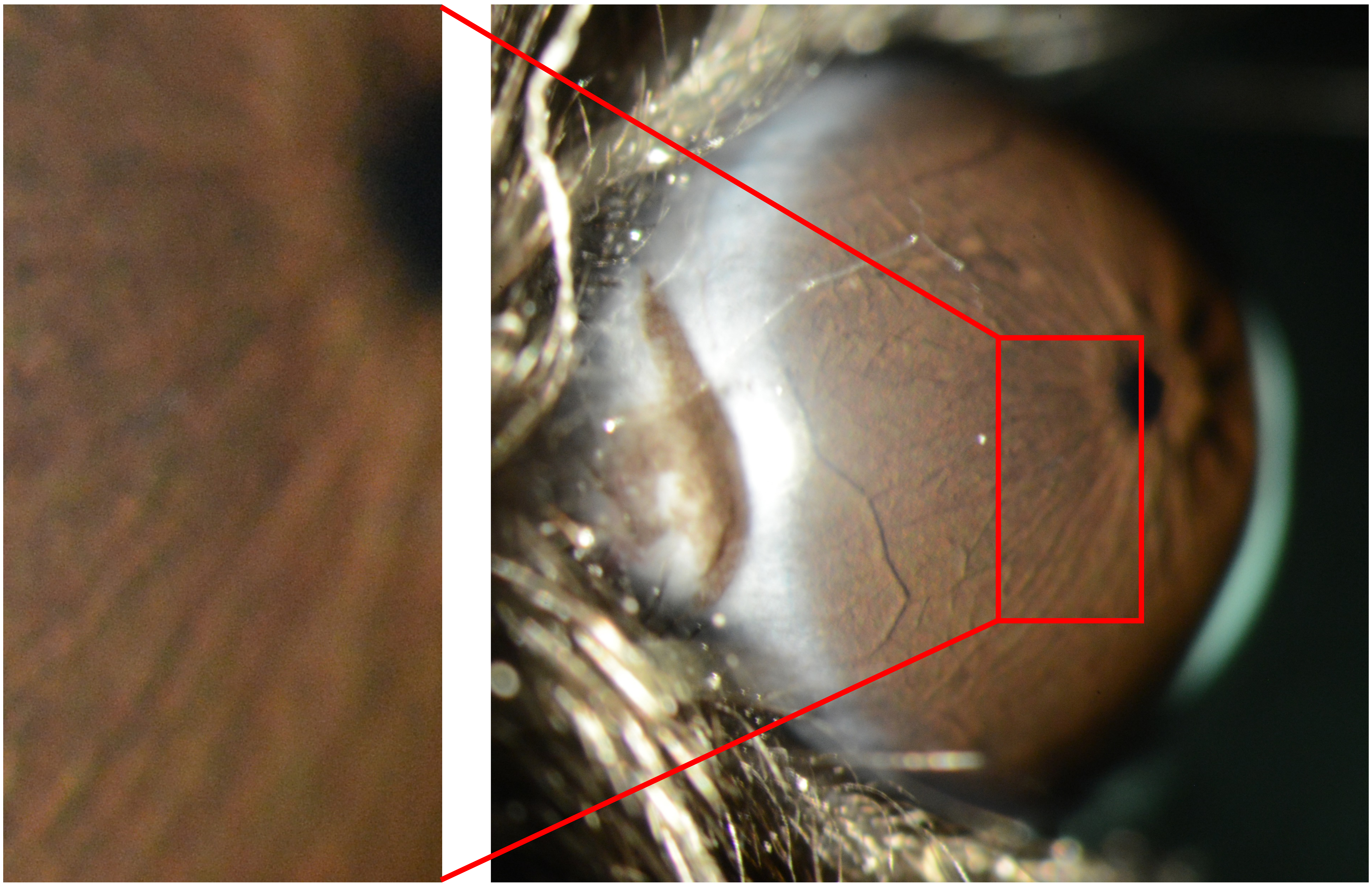

**OD**

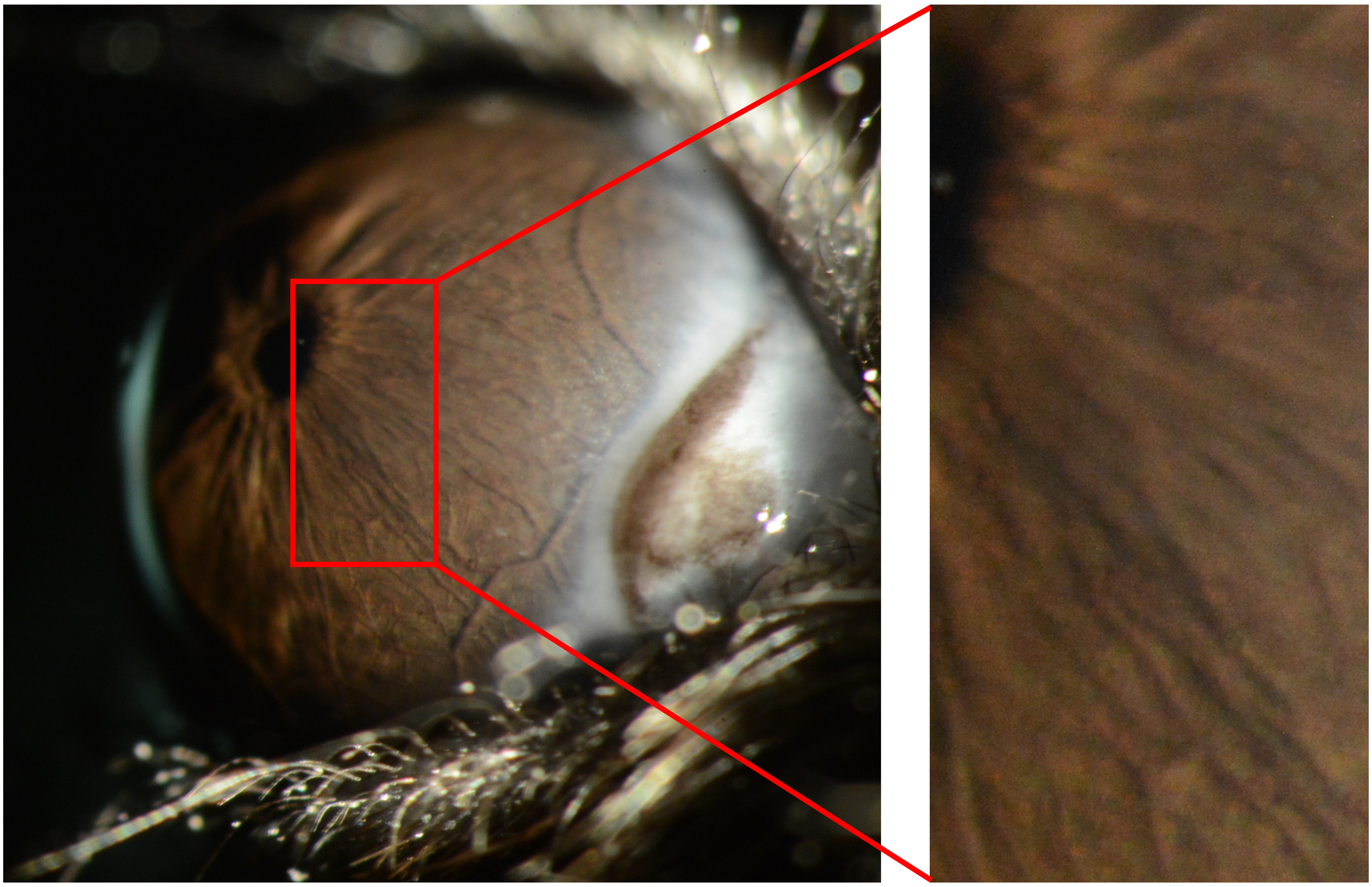

Appendix 8
Mouse ID: 28489
10-week timepoint

Masked assignment: **Naïve**
Actual group: **Naïve**

**OS**

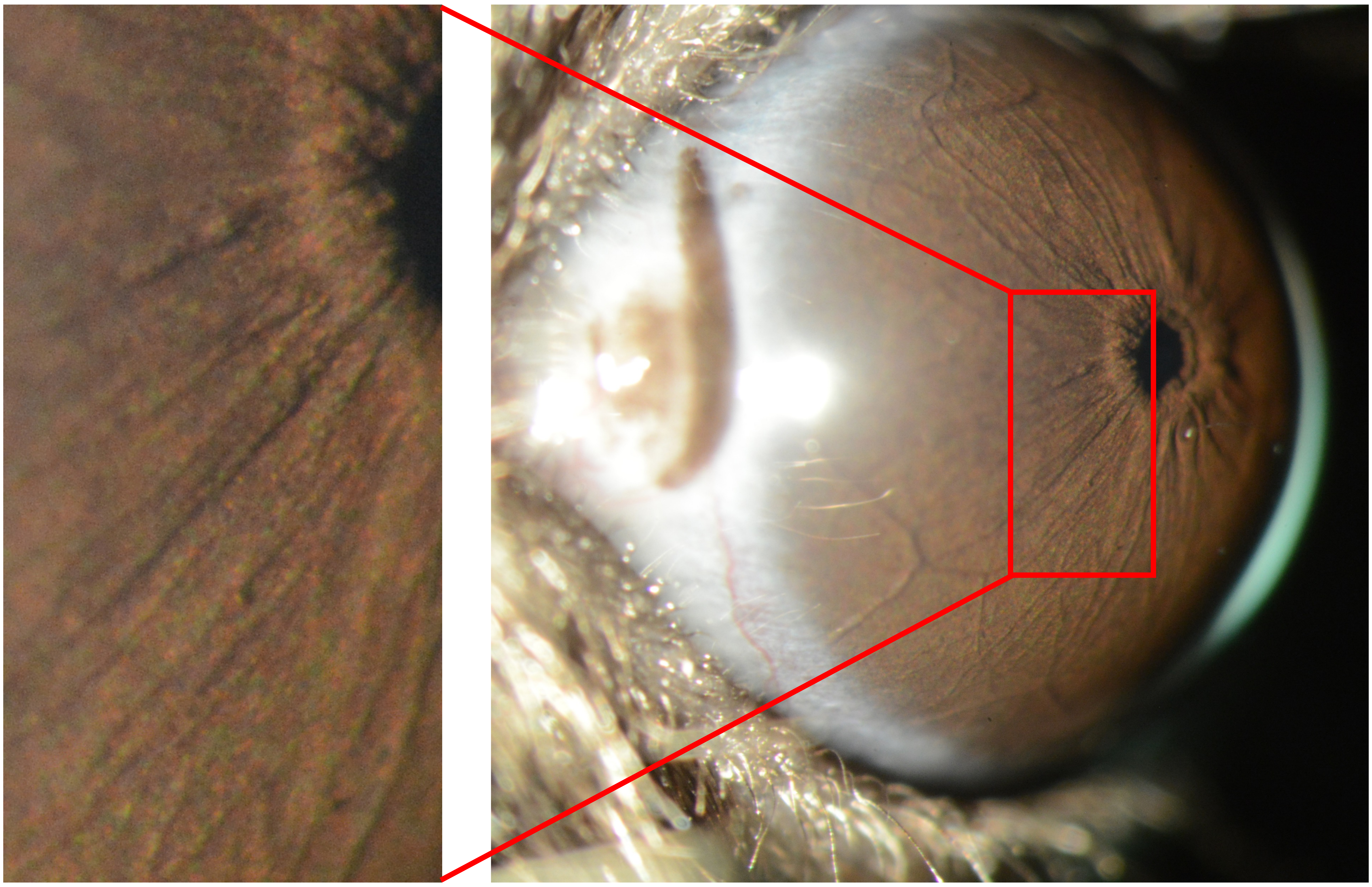

**OD**

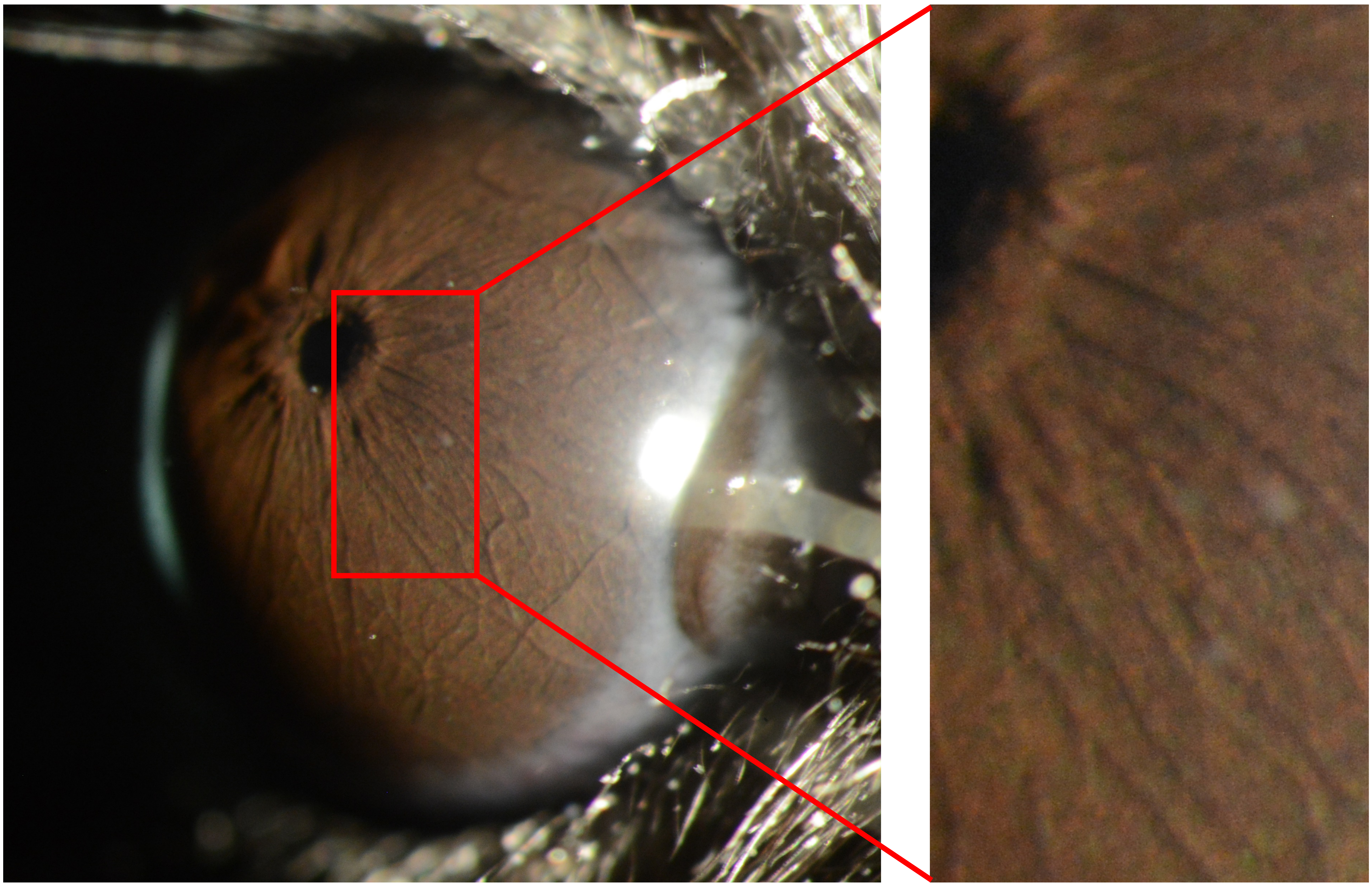
