## Appendix 9 Virus 25X for "Recombinant adenovirus causes prolonged mobilization of macrophages in the anterior chamber of mice"

Appendix 9
Mouse ID: 36546

Masked assignment: **Virus**
Actual group: **Virus** (OS injected)

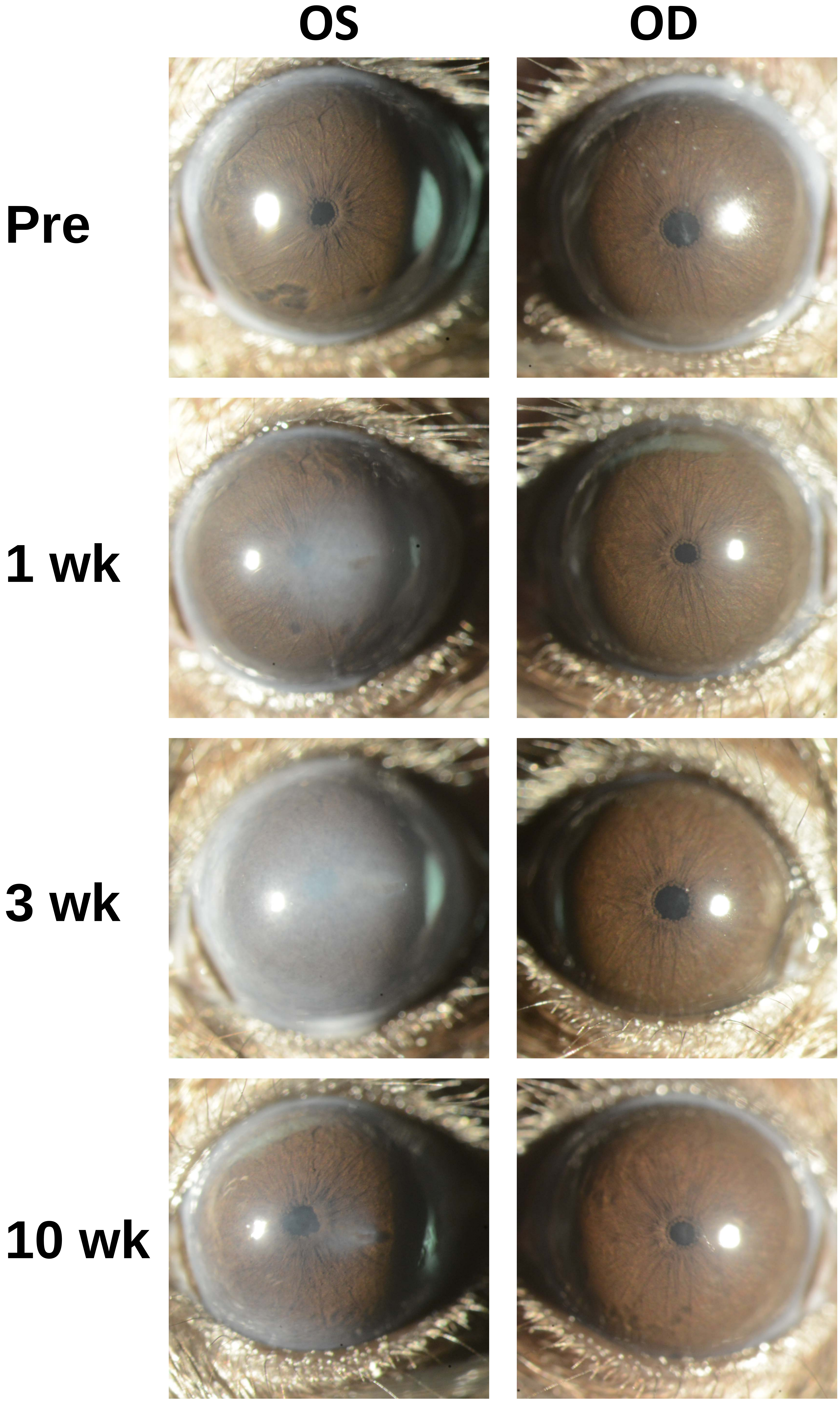

Appendix 9
Mouse ID: 36549

Masked assignment: **Virus**
Actual group: **Virus** (OS injected)

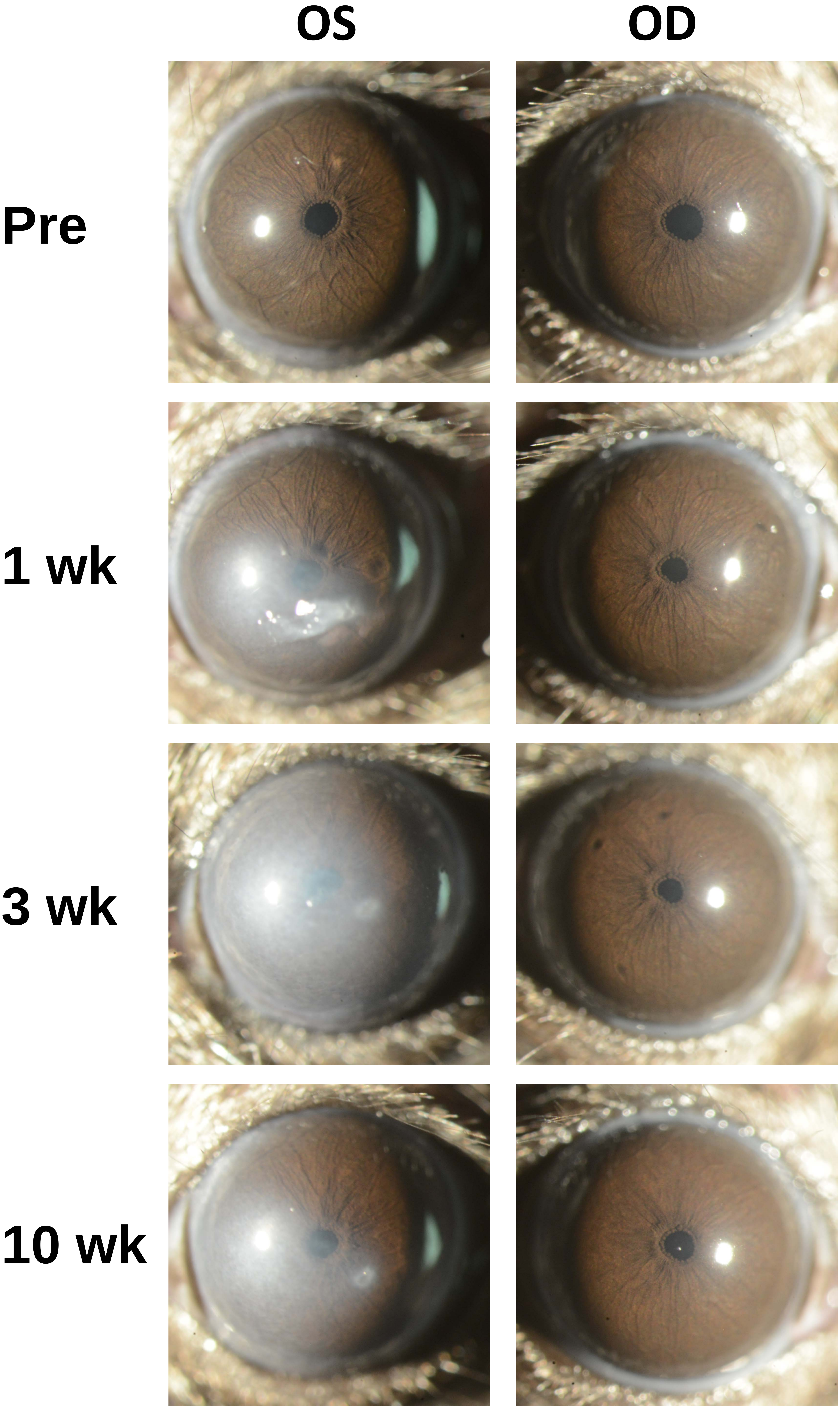

Appendix 9
Mouse ID: 36544

Masked assignment: **Virus**
Actual group: **Virus** (OD injected)

**OS** **OD**

**Pre**

**1 wk**

**3 wk**

**10 wk**

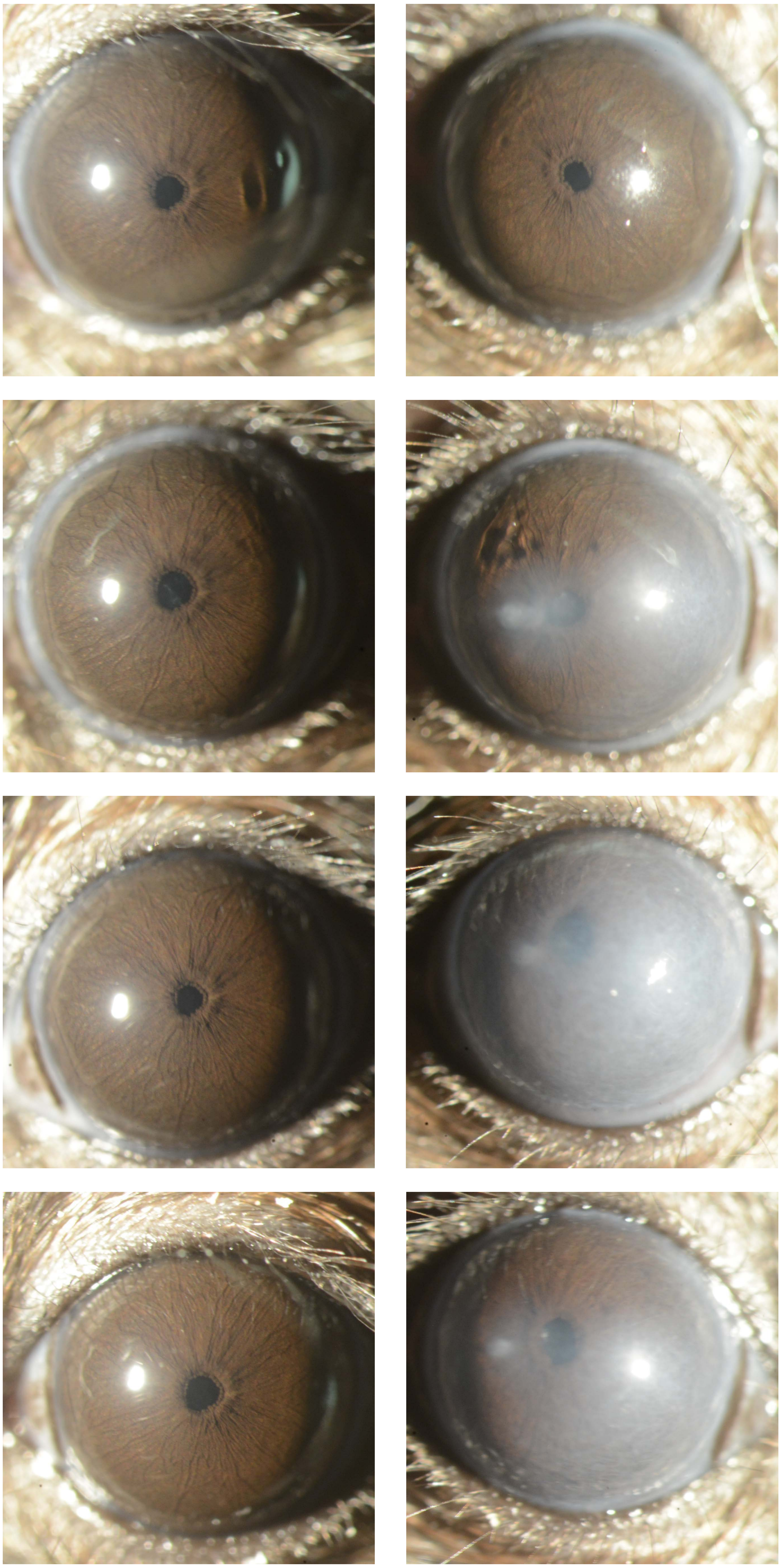

Appendix 9
Mouse ID: 36552

Masked assignment: **Virus**
Actual group: **Virus** (OD injected)

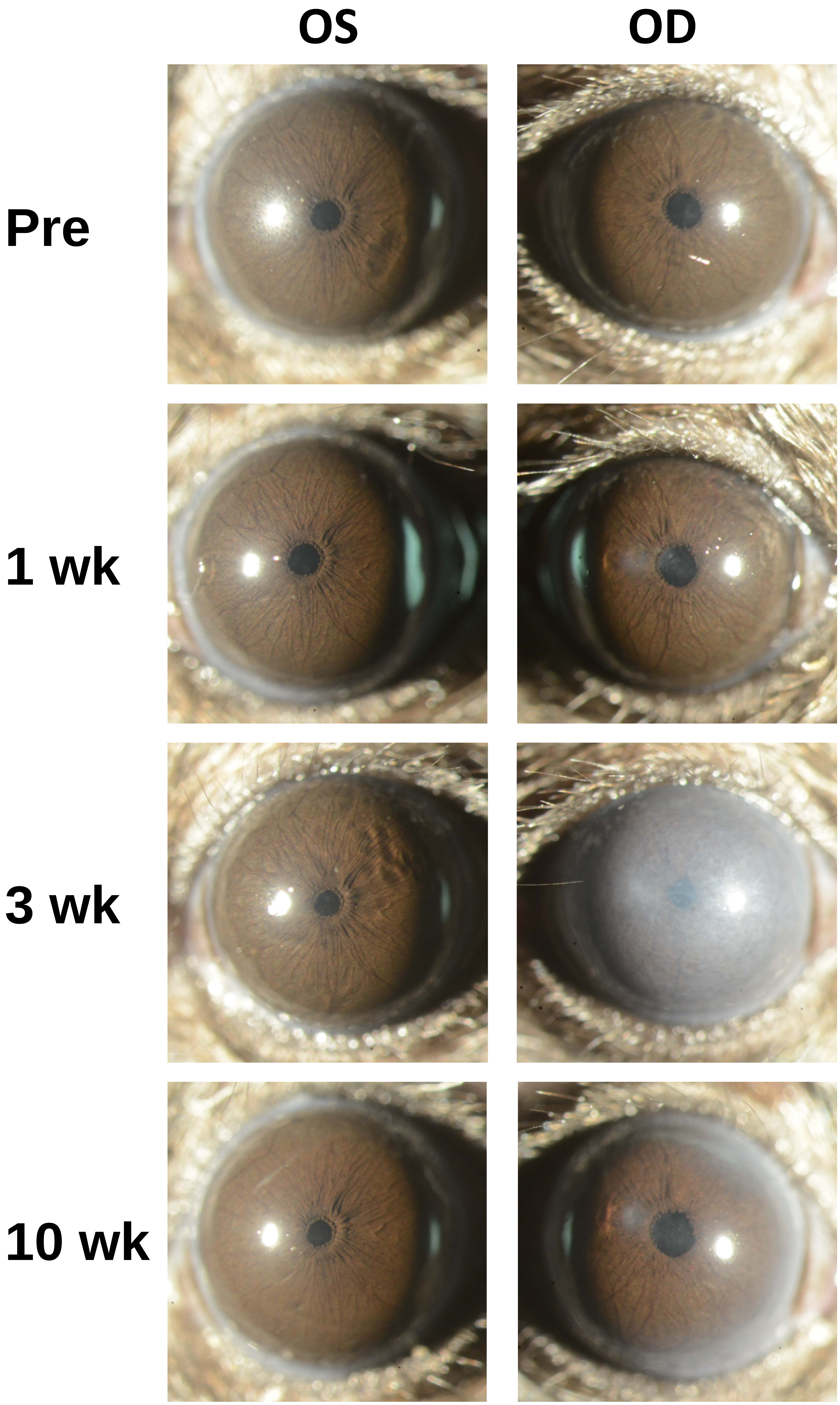
