## Appendix 10 Virus 40X for "Recombinant adenovirus causes prolonged mobilization of macrophages in the anterior chamber of mice"

Appendix 10
Mouse ID: 36546
10-week timepoint

Masked assignment: **Virus**
Actual group: **Virus** (OS injected)

**OS**

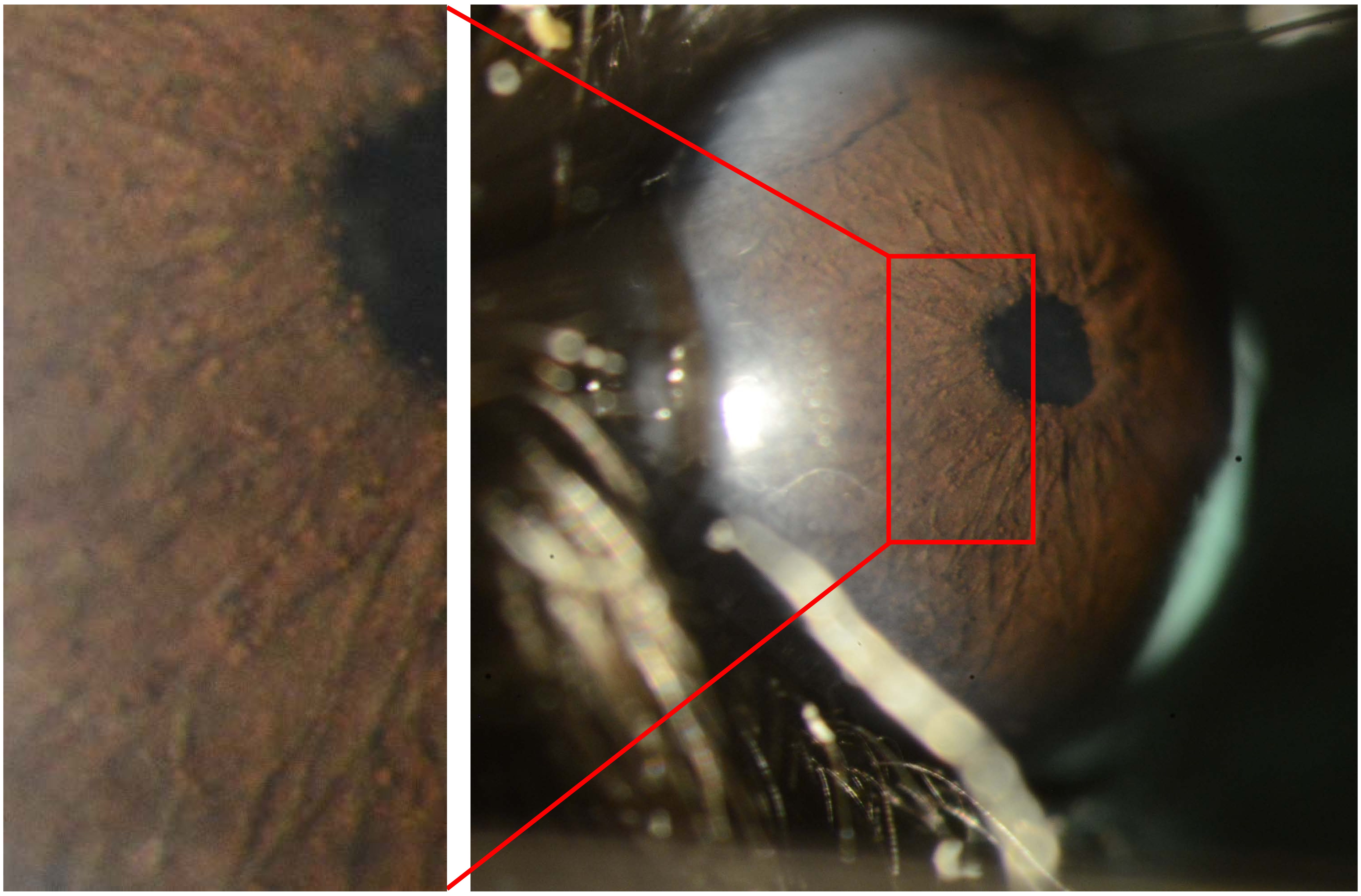

**OD**

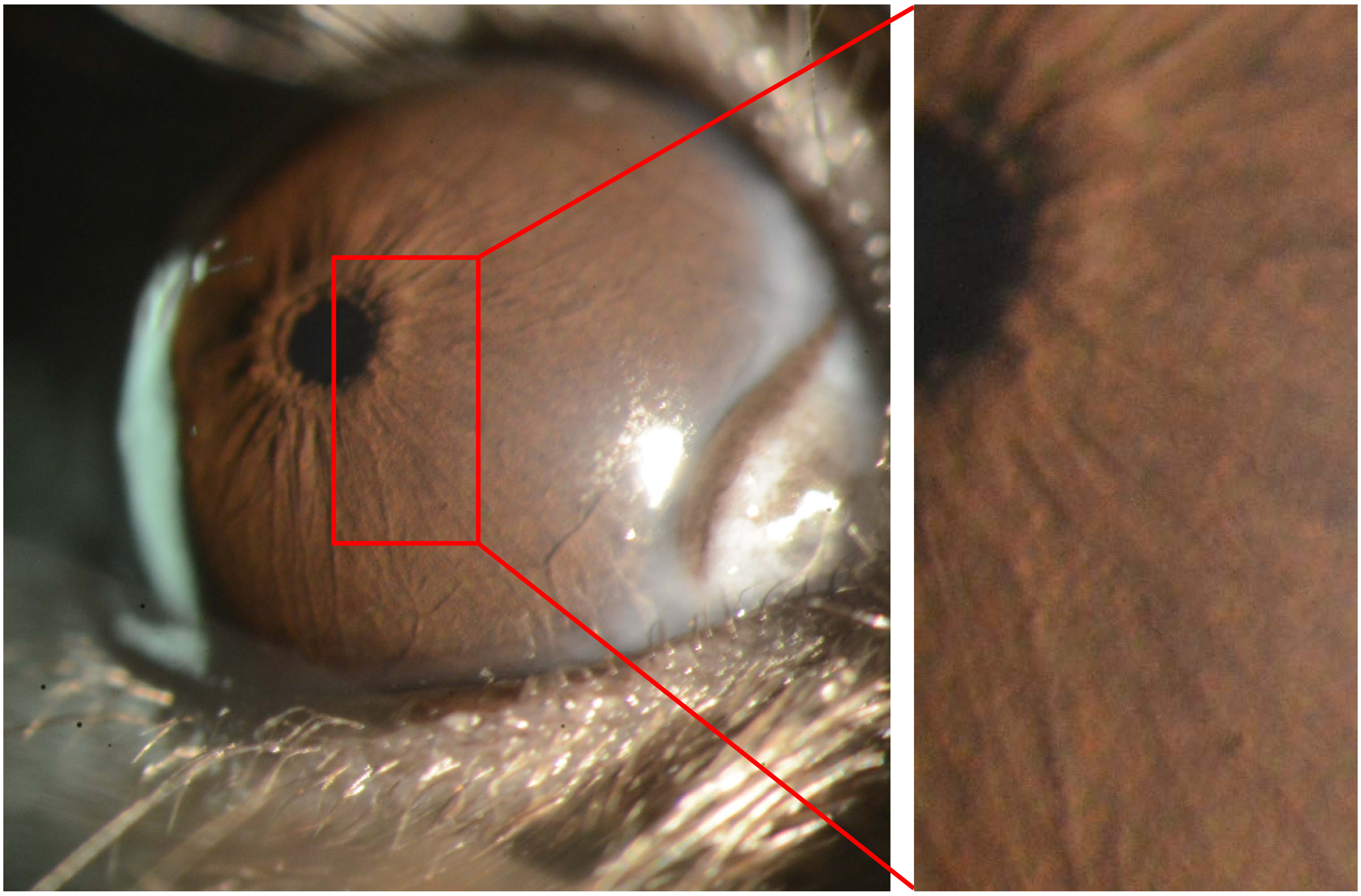

Appendix 10
Mouse ID: 36549
10-week timepoint

Masked assignment: **Virus**
Actual group: **Virus** (OS injected)

**OS**

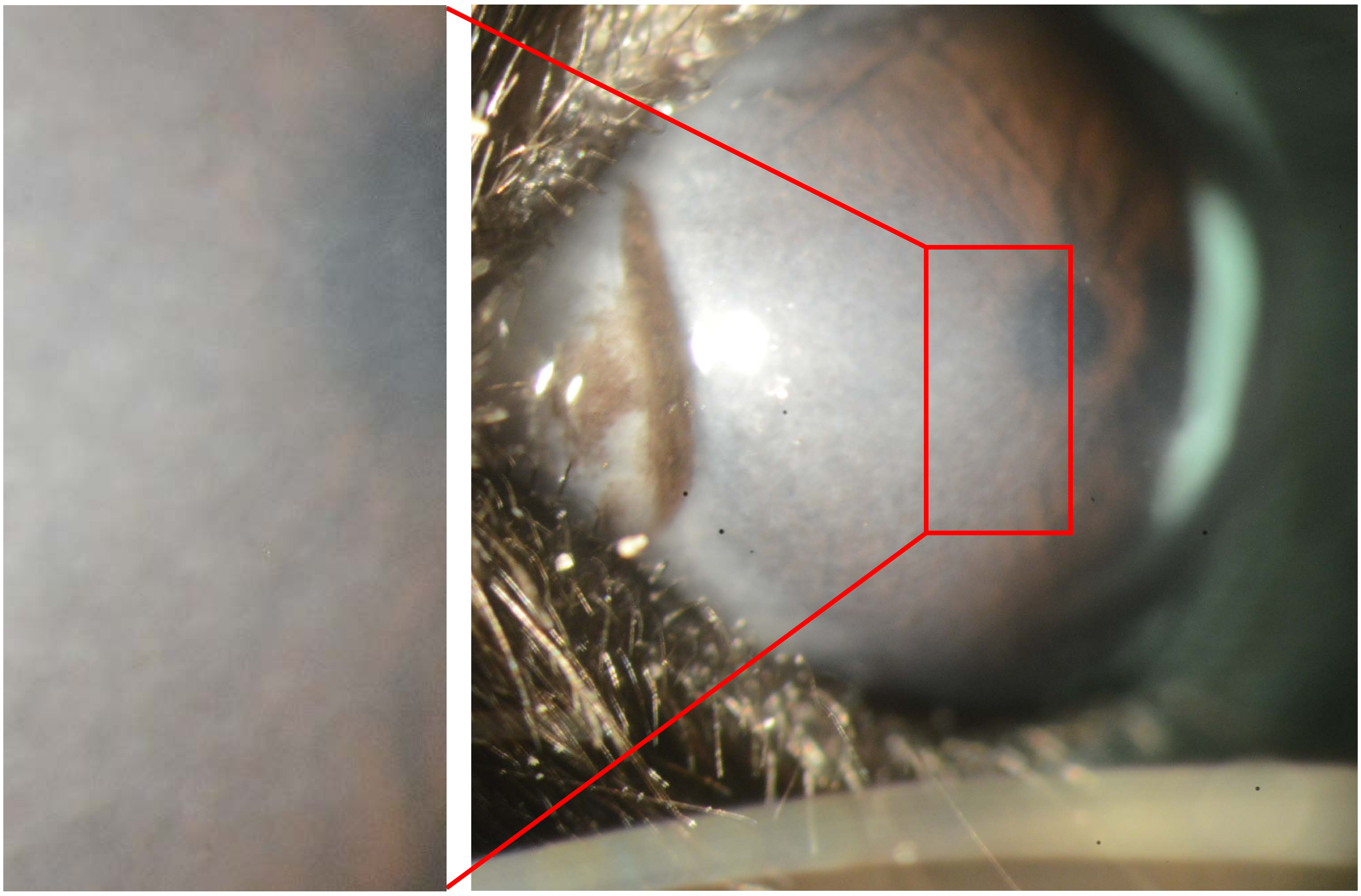

**OD**

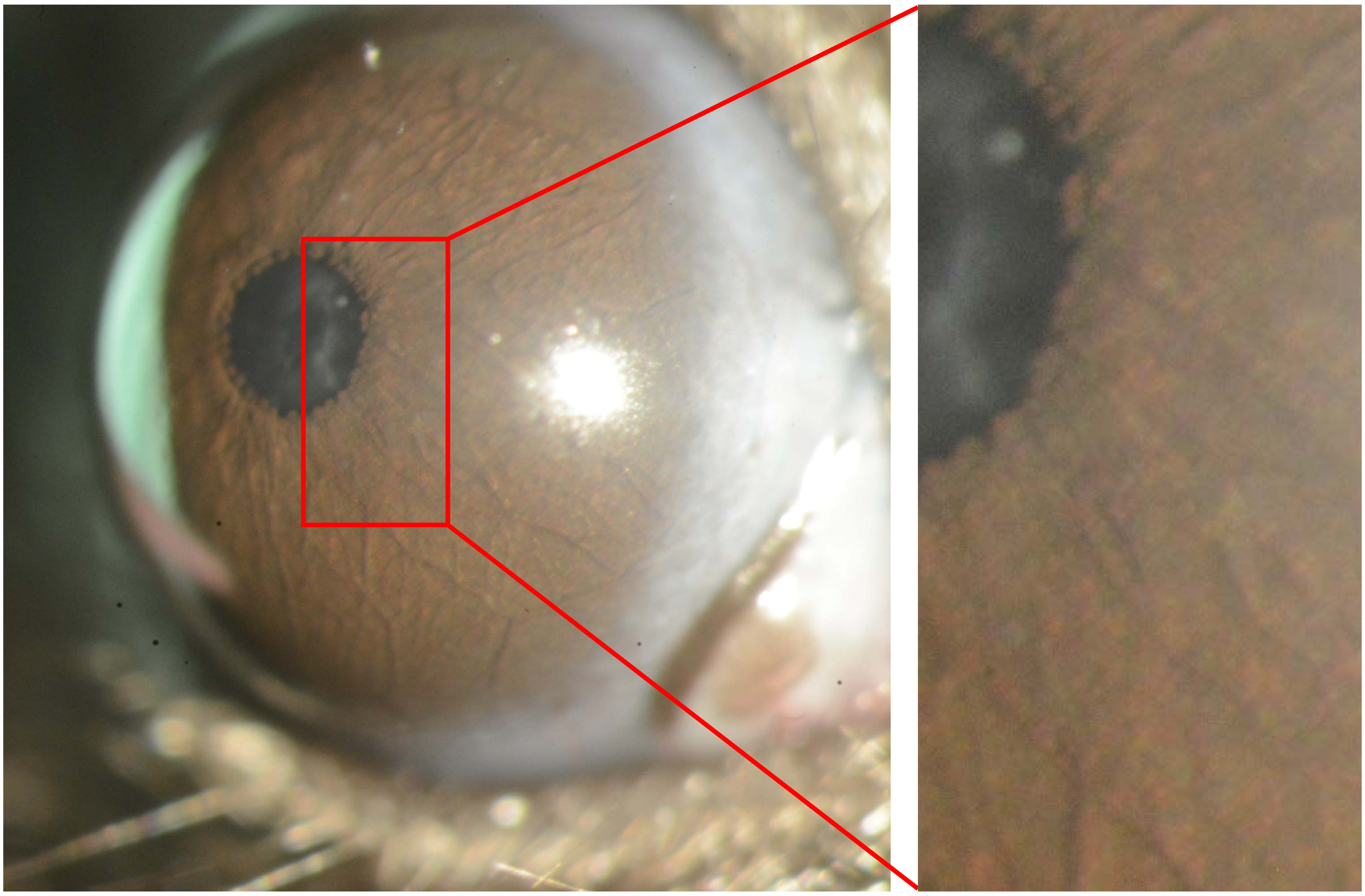

Appendix 10
Mouse ID: 36544
10-week timepoint

Masked assignment: **Virus**
Actual group: **Virus** (OD injected)

**OS**

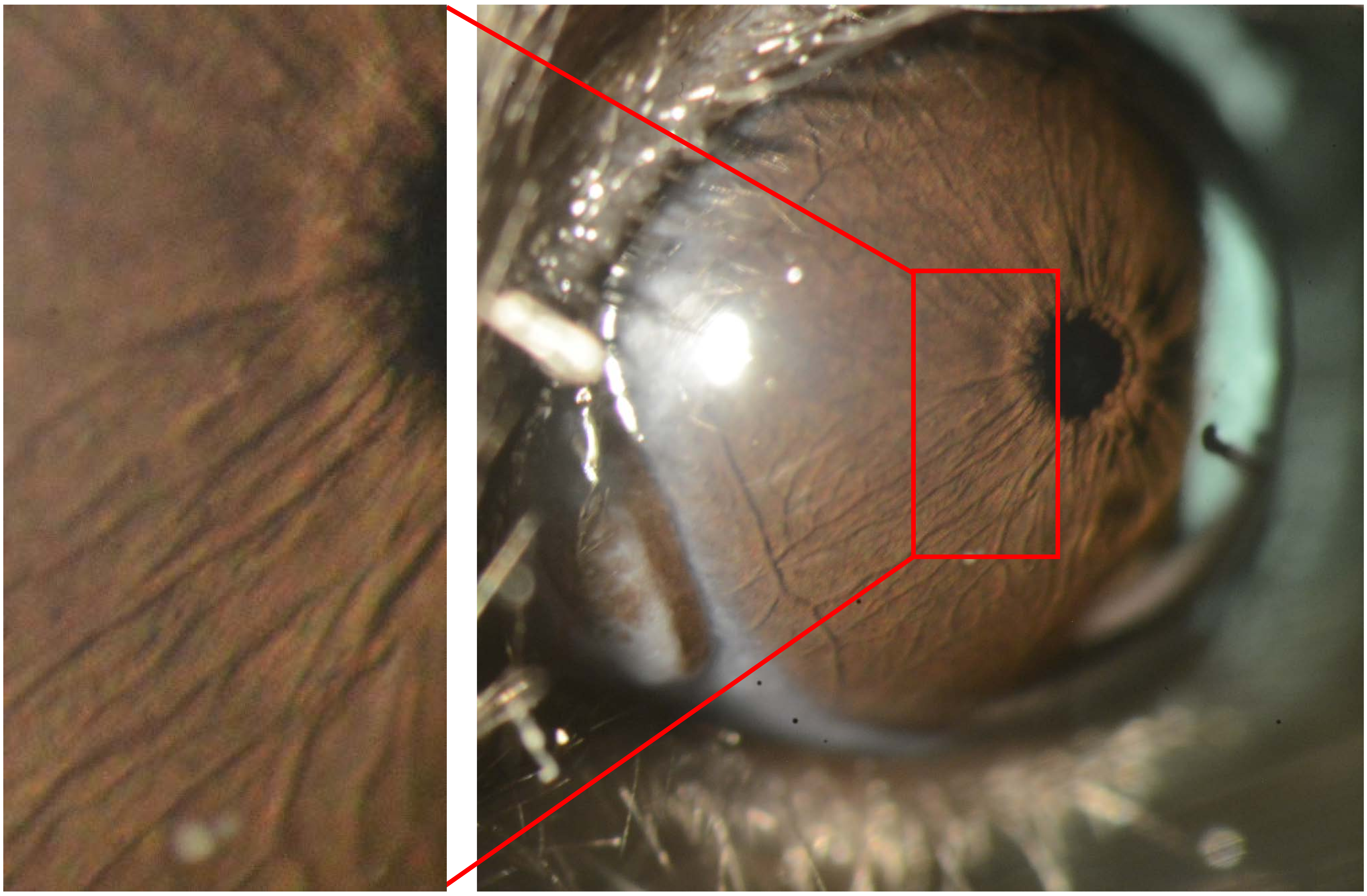

**OD**

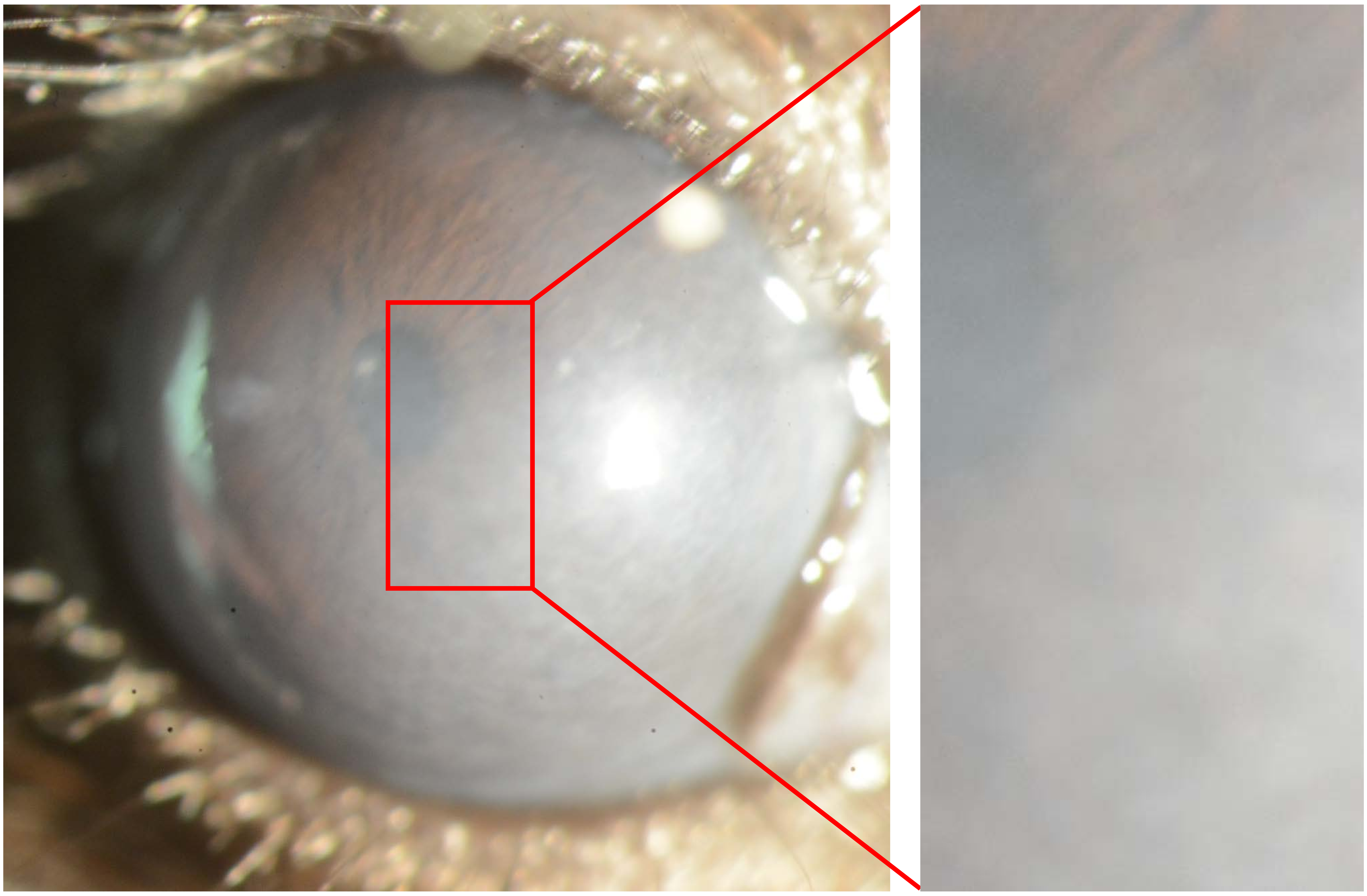

Appendix 10
Mouse ID: 36552
10-week timepoint

Masked assignment: **Virus**
Actual group: **Virus** (OD injected)

**OS**

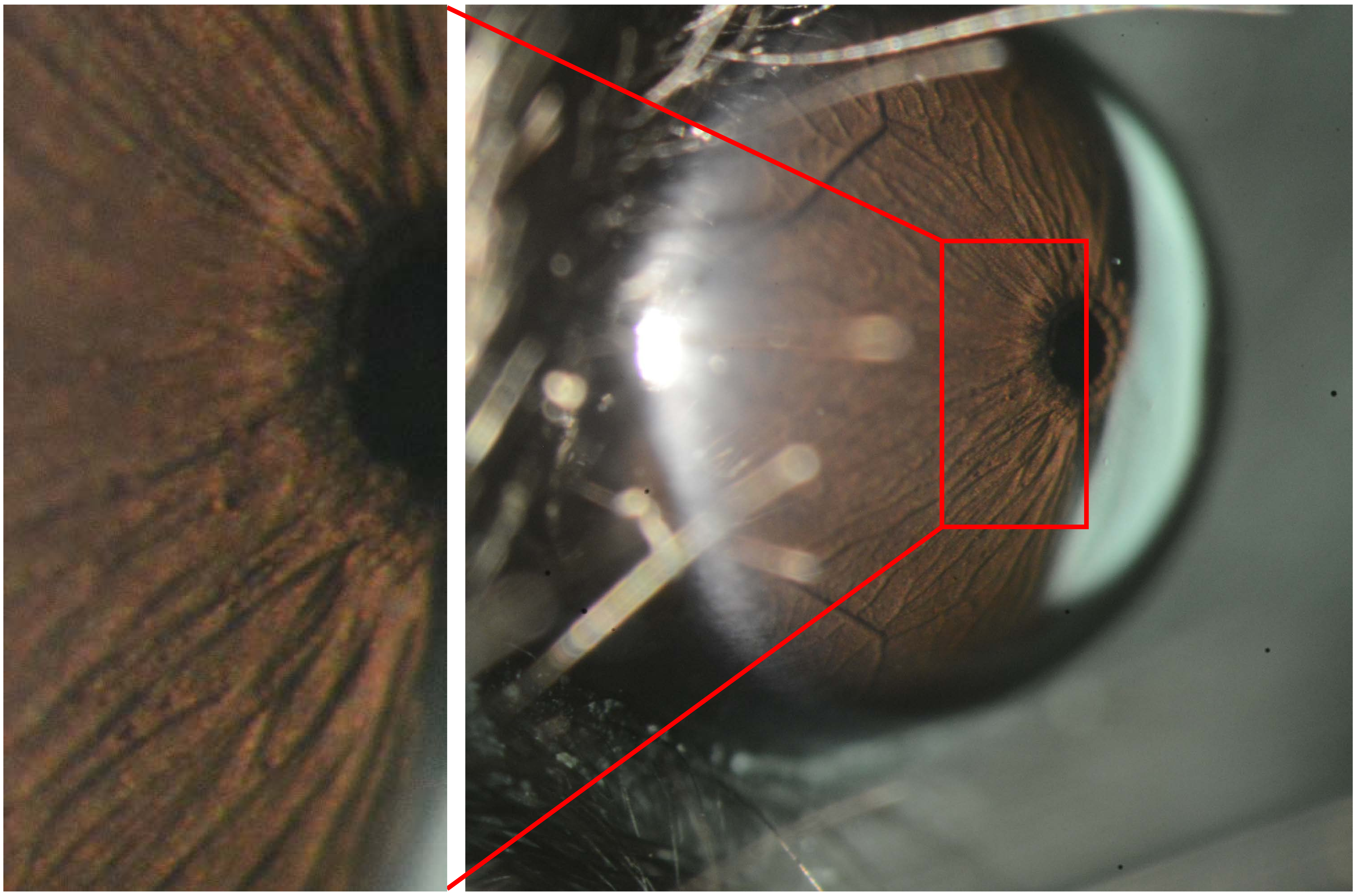

**OD**

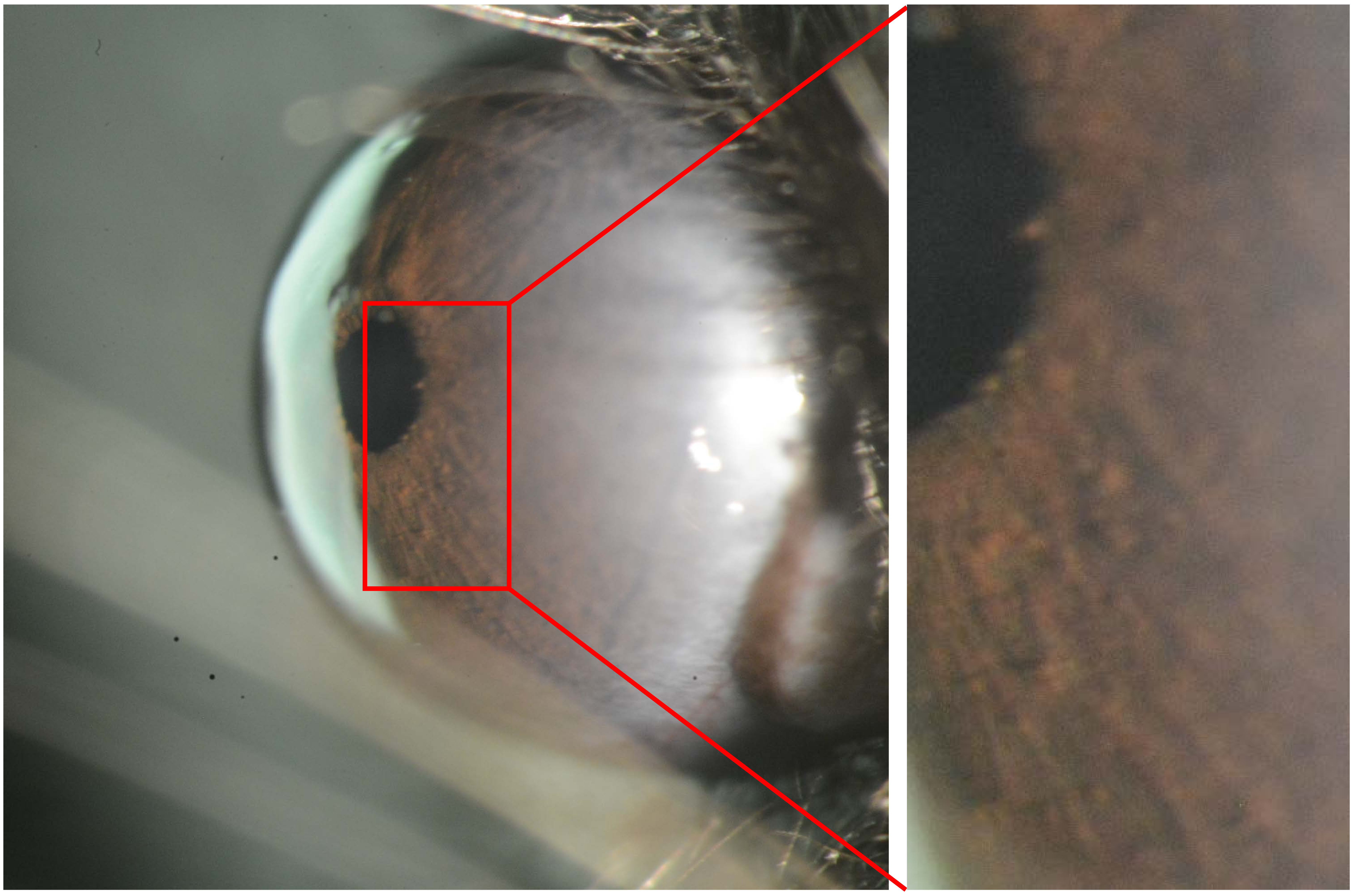
