## Appendix 11 Buffer 25X for "Recombinant adenovirus causes prolonged mobilization of macrophages in the anterior chamber of mice"

Appendix 11
Mouse ID: 36543

Masked assignment: **Buffer**
Actual group: **Buffer** (OD injected)

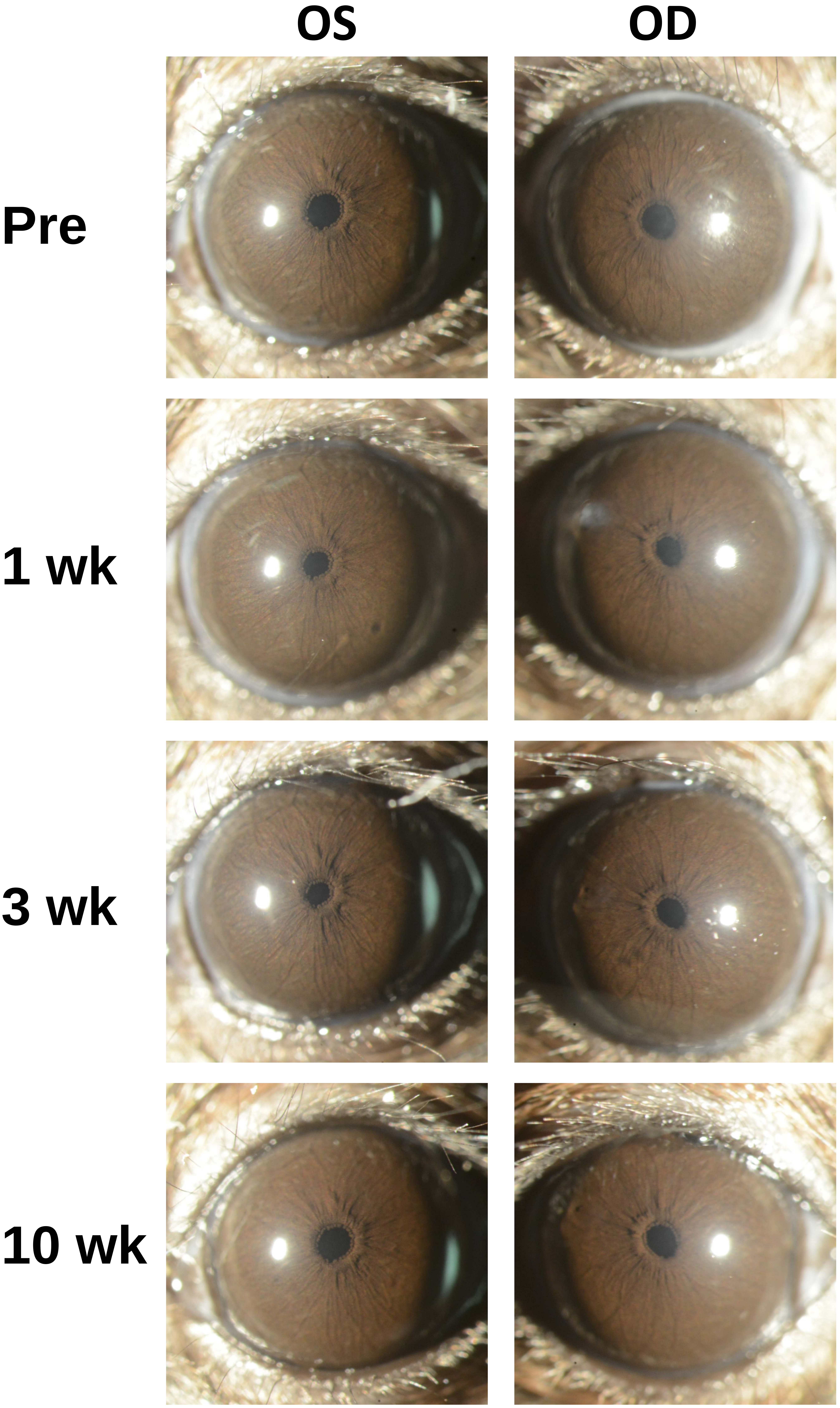

Appendix 11
Mouse ID: 36545

Masked assignment: **Buffer**
Actual group: **Buffer** (OS injected)

| | OS | OD |
|---|---|---|
| **Pre** | | |
| **1 wk** | | |
| **3 wk** | | |
| **10 wk** | | |

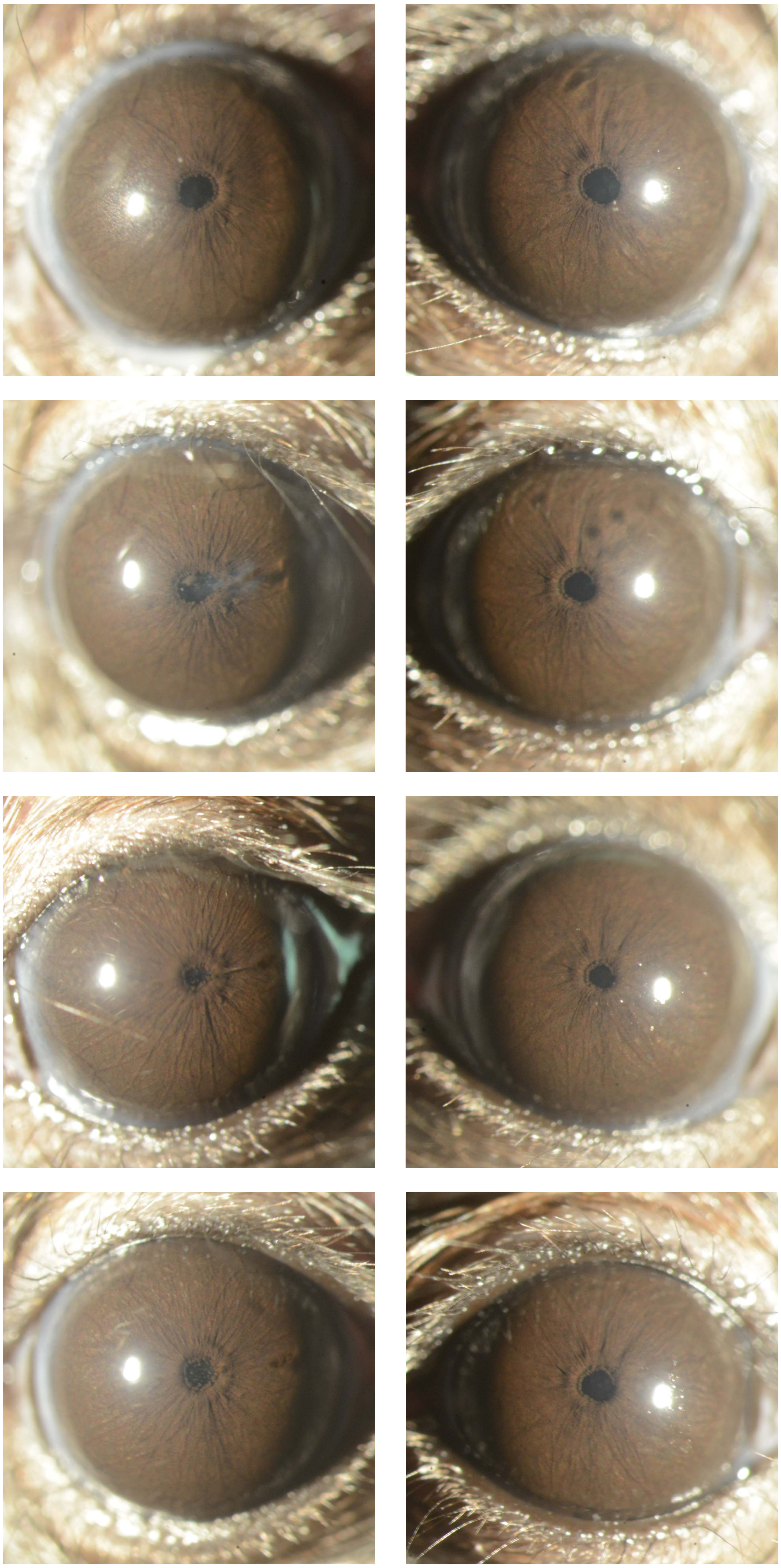

Appendix 11
Mouse ID: 36553

Masked assignment: **Buffer**
Actual group: **Buffer** (OS injected)

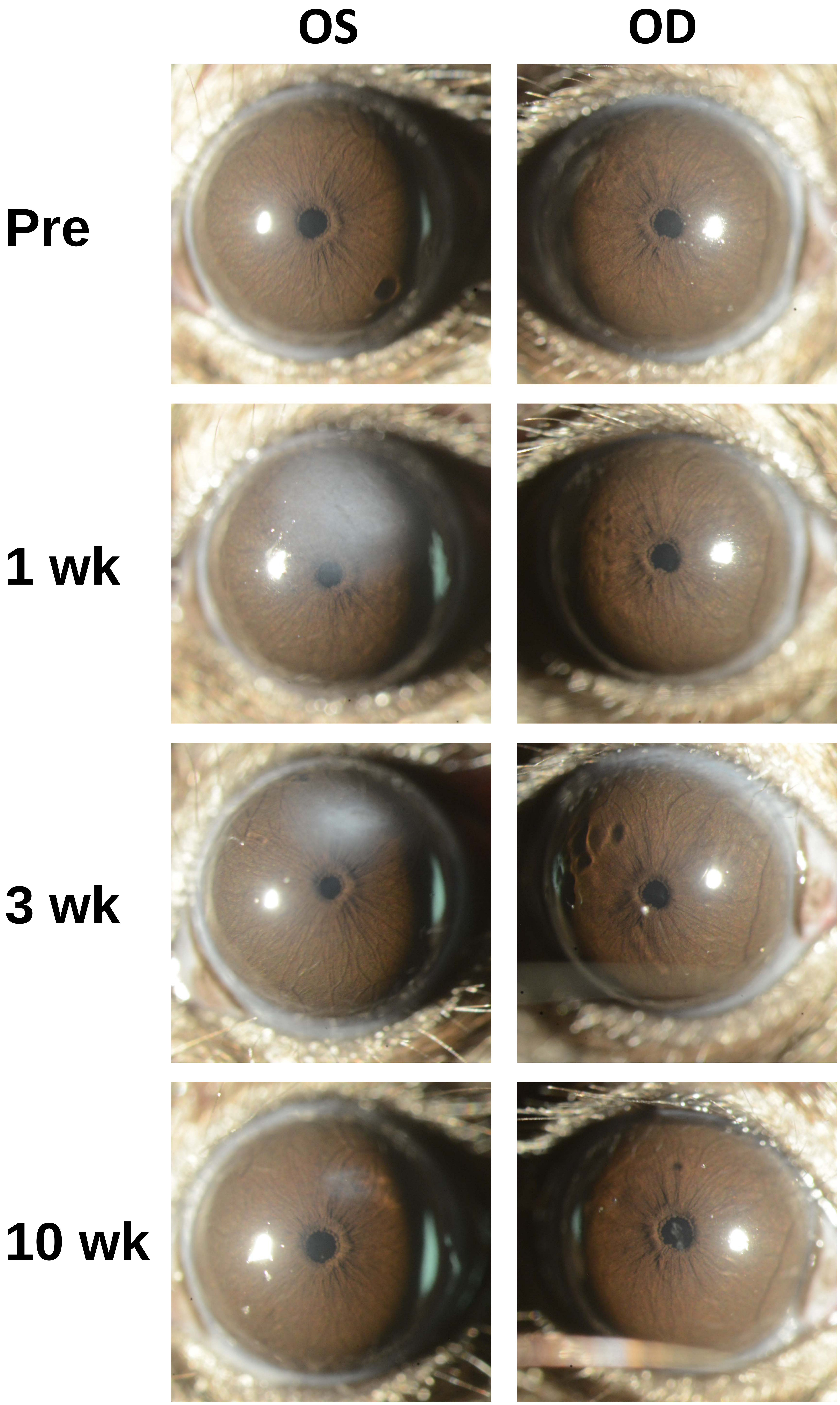

Appendix 11
Mouse ID: 36547

Masked assignment: **Buffer**
Actual group: **Buffer** (OD injected)

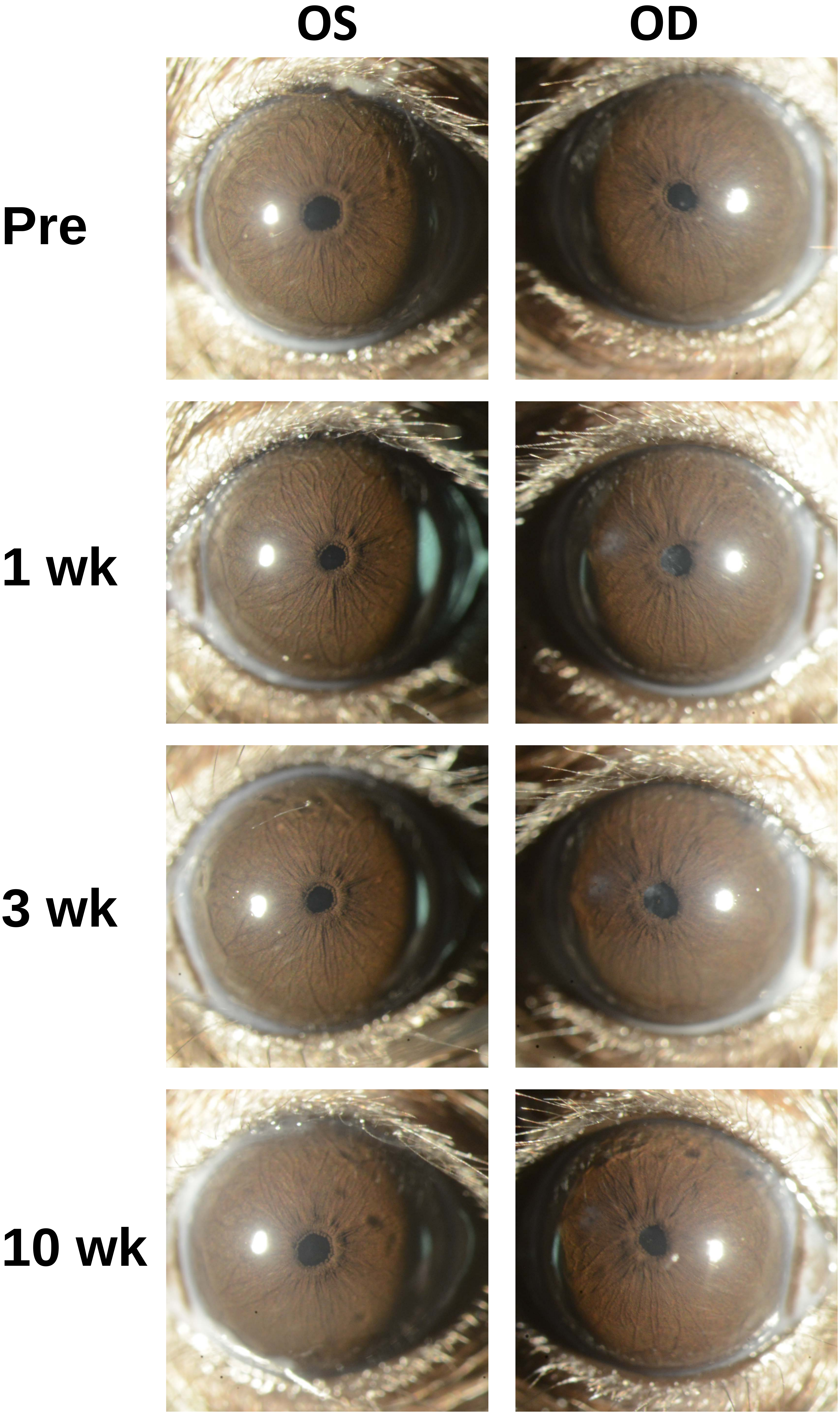
