## Appendix 12 Buffer 40X for "Recombinant adenovirus causes prolonged mobilization of macrophages in the anterior chamber of mice"

Appendix 12
Mouse ID: 36543
10-week timepoint

Masked assignment: **Buffer**
Actual group: **Buffer** (OD injected)

**OS**

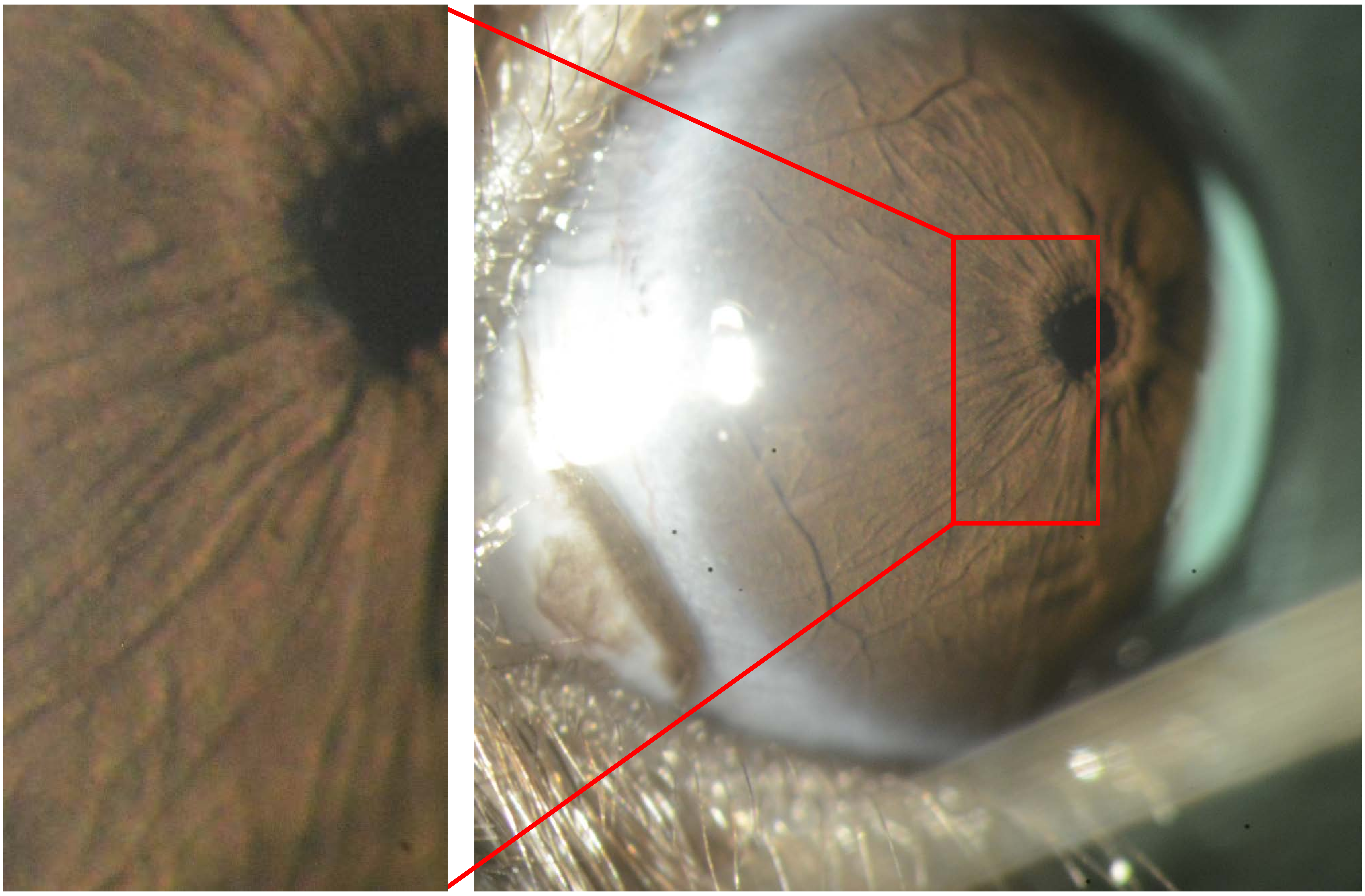

**OD**

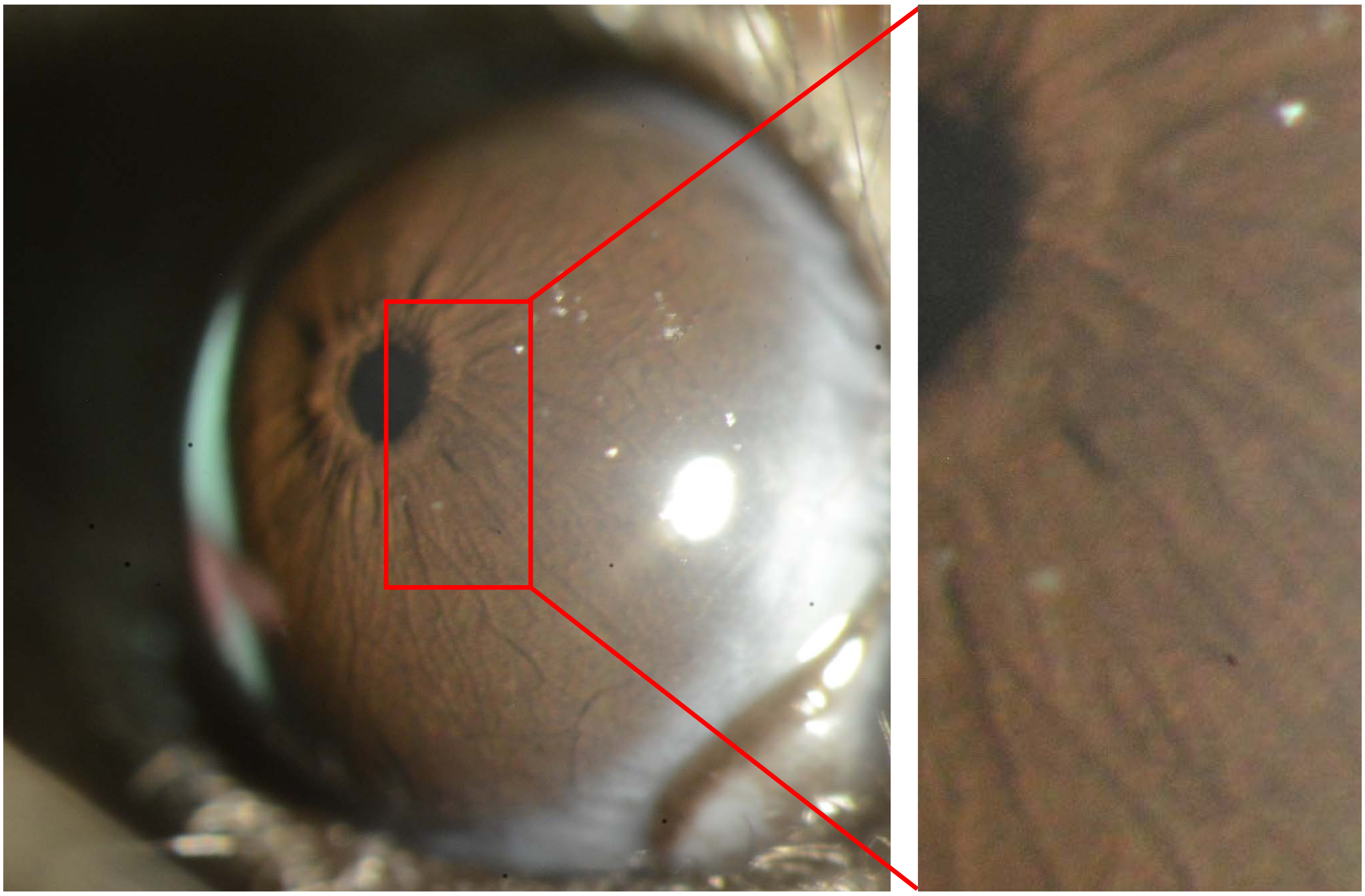

Appendix 12
Mouse ID: 36545
10-week timepoint

Masked assignment: **Buffer**
Actual group: **Buffer** (OS injected)

**OS**

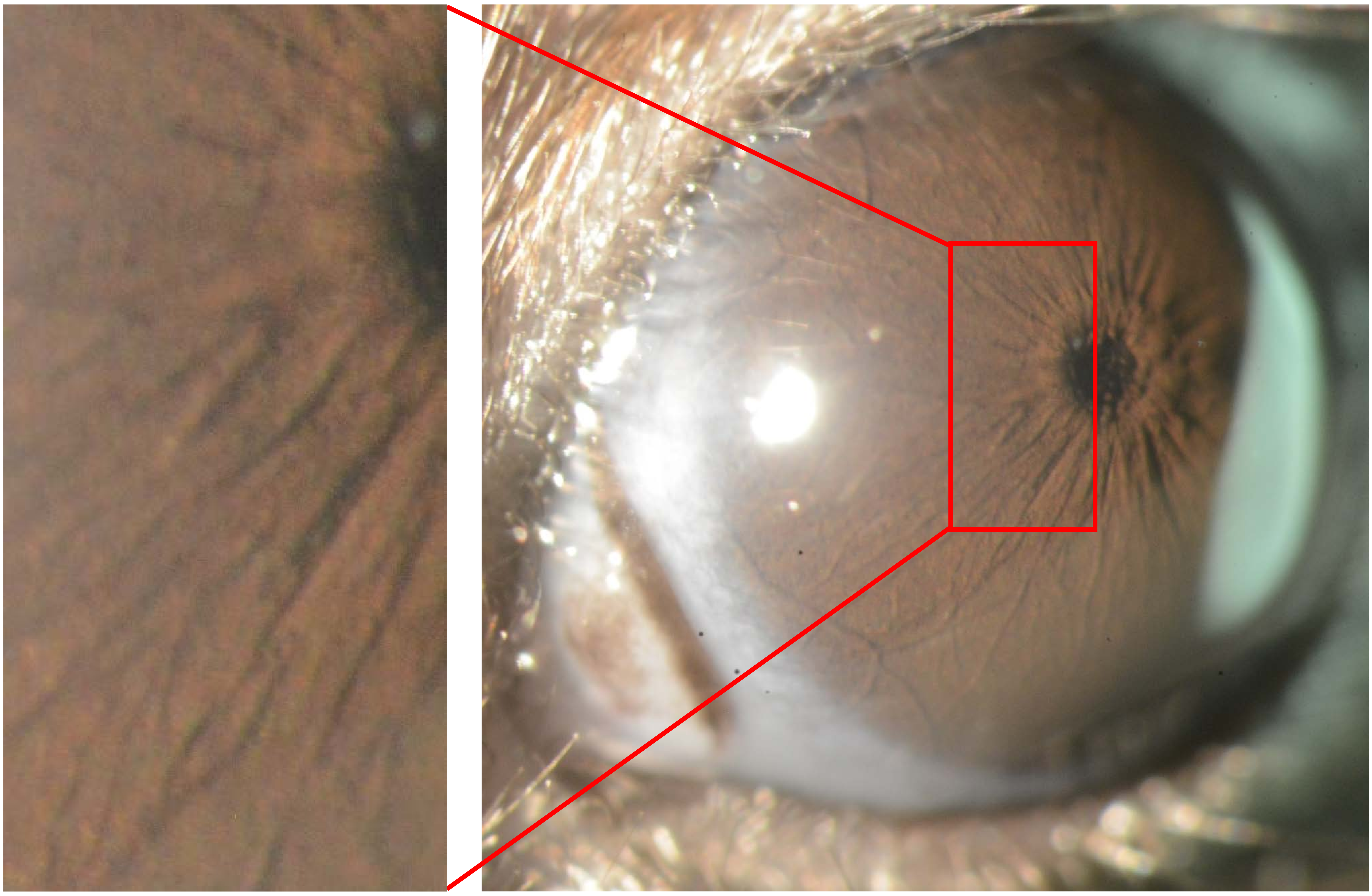

**OD**

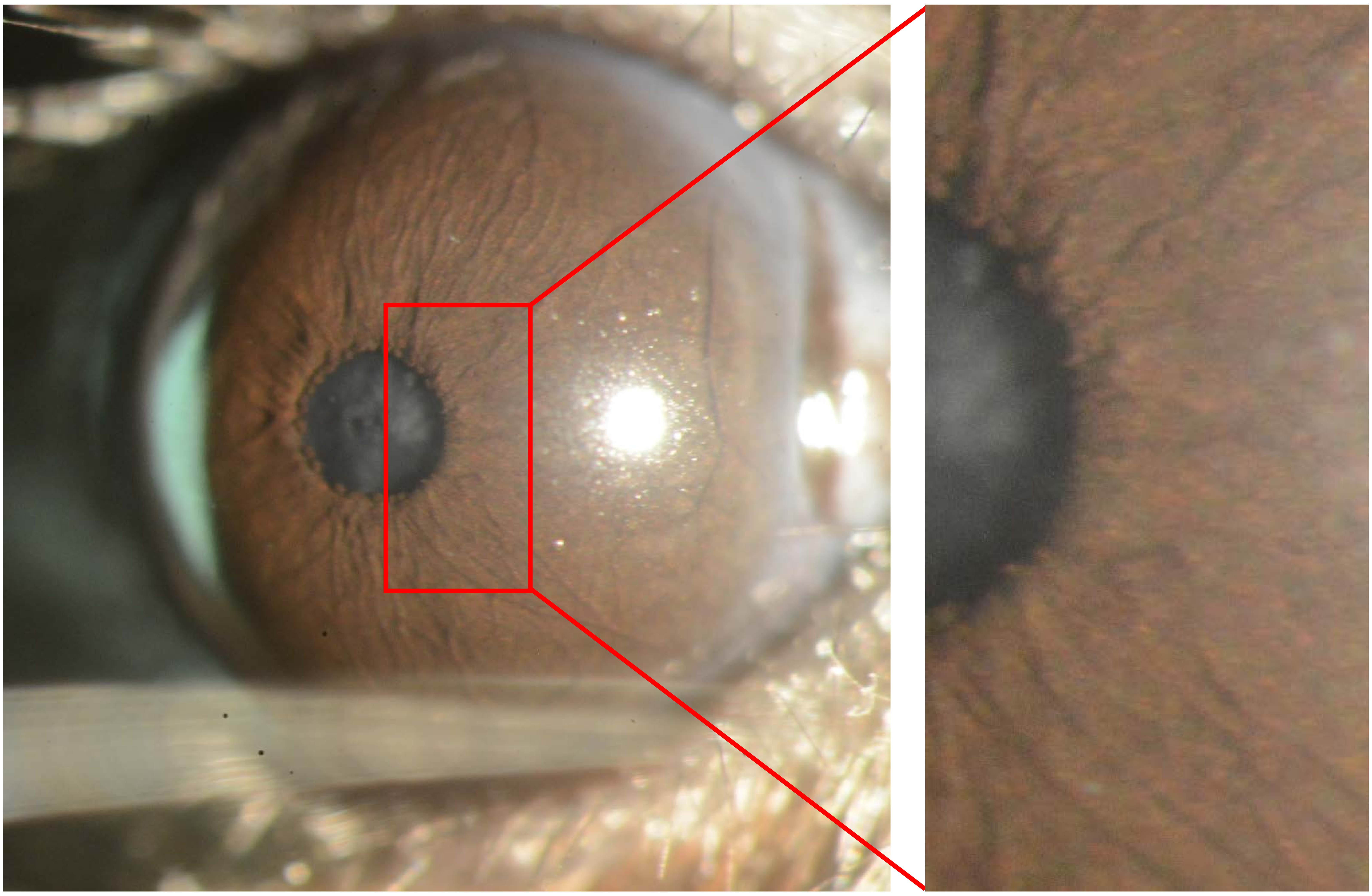

Appendix 12
Mouse ID: 36553
10-week timepoint

Masked assignment: **Buffer**
Actual group: **Buffer** (OS injected)

**OS**

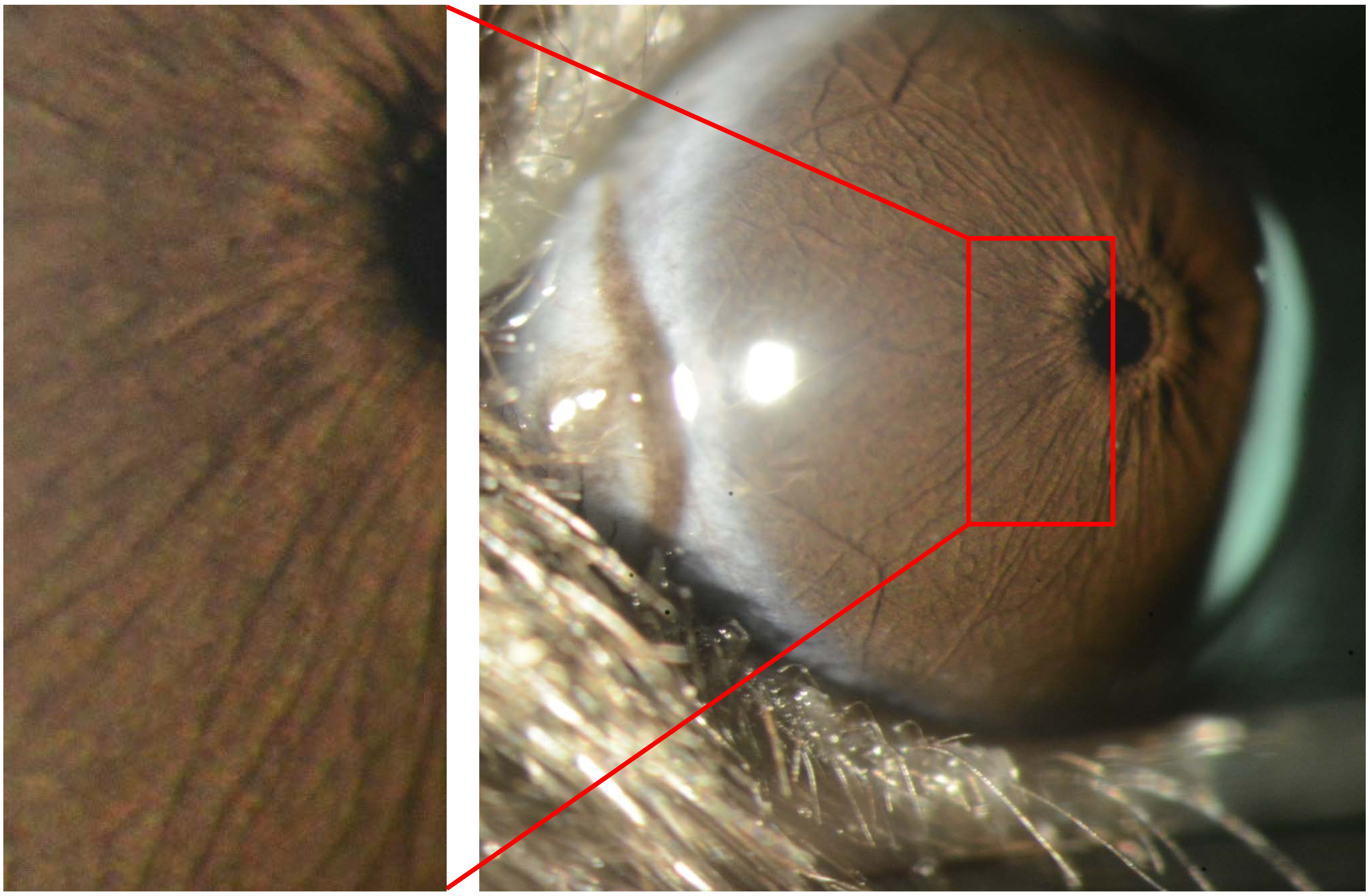

**OD**

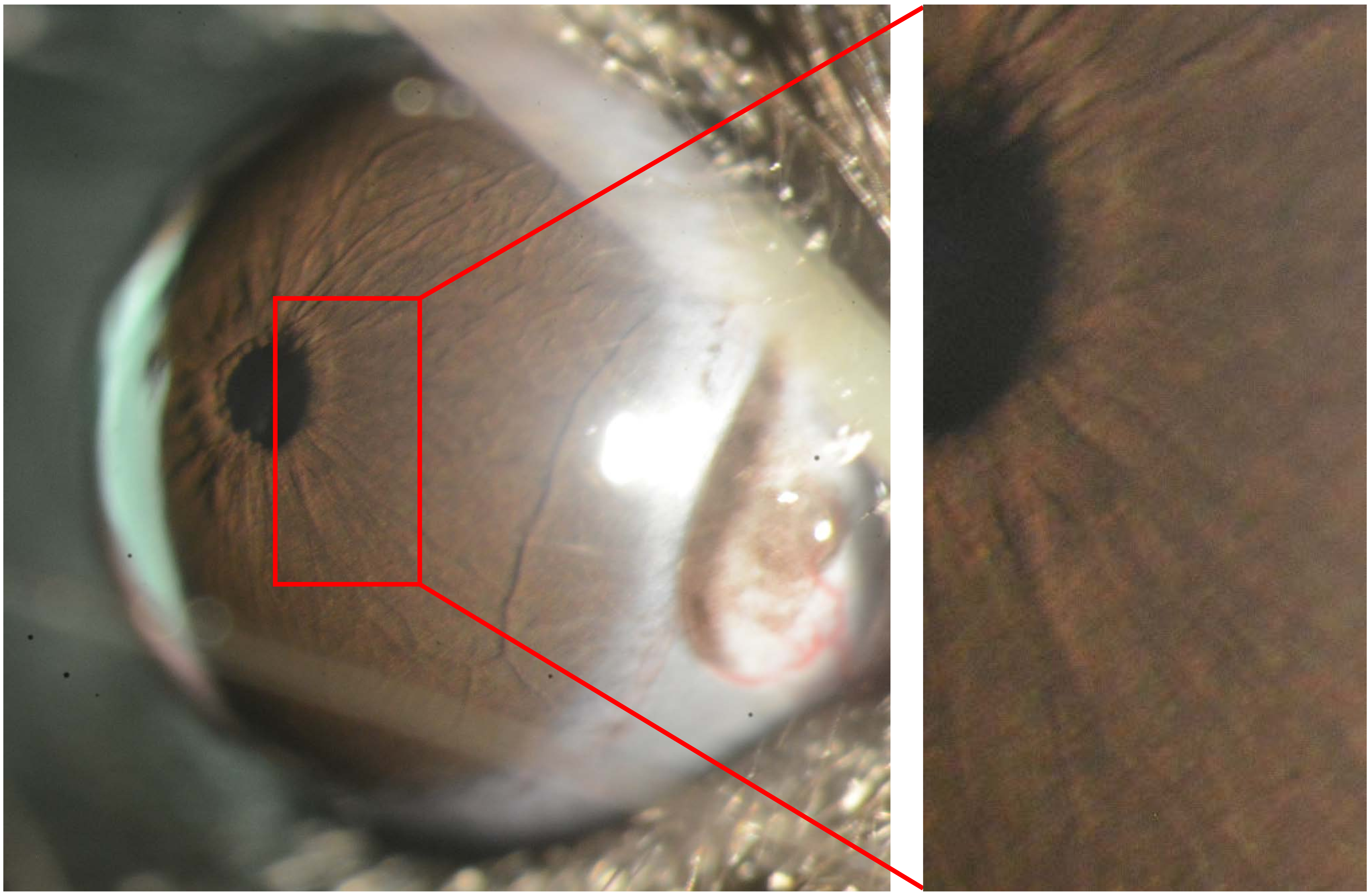

Appendix 12
Mouse ID: 36547
10-week timepoint

Masked assignment: **Buffer**
Actual group: **Buffer** (OD injected)

**OS**

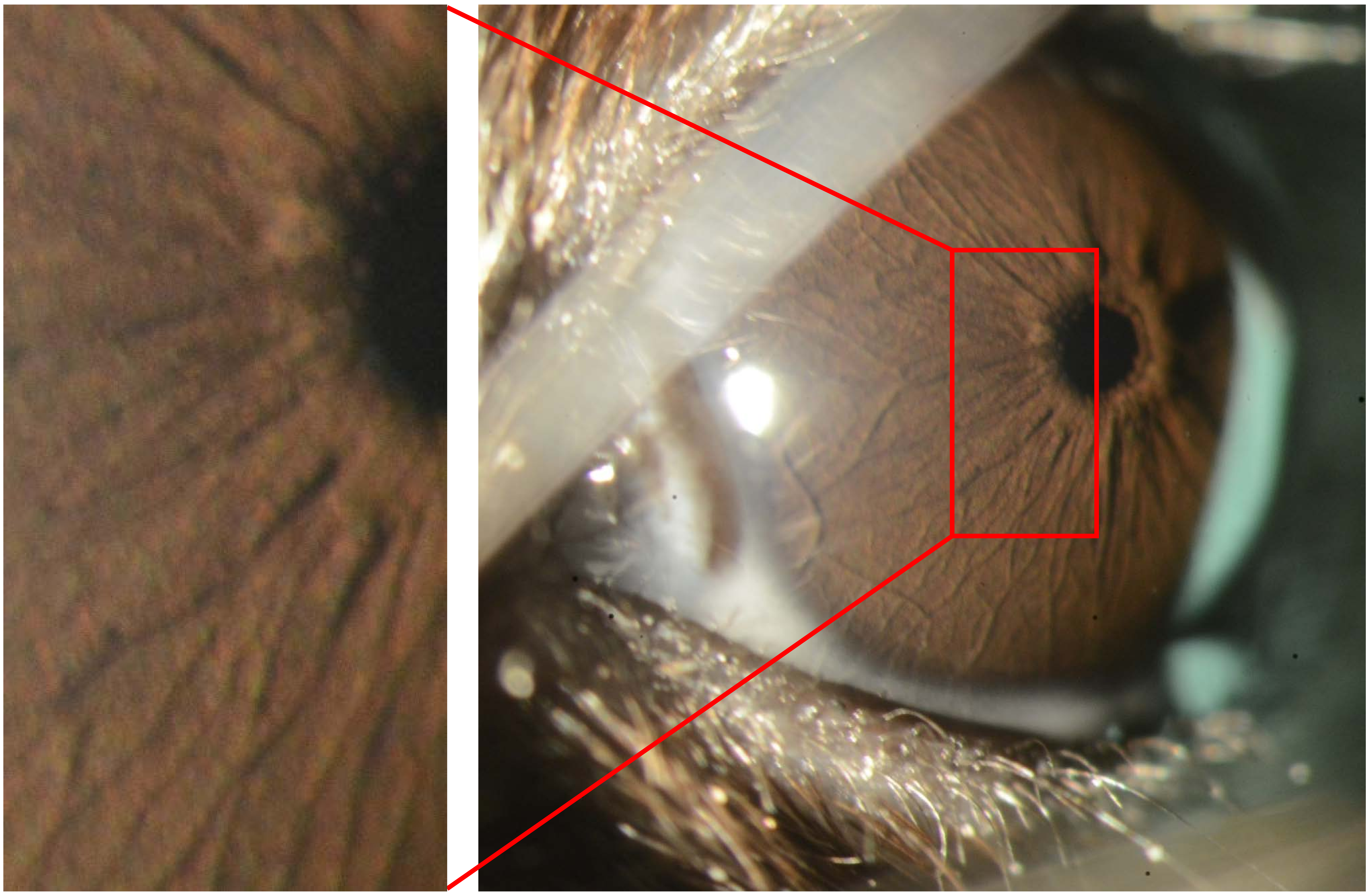

**OD**

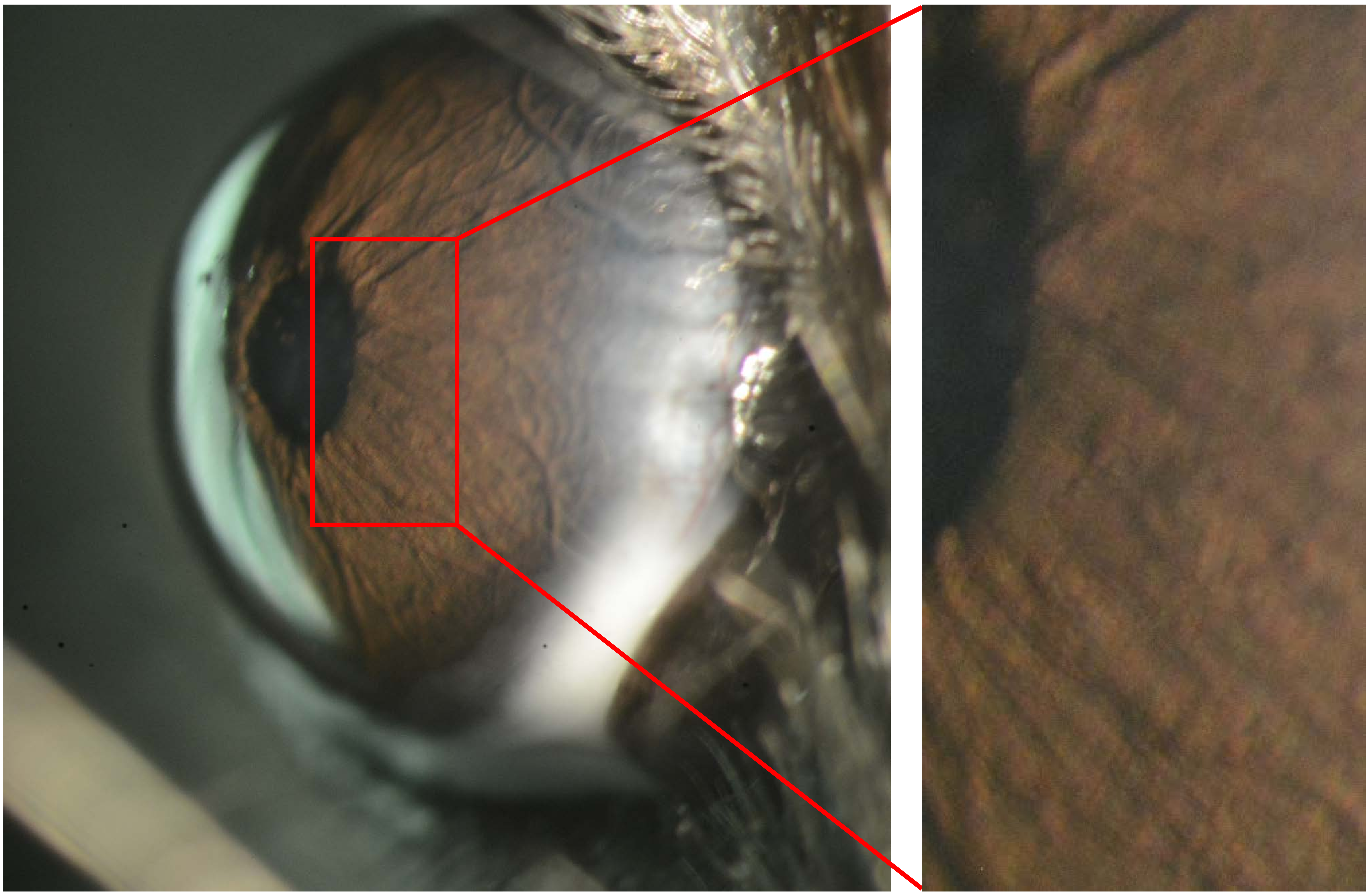
