## Appendix 13 Intravitreal 25X for "Recombinant adenovirus causes prolonged mobilization of macrophages in the anterior chamber of mice"

Appendix 13
Mouse ID: 36542

Masked assignment: **Intravitreal**
Actual group: **Intravitreal** (OS injected)

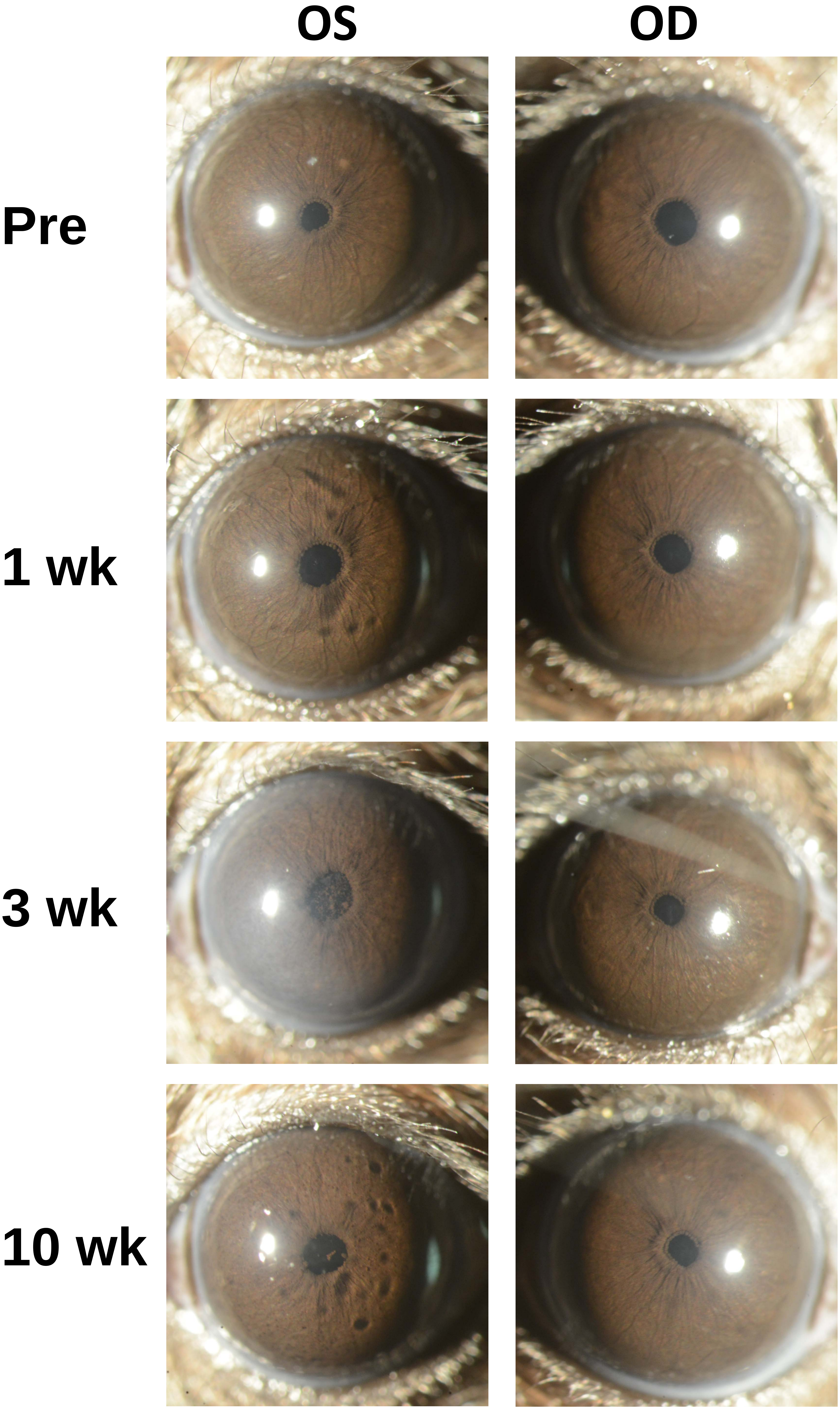

Appendix 13
Mouse ID: 36548

Masked assignment: **Intravitreal**
Actual group: **Intravitreal** (OD injected)

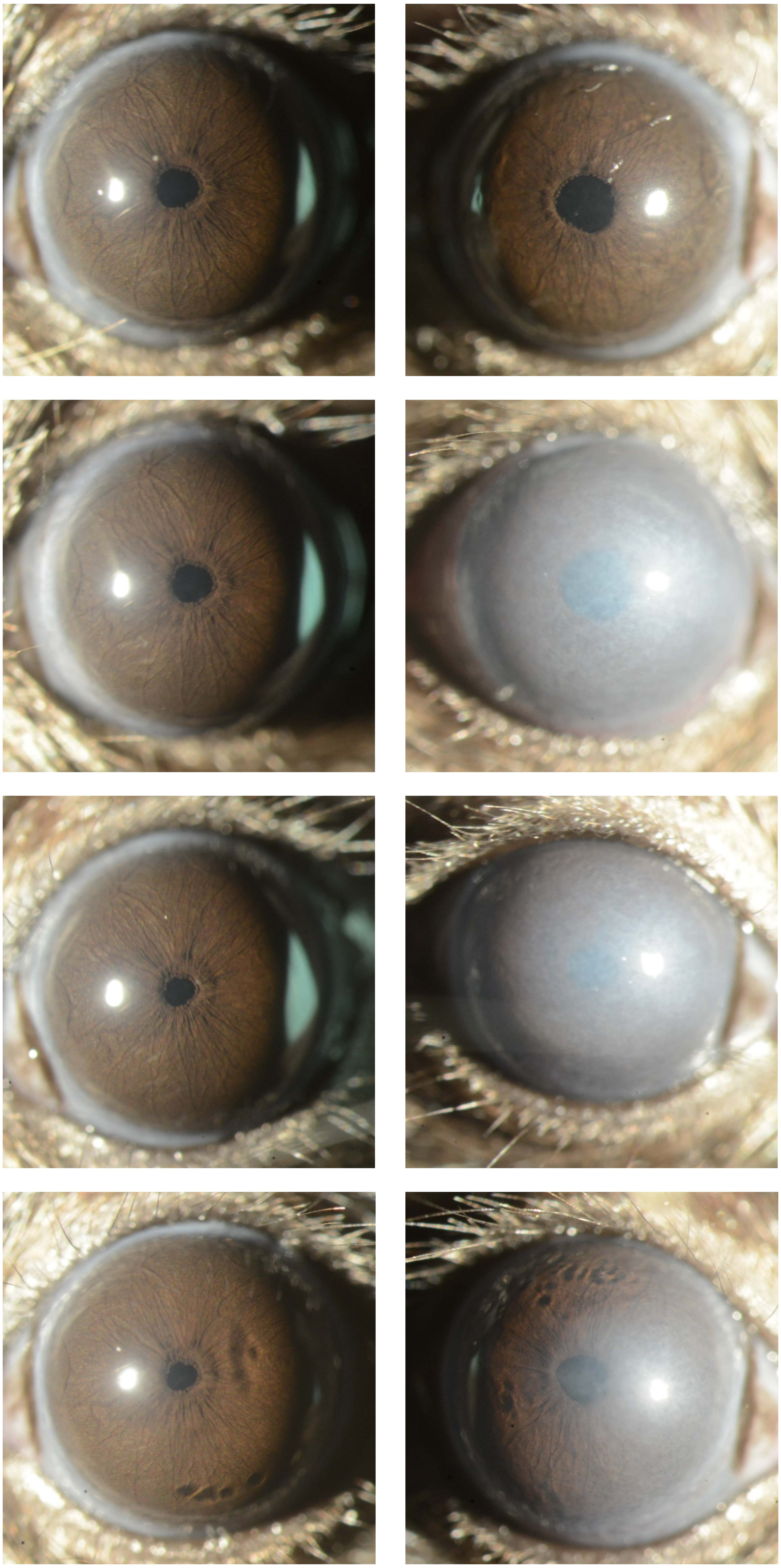

Appendix 13
Mouse ID: 36551

Masked assignment: **Intravitreal**
Actual group: **Intravitreal** (OS injected)

**OS** **OD**

**Pre**

**1 wk**

**3 wk**

**10 wk**

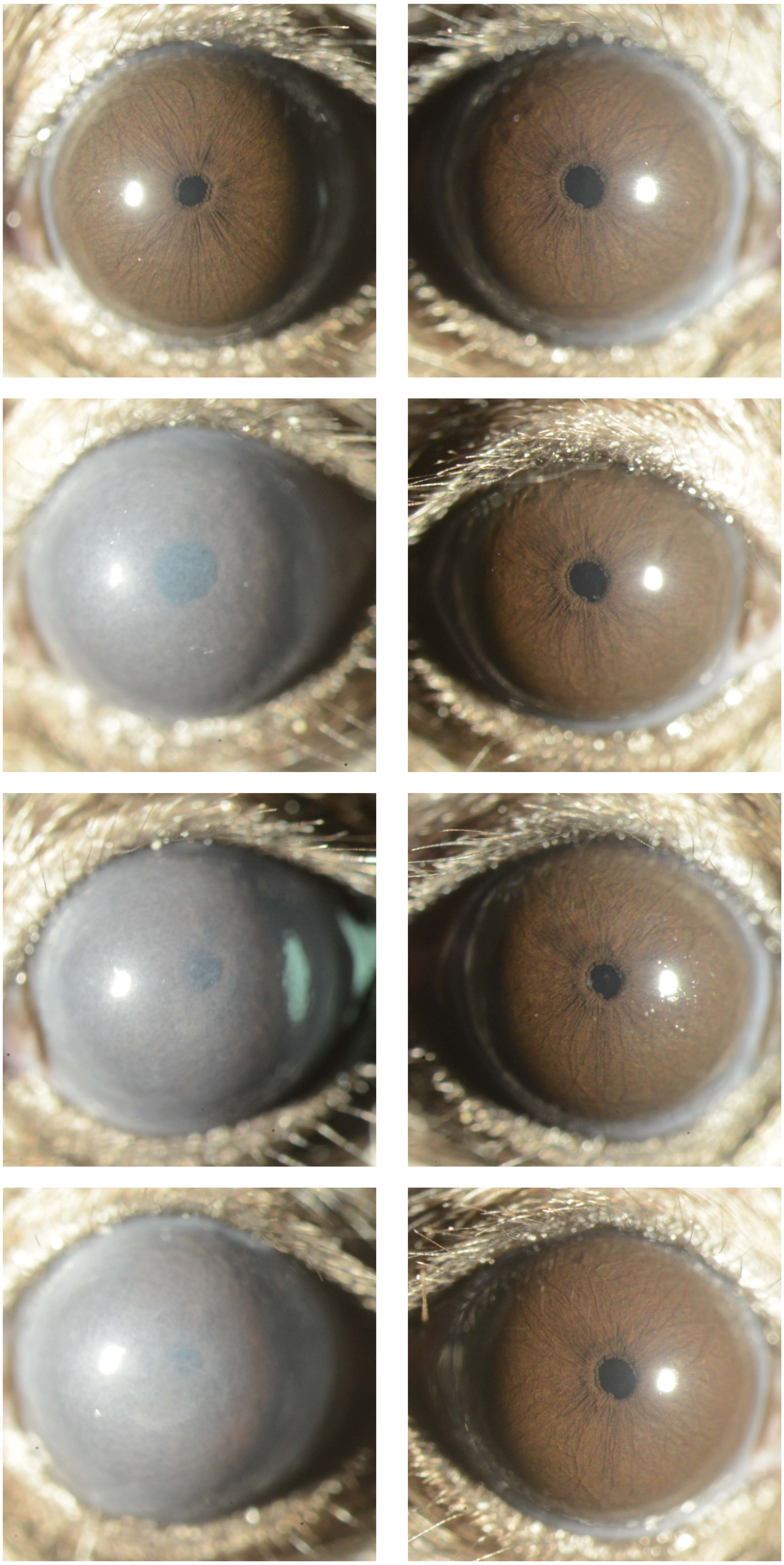

Appendix 13
Mouse ID: 36550

Masked assignment: **Intravitreal**
Actual group: **Intravitreal** (OD injected)

**OS** **OD**

**Pre**

**1 wk**

**3 wk**

**10 wk**

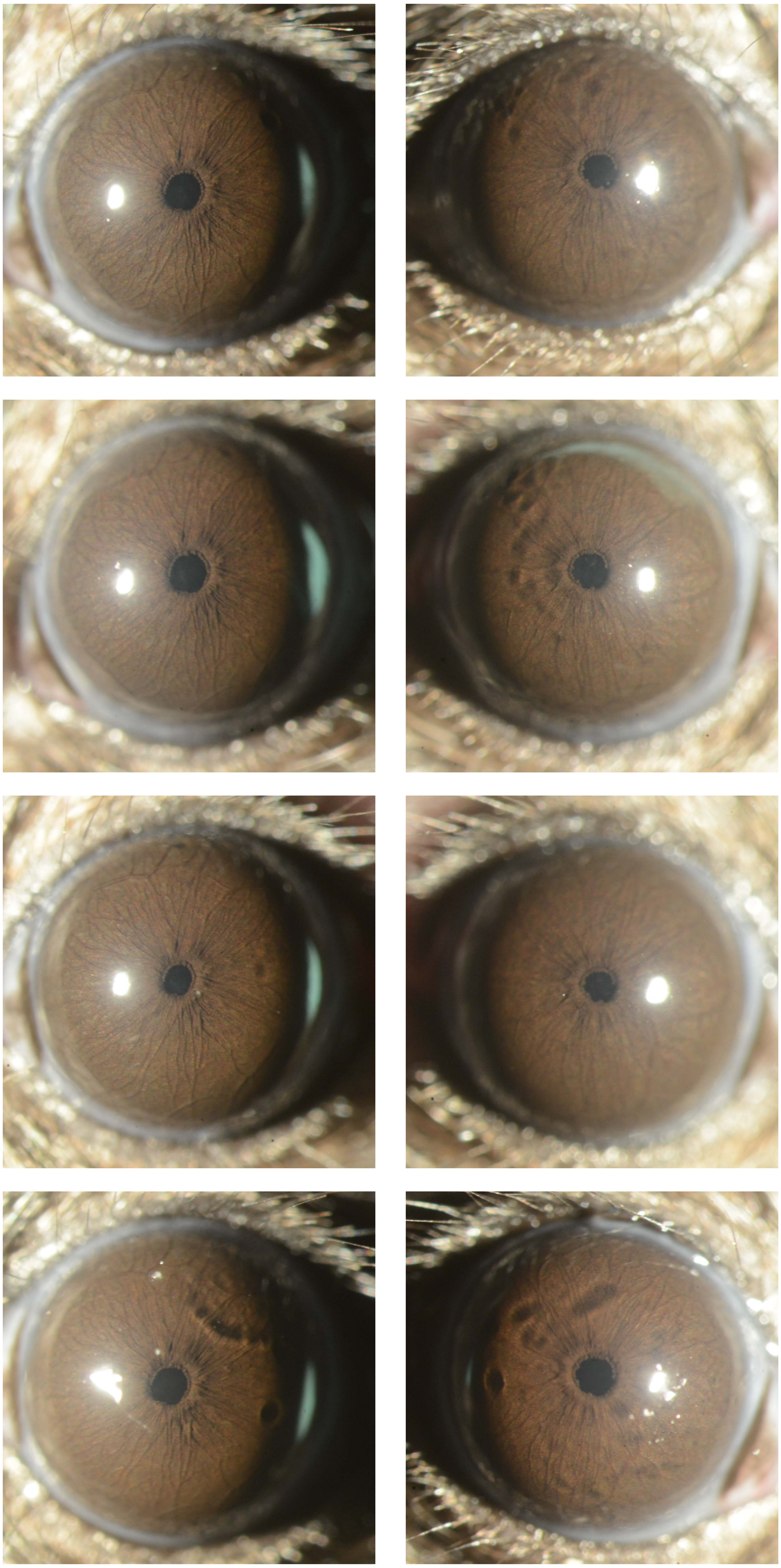
