## Appendix 14 Intravitreal 40X for "Recombinant adenovirus causes prolonged mobilization of macrophages in the anterior chamber of mice"

Appendix 14
Mouse ID: 36542
10-week timepoint

Masked assignment: **Intravitreal**
Actual group: **Intravitreal** (OS injected)

**OS**

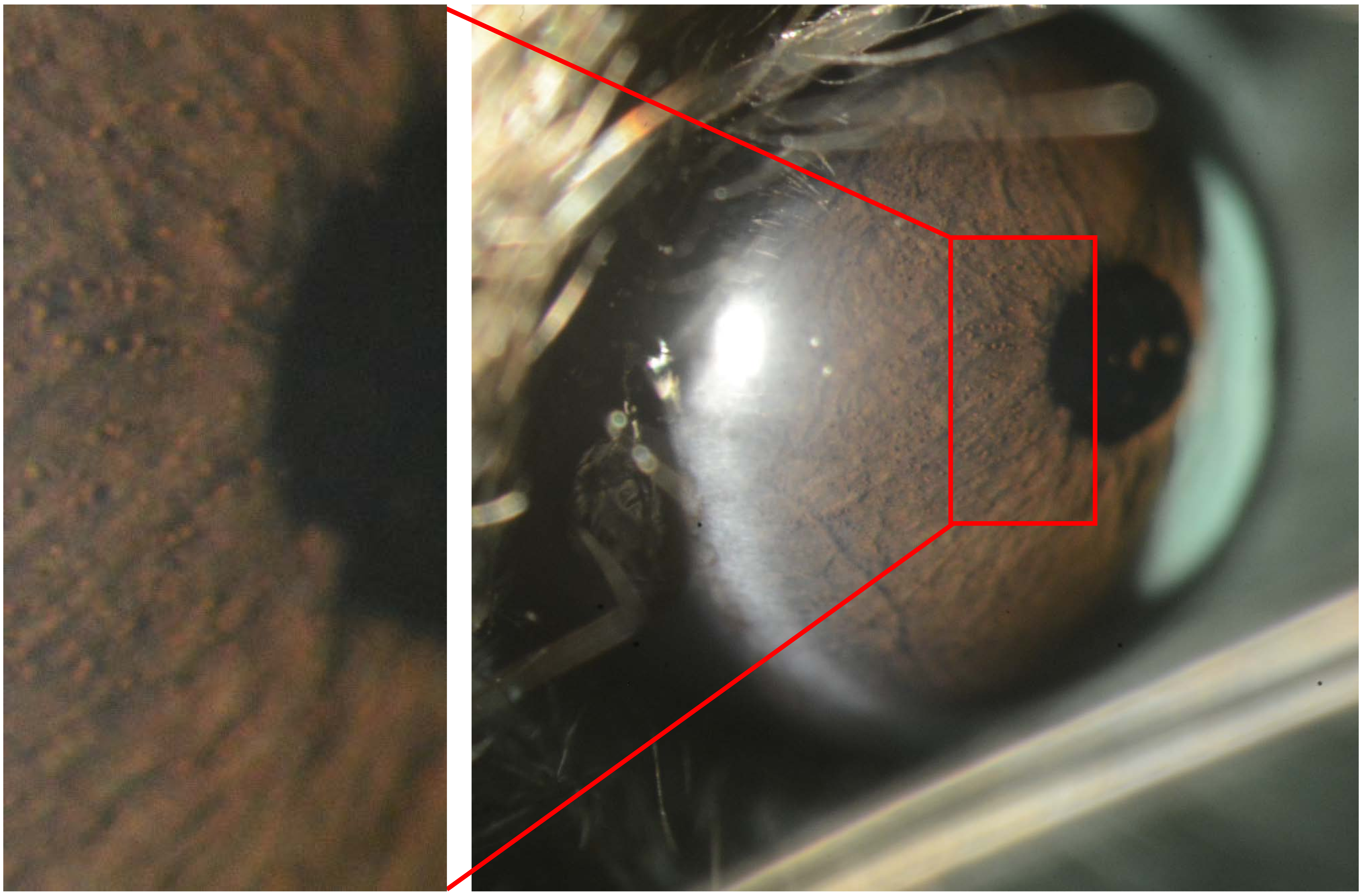

**OD**

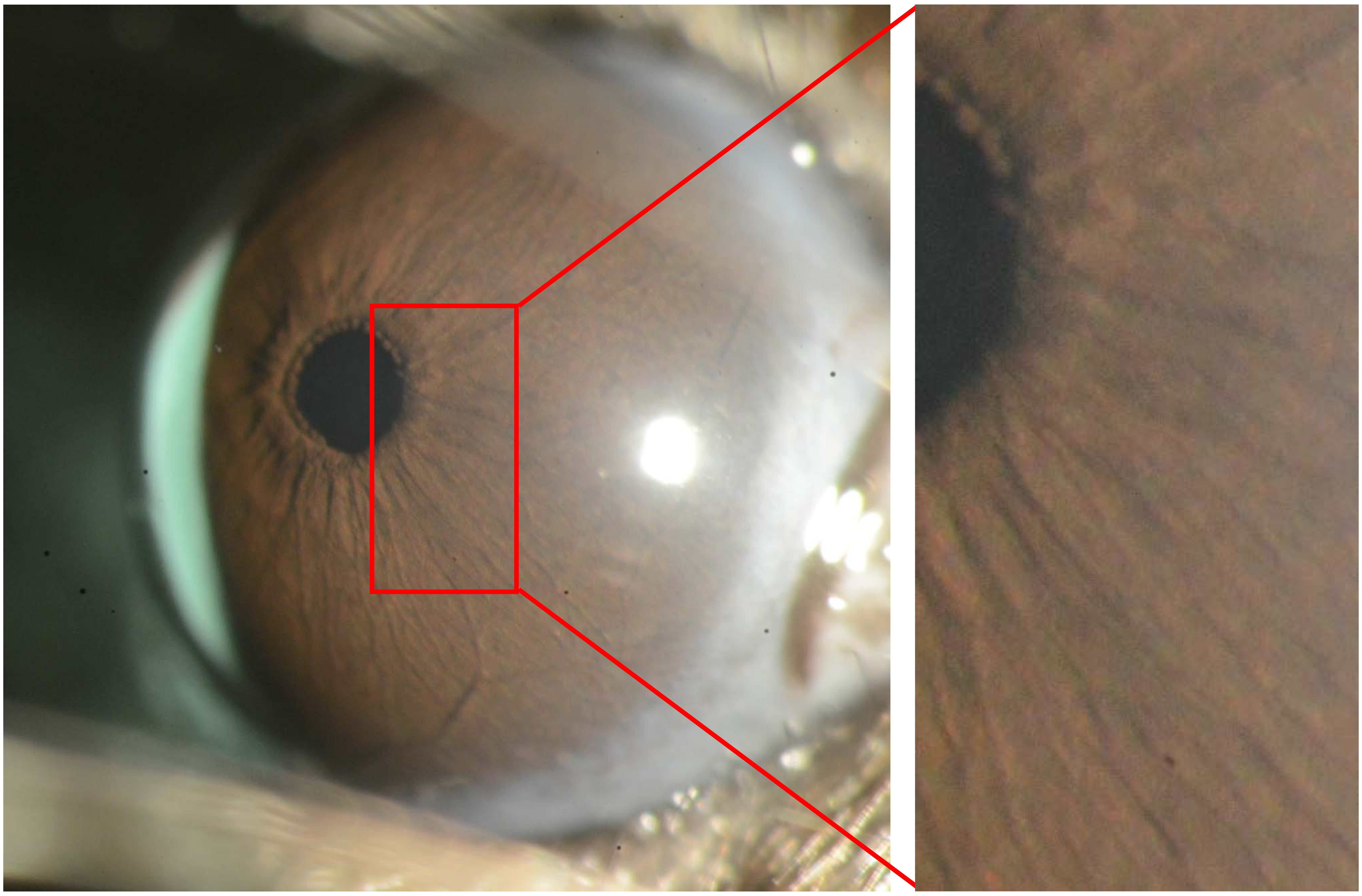

Appendix 14
Mouse ID: 36548
10-week timepoint

Masked assignment: **Intravitreal**
Actual group: **Intravitreal** (OD injected)

**OS**

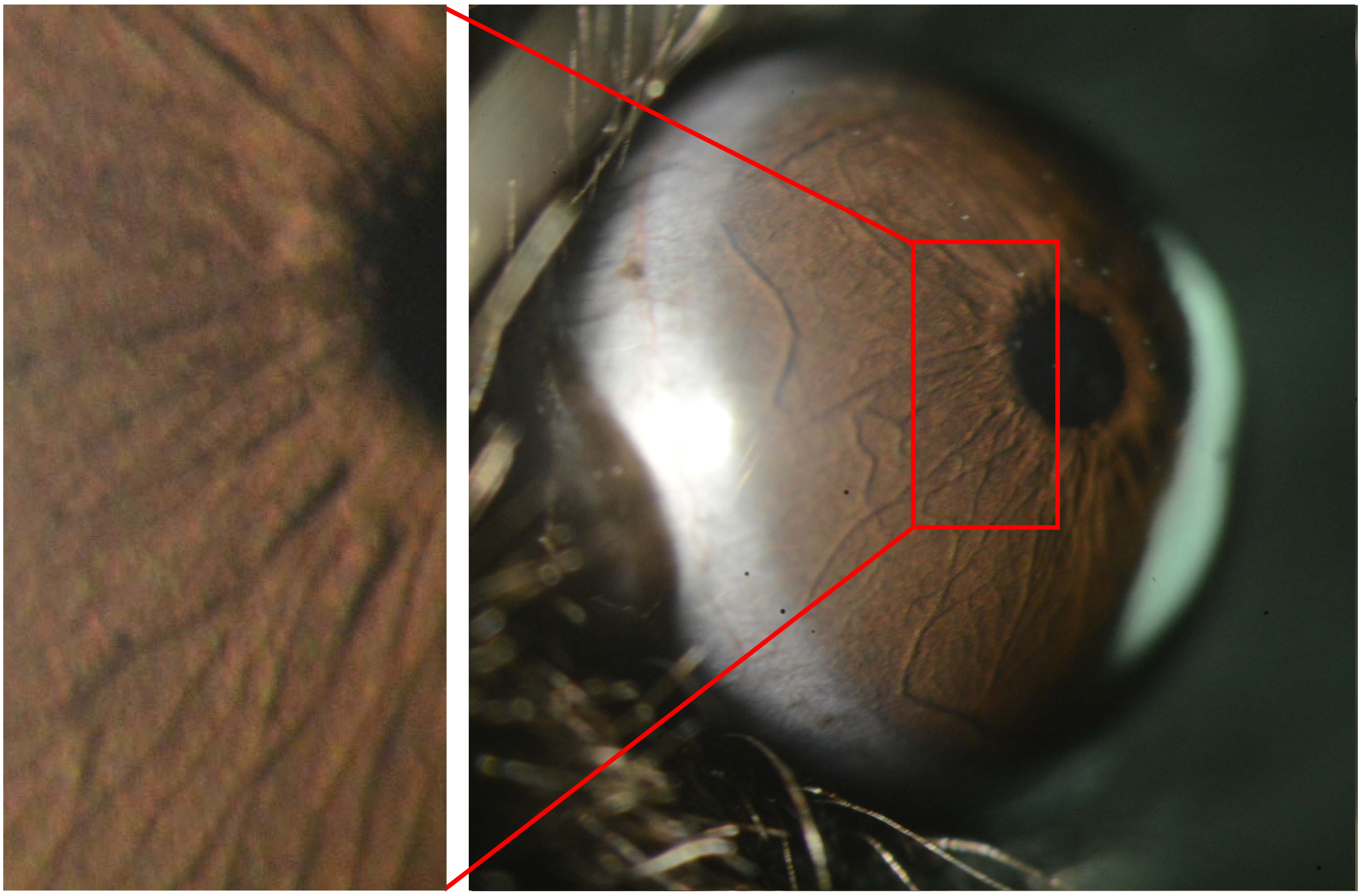

**OD**

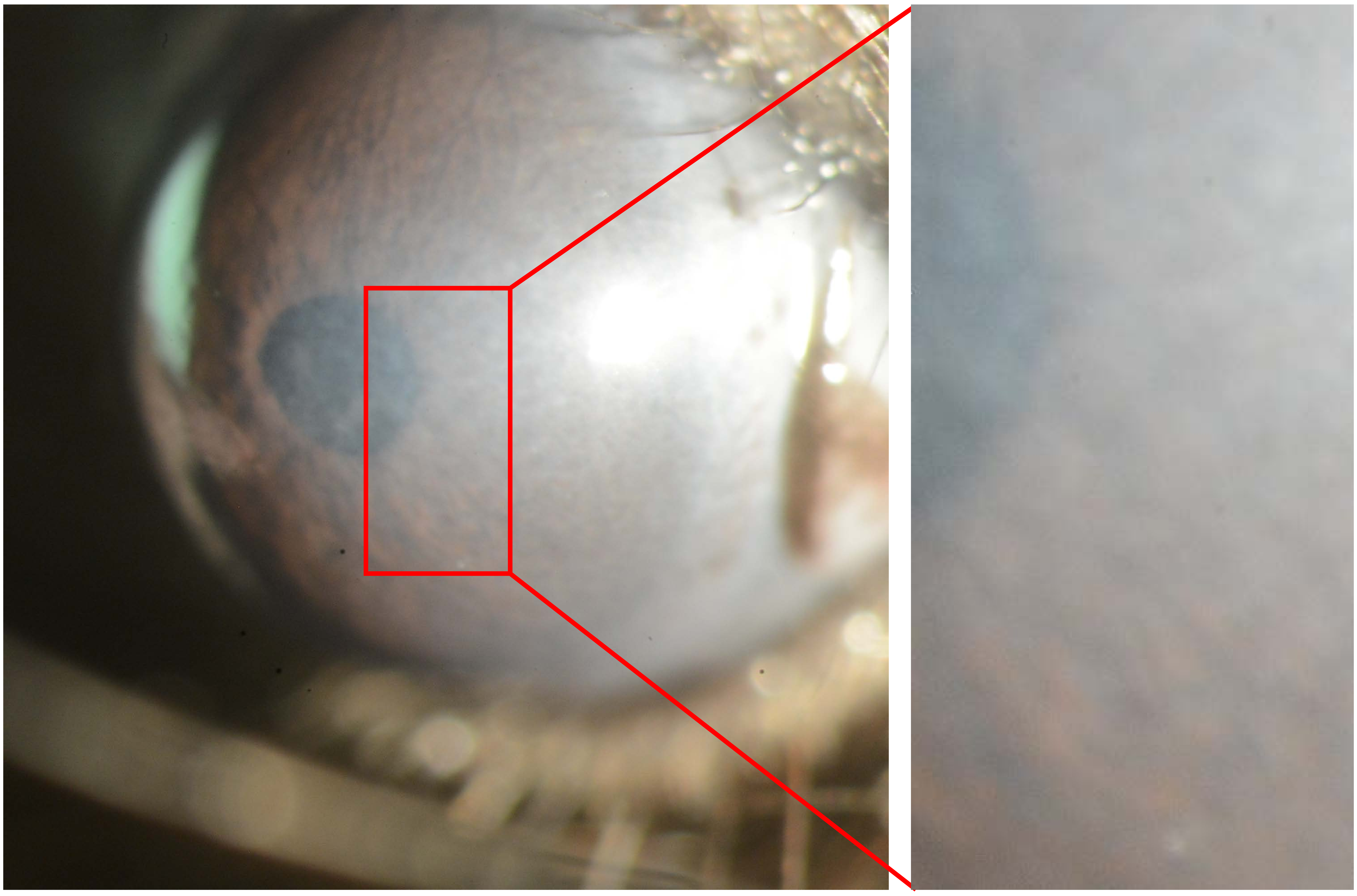

Appendix 14
Mouse ID: 36551
10-week timepoint

Masked assignment: **Intravitreal**
Actual group: **Intravitreal** (OS injected)

**OS**

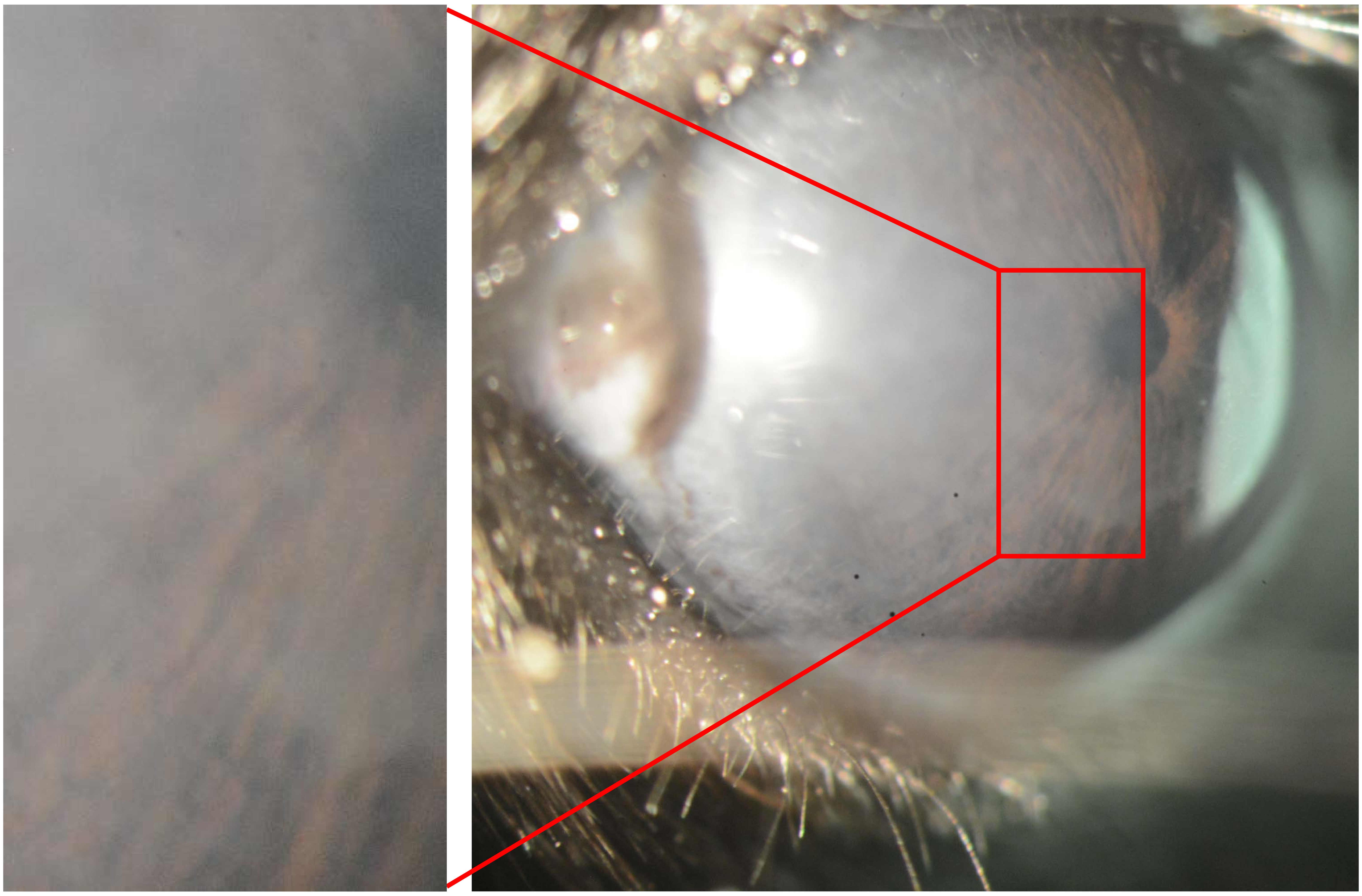

**OD**

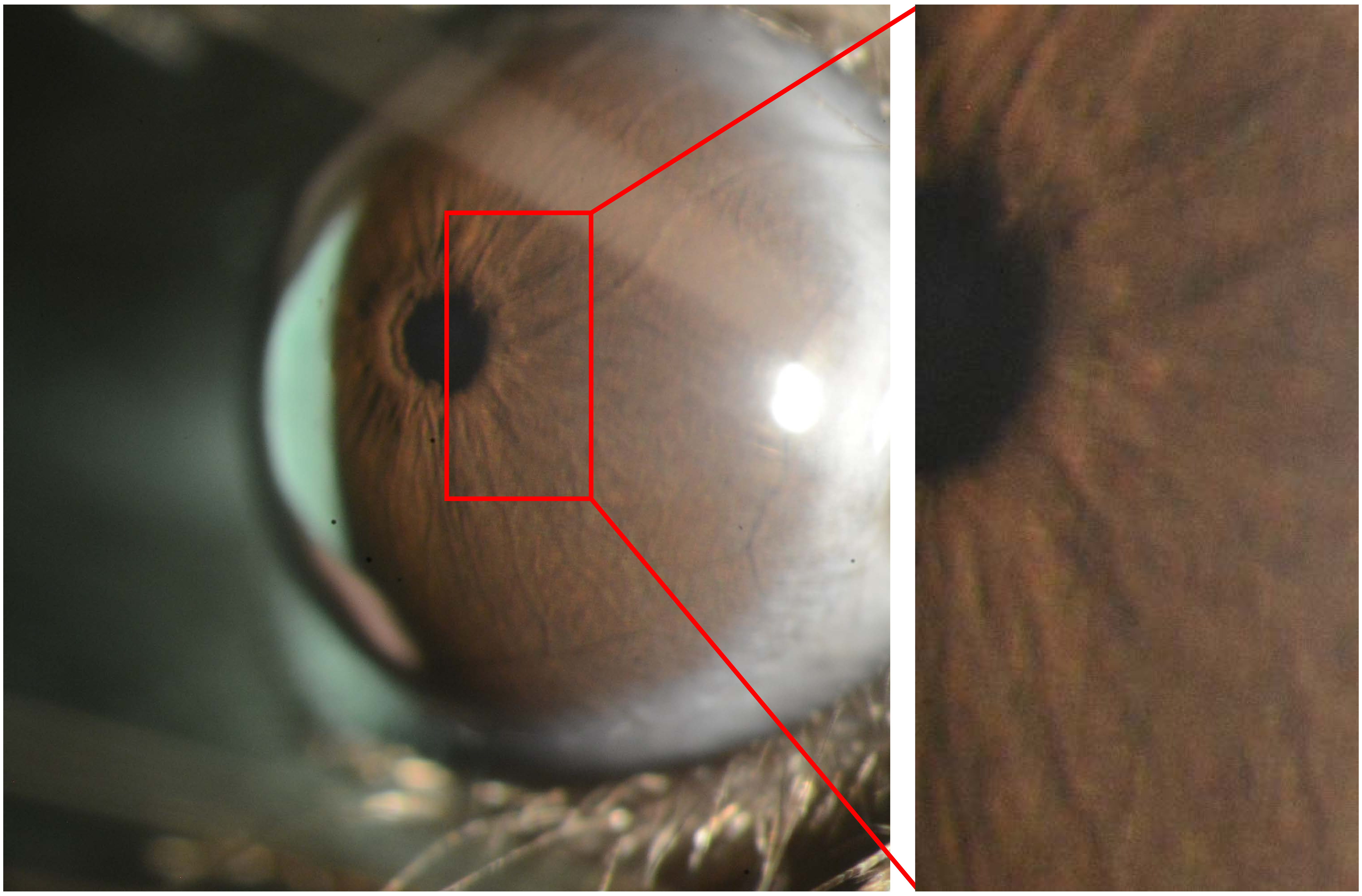

Appendix 14
Mouse ID: 36550
10-week timepoint

Masked assignment: **Intravitreal**
Actual group: **Intravitreal** (OD injected)

**OS**

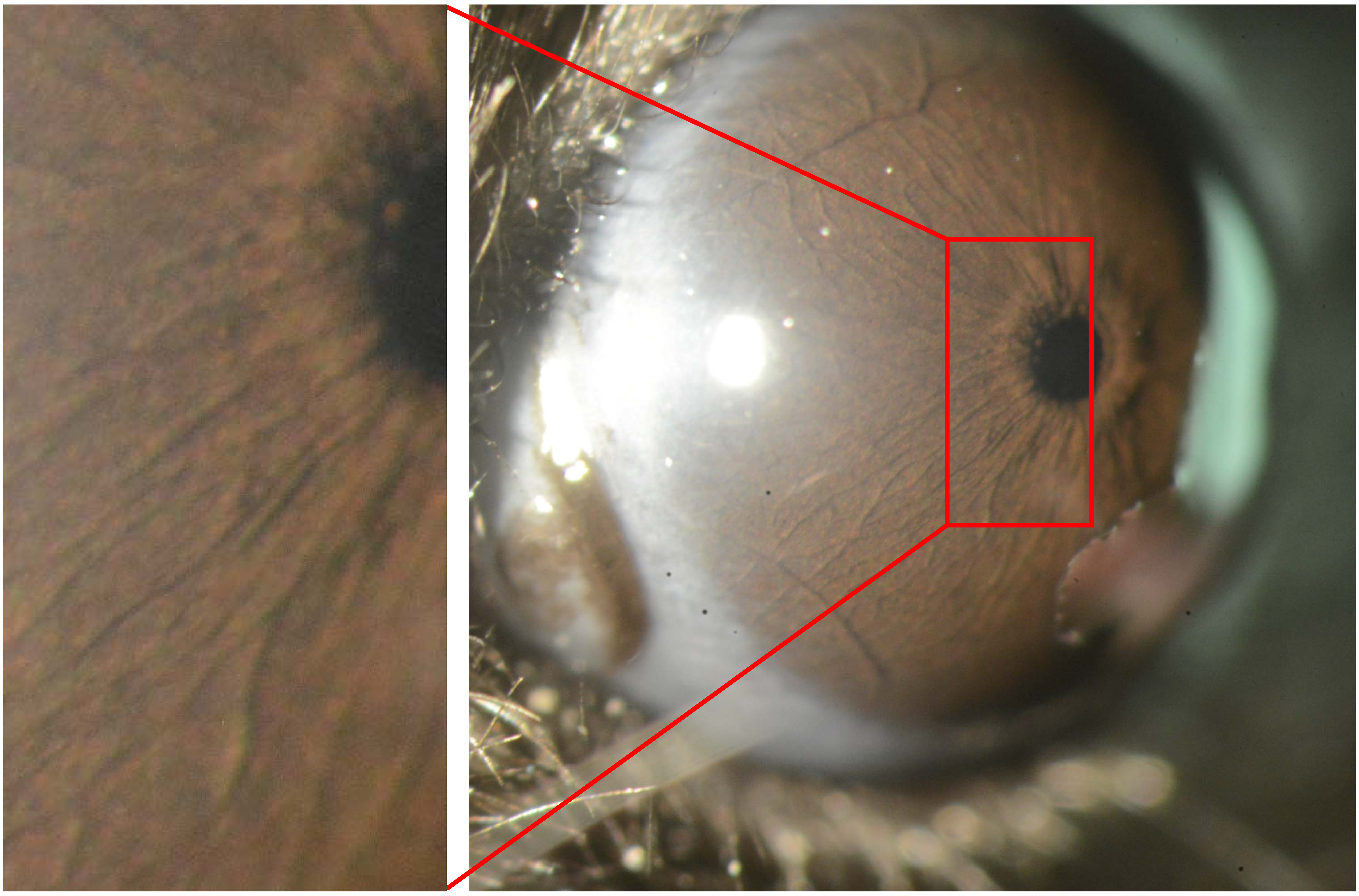

**OD**

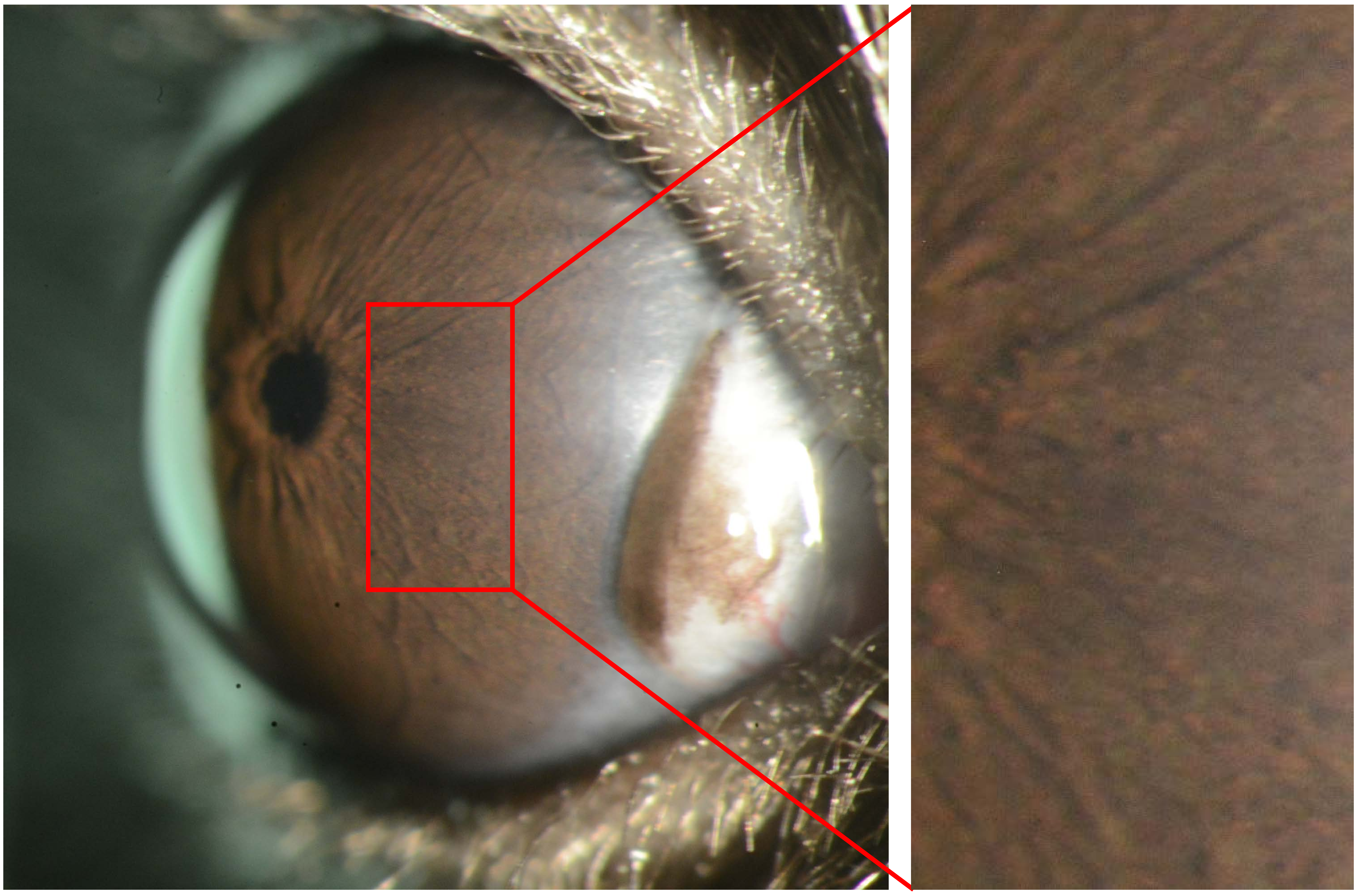
