## Appendix 15 Aged 25X for "Recombinant adenovirus causes prolonged mobilization of macrophages in the anterior chamber of mice"

Appendix 15
Mouse ID: 35787

Masked assignment: **Aged**
Actual group: **Aged**
OD anterior chamber injection

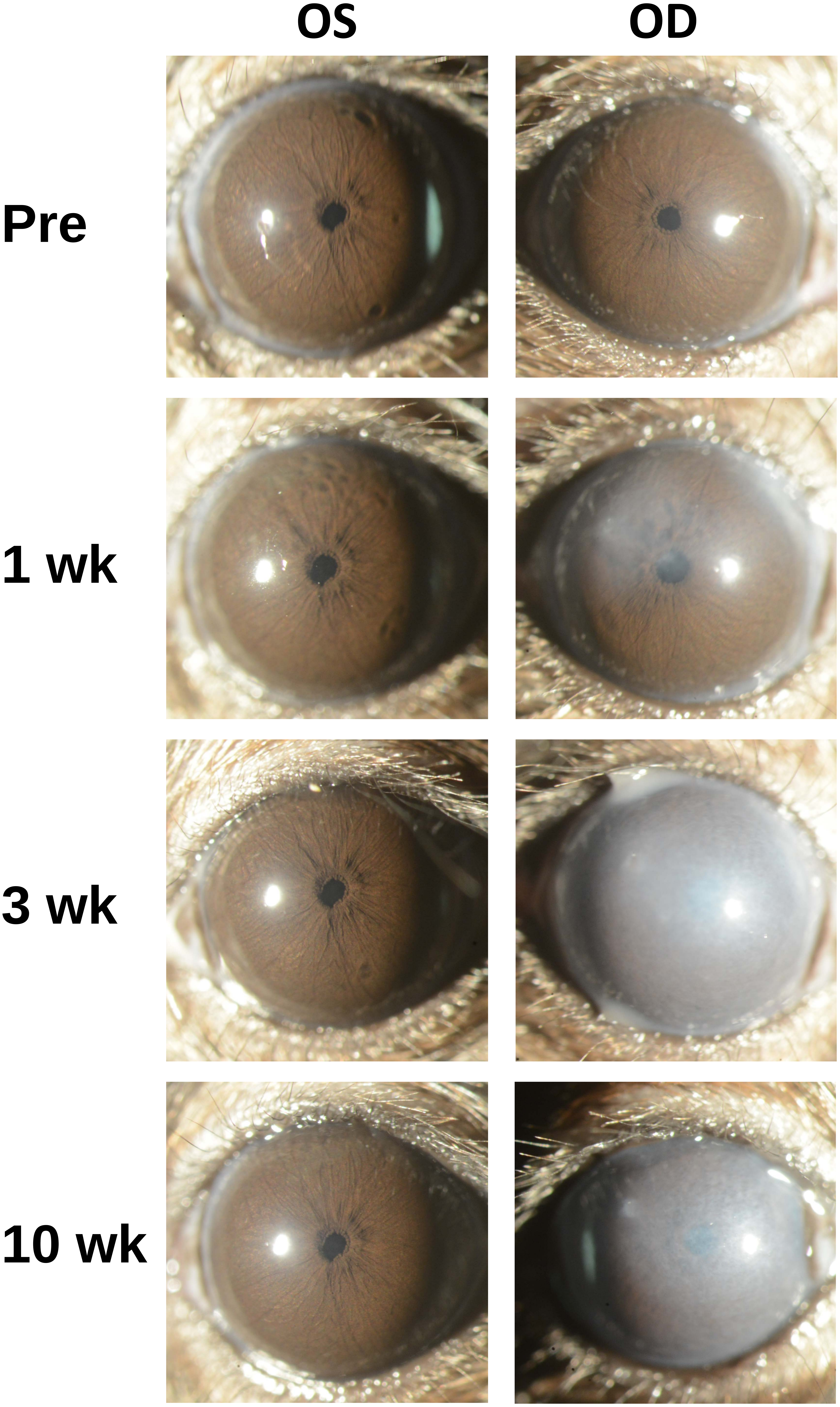

Appendix 15
Mouse ID: 35788

Masked assignment: **Aged**
Actual group: **Aged**
OD intravitreal injection

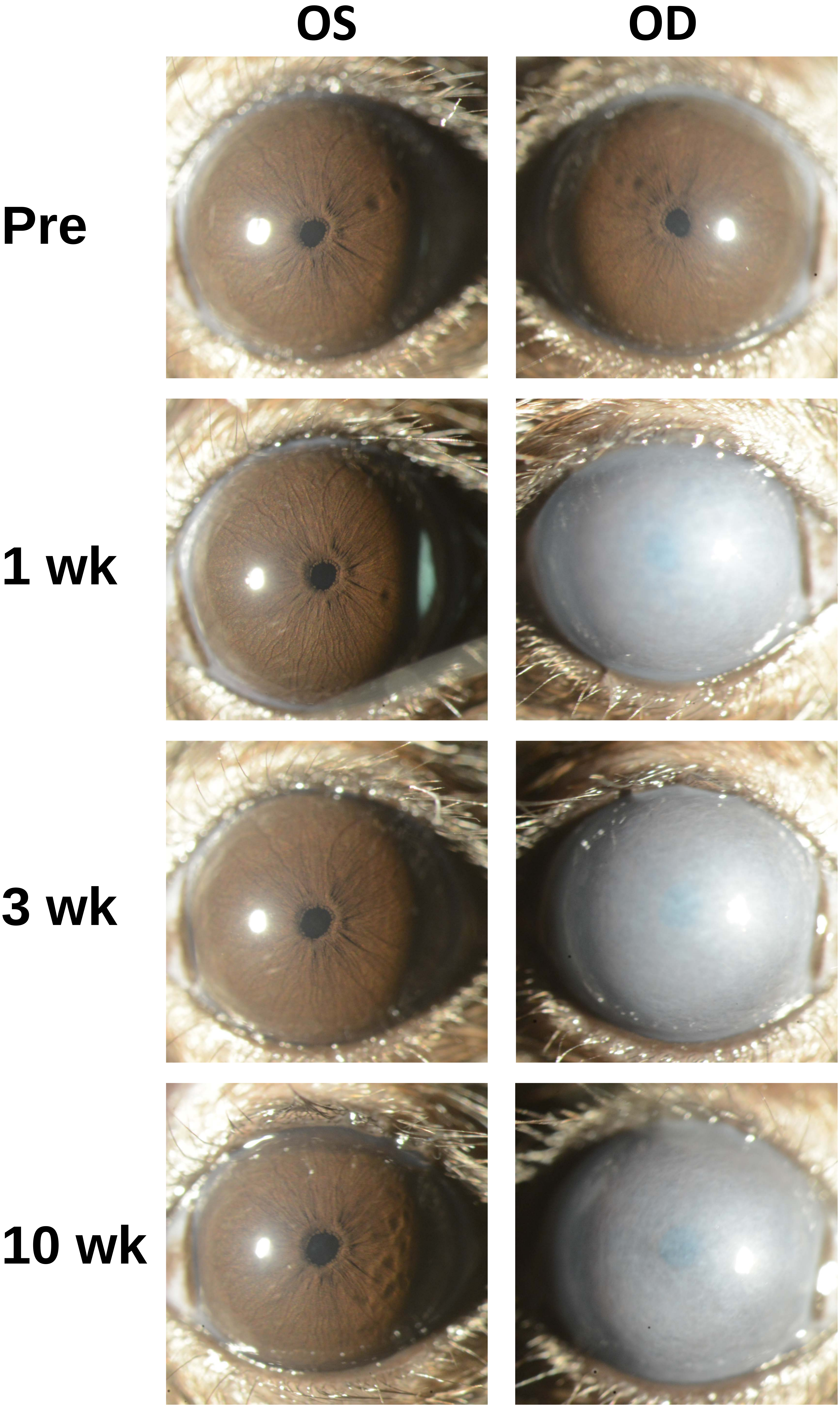

Appendix 15
Mouse ID: 35789

Masked assignment: **Aged**
Actual group: **Aged**
OD anterior chamber injection

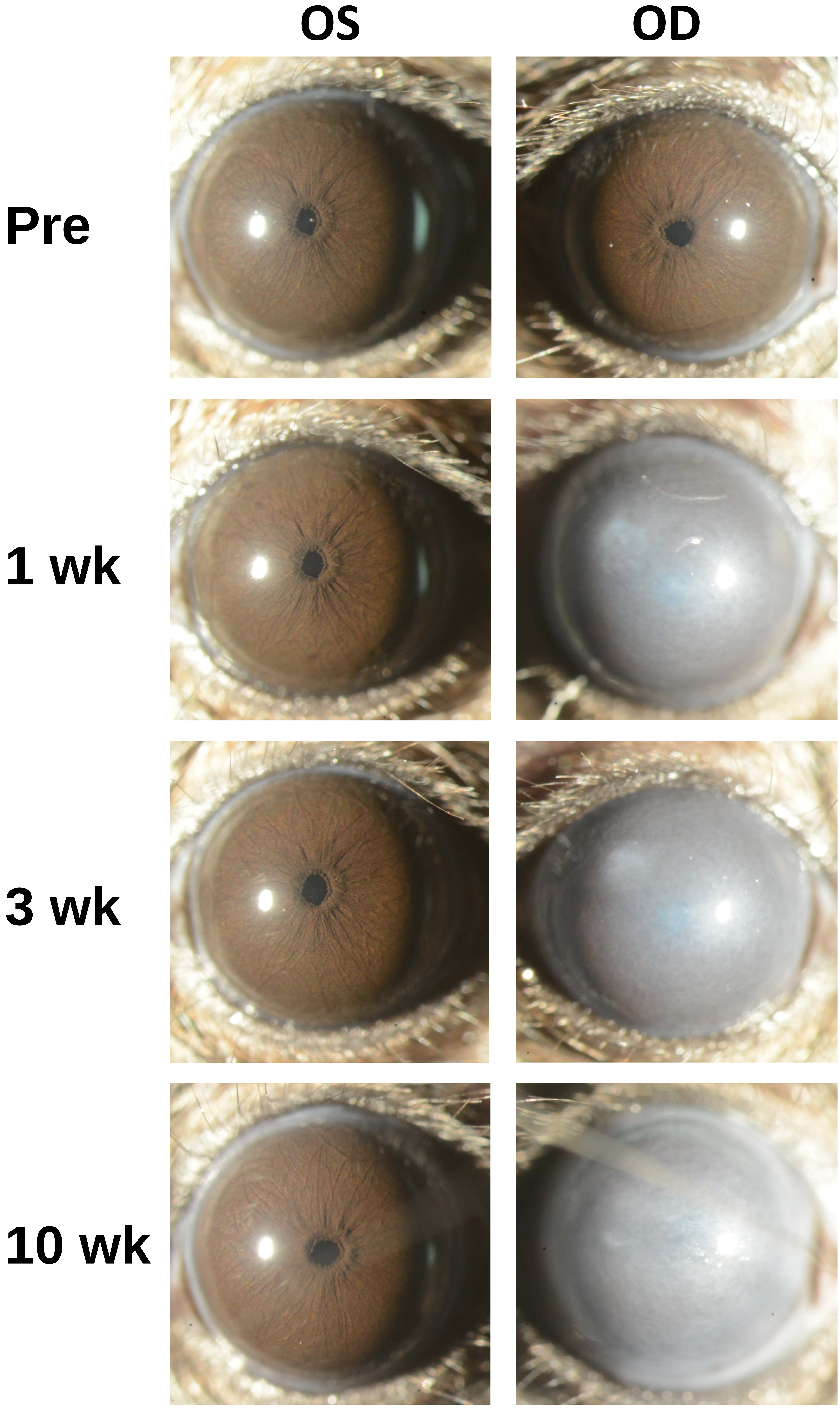

Appendix 15
Mouse ID: 35790

Masked assignment: **Aged**
Actual group: **Aged**
OD anterior chamber injection

**OS** **OD**

**Pre**

**1 wk**

**3 wk**

**10 wk**

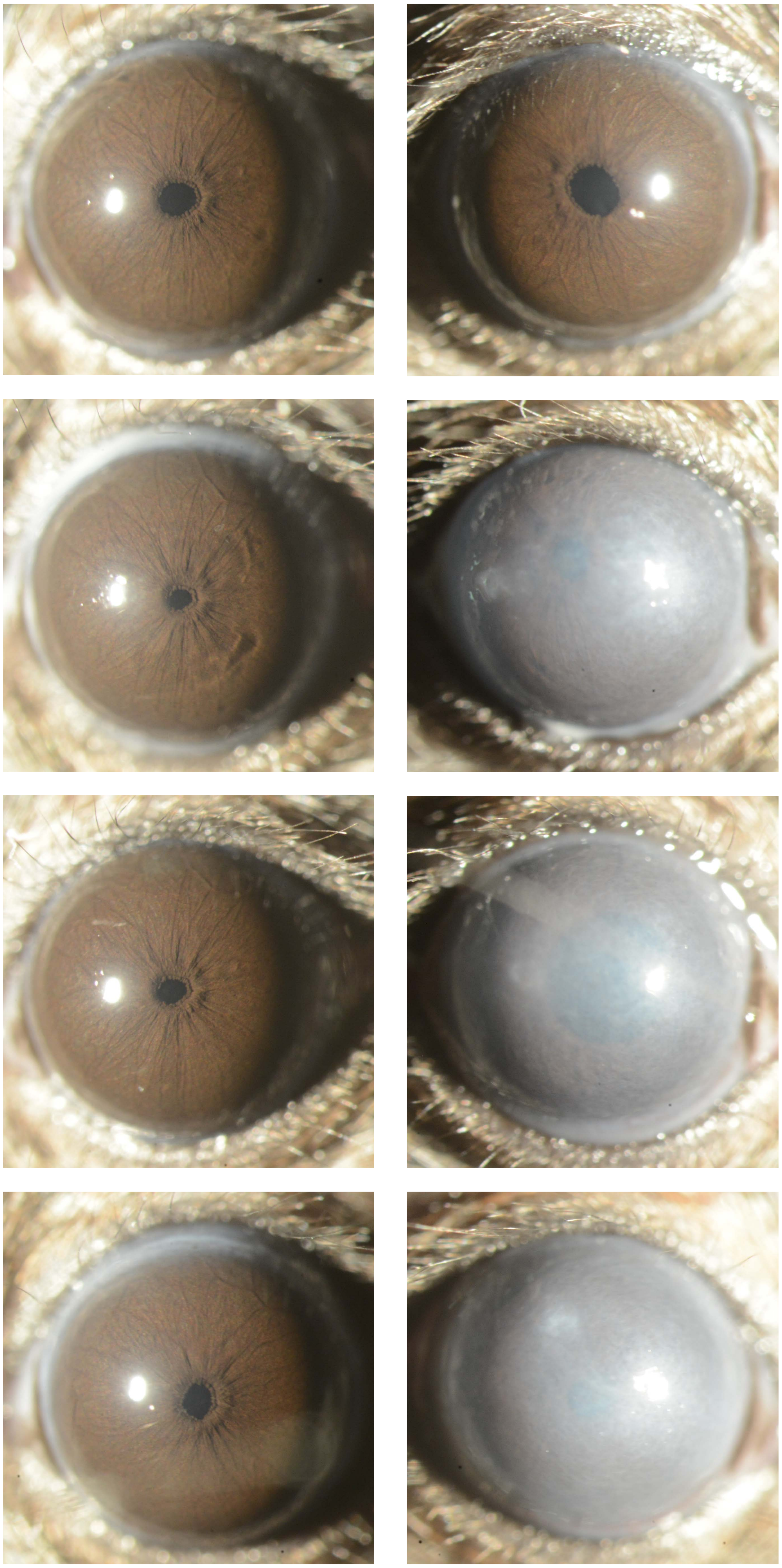
