## Appendix 16 Aged 40X for "Recombinant adenovirus causes prolonged mobilization of macrophages in the anterior chamber of mice"

Appendix 16
Mouse ID: 35787
10-week timepoint

Masked assignment: **Aged**
Actual group: **Aged**
(OD anterior chamber injection)

**OS**

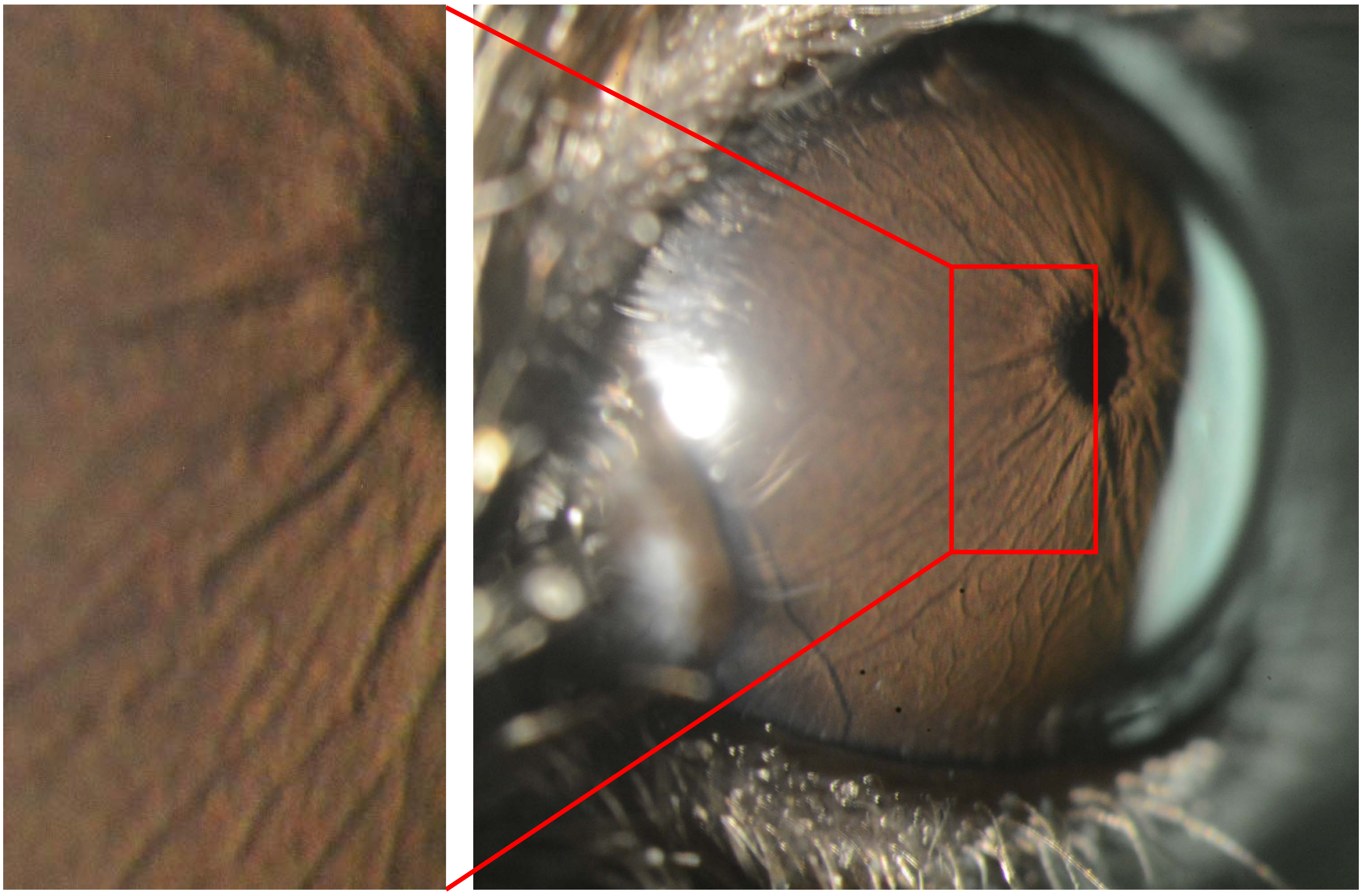

**OD**

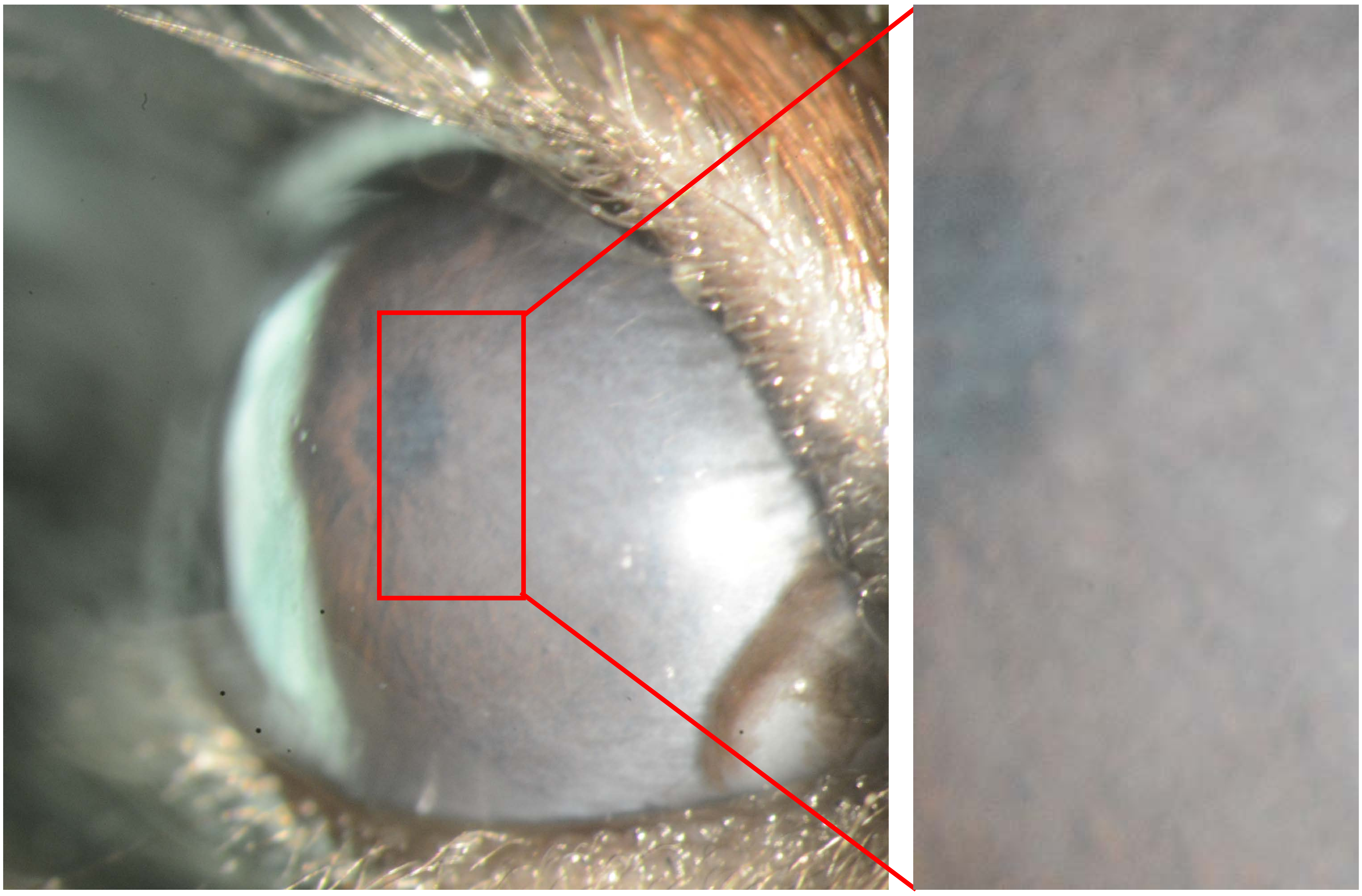

Appendix 16
Mouse ID: 35788
10-week timepoint

Masked assignment: **Aged**
Actual group: **Aged**
(OD intravitreal injection)

**OS**

**OD**

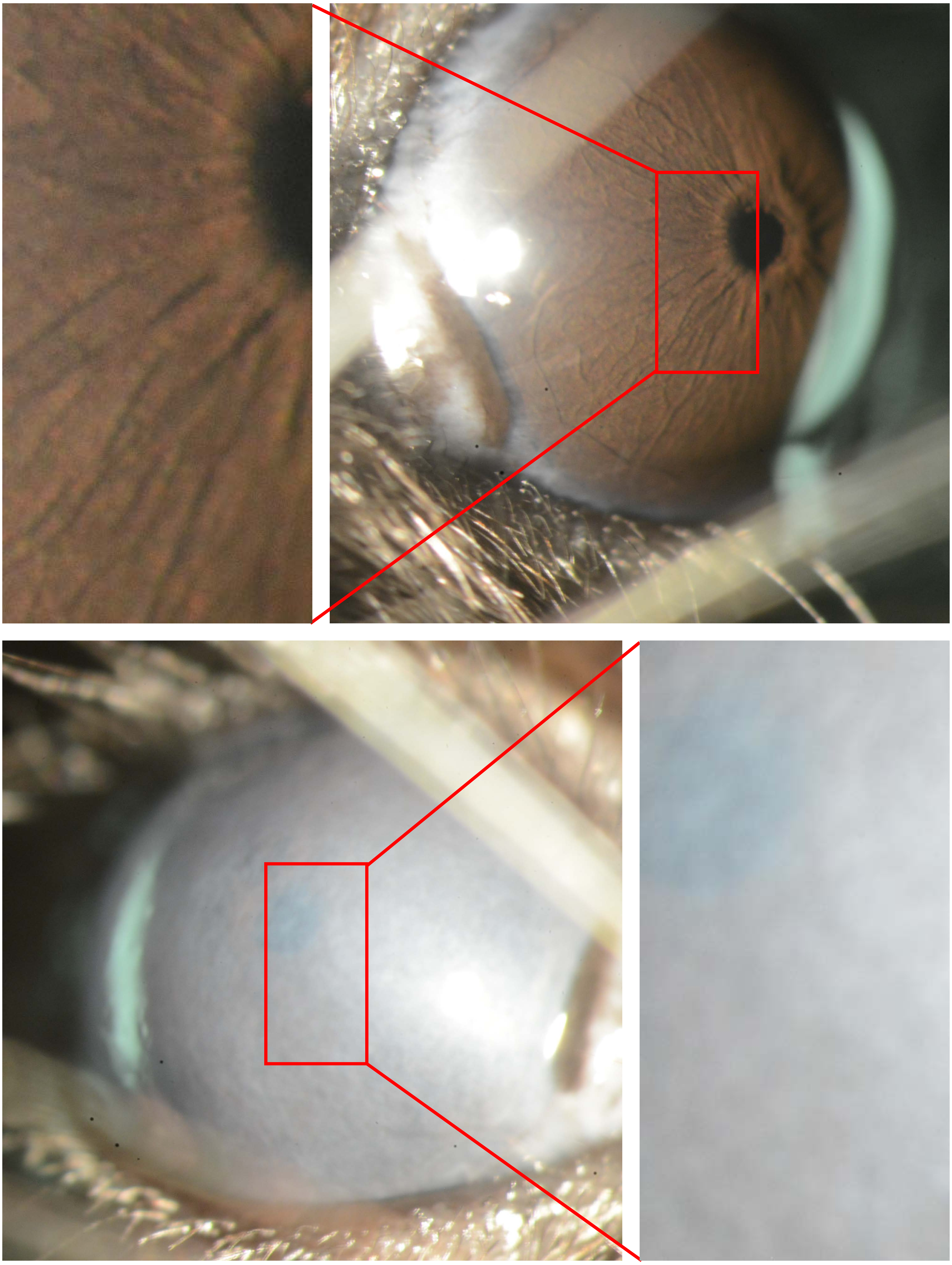

Appendix 16
Mouse ID: 35789
10-week timepoint

Masked assignment: **Aged**
Actual group: **Aged**
(OD anterior chamber injected)

**OS**

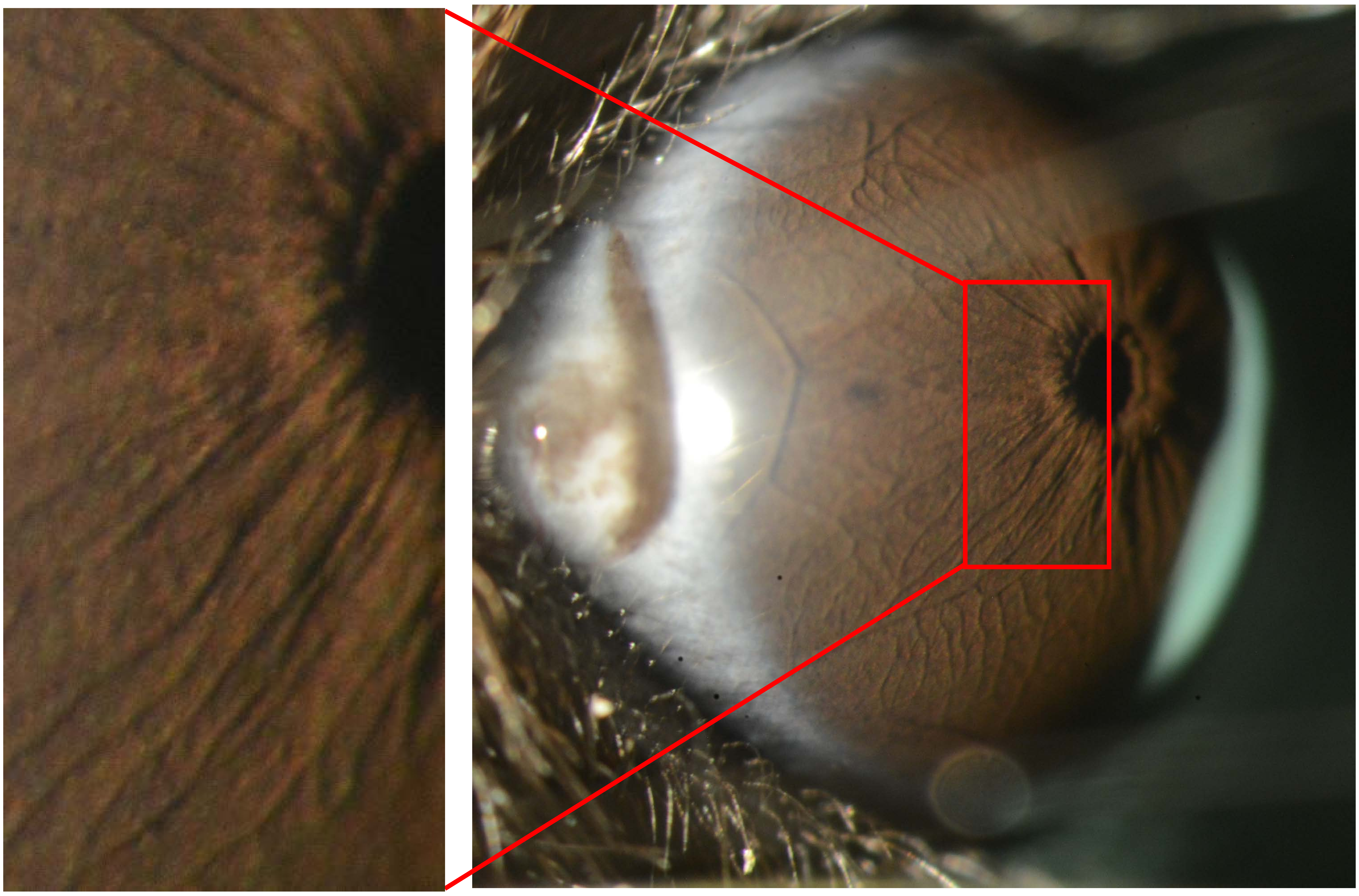

**OD**

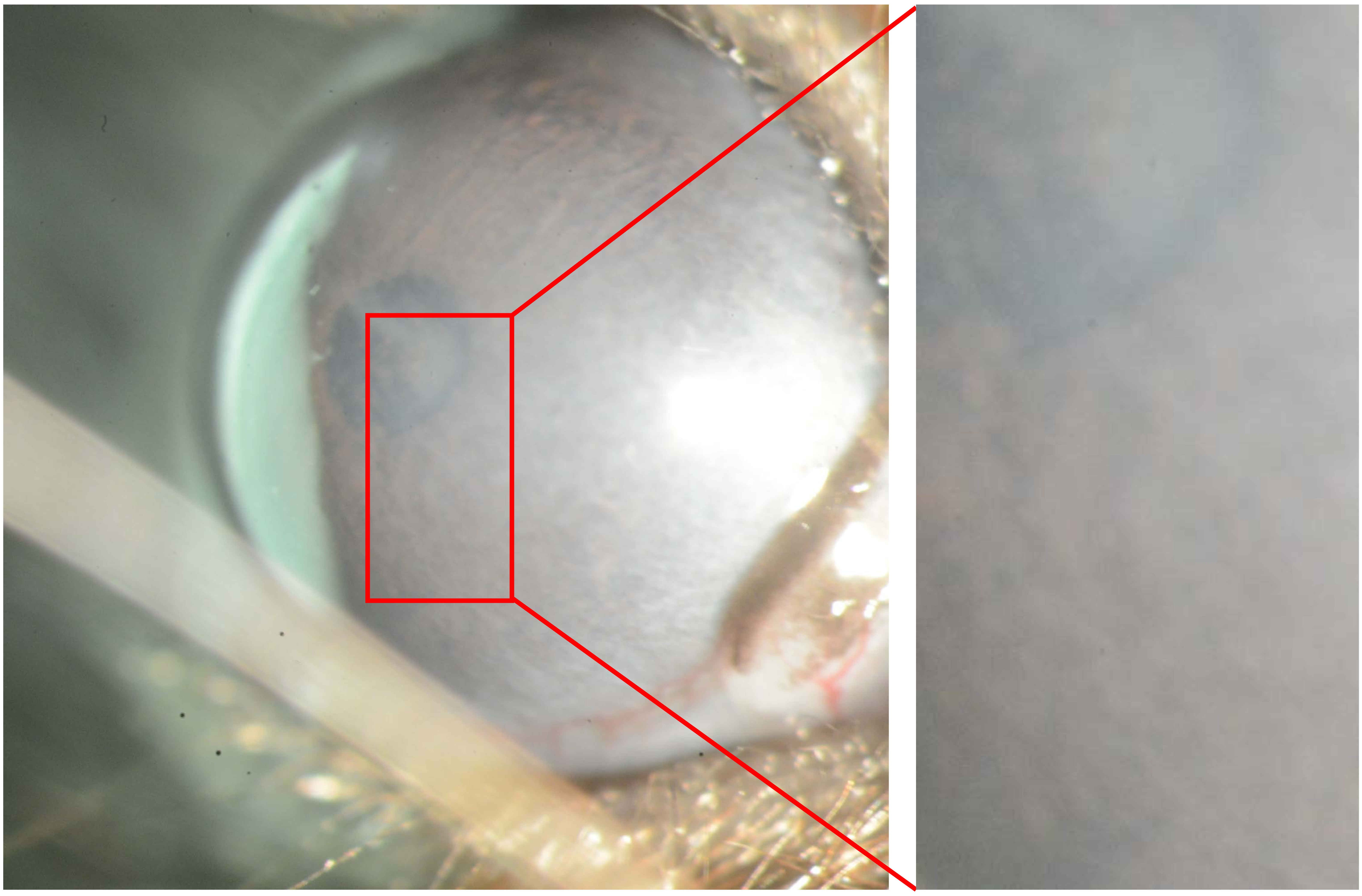

Appendix 16
Mouse ID: 35790
10-week timepoint

Masked assignment: **Aged**
Actual group: **Aged**
(OD anterior chamber injected)

**OS**

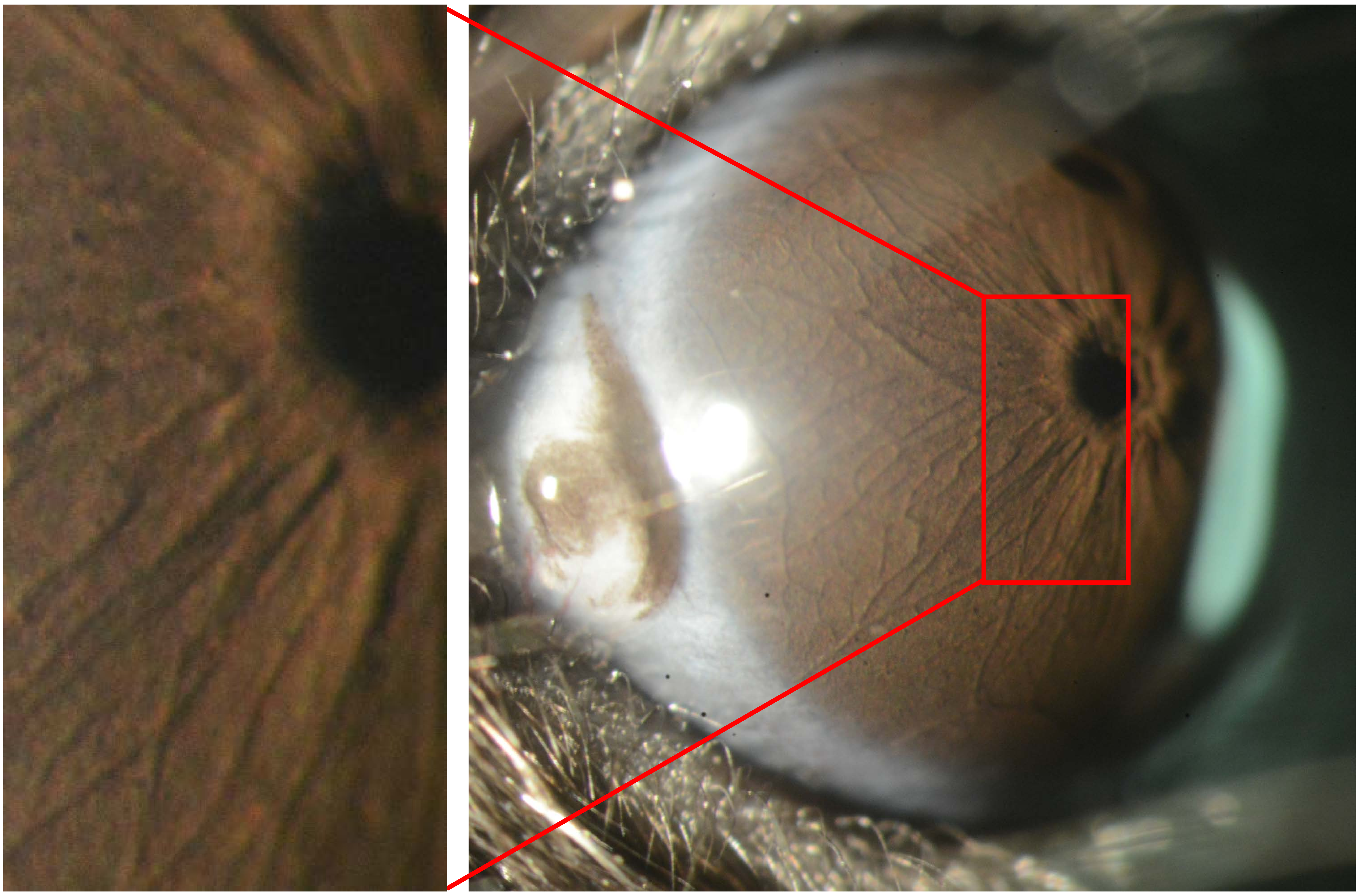

**OD**

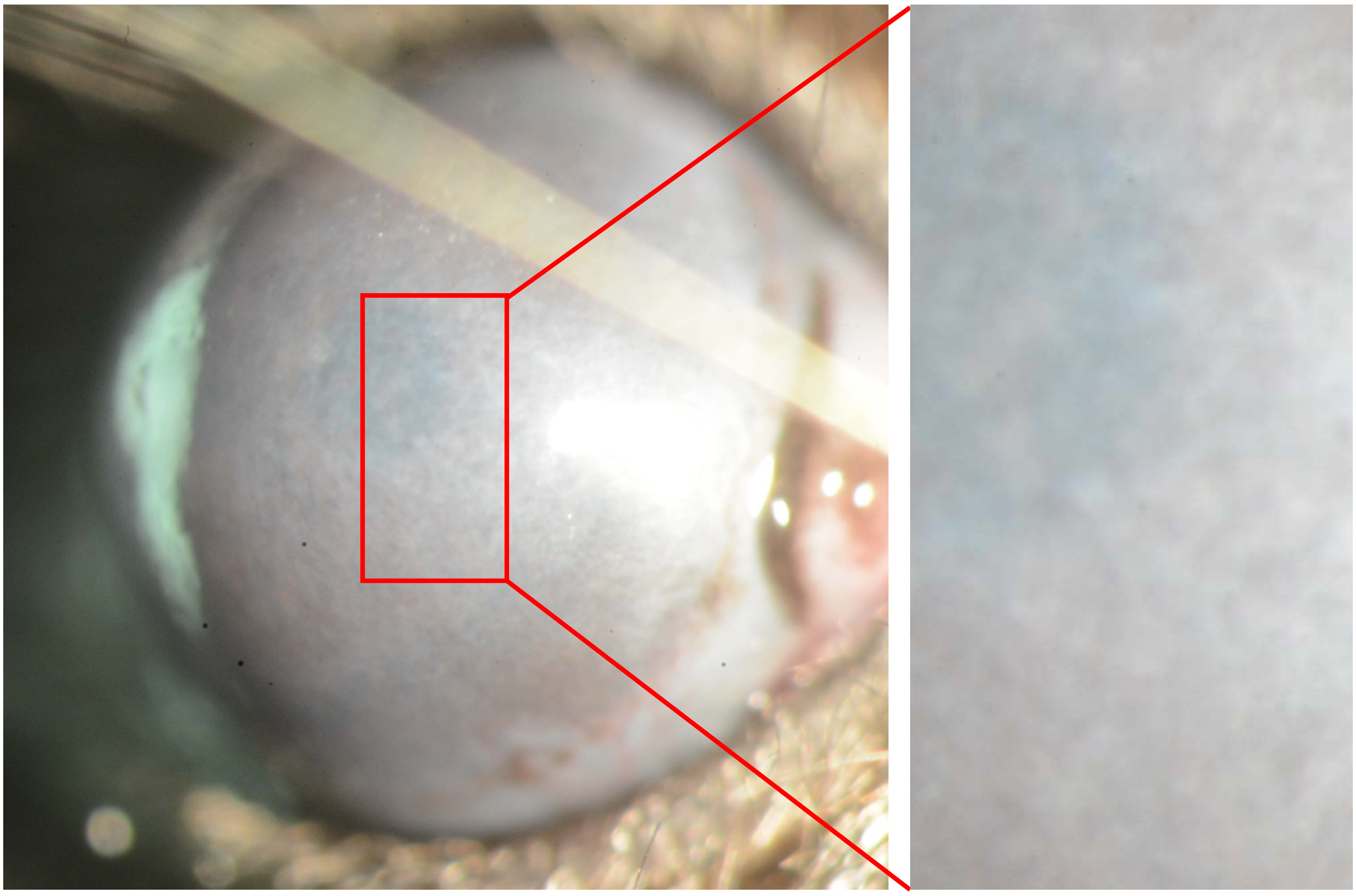
